# GTPBP8 is required for mitoribosomal biogenesis and mitochondrial translation

**DOI:** 10.1101/2023.05.26.542385

**Authors:** Liang Wang, Taru Hilander, Xiaonan Liu, Hoiying Tsang, Ove Eriksson, Christopher B. Jackson, Markku Varjosalo, Hongxia Zhao

## Abstract

Mitochondria contain a multi-copy genome and a distinct set of ribosomes devoted to the exclusive synthesis of proteins that are essential for oxidative phosphorylation. The assembly of mitoribosomes for mitochondrial translation is a poorly understood process. In this study, we identify the uncharacterized GTP-binding protein 8 (GTPBP8) as a mitoribosomal assembly factor that specifically associates with the mitoribosomal large subunit. Genetic depletion of GTPBP8 causes an aberrant accumulation of the large mitoribosomal subunit at a late assembly stage and reduces the level of fully assembled 55S mitoribosomes, resulting in impaired mitochondrial translation and function. Together, our findings uncover an important role for human GTPBP8 in the processes of mitoribosomal assembly and mitochondrial translation.

**Key points:**

1. GTPBP8 is a novel GTPase residing in the matrix peripherally bound to the inner mitochondrial membrane
2. GTPBP8 specifically associates with the large mitoribosome subunit through interactions with large mitoribosomal proteins assembled at late stages.
3. GTPBP is essential for maturation of the mitoribosomal large subunit and monosome formation

## Introduction

Mitochondria have a genome of their own coding for 13 proteins that are essential components of the oxidative phosphorylation (OXPHOS) complexes. In mammals, these largely hydrophobic proteins are synthesized by specialized mitochondrial ribosomes (mitoribosomes) located in the matrix and associated with the inner mitochondrial membrane to facilitate co-translational insertion of newly synthesized proteins (1). At the molecular level, the mechanism of mitochondrial protein synthesis is homologous to that of protein synthesis in bacteria. As a result, antibiotics disrupting protein synthesis on bacterial ribosomes also have similar effects on mammalian mitoribosomes (2–4). However, mitoribosomes are distinct from their prokaryotic ancestors in several ways (5–7). The bacterial ribosome, a 70S particle composed of a 30S small ribosomal subunit and a 50S large ribosomal subunit, possesses a very compact structure with approximately 70% ribosomal RNA (rRNA) and 30% proteins (7). In contrast, the mammalian mitoribosome has a sedimentation coefficient of 55S with an inverse protein to RNA ratio, having acquired both additional proteins and N- and C-terminal extensions to pre-existing ribosomal proteins during evolution (1, 5, 6). The 55S mitoribosome consists of two subunits, the large subunit (mt-LSU, 39S) and the small subunit (mt-SSU, 28S) which contain 16S and 12S mitoribosomal RNAs (mt-rRNA), respectively, and more than 80 mitoribosomal proteins (MRPs) (3). Recent advances in cryo-electron microscopy have solved the high-resolution structures of mammalian mitoribosomes (3, 5, 6, 8–14). The biogenesis of each subunit proceeds through an intricate and hierarchical process and forms the mature monosome during the initiation phase of mitochondrial translation. The mt-rRNAs are encoded by the mitochondrial DNA (mtDNA), while all the MRPs are encoded in the nucleus and imported into the mitochondrion following synthesis in the cytosol. Therefore, mitoribosome assembly is a highly coordinated process that involves the import, processing, and assembly of nuclear-encoded MRPs as well as post transcriptional processing of the mt-rRNAs (1, 15–18).

Robust models for bacterial ribosome assembly have been established earlier (19–21). However, this information is not readily transferable to eukaryotic mitoribosomes because of the substantial structural and compositional alterations that have occurred during evolution. The assembly of the mitoribosome is assisted by many assembly factors transiently associating with the nascent ribosome to enable its accurate and efficient construction (15, 22). These assembly factors include rRNA processing and modifying enzymes, RNA helicases, chaperones, and GTPases (15, 21, 23, 24). In bacteria, GTPases represent the largest class of essential ribosome assembly factors (23, 25). The GTPases involved in the ribosome assembly have mostly been studied in bacteria and yeast. The available information is scarcer regarding their roles in the assembly of the mammalian mitochondrial ribosomes. Only a few GTP binding proteins (GTPBPs) of mitoribosomes have been identified and characterized thus far for the assembly of mt-LSU and mt-SSU. Two human GTPases MTG3 (C4orf14, NOA1, YqeH) and ERAL1 (bacterial Era) are known to be important for the assembly of the mt-SSU (26, 27). GTPBP5 (also known as MTG2/OBGH1, bacterial ObgE), GTPBP6 (also known as bacterial HflX), GTPBP7 (also known as MTG1, bacterial RbgA) and GTPBP10 (also known as OBGH2) (22, 26–30) facilitate the assembly of mt-LSU at different time points (18). Both GTPBP5 and GTPBP10 associate with the mt-LSU but they display distinct and not mutually compensable functions in the mt-LSU assembly (26–28, 30). GTPBP10 mostly associates with the mtLSU in a GTP-dependent manner (26, 30). GTPBP5 facilitates the methylation of 16S rRNA securing coordinate subunit joining and release of late-stage mtLSU assembly factors (27, 28). GTPBP7 binds the 16S rRNA of mt-LSU to promote the incorporation of late-assembling MRPs and couples the mt-LSU assembly with the formation of inter-subunit bridges (28, 29). GTPBP6 acts during the last steps of the mt-LSU assembly and assists in the dissociation of the ribosomal subunits for recycling (22).

The GTP binding protein 8 (GTPBP8), a previously uncharacterized protein, was recently found in the DDX28 (an assembly factor of mt-LSU) interactome from mitochondrial RNA granules (28). The bacterial orthologue of GTPBP8, Ysxc/Yiha, sharing ~20% sequence identity with GTPBP8, is associated with the bacterial ribosomal large subunit and required for ribosome biogenesis (29–31). Yet little is known regarding its mitochondrial GTPBP8 counterpart, particularly in human cells. In this study, we characterize the cellular function of human GTBBP8 in mitochondria. Our results reveal that GTPBP8 is imported into mitochondria by an N-terminal pre-sequence and resides in the mitochondrial matrix, peripherally bound to the inner membrane. GTPBP8 specifically associates with the 39S mt-LSU and plays an essential role in the maturation of the 39S mt-LSU subunit at a late stage. Loss of GTPBP8 leads to an abnormal accumulation of mt-LSU and a marked reduction of fully assembled 55S monosomes, resulting in impaired mitochondrial translation and respiration. Taken together, our findings uncover human GTPBP8 to be an essential assembly factor of the mitoribosome and critical for mitochondrial translation.

## Results

### GTPBP8 is a GTPase localized to mitochondria

Primary sequence analysis of the GTPBP8 in eukaryotes, *E.coli* (YihA), and *B.subtilis* (YsxC) reveals that GTPBP8 has five evolutionary conserved GTPase domains (G1-G5; Figure S1A, B). To examine whether human GTPBP8 possesses GTPase activity, we purified His-tagged GTPBP8 and assayed its GTPase activity in vitro. The result confirmed GTPBP8 to exhibit a low intrinsic GTPase activity, with a K*_M_* and K*_cat_* of 0.20±0.04 mM and 0.0079 ± 0.0033 min^-1^ (n=3), respectively (Figure S1C), which are comparable to those of other mitoribosomal GTPases (44, 45). This demonstrates that human GTPBP8 is an evolutionarily conserved guanosine triphosphate (GTP) binding protein.

We further examined the intracellular localization of GTPBP8 by immunocytochemistry using an anti-GTPBP8-antibody. The endogenous GTPBP8 co-localized with the mitochondrial protein TOM20 (Figure 1A), suggesting that GTPBP8 resides within the mitochondria. Expression of myc- or green fluorescent protein (GFP)-tagged GTPBP8 (Figures 1A, 1B, S2A, and S2B) in human osteosarcoma cells (U2OS) displayed a similar intracellular distribution, co-localizing with the mitochondrial proteins TOM20 and COXIV (Figure 1A). Furthermore, subcellular fractionation and subsequent immunoblotting of both endogenous GTPBP8 (Figure 1C) and GTPBP8-myc (Figure S2C) were exclusively detected in the mitochondrial fraction, as indicated by the mitochondrial outer membrane (MOM) protein TOM40, the mitochondrial intermembrane space (IMS) protein cytochrome C, and the matrix-localized mitochondrial ribosomal protein uL11m. GTPBP8 was not detected in the cytosolic fraction, revealing that GTPBP8 localized exclusively to mitochondria.

**Figure 1.**
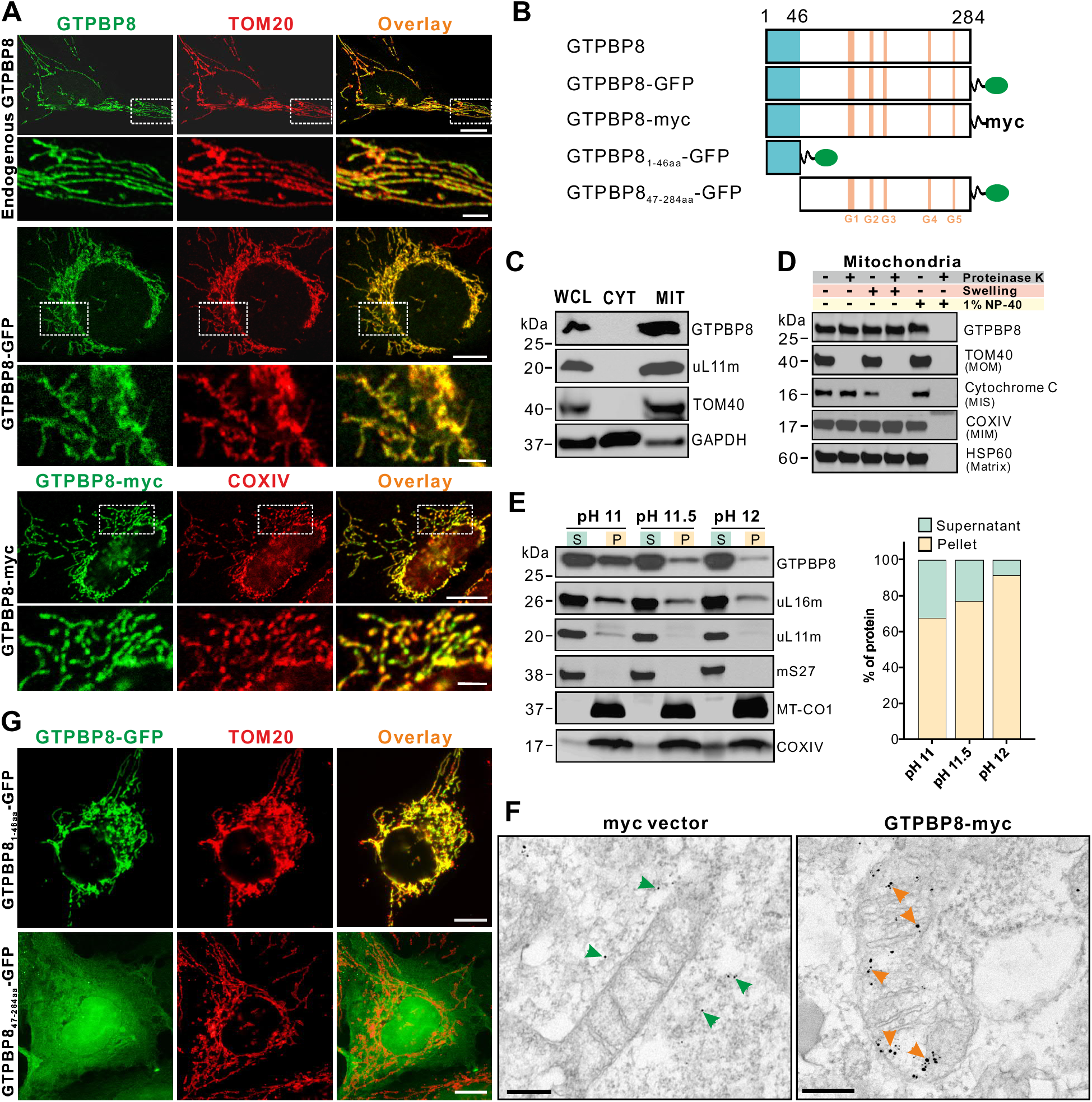
GTPBP8 resides in the mitochondrial matrix peripherally associated with the inner mitochondrial membrane. (**A**) Immunofluorescence microscopy analysis of cellular localization of endogenous GTPBP8, GTPBP-GFP, and GTPBP8-myc. Scale bars: 10 µm. Scale bars in insets: 2.5 µm. (**B**) Schematic diagram of GTPBP8 constructs. (**C**) Immunoblotting analysis of the subcellular localization of GTPBP8. U2OS cells were fractionated, and the endogenous GTPBP8 was detected in the whole-cell lysate (WCL), cytoplasm (CYT) and isolated mitochondria (MIT). Antibodies against the mitochondrial proteins TOM40 and uL11m, and a cytosolic protein GAPDH were used as controls. (**D**) Immunoblotting analysis of the submitochondrial localization of endogenous GTPBP8 using proteinase K digestion of isolated mitochondria from U2OS cells. (**E**) Mitochondrial protein extraction by sodium carbonate using isolated mitochondria from U2OS cells. Quantification of GTPBP8 protein abundance in the supernatant and pellet was shown on the right panel. Mitoribosomal proteins uL16m, uL11m, and mS27 and transmembrane mitochondrial proteins MT-CO1 and COXIV were used as controls. (**F**) Immuno-electron microscopic analysis of GTPBP8-myc localization in U2OS cells. The nano-gold particles in the cytoplasm and mitochondria were indicated by green and orange arrows, respectively. Scale bars: 200 nm. (**G**) Representative immunofluorescence images of U2OS cells transfected with GTPBP8_1-46aa_-GFP and GTPBP8_47-284aa_-GFP for 24 h. The transfected cells were immunostained with an anti-TOM20 antibody. Scale bars: 10 µm.

Mitochondria are enclosed by two membrane defining distinct functional and morphological subdomains. To determine the sub-mitochondrial localization of GTPBP8, we purified mitochondria and treated them with proteinase K in order to degrade proteins exposed on the external surface of the outer membrane. Immunoblotting revealed that both endogenous GTPBP8 (Figure 1D) and GTPBP8-myc (Figure S2D) were not degraded by proteinase K but instead became accessible to enzyme digestion when the mitochondrial membranes were solubilized by 1% NP-40, similar to the integral mitochondrial inner membrane (MIM) protein COXIV and the mitochondrial matrix protein HSP60. In contrast, TOM40 localized on the MOM was completely digested in the absence of detergent, whereas the IMS protein cytochrome C was degraded only when the MOM was ruptured by hypo-osmotic swelling, indicating that the differences in their accessibility to proteinase K were due to their differing sub-mitochondrial localization. These data suggest that GTPBP8 resides either on the inner surface of MIM or in the mitochondrial matrix. Thus, to determine whether GTPBP8 is a mitochondrial soluble, peripheral, or integral membrane protein, mitochondria were subjected to alkaline carbonate treatment at different pH values. Following treatment at pH 11, GTPBP8 resided mainly in the supernatant and partially in the pellet/membrane fraction (Figure 1E). However, at a pH of 12, GTPBP8 was almost completely released to the soluble fraction, suggesting that GTPBP8 is a mitochondrial matrix protein, with a fraction of GTPBP8 peripherally interacting with the inner mitochondrial membrane similar to other mitochondrial GTPases, including GTPBP5, GTPBP6, or GTPBP10 (22, 46, 47). To corroborate these findings, we performed immuno-electron microscopy of GTPBP8-myc-expressing cells using an anti-myc antibody. The electron micrographs showed that GTPBP8-myc resided in the mitochondrial matrix in close proximity to the inner membrane (Figure 1F), while in the control myc-expressing cells the colloidal gold particles were localized in the cytoplasm. Taken together, these results demonstrate that GTPBP8 is a mitochondrial protein in the matrix that is peripherally associated with the inner mitochondrial membrane.

GTPBP8 is a nuclear DNA (nDNA)-encoded protein and is imported into the mitochondria after synthesis. In addition to the highly conserved GTPase domains, human GTPBP8 possesses a specific N-terminal extension of 84 amino acids predicted to contain a putative mitochondrial import sequence (Figure S1B). Analysis for mitochondrial import probability using the Mitominer algorithm revealed a putative mitochondrial import sequence at the N-terminus of human GTPBP8 (the IMPI and Target P scores were 1 and 0.815, respectively) (http://mitominer.mrc-mbu.cam.ac.uk/release-4.0/report.do?id=1021783&trail=%7c1021783). Thus, we tagged the N-terminal 1-46 amino acids containing positively charged clusters or a mutant lacking the N-terminal 1-46 amino acids to GFP (GTPBP8_1-46aa_-GFP, Figures 1B and S1B). Transient expression of GTPBP8_1-46aa_-GFP confirmed unambiguous mitochondrial localization (Figure 1G). On the other hand, deletion of the N-terminal 1-46aa (GTPBP8_47-284aa_-GFP) resulted in a diffuse cellular localization signal (Figure 1G). These data demonstrate that the N-terminal 46aa extension is essential for mitochondrial import of human GTPBP8.

### Depletion of GTPBP8 impaired mitochondrial bioenergetics

To study the cellular role of GTPBP8, we performed loss-of-function experiments using siRNA silencing. All four tested siRNAs significantly depleted the mRNA level of GTPBP8 (Figure 2A) and showed both abrogated GTPBP8 protein levels in whole-cell and isolated mitochondrial lysates (Figure 2B). Loss of GTPBP8 decreased cell proliferation significantly (Figure 2C). To reveal the underlying mechanism by which GTPBP8 depletion affected cell proliferation, mass spectrometry (MS)-based proteomics analysis was performed. Reactome network analysis of all identified mitochondrial proteins revealed several notably changed biological processes and molecular pathways in mitochondria after depletion of GTPBP8, with substantial clustering in mitochondrial translation and electron transport/oxidative phosphorylation (Figures 2D and S3A). Further analysis with mitochondrial proteins having significant changes after depletion of GTPBP8 showed that mitochondrial translation is the most affected process among the enriched biological processes (Figure 2E). Additionally, most of the involved mitochondrial proteins displayed reduced abundance following GTPBP8 depletion (Figure 2E, right panel), suggesting that loss of GTPBP8 could have a profound effect on mitochondrial function. Thus, we measured bioenergetics using high-resolution respirometry. The results showed that downregulation of GTPBP8 resulted in a clearly diminished oxygen consumption rate and respiratory capacity, significantly lower basal and maximum respiration, as well as ATP-linked respiration (Figures 2F and 2G). In agreement with these findings, the level of total cellular ATP was reduced (Figure 2H). Collectively, these data demonstrated that silencing of GTPBP8 caused mitochondrial respiration defects and reduced cellular ATP production, resulting in decreased cell proliferation.

**Figure 2.**
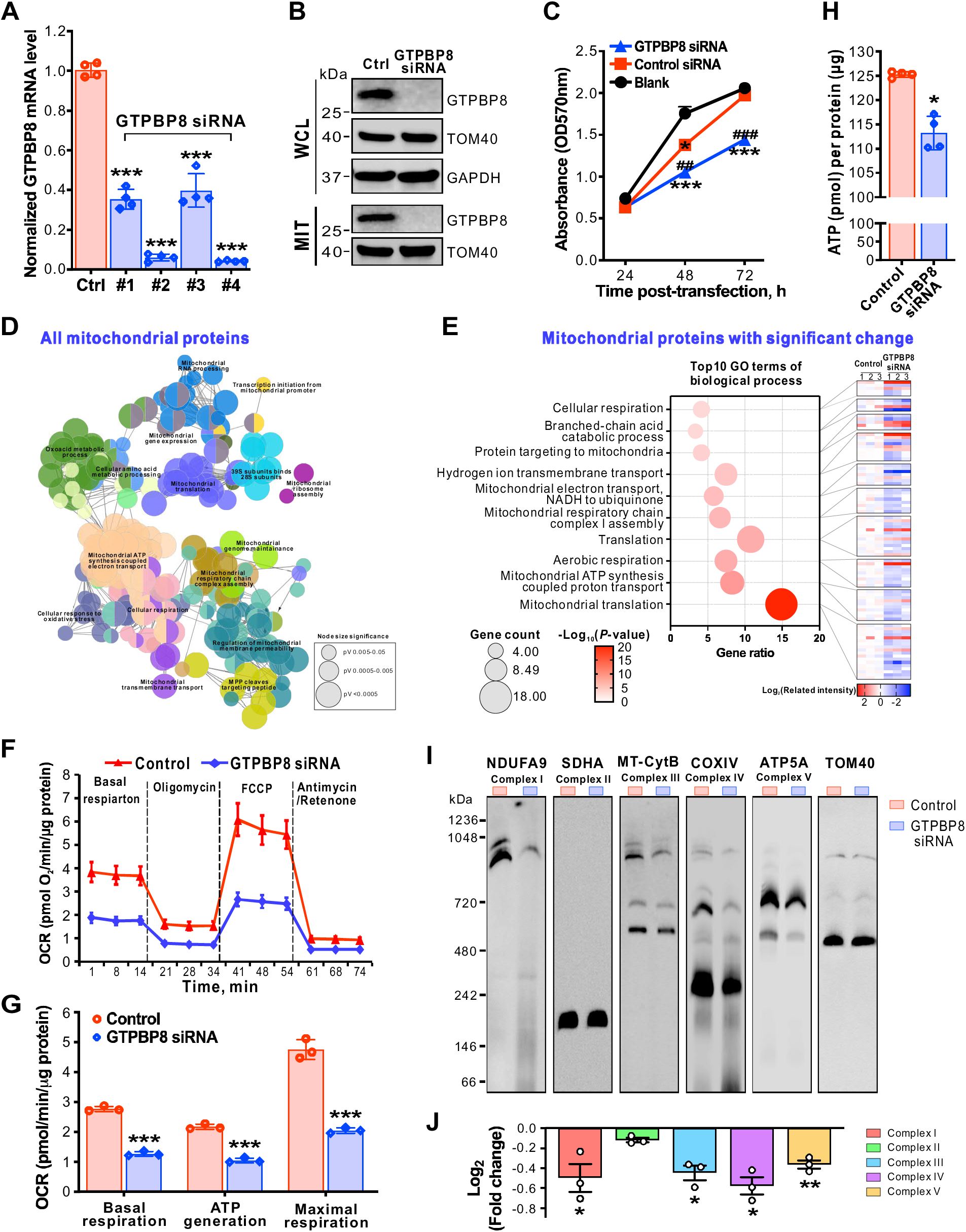
Depletion of GTPBP8 impairs mitochondrial respiration and the assembly of OXPHOS complexes. (**A**) Steady-state mRNA levels of GTPBP8 in U2OS cells after treatment with control or GTPBP8 siRNA detected by RT-qPCR. Error bars represent the mean ± SD of four independent experiments. *** P≤0.001, One-way Anova. (**B**) Immunoblotting analysis of endogenous GTPBP8 protein levels in the whole-cell lysate (WCL) and isolated mitochondria (MIT). TOM40 and GAPDH were used as loading controls. (**C**) Determination of cell proliferation/cell viability by a MTT assay at the indicated times after treatment with control or GTPBP8 siRNA for 5 days. Blank does not have any siRNA treatments. * P≤0.05 and *** P≤0.001 are GTPBP8 siRNA treatment results compared to the blank. ## P≤0.01 and ### P≤0.001 are GTPBP8 siRNA treatment results compared to control siRNA, Two-way ANOVA. (**D**) Grouping of network based on functionally enriched GO terms and pathways after depletion of GTPBP8. The functionally grouped network of enriched categories of all identified mitochondrial proteins was generated using ClueGO/CluePedia. All significant enriched GO terms are represented as nodes based on their kappa score level 0.4. Only the networks and pathways with p<0.05 are shown. Functionally grouped networks are linked to their biological function, where only the representative term for each group is labeled. The node size presents the significance level of the enriched GO term. Functionally related groups partially overlap. Visualization was carried out using Cytoscape 2.8.2. (**E**) Bubble plot of top10 GO terms of significantly changed mitochondrial proteins after depletion of GTPBP8. The heatmap in the right panel presents the related intensity of individual proteins in the corresponding GO term. (**F**) Oxygen consumption rate (OCR), measured using a Seahorse XF96^e^ Analyzer after depletion of GTPBP8. The OCR was determined by adding mitochondrial function inhibitors sequentially at the indicated times. The OCR was normalized to the total protein amount used for the assay. (**G**) Quantification of the basal, ATP linked and maximal respiration capacities from the OCR measurement in panel F. (**H**) Quantification of cellular ATP level after GTPBP8 depletion. (**I**) Effects of GTPBP8 depletion on the OXPHOS complex assembly assessed by BN-PAGE. (**J**) Quantification of OXPHOS complexes in Panel I. Normalization was done against TOM40 complex. In panels B-J, U2OS cells were treated with control or GTPBP8 siRNA for 5 days. Error bars represent the mean ± SD of three independent experiments. * P≤ 0.05, ** P≤ 0.01 and *** P≤0.001, Student’s t-test.

### GTPBP8 is important for mitochondrial translation

Mitochondrial bioenergetics relies on the oxidative phosphorylation (OXPHOS) complexes, which are multimeric and, except complex II, composed of the subunits encoded by both mitochondrial DNA (mtDNA) and nDNA. To examine whether the bioenergetic defect caused by the depletion of GTPBP8 was a consequence of perturbed OXPHOS complexes, we purified mitochondria to separate protein complexes by native-PAGE followed by immunoblotting. Indeed, depletion of GTPBP8 reduced the levels of OXPHOS complexes, in particular complexes I, III, IV, and V, whereas the abundance of the exclusively nuclear-encoded complex II was not affected (Figures 2I, 2J, and S3B). To determine whether the proteins encoded by mtDNA or nDNA contributed to the reduced complexes I, III, IV, and V, the steady-state levels of proteins in the OXPHOS complexes were measured by immunoblotting. Strikingly, only the mtDNA-encoded protein levels of MT-CO1 and MT-CytB were reduced after GTPBP8 silencing (Figure 3A). In contrast, the protein abundance of a set of nDNA-encoded proteins, including NDUFA9, SDHA, COXIV, and ATP5A in complexes I, II, IV, and V, respectively, was not affected by GTPBP8 silencing (Figure 3A). Interestingly, the reduced protein levels of MT-CO1 and MT-CytB were neither caused by their transcription levels (Figure 3B) nor the mtDNA content (Figure 3C), suggesting a general mitochondrial translation deficiency.

**Figure 3.**
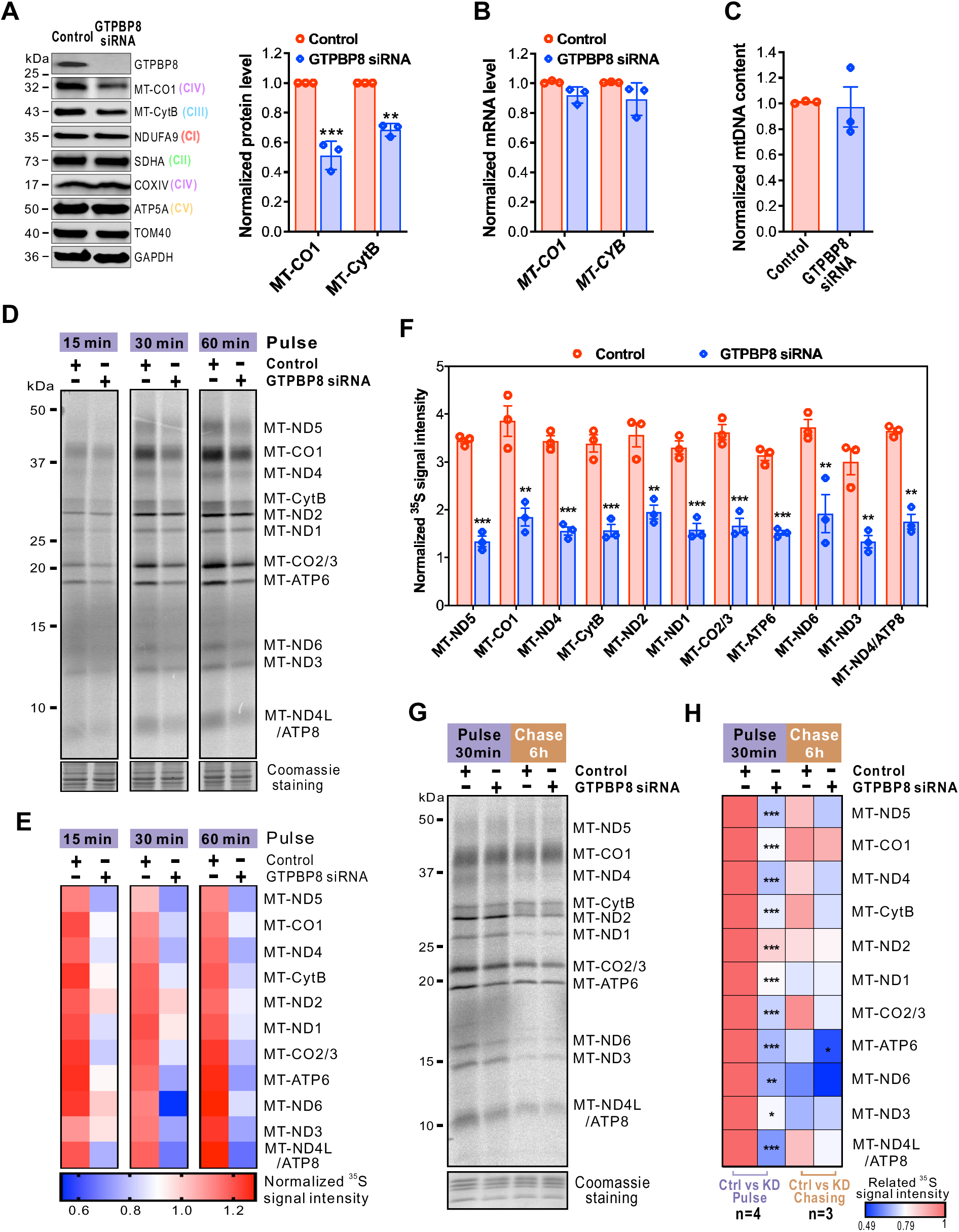
Loss of GTPBP8 impairs mitochondrial translation. (**A**) Steady state levels of oxidative phosphorylation complex proteins encoded by mitochondrial DNA (mtDNA) and nuclear DNA (nDNA) after depletion of GTPBP8. The right panel shows quantification of the relative MT-CO1 and MT-CytB protein abundance normalized to TOM40. (**B**) The mRNA expression levels of MT-CO1 and MT-CytB normalized to 18S rRNA. (**C**) Steady-state levels of mtDNA content detected by quantitative PCR. The total mtDNA levels were normalized to β-globin. (**D**) Metabolic labeling of mitochondrial translation products with [^35^S]-methionine/cysteine for the indicated time in U2OS cells. (**E**) The heatmap depicts the quantified ^35^S intensity in panel D for mitochondrially-translated proteins relative to the corresponding intensity in Coomassie staining. The color bar represents the normalized ^35^S intensity value. (**F**) Quantification of ^35^S intensity for mitochondrially-translated 13 proteins after depletion of GTPBP8. The values were normalized to the corresponding intensity in Coomassie staining. Error bars represent the mean ± SD of three independent experiments. ** P≤ 0.01 and *** P≤ 0.001, Student’s t-test. Metabolic labeling of mitochondrial translation products with [^35^S]-methionine/cysteine was performed for 30 min in U2OS cells after depletion of GTPBP8. The cells were treated with control and GTPBP8 siRNA for 5 days. (**G**) Metabolic labeling of mitochondrially-translated proteins with [^35^S]-methionine/cysteine for 30 min with or without 6 h of chasing in U2OS cells. In Panels A, B, and C, error bars represent the mean ± SD of three independent experiments. Student’s t-test. (**H**) The heatmap depicts the quantified ^35^S intensity in panel G for mitochondrially-translated proteins relative to the corresponding intensity in Coomassie staining. The color bar represents the normalized ^35^S intensity value.

To corroborate the importance of GTPBP8 in mitochondrial translation, we assessed the *de novo* protein translation rate in GTPBP8-depleted cells. Pulse-labelling with [^35^S] methionine/cysteine at different time points in the presence of the cytosolic protein synthesis inhibitor anisomycin revealed an overall reduction of newly synthesized mtDNA-encoded proteins in GTPBP8-depleted cells (Figures 3D-3F, S4A, and S4B). To further examine whether the stability of mitochondrial nascent proteins was impaired, pulse-chase labeling was applied. After 6 hours of chasing, there was no significant difference in protein turnover between control and GTPBP8 siRNA treated cells (Figures 3G and 3H), indicating that the nascent chains were stable. These data indicated that the reduction in mtDNA-encoded protein abundance in GTPBP8-silenced cells was caused by impaired mitochondrial protein synthesis, suggesting a critical role for GTPBPT8 in mitochondrial translation.

### GTPBP8 associates with the mitoribosomal large subunit (mt-LSU)

To assess how GTPBP8 functions in mitochondrial translation, we applied proximity labeling by fusing GTPBP8 to BioID (48) to identify the interaction partners of GTPBP8. The comprehensive interactome analysis revealed that 11 mitochondrial translation-related proteins, including 8 mt-LSU proteins, two mitochondrial aminoacyl-tRNA synthetases (YARS2 and IARS2), and a methyltransferase TRMT61B (an assembly factor of the mitroribosome), were pulled down with GTPBP8 (Figure 4A, Table S1), indicating that GTPBP8 may be associated with mitoribosomes. To further assess the link between GTPBP8 and mitoribosomes, the effects of GTPBP8 depletion on the abundance of mitoribosomal proteins were examined. Altogether, 74 out of 82 MRPs and 28 factors involved in mitoribosomal assembly were detected (Figure S5, Table S2). Clearly, the abundance of most of the MRPs was reduced after the depletion of GTPBP8 (Figure S5, Table S2). Consistently, western blot analysis revealed that depletion of GTPBP8 significantly decreased the steady-state levels of tested MRPs (Figures 4B, 4C, and S5C).

**Figure 4.**
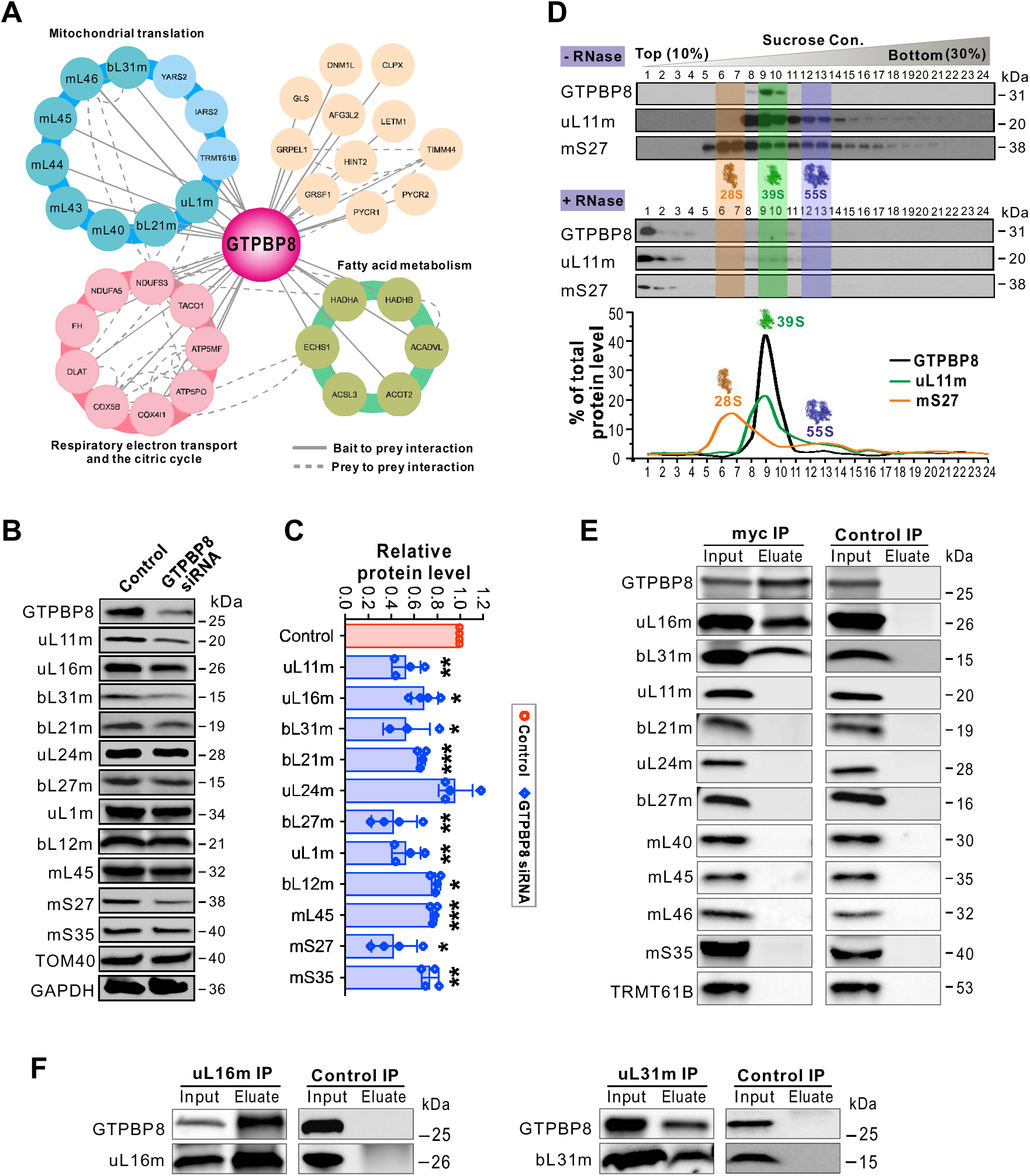
GTPBP8 associates with the large mitoribosomal subunit. (**A**) The comprehensive interactome of GTPBP8 using a proximity-dependent biotin identification (BioID) analysis. The protein complex/functional groups were indicated by different colors. The mitoribosomal large subunit (mt-LSU) proteins are highlighted in cyan. Bait to bait interactions are shown with solid lines, and prey-to-prey interactions are shown with dashed lines. (**B**) Steady state levels of mitoribosomal proteins after GTPBP8 depletion. (**C**) Quantification of the steady state levels of mitoribosomal proteins in panel B. The protein level was normalized to TOM40. Error bars represent the mean ± SD of three independent repetitions. * P≤ 0.05, ** P≤ 0.01, and *** P≤ 0.001, Student’s t-test. (**D**) Sucrose gradient sedimentation profile of GTPBP8 compared with the mt-LSU protein uL11m and small subunit protein mS27. RNAase-treated samples are in the lower panel. The mt-SSU, mt-LSU, and monosome fractions are highlighted by yellow, green, and purple, respectively. (**E and F**) Co-immunoprecipitation analysis of GTPBP8-associated proteins using antibody-coated Protein G Dynabeads^TM^. U2OS cells transfected with GTPBP8-myc for 48 h were used for the co-immunoprecipitations. The input and eluate fractions were analyzed by SDS-PAGE followed by immunoblotting with the indicated antibodies.

To corroborate the interaction of GTPBP8 with mitoribosomes, the mitoribosomal protein profile was examined by an isokinetic sucrose density gradient. Interestingly, GTPBP8 was predominantly detected in the mt-LSU fraction, corresponding to factions 9 and 10 in Figure 4B (39S mt-LSU), but not in the 55S monosome or the mitoribosomal small subunit (28S, mt-SSU) fraction (Figure 4D). To confirm that GTPBP8 was specifically associated with the mt-LSU and not a protein complex of similar density, we pre-treated mitochondrial lysates with ribonuclease A (RNaseA) to digest mitochondrial ribosomal RNA (mt-rRNA) and disrupt the mitoribosomal integrity. RNaseA treatment shifted the detected mt-LSU protein uL11m and the mt-SSU protein bL27m to less dense fractions together with the majority of GTPBP8, confirming that GTPBP8 associated with the mt-LSU (Figure 4D). To understand the molecular details of GTPBP8’s interaction with mt-LSU, we performed co-immunoprecipitation (co-IP) experiments. A previous finding showed that downregulation of the bacterial GTPBP8 homolog YsxC resulted in immature ribosomal particles lacking some large ribosomal subunit proteins (49). Combining with our BioID results, several proteins, including uL11m, uL16m, bL21m, uL24m, bL27m, bL31m, mL40, mL45, mL46, TRMT61B, and an mt-SSU protein mS35, were selected as prey for the immunoprecipitation experiment. Our data revealed that uL16m and bL31m were clearly co-precipitated with GTPBP8, but not with the other tested proteins (Figure 4E). To confirm the association of GTPBP8 with uL16m and bL31m, we performed reverse co-immunoprecipitation experiments using uL16m or bL31m as bait. GTPBP8-myc can be mutually immunoprecipitated with uL16m and bL31m (Figure 4F), demonstrating that GTPBP8 is specifically associated with uL16m and bL31m.

### GTPBP8 is required for the mt-LSU assembly at a late stage

To understand the role of GTPBP8 in association with mitoribosomes, we analyzed the biogenesis of 39S mt-LSU, 28S mt-SSU, and 55S monosomes in GTPBP8-depleted cells (Figure 5). Strikingly, silencing of GTPBP8 led to a profound decrease of 55S monosomes and the concomitant accumulation of the mt-LSU subunit (Figures 5A-5D), suggesting that the immature mt-LSU particles could not be assembled into monosomes. The changes were similar for all the MRPs tested, including uL11m, uL24m, uL16m, mS27, and mS35. In contrast, the gradient sedimentation profile of the 28S mt-SSU was not affected by the depletion of GTPBP8, except for the reduction of the 55S monosome level (Figures 5A-5D). The rRNA profiles of the gradients showed that the level of 12S rRNA in the mtSSU fraction was similar in WT and GTPBP8-depleted cells, but the 16S rRNA level increased in the mt-LSU fraction but decreased in the monosome fraction in the GTPBP8-depleted cells, reflecting the abnormal accumulation of mt-LSU and impaired monosome formation (Figures 5E and 5F). The overall abundance of mitochondrial rRNAs was not significantly altered in GTPBP8-depleted cells (Figure 5G), which was also true for tRNA^Val^ (Figure 5H). The comparable sedimentation patterns of the mt-LSU protein markers in WT and GTPBP8 depleted cells indicate that GTPBP8 acts in the late stage of mt-LSU maturation. This is consistent with the result of GTPBP8 specifically binding to the late assembly proteins uL16m and bL31m (Figures 4E and 4F). Taken together, silencing of GTPBP8 results in the accumulation of late mt-LSU assembly intermediates and a profound decrease of the fully assembled 55S monosomes, indicating that GTPBP8 is involved in the late stage of mitoribosomal maturation.

**Figure 5.**
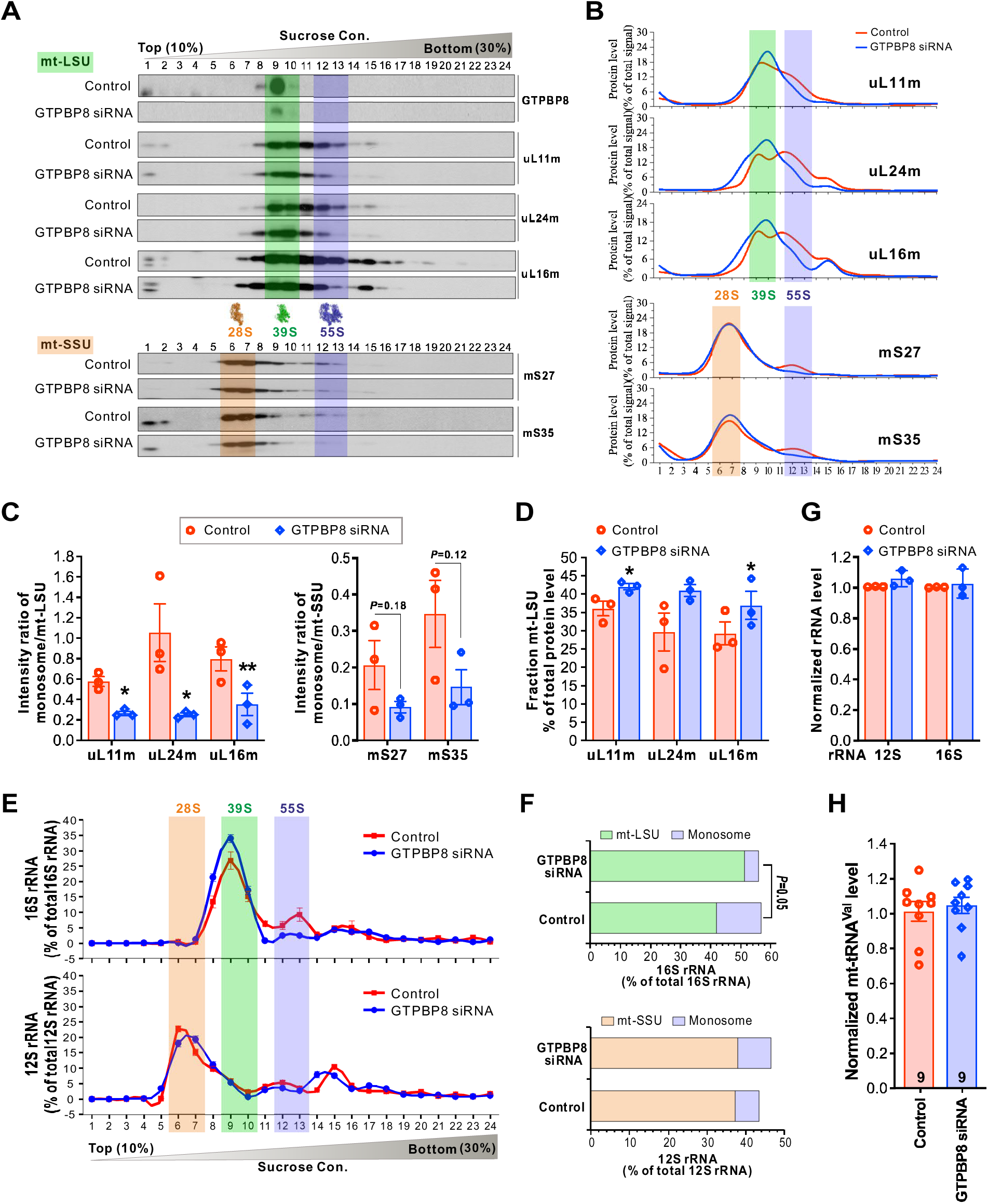
Loss of GTPBP8 causes aberrant accumulation of the large mitoribosomal subunit and reduction of fully assembled 55S monososmes. (**A**) Effects of GTPBP8 depletion on the sucrose gradient sedimentation profiles of the mitoribosomal proteins. Total cell lysates (900 μg protein) were separated on a 10–30% isokinetic sucrose gradient. 24 equal fractions were taken and analyzed by western blotting with the indicated antibodies. The mt-SSU, mt-LSU, and monosome fractions are highlighted by orange, green, and purple, respectively. (**B**) Quantified gradient distribution of indicated mitoribosomal proteins upon GTPBP8 depletion. Lines represent the mean of three independent repetitions. The mt-SSU, mt-LSU, and monosome fractions are highlighted by orange, green, and purple, respectively. (**C**) The intensity ratios of the monosome (fractions 12 and 13) against the mitoribosomal large subunit (fractions 9 and 10) or the mitoribosomal small subunit (fractions 6 and 7). (**D**) The protein levels in the mitoribosomal large subunit fractions of the sucrose gradient, normalized to the total protein. (**E**) The 16S and 12S rRNA profiles in the individual sucrose gradient fractions in WT and GTPBP8-depleted cells. The mt-SSU, mt-LSU, and monosome fractions are highlighted by orange, green, and purple, respectively. (**F**) The quantified relative levels of rRNA in the mt-LSU (fractions 9 and 10), mt-SSU (fractions 6 and 7), and monosome fractions (fractions 12 and 13). (**G**) Steady-state levels of mitochondrial rRNAs detected by quantitative RT-PCR. Cytoplasmic 18S rRNA was used as a loading control. The relative 12S and 16S rRNA levels were normalized to 18S rRNA. (**H**) Steady-state level of mitochondrial transfer RNA valine (mt-tRNA^Val^) detected by quantitative RT-PCR. The total mt-tRNA^Val^ levels were normalized to 18S rRNA. All the experiments in Figure 5 were done in U2OS cells treated with siRNA for 5 days. Error bars represent the mean ± SD of three independent repetitions. * P≤ 0.05 and ** P≤ 0.01, Student’s t-test.

To gain further evidence for the role of GTPBP8 in the hierarchical incorporation of the mitoribosomal proteins with the growing mt-LSU particles, we individually knocked down eight mt-LSU proteins assembled at distinct stages, and GTPBP8 (Figures 6A and 6B). Depletion of mL45, mL64, and bL36 decreased the level of most of the tested MRPs, while downregulation of the other MRPs affected the MRP levels in a less dramatic manner (Figures 6A and 6B). Loss of GTPBP8 significantly reduced the abundance of most tested MRPs, including uL16m, further demonstrating its critical role in mitoribosomal assembly. Interestingly, the GTPBP8 level tended to increase when the late assembly mt-LSU proteins uL16m and bL31m were depleted and to decrease slightly when the other mt-LSU proteins were depleted (Figures 6A and 6B). This result supports the role of GTPBP8 in the late assembly steps of mt-LSU, suggesting that GTPBP8 acts before the late assembly proteins uL16m and bL31m. This is consistent with the recent findings from the structures of the assembly intermediates of the mitoribosomal large subunit in the parasite *Trypanosoma brucei* (50), showing that uL16m and bL31m were missing in the structure of the assembly intermediate containing mt-EngB (the parasite homolog of GTPBP8).

**Figure 6.**
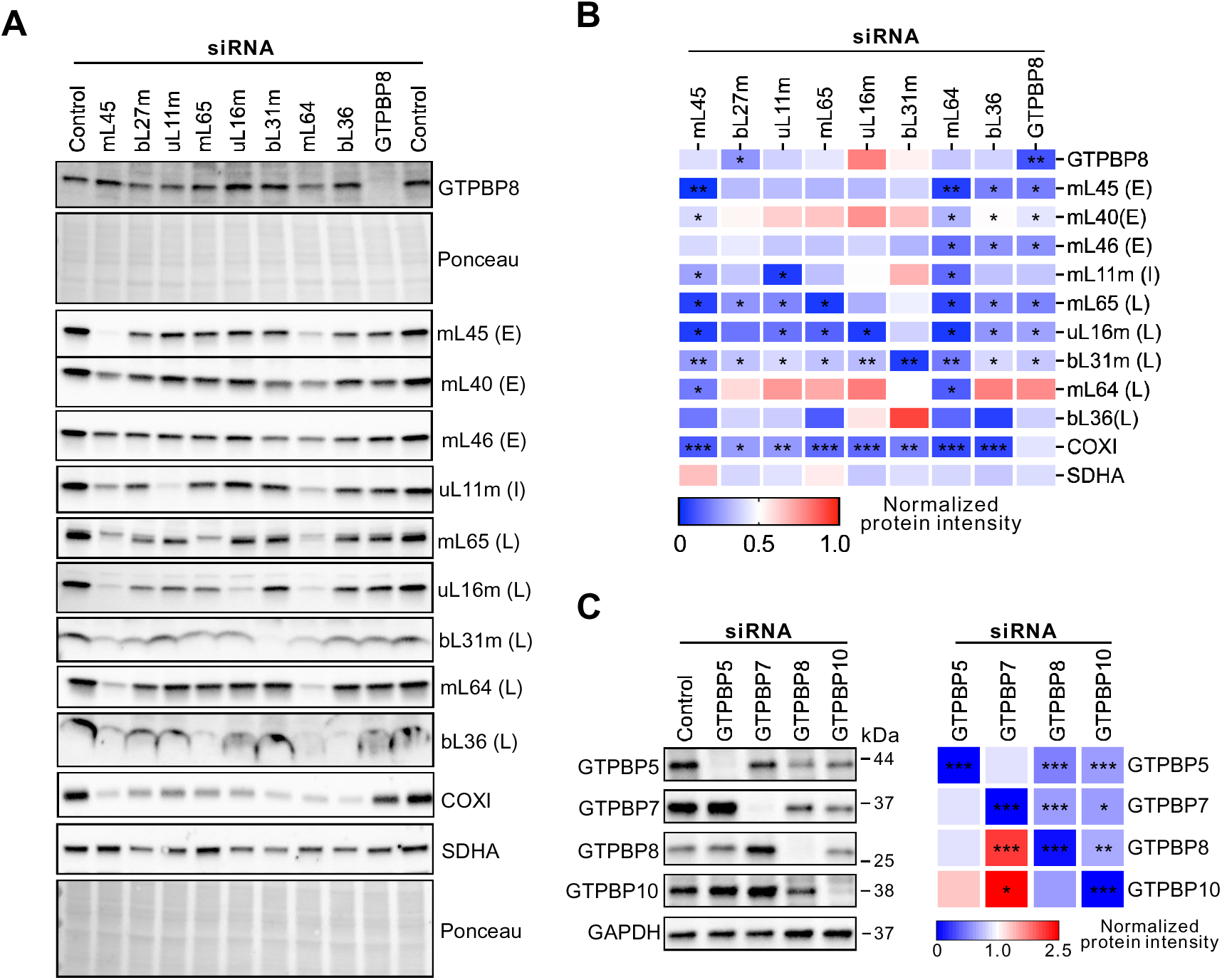
Changes of the mitoribosomal protein abundance after depletion of GTPBP8 or target proteins. (**A**) Representative images of immunoblot analysis of the steady-state levels of GTPBP8 and mitoribosome proteins after depletion of target proteins in U2OS cells. Antibodies are listed on the right side. E represents early, I intermediate and L late assembly MRPs. (**B**) Quantification of the steady state levels of indicated proteins in panel A. Error bars represent the mean ± SD of three independent repetitions. * P≤ 0.05, ** P≤ 0.01, and *** P≤ 0.001, Student’s t-test. (**C**) Changes of the steady-state GTPase levels after 5 days of siRNA treatment. The quantification results on the right were normalized against GAPDH and represented three independent experiments. *P≤0.05, **≤0,01, and ***≤0,001, Student’s t-test.

The so far characterized four mitochondrial GTPases critical for the mt-LSU assembly include GTPBP5, GTPBP6, GTPBP7, and GTPBP10 (18, 22, 28, 44, 46, 47, 51, 52). To examine the functional link among these GTPases, we knocked down each GTPase individually and detected the level changes of the others (Figure 6C). GTPBP6 was not studied due to the poor quality of the antibody. The result revealed that GTPBP8 abundance increased when GTPBP7 was depleted but decreased when GTPBP10 was depleted. Knockdown of GTPBP10 decreased the levels of all four GTPases, whereas GTPBP5 depletion did not affect the abundance of the other GTPases significantly although the levels of GTPBP7 and GTPBP8 tended to decrease and GTPBP10 level increased. Moreover, downregulation of GTPBP7 increased the GTPBP10 level, but silencing of GTPBP10 decreased the GTPBP7 level. Collectively, our data revealed that the abundance of these GTPases is mutually affected in cells, suggesting that they are functionally linked. Depletion of GTPBP8 affected the abundance of the other GTPases involved in mt-LSU maturation, further supporting its role in mt-LSU assembly.

### Rescue of defects in mitoribosomal assembly and mitochondrial translation by GTPBP8

To assess whether the defects in mitoribosomal assembly and mitochondrial protein translation were specific consequences of GTPBP8 silencing, we performed rescue experiments (Figures 7 and S6B-S6C). The impaired biogenesis of 55S monosomes and aberrant accumulation of mt-LSU in fraction 10 as a consequence of defective maturation were rescued by the expression of GTPBP8 in GTPBP8-depleted cells (Figures 7A and 7B). We quantified and confirmed rescue in GTPBP-myc-expressing conditions by running relevant sucrose gradient fractions corresponding to mt-LSU, mt-SSU, and monosome on denaturing gels (Figure 7C). Moreover, the newly synthesized mtDNA-encoded proteins reflecting mitochondrial translation were rescued by the expressed GTPBP8-myc in siRNA-treated cells (Figures 7D, 7E, S6B, and S6C). In line with the rescue of monosome abundance and mitochondrial translation, the reduced protein levels of MT-CO1, Mt-CytB, uL16m, and uL11m were also significantly rescued (Figures 7F and 7G). These results demonstrate that the defects in mitoribosomal assembly and mitochondrial translation are specific to GTPBP8 depletion. In conclusion, our results reveal that GTPBP8 is essential for the maturation of mt-LSU and the biogenesis of mitoribosomes, which are required for mitochondrial translation and mitochondrial respiration.

**Figure 7.**
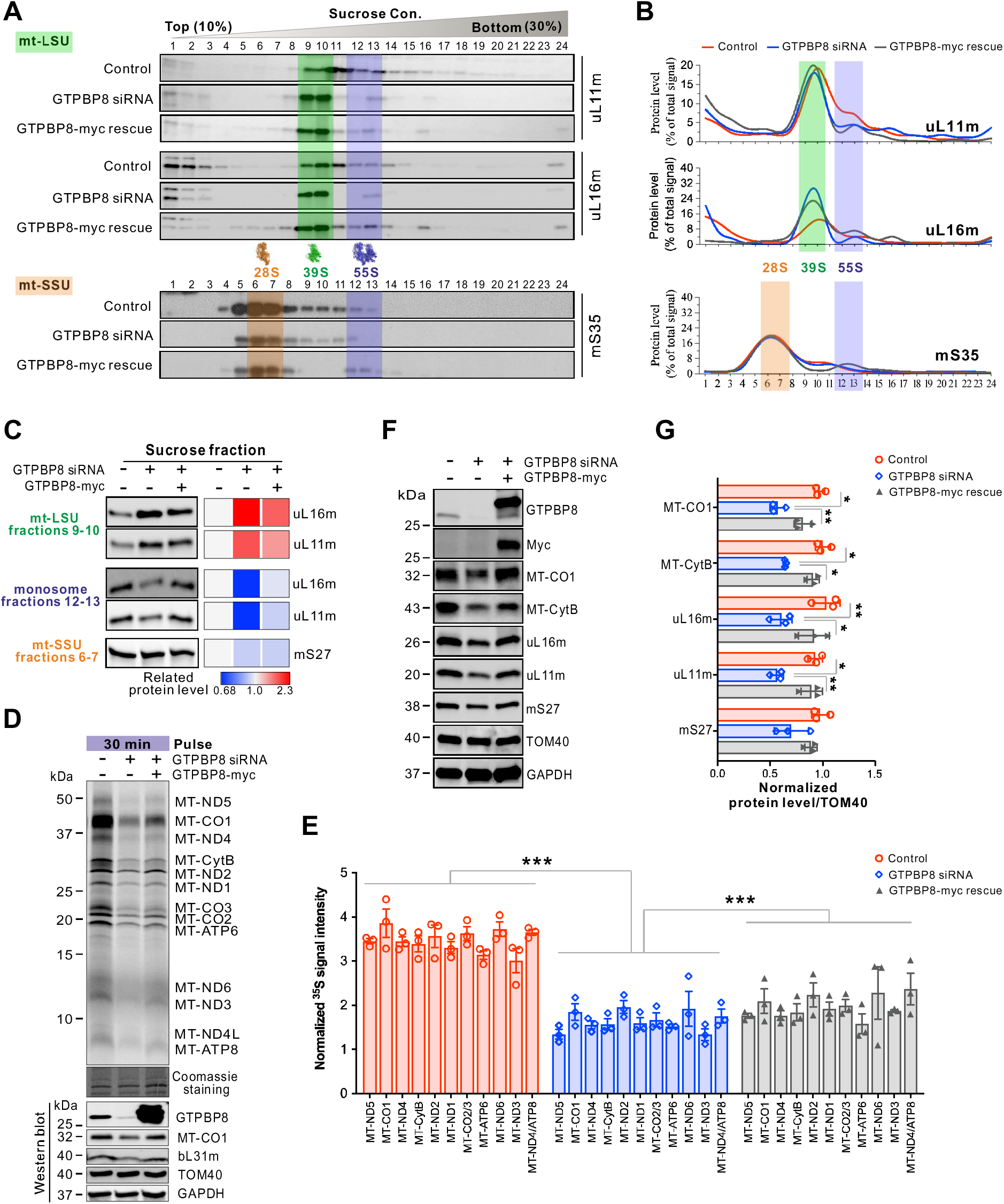
Defects in mitoribosomal assembly and mitochondrial translation caused by GTPBP8 depletion can be rescued by GTPBP8 expression. (**A**) The representative mitoribosome profiles in GTPBP8-depleted and GTPBP8-myc-expressed cells. The mt-SSU, mt-LSU, and monosome fractions are highlighted by orange, green, and purple, respectively. (**B**) Quantified gradient distribution of indicated mitoribosomal proteins upon GTPBP8-silencing and GTPBP8-myc-expression. Lines represent the mean of three independent experiments. The mt-SSU, mt-LSU, and monosome fractions are highlighted by orange, green, and purple, respectively. (**C**) Rescue of the indicated protein levels in the mt-LSU (fractions 9 and 10), mt-SSU (fractions 6 and 7) and monosome fractions (fractions 12 and 13). Quantification of mitoribosomal protein abundance was show on the right. (**D**) Rescue of mitochondrial translation defect caused by GTPBP8 depletion. Top: Metabolic labeling of mitochondrial translation products with [^35^S]-methionine/cysteine for 30 min in control, GTPBP8-depleted or rescued cells. Bottom: immunoblotting for the steady state level of GTPBP8. (**E**) Quantification of ^35^S intensity for mitochondrially-translated proteins relative to the corresponding intensity in Coomassie staining. Error bars represent the mean ± SD of three independent repetitions. * P≤ 0.05, ** P≤ 0.01, and *** P≤ 0.001, Student’s t-test. (**F**) Steady state levels of MT-CO1 and mitoribosomal proteins in control, GTPBP8 depleted, and GTPBP8-myc rescued cells. TOM40 and GAPDH were used as loading controls. (**G**) Quantification of protein levels in Panel F. * P≤ 0.05, ** P≤ 0.01, and *** P≤ 0.001, Student’s t-test.

## Discussion

Mitoribosome biogenesis is a coordinated and hierarchical process that requires a significant fraction of cellular resources devoted to macromolecular synthesis and trafficking. The mtDNA contributes only to the RNA components of the mitoribosomes, and mitoribosomal biogenesis requires the synthesis of 82 human mitoribosomal proteins encoded by nuclear DNA, translated on cytoplasmic ribosomes, and individually imported into mitochondria. These processes occur in a highly orchestrated and regulated assembly line, which is coordinated by a number of assembly factors. While the high-resolution cryo-EM structures of the mammalian mitoribosomes have recently become available, knowledge of the molecular details of the mammalian mitoribosomal assembly, specific factors involved, and sequence is still very limited. In this study, we demonstrate that GTPBP8 is a mitochondrial protein required for the late assembly of the large mitoribosomal subunit (39S mt-LSU) and is essential for mitochondrial translation. GTPBP8 is imported into mitochondria by an N-terminal pre-sequence and binds specifically to the 39S mt-LSU. Downregulation of GTPBP8 caused an abnormal accumulation of late mt-LSU immature particles, which led to a sharp reduction of functional 55S monosomes. Regulation of mitoribosome assembly and biogenesis is paramount to mitochondrial respiration and, thus, to cell viability, proliferation, and differentiation. Indeed, our data showed that the loss of GTPBP8 led to defective mitochondrial translation, which consequently impaired the assembly of OXPHOS complexes and mitochondrial respiration. As a result, cellular ATP production was reduced, resulting in decreased cell proliferation.

The mitochondrial translation system is of prokaryotic origin. Thus, many proteins involved in mitoribosomal biogenesis have bacterial homologs (15). The bacterial homolog of GTPBP8, Ysxc/Yiha, has been reported to be essential for *B. subtilis, E. coli, S. pneumoniae, H. influenzae*, and *M. genitalium* (29–31, 49). Depletion of YsxC resulted in defects in ribosome biogenesis with the accumulation of aberrant large subunit intermediates, ultimately causing severe growth deficiency (29–31, 49). This role in ribosomal assembly is conserved for human GTPBP8. Our data show that human GTPBP8 exclusively associates with the mt-LSU, neither with the mt-SSU nor the monosome, suggesting that GTPBP8 transiently binds the 39S mt-LSU during mt-LSU assembly but detaches from the mt-LSU before formation of the monosome. The comparable sedimentation pattern of the mt-LSU protein markers in GTPBP8-depleted cells indicates a role for GTPBP8 at the late stage of the mt-LSU assembly. Furthermore, GTPBP8 interacts with the mitoribosomal proteins uL16m and bL31m, which both are assembled into the 39S mt-LSU at late stages. uL16m in the A site and bL31m in the central protuberance were found missing in YsxC-depleted cells (49), suggesting the importance of GTPBP8 for the assembly of uL16m and bL31m. Consistently, uL16m was absent from the large subunit assembly intermediate of the parasite *Trypanosoma Brucei*, in which mt-EngB (human GTPBP8), Mtg1 (human GTPBP7), and mt-EngA were present (50), indicating that uL16m is assembled to the large subunit after the recruitment of these GTPases. uL16 was also found missing in the bacterial immature ribosomal intermediates upon depletion of RbgA (GTPBP7) (49, 53, 54), supporting the notion that both GTPBP8 and GTPBP7 are indispensable for the assembly of uL16m. Moreover, a previous study reveals that a mitochondrial assembly of ribosomal large subunit protein 1 (MALSU1) forms a preassembly complex with uL14m, making the mt-LSU binding site accessible for the immediate association of GTPBP7, which further enables the binding of uL16m to the mt-LSU (51).

Ribosome biogenesis is assisted by an increasing number of assembly factors, and the guanosine triphosphate hydrolases (GTPases) are the most abundant class. In mammalian mitoribosomes, only a few GTPases have been characterized so far. Two GTPases, the mitochondrial ribosome-associated GTPase 3 (MTG3) [also known as nitric oxide-associated protein 1 (NOA1) or C4ORF14] and Era G-protein-like 1 (ERAL1), are involved in mt-SSU biogenesis (27, 55–57). Four GTPases are required for mt-LSU biogenesis (18), including GTPBP5 (also known as MTG2/OBGH1, bacterial ObgE), GTPBP6 (bacterial HflX), GTPBP7 (also known as MTG1, bacterial RbgA), and GTPBP10 (also known as OBGH2) (22, 28, 44, 46, 47, 51). GTPBP7 (MTG1) was the first GTPase to be thoroughly studied (44, 51). Similar to GTPBP8, GTPBP7 depletion lead to abnormal accumulation of mt-LSU and loss of the 55S monosome (51). GTPBP7 facilitates the incorporation of late assembly MRPLs, at least bL36m and bL35m, into the mt-LSU and remains bound to the mt-LSU until the maturation of the mtLSU is completed (51). GTPBP5 (MTG2, OBGH1) and GTPBP10 (OBGH2) are homologs of the Obg proteins in bacteria. Both GTPBP5 (MTG2, OBGH1) and GTPBP10 (OBGH2) are involved in the late assembly stages of mt-LSU and the formation of monosomes, but they act at distinct time points (28, 46, 47, 52). In contrast, loss of GTPBP6 stalled mtLSU biogenesis at a very late assembly stage when all of the 52 MRPs, including the last assembly protein bL36m, as well as GTPBP5, GTPBP7, and GTPBP10, were bound to the mt-LSU complex (22), demonstrating that GTPBP6 is required for the very final steps of mt-LSU maturation. Accordingly, a sequential action of these GTPases on the large mitoribosomal subunit assembly was proposed (18), with GTPBP10 being the first assembly factor, followed by GTPBP5, GTPBP7, and GTPBP6. Recently, the parasite homolog of GTPBP8, mt-EngB, was found to bind a LSU assembly intermediate together with the other two GTPases, mt-EngA and Mtg1 (GTPBP7) (50). mt-EngB is recruited only to a later assembly intermediate complex, where mt-EngA interacts with Mtg1 to form a docking site for mt-EngB (50), suggesting that mt-EngB (GTPBP8) acts downstream of Mtg1 (GTPBP7) and mt-EngA. Interestingly, depletion of bacterial YsxC (GTPBP8) or RbgA (GTPBP7) resulted in similar extensive structural changes in several functional domains, including the central protuberance, the GTPase associated region, and key RNA helices in the A, P, and E sites of the bacterial 50S large subunit (49, 50), indicating that YsxC and RbgA may share functional roles in the maturation of these sites. Structurally, mt-EngB (GTPBP8) and Mtg1 (GTPBP7) are in contact with each other in the assembly intermediate (50). A conformational change of Mtg1 (GTPBP7) was observed in the assembly intermediate when mt-EngB (GTPBP8) was bound (50). Simultaneous binding of these GTPases to an assembly intermediate suggests that GTPBP8 works in conjunction with the other GTPases, particularly GTPBP7, during mt-LSU assembly. However, the molecular mechanism underlying their cooperative interplay in the maturation of mt-LSU requires further studies.

In conclusion, our data uncover human GTPBP8 to be an important mitochondrial GTPase involved in the late-stage assembly of the large mitoribsomal subunit and essential for the maturation of the translationally competent monosomes. The previous studies, together with our work, have expanded the model for mitoribosome assembly facilitated by the mt-LSU associated GTPases (18). These steps for mt-LSU maturation are critical because depletion of these GTPases results in immature mt-LSU intermediates and impaired monosome formation, and subsequently in translational deficiency.

## Materials and Methods

### Plasmid Construction

Full-length human GTPBP8 coding sequence (GenBank accession no. NM_014170.2) in the pENTR221 vector (ORFeome Library; Biocentrum Helsinki Genome Biology Unit) was cloned with XhoI and BamHl restriction sites into the pEGFP-N1 vector (Takara Bio Inc.). GTPBP8 was cloned into the pHAT vector (provided by the EMBL Protein Expression and Purification Facility) with BcuI and BamHI cloning sites. GTPBP8-myc was constructed from GTPBP8-GFP with GFP replaced by myc. GTPBP8_1-46aa_-GFP and GTPBP8_47-284aa_-GFP were subcloned into the pEGFP-N1 vector. Inserts were sequenced to confirm the cloning. GTPBP8 was cloned into the pGAT2 vector using NcoI and XhoI restriction sites. LR recombination was performed between the GTPBP8 entry clone and the MAC-C destination vector (32) to generate the MAC-tagged GTPBP8 expresion vector.

### GTPase Activity Assay

The GTPase activity of purified GTPBP8 protein was assessed by using the Enzcheck assay kit (E6646, Life Technologies) according to the manufacturer’s instructions. GTPBP8 (12 μM) was incubated with various concentrations of GTP (0, 0.094, 0.1875, 0.375, 0.75, 1.5, and 3 mM) in the kit’s reaction buffer at 25 °C. Absorbance at 360 nm was immediately recorded every 4 min for a duration of 60 min using the VarioskanTM LUX multimode microplate reader (Thermo Fisher Scientific) with SkanIt Software 5.0. The amount of inorganic phosphate released from GTP hydrolysis at each time point was determined by extrapolation using a phosphate standard curve. Kinetic parameters were determined by fitting the data to the Michaelis-Menten equation using OriginPro 2018.

### Cell Culture and Transient Transfections

Human osteosarcoma (U2OS) cells, which regularly tested negative for mycoplasma with MycoAlert Plus (Lonza), were cultured in high-glucose DMEM containing 10% FBS (12-614F, Lonza), 10 U/ml Penicillin, 10 µg/ml Streptomycin and 20 mM L-glutamine (10378-016; Gibco) in a humidified 95% air/5% CO_2_ incubator at 37 °C. For transient transfections, cells were plated on culture plates and transfected the next day with DNA constructs using FugeneHD reagent (#E2311, Promega), according to the manufacturer’s instructions.

For silencing experiments, cells were transfected with 30 pmol of target gene or control siRNA directly after replating on the 6 cm culture plate using Lipofectamine® RNAiMAX (#13778-075, Thermo Fisher), according to the manufacturer’s instructions. Cells were transfected again after 48 h with the same amount of siRNA and incubated for another 72 h. All the samples were subjected to mass spectrometry and western blot analysis. The following siRNA target sequences were used: GTPBP5 siRNA: 5’-CGGUGGACACGUCAUUCUGTT-3’ (134621, Ambion); GTPBP7 siRNA: 5’-GCAACACUUAGAAGGAGAAGGCCUA-3’ (1362318, Invitrogen); GTPBP8 siRNA#1: 5’-AAAGTTACTATGTAAGCCTAA-3’ (SI04257974); siRNA #2: 5’-AGCGACTGAGCCGCTATAATA-3’ (SI04232011, Qiagen); #3: 5’-AAGCATCGATAGGTAAGTTGA-3’ (SI03130008, Qiagen); #4: 5’-CCGGTTTAGCTGAAGATTCAA-3’ (SI00443779, Qiagen), GTPBP10 siRNA 5’-TTGCGTGTTGTTCAGAAAGTA-3’ (SI04308647, Qiagen), uL16m siRNA: 5’ TACGGAGTTTACAGAAGGCAA-3’ (SI00648291, Qiagen) and Negative control siRNA (SI03650318, Qiagen), MRPL45 siRNA: 5’-CTGGAGTATGTTGTATTCGAA-3’ (SI00649005, Qiagen), bL27m siRNA 5’CAGGCAGACGCCAAGGCATTA-3’, uL11m siRNA 5’-AGGAAGGAGGTCACACCAATA-3’ (SI04135684, Qiagen), mL65 siRNA 5’CAAGCUAUGUAUCAAGGAUtt3’ (s21375, ThermoFisher Scientific) bL31m siRNA 5’-CCAGGCTTATGCACGACTCTA-3’ (SI00649271, Qiagen), mL64 siRNA 5’-AAGAACGCGAATGGTACCCGA-3’ (SI02652349, Qiagen), and bL36m siRNA 5’-CGGTGGTACGTCTACTGTAAA-3’ (SI04156299, Qiagen).

In the rescue experiments, U2OS cells were treated with either control or GTPBP8 siRNA altogether for 8 days (siRNA transfection on the 1^st^ and 3^rd^ day on replated cells) out of which the last 48 hours the GTPBP8-depleted cells were rescued with the GTPBP8-myc construct using FugeneHD reagent according to the manufacturer’s instructions. The control cells were replated after 5 days of AllStars negative-control siRNA (QIAGEN) treatment, and all the control, GTPBP8 depleted, and GTPBP8 rescued cells were collected at the same time on the 8^th^ day.

### Immunofluorescence Microscopy

U2OS cells were grown on coverslips and fixed with 4% paraformaldehyde for 20 min, washed three times with PBS, and permeabilized with 0.1% Triton X-100 in PBS for 5 min. For antibody staining, permeabilized cells were incubated with primary antibody for 1 h at 37°C. Coverslips were washed with Dulbecco plus 0.2% BSA and incubated with fluorescence-conjugated secondary antibody for 1h at RT. Cells were mounted in Mowiol supplemented with 1,4-Diazabicyclo[2.2.2]octane (DABCO). The following antibodies were used: rabbit anti-GTPBP8 (1:100, HPA034831, Sigma), mouse anti-TOM20 (1:200, sc-17764, Santa Cruz Biotechnology), mouse anti-myc (1:100, sc-40, Santa Cruz Biotechnology), rabbit anti-COXIV (1:100, MA5-15078, ThermoFisher Scientific). Cells were imaged with a DM6000B microscope (Leica Biosystems) equipped with a Hamamatsu Orca-Flash 4.0 V2 sCMOS camera and LAS X software (Leica), using 63x/1.4-0.60 HCX PL Apochromat objective and brightline filters (Semrock): GFP-4050B (excitation, 466/40; emission, 525/50), TRITC-B (excitation, 543/22; emission, 593/40).

### Immunoblotting and Antibodies

Cells were solubilized in PBS containing 1% dodecyl maltoside (DDM), 1 mM PMSF, 10 mM sodium azide, 10 mM sodium ascorbate, and 5 mM Trolox. Protein concentrations were measured by the Bradford assay (Bio-Rad Laboratories). Equal amounts of proteins were loaded and separated by Tris-glycine SDS-PAGE, and transferred to PVDF membranes. Primary antibodies were incubated overnight at 4°C and detected the following day with HRP-linked secondary antibodies (1:10000) using the ECL reagent. The following primary antibodies were used: mouse anti-ATP5A (1:1000, 14748, Abcam), rabbit anti-GFP (1:4000, 50430-2-AP, Proteintech), mouse anti-MT-CO1 (1:2000, 14705, Abcam), rabbit anti-COXIV (1:1000, MA5-15078, ThermoFisher Scientific), rabbit anti-CytB (1:3000, 55090-1-AP, Proteintech), mouse anti-CytC (1:10000, 05-479, Millipore), goat anti-GAPDH (1:1000, sc-20357, Santa Cruz Biotechnology), rabbit anti-GTPBP5 (1:2000, 20133-1-AP, Proteintech), rabbit anti-GTPBP7 (1:4000, 13742-1-AP, Proteintech), rabbit anti-GTPBP8 (1:500, HPA034831, Sigma), rabbit anti-GTPBP10 (1:500, NBP1-85055, Novus Biologicals), mouse anti-HSP60 (1:20000, SMC-110, StressMarq Biosciences), rabbit anti-uL1m[MRPL1] (1:2000, 16254-1-AP, Proteintech), rabbit anti-uL2m[MRPL2] (1:2000, 16492-1-AP, Proteintech), rabbit anti-uL11m[MRPL11] (1:6000, 15543-1-AP, Proteintech), rabbit anti-bL12m[MRPL12] (1:4000, 14795-1-AP, Proteintech), rabbit anti-uL16m[MRPL16] (1:500, HPA054133, Sigma), rabbit anti-bL21m[MRPL21] (1:3000, PA5-31939, ThermoFisher Scientific), rabbit anti-uL24m[MRPL24] (1:2000, 16224-1-AP, Proteintech), rabbit anti-bL27m[MRPL27] (1:1000, 14765-1-AP, Proteintech), rabbit anti-mS27[MRPS27] (1:1000, 17280-1-AP, Proteintech), rabbit anti-mL65[MRPS30] (1:1000, 18441-1-AP, Proteintech), rabbit anti-mS35[MRPS35] (1:10000, 16457-1-AP, Proteintech), rabbit anti-bL36m[MRPL36] (1:1000, ab126517, Abcam), rabbit anti-mL40[MRPL40] (1:1000, HPA006181, Sigma), rabbit anti-mL45[MRPL45] (1:4000, 15682-1-AP, Proteintech), rabbit anti-mL46[MRPL46] (1:2000, 16611-1-AP, Proteintech), rabbit anti-bL31m[MRPL55] (1:1000, 17679-1-AP, Proteintech), rabbit anti-mL64[MRPL59] (1:500, 16260-1-AP, Proteintech), mouse anti-myc (1:1000, sc-40, Santa Cruz Biotechnology), mouse anti-NDUFA9 (1:1000, 20312-1-AP, Proteintech), mouse anti-TOM20 (1:1000, sc-17764, Santa Cruz Biotechnology), rabbit anti-TOM40 (1:1000, 18409-1-AP, Proteintech), rabbit anti-TRMT61B (1:500, 26009-1-AP, Proteintech), mouse anti-SDHA (1:1000, 14715, Abcam).

### Mitochondrial Isolation and Submitochondrial Fractionation

Mitochondria were isolated and purified as described previously (33). Briefly, cells were collected and resuspended in a mitochondrial isolation buffer (10 mM Tris-MOPS, 1 mM EGTA, 200 mM sucrose, pH 7.4) and homogenized with a Teflon potter (Potter S, Braun). Cells that had not been lysed were sedimented with 600 xg for 5 minutes at 4°C before being discarded. The supernatant was then centrifuged for 10 minutes at 4 °C at 9,000 x g. The resulting pellet was dissolved in ice-cold mitochondrial isolation buffer and centrifuged at 9000 xg for 10 min at 4°C. The pellet includes the mitochondrial fraction.

For identification of the sub-mitochondrial localization of GTPBP8, the isolated mitochondria were sub-fractionated to obtain mitoplasts using a phosphate swelling-shrinking method (34, 35). Briefly, purified mitochondrial pellets were resuspended in a swelling buffer (10 mM KH_2_PO_4_, pH 7.4) and incubated for 20 min with gentle mixing. In order to keep the mitoplasts intact, mitochondria were mixed with an equal volume of shrinking buffer (10 mM KH_2_PO_4_, pH 7.4, 32% sucrose, 30% glycerol, and 10 mM MgCl_2_) for another 20 min. The purified mitochondria and mitoplast were resuspended in a homogenization buffer (10 mM Tris-MOPS, 1 mM EGTA, 200 mM sucrose, pH 7.4) and treated with 0.2 mg/mL proteinase K with or without 1% NP-40 for 30 min. Proteinase K activity was quenched with 2 mM PMSF for 10 min. All steps were carried out on ice. 1% SDS was added to solubilize the mitochondrial proteins and immunoblotted as indicated.

To detect the mitochondrial membrane binding of GTPBP8, mitochondria were subjected to alkaline extraction in freshly prepared 0.1 M Na_2_CO_3_ at pH 11, 11.5, and 12. Mitochondrial membranes were pelleted at 72000 xg for 30 min at 4°C using an optimaTM ultracentrifuge (Beckman) with Beckman rotor (TLA 120). The membrane pellets (P) were dissolved in Laemmli loading buffer (2% SDS, 10% glycerol, 60 mM Tris-HCl, 0.005% bromophenol blue). Supernatants (S) were further precipitated using trichloroacetic acid (TCA) to a final concentration of 13% on ice for 30 min. After centrifugation at 20,000 xg for 30 min, pellets were washed twice with ice-cold acetone and dissolved in Laemmli loading buffer. Both pellets (P) and supernatant (S) were subjected to SDS-PAGE and western blot analysis.

### Transmission Electron Microscopy

For pre-embedding immuno-EM, cells were grown on coverslips and fixed with PLP fixative (2% formaldehyde, 0.01 M periodatem, 0.075 M lysine-HCl in 0.075 M phosphate buffer, pH 7.4) for 2 h at room temperature, as described previously (36). Cells were permeabilized with 0.05% saponin before being immunolabeled with anti-myc antibody in a 1:50 dilution, followed by Nano-gold mouse IgG in a 1:60 dilution. Nano-gold was silver enhanced using the HQ Silver kit for 5 min and gold toned with 0.05% gold chloride. After washing, the cells were further processed for embedding.

### Blue Native (BN)-PAGE

The abundance of respiratory chain complexes was analyzed by Blue Native polyacrylamide gel electrophoresis (BN-PAGE) using a NativePAGETM Novex® Bis-Tris Gel System according to manufacturer’s instructions. Briefly, mitochondria were suspended in 1% DDM in PBS on ice for 30 min and centrifuged at 22,000 xg for 30 min. The supernatant was supplemented with 0.2% Coomassie G250 and loaded on a 3-12% gradient gel. Proteins were transferred to the polyvinylidene fluoride (PVDF) membrane (Millipore, USA) which was subsequently fixed with 8% acetic acid for 15 min, washed with milliQ and methanol. The proteins were probed with the indicated antibodies.

### Viability Assays

Cell proliferation/viability was determined using the Thiazolyl Blue Tetrazolium Bromide (MTT) reagent. Briefly, MTT (M2128, Sigma-Aldrich) solution was added to the cell culture medium with a final concentration of 0.5 mg/ml at 24, 48, and 72 h during siRNA treatment. The medium was discarded after the incubation with MTT at 37°C for 4h. Subsequently, 150 µl DMSO was added to dissolve the formazan, and the quantity of formazan was measured by the absorbance at 570 nm using a Varioskan™ LUX multimode microplate reader (Thermo Fisher Scientific) with SkanIt Software 5.0.

### Measurement of Oxygen Consumption Rate (OCR) and ATP production

After treatment with control or GTPBP8 siRNA for 5 days, mitochondrial function was assessed by detecting OCR and ATP production. Cellular oxygen consumption rates were measured using a Seahorse XF96^e^ analyzer (Seahorse Bioscience, Agilent Technologies) as described before (37). The XF96 sensor cartridge was activated with 200 µl of XF96 calibration solution per well for 12 h at 37°C. U2OS cells were seeded onto XF96 cell culture microplates at 1×10^4^ cells per well and treated with GTPBP8 or control siRNA for 5 days. One hour prior to measurement, the culture medium was changed to serum-free and bicarbonate-free Dulbecco’s Modified Eagle’s medium supplemented with 10 mM glucose, 5 mM pyruvate and 5 mM glutamine. After incubation for 1 h at 37°C in a non-CO_2_ incubator, steady-state and post-intervention analyses were performed. Respiration was assessed by the injection of oligomycin (1 μM) to inhibit the mitochondrial ATP synthase, carbonyl cyanide-p-trifluoromethoxy-phenylhydrazone (FCCP, 1 μM) to collapse the mitochondrial membrane potential, and rotenone (1 μM) and antimycin A (1 μM) to inhibit the respiratory chain. OCR was measured before and after the addition of inhibitors at the indicated time points. The OCRs were normalized to total protein amounts to account for alterations in cell density.

Cellular ATP levels were detected by an ATP determination kit (A22066, ThermoFisher Scientific) according to the manufacturer’s instructions. After treatment with GTPBP8 siRNA for 5 days, cells were harvested and lysed on ice for 10 minutes using the DDM lysis buffer without proteinase inhibitors. The cell lysates were briefly sonicated and centrifuged at 16,000 xg for 10 minutes at 4°C. 10 µL supernatant was mixed with 90 µL reaction solution and luminescence measured using a Varioskan™ LUX multimode microplate reader (Varioskan Flash, Thermo Fisher Scientific, Inc., Waltham, MA USA). The amount of ATP were calculated from the standard curve and normalized to the total protein amount.

### Radioisotope Labeling of Mitochondrial Translation

Mitochondrial protein synthesis in cultured cells was detected through metabolic labeling with [^35^S] methionine/cysteine in the presence of anisomycin to inhibit cytoplasmic ribosomes (38). After treatment with targeted or control siRNA for 5 days, cells were washed three times with PBS and pretreated with 100 µg/ml anisomycin for 5 min to inhibit cytoplasmic translation. Subsequently, [^35^S] methionine/cysteine (EasyTag, PerkinElmer) was added with a final concentration of 400 μCi for pulse labeling assays. After pulse labeling for 30 min, cells were washed twice either with PBS (pulse samples) or medium without [^35^S] methionine/cysteine (chase labeling). After 6 h of chasing, cells were scraped and treated with benzonase^®^ Nuclease (E1014, Sigma) according to the manufacturer’s instructions. Gel loading buffer (186-mM Tris-HCl, pH 6.7, 15% glycerol, 2% SDS, 0.5 mg/ml bromophenol blue, and 6%-mercaptoethanol) was added to the samples before loading onto a 12– 20% gradient SDS-PAGE gel to separate proteins. The gel was then dried for exposure to a phosphor screen and scanned with a FUJIFILM FLA-5100. Gels were rehydrated in water and Coomassie-stained to confirm equal loading.

### Isokinetic Sucrose Gradient Assays

The sedimentation properties of GTPBP8 and the ribosomal proteins in sucrose gradients were analyzed as described before (36). Cells or mitochondria were lysed in 1% DDM lysis buffer (50 mM Tris, pH 7.2, 10 mM Mg(Ac)_2_, 40 mM NH4Cl, 100 mM KCl, 1% DDM, and 1 mM PMSF) for 20 minutes on ice, then centrifuged at 20,000 xg for 20 minutes at 4°C. A supernatant containing 900 g of total protein was loaded onto a 16 ml linear 10–30% sucrose gradient (50 mM Tris, pH 7.2, 10 mM Mg(Ac)_2_, 40 mM NH4Cl, 100 mM KCl, and 1 mM PMSF) and centrifuged for 15 hours at 4°C with 74,400 xg (SW 32.1 Ti; Beckman Coulter). 24 equal volume fractions were collected from the top and subjected to TCA precipitation. Samples were separated by SDS-PAGE for subsequent immunoblotting. For RNaseA treatments, RNaseA (ThermoFisher Scientific) was added to the cell lysate at a final concentration of 600 U/ml at the stage of protein sample preparation, and sucrose gradients were prepared as described.

### RNA and DNA isolation and Real-time Quantitative PCR (RT-qPCR)

Total RNA was extracted from the whole cells and sucrose fractions using the GeneJET RNA purification kit (K0731; Thermo Fisher Scientific) and Trizol reagent (Invitrogen), respectively. Single-stranded cDNA was synthesized from extracted mRNA with the Maxima first strand cDNA synthesis kit (K1671; Thermo Fisher Scientific). For quantification of mtDNA copy numbers, total DNA was extracted from cells using a total DNA, RNA, and protein isolation kit (Macherey-Nagel) following the manufacturer’s instructions, and mtDNA copy numbers were assessed using quantitative PCR analyses. β-globin was used to normalize the copy numbers of mtDNA. Quantitative PCR reactions were performed with Maxima SYBR Green/ROX (K0221; Thermo Fisher Scientific) by Bio-Rad Laboratories CFX96. GAPDH and 18S rRNA were used as controls for genes encoded in the nucleus and mitochondria, respectively. Changes in expression levels were calculated with the 2^−ΔΔCt^ method. Primers used in this study were: GTPBP8 F: 5’-GCGGCCAGAGGTGTGTTTTA-3’, GTPBP8 R: 5’-CCATAACCTGGCATGTCCAC-3’; GAPDH F: 5’-TGCACCACCAACTGCTTAGC-3’, GAPDH R: 5’-GGCATGGACTGTGGTCATGAG-3’; MT-CO1 F: 5’-CTCTTCGTCTGATCCGTCCT-3’, MT-CO1 R: 5’-ATTCCGAAGCCTGGTAGGAT-3’; MT-CytB F: 5’-TAGACAGTCCCACCCTCACA-3’, MT-CytB R: 5’-CGTGCAAGAATAGGAGGTGGA-3’;; mt-tRNAval F: 5’-TAGACAGTCCCACCCTCACA-3’, mt-tRNAval R: 5’-CGTGCAAGAATAGGAGGTGGA-3’; 12S rRNA F: 5’-TAGAGGAGCCTGTTCTGTAATCGA-3’, 12S rRNA R: 5’-TGCGCTTACTTTGTAGCCTTCAT-3’; 16S rRNA F: 5’-AGAGAGTAAAAAATTTAACACCCAT-3’, 16S rRNA R: 5’-TTCTATAGGGTGATAGATTGGTCC-3’; 18S rRNA F: 5’-CCAGTAAGTGCGGGTCATAAGC-3’, 18S rRNA R: 5’-CCTCACTAAACCATCCAA TCGG-3’; mtDNA F: 5’-ACCACAGTTTCATGCCCATCGT-3’, mtDNA R: 5’-TTTATGGGCTTTGGTGAGGGAGGT-3’; β-globin F: 5’-GGTGAAGGCTCATGGCAAGAAAG-3’ and β-globin R: 5’-GTCACAGTGCAGCTCACTCAGT-3’.

### Co-immunoprecipitation Analysis

Immunoprecipitation of expressed GTPBP8-myc was performed using Dynabeads^TM^ Protein G (10004D, Invitrogen) according to the manufacturer’s instructions. Briefly, 50 µl Dynabeads^TM^ Protein G beads were first coated with 1 µg mouse anti-MYC antibody (sc-40, Santa Cruz Biotechnology) or normal mouse IgG1 (sc-3877, Santa Cruz Biotechnology). After transient expression of GTPBP8-myc in U2OS cells for 48 h, mitochondria were extracted and lysed in ice-cold 1% DDM in PBS buffer containing a complete protease inhibitor cocktail. After 60 min of incubation on ice, the insolubilized material was removed by centrifugation at 20,000 xg for 20 min at 4°C. The supernatants were subsequently incubated with antibody-bound Dynabeads^TM^ Protein G overnight at 4 °C. After washing, the protein G Dynabeads-antibody-protein was eluted with 50 mM glycine (pH 2.8) to dissociate the complex at RT for 2 min. The samples were subjected to SDS-PAGE and immunoblotting.

### Proximity Protein Purification

U2OS were transfected with the MAC-tagged GTPBP8 expression vector or a control MAC-tagged GFP vector using Fugene 6 (Invitrogen) according to the manufacturer’s instructions. Cells were treated with 50 μM biotin for 24 h before harvesting, and one biological sample contains cells derived from 5 × 150 mm fully confluent dishes (approximately 5 × 10^7^ cells). Each sample has three biological replicates. Samples were frozen at − 80 °C until further use. For proximity purification (BioID), cells were lysed and loaded onto columns (Bio-Rad) containing the Strep-Tactin matrix (IBA, GmbH). The detailed procedures are described in (39).

### Mass Spectrometry

Protein lysate or pulldown samples were processed for a reduction of the cysteine bonds with 5 mM Tris(2-carboxyethyl)phosphine (TCEP) for 30 min at 37 °C and alkylation with 10 mM iodoacetamide. The proteins were then digested into peptides with sequencing grade modified trypsin (Promega V5113) at 37 °C overnight. Digested peptides were used for mass spectrometry analysis. The analysis was performed on a Q-Exactive mass spectrometer using Xcalibur version 3.0.63 coupled with an EASY-nLC 1000 system via an electrospray ionization sprayer (Thermo Fisher Scientific). Mass spectrometry was in a data-dependent acquisition mode using FTMS full scan (300-1700 m/z) resolution of 60,000 and collision-induced dissociation (CID) scan of the top 20 most abundant ions (40). BioID samples were run in quadruplicates, GTPBP8 knockdown samples in triplicates, respectively.

### Data Processing for Mass Spectrometry

For protein identification, Thermo.RAW files were uploaded into the Proteome Discoverer 1.4 (Thermo Scientific) and searched against the Sequest search engine of the selected human part of UniProtKB/SwissProt database (http://www.uniprot.org). All reported data were based on high-confidence peptides assigned in Proteome Discoverer (Thermo Scientific) with a 5% FDR by Percolator. The high confidence protein-protein interactions (HCIs) were identified using stringent filtering against GFP control samples and the contaminant Repository for Affinity Purification (CRAPome, http://www.crapome.org/) database (41). HCIs data were imported into Cytoscape 3.4.0 for the visualization. The known prey-prey interaction data were obtained from the IMEx database (http://www.imexconsortium.org/) (42). Gene ontology classification analysis was based on the DAVID bioinformatics resource (https://david.ncifcrf.gov/) (43). Hierarchical cluster was performed by centered correlation using Cluster 3.0 and the clusters were visualized with Tree View 1.1.6, and the matrix2png web server (http://www.chibi.ubc.ca/matrix2png/).

## Statistical Analysis

All of the experiments were done in triplicates or otherwise indicated. All data is reported as mean ± SD as indicated in the figure legends. Differences among groups were analyzed using the unpaired student t-test, one-way or two-way Anova.

## Data Availability

Mass spectrometry data are available at the MassIVE (https://massive.ucsd.edu/ProteoSAFe/dataset.jsp?task=b43e62d4e9894bca867e548485f7d9fa) with the DOI: doi:10.25345/C5FR38 and FTP download link (ftp://massive.ucsd.edu/MSV000086865/).

## Acknowledgements

We thank Biocentrum Helsinki Genome Biology Unit for providing gene constructs. We are grateful to the Light and Electron Microscopy Unit in University of Helsinki (Biocenter Finland) for their kind service in imaging. This work was supported by grants from the Academy of Finland (H.Z. decision No.266846; C.B.J. decision No.330098), Jane and Aatos Erkko Foundation (H.Z.).

## Author Contributions

H.Z. conceived the project. X.L. and M.V. performed the mass spectrometry analysis. L.W. performed the sea horse experiment together with O.E. T.H. performed the MRP and GTPBP10 depletion experiments and assisted in the Co-IPs, pull downs as well as in the rescue experiments. C.B.J. performed partial analysis of mass spectroscopy data after depletion of GTPBP8. L.W. carried out all the rest experiments and data analysis. H.Z. and L.W. wrote the manuscript with input from T.H., O.E., X.L., and C.B.J. The authors declare no competing financial interests.

