## Supplementary figures and tables for "GTPBP8 is required for mitoribosomal biogenesis and mitochondrial translation"

### Supplementary Information

#### Supplementary figure legends

**Figure S1. GTPBP8 is an evolutionarily conserved GTPase.** (**A**) Phylogenetic analysis of GTPBP8. Phylogenetic tree was built using Geneious 6.1.8. (**B**) Multiple sequence alignment of GTPBP8 orthologue in prokaryotes (P38424 for *B. subtilis* Ysxc and P0A6P7 for *E.coli* Yiha) and eukaryotes (Q8N3Z3 for human GTPBP8) using clustal omega server. The GTPase domains (G1-G5) were indicated in red boxes. The N-terminal mitochondrial targeting sequence (1-46aa) was indicated by a magenta bar. (**C**) The intrinsic GTPase activity of GTPBP8. Data is an average of three independent measurements with error bars (± SD) indicated.

**Figure S2. GTPBP8 is a mitochondrial protein.** (**A**) Immunoblotting detection of the expressed GTPBP8-myc in U2OS cells using an anti-myc antibody. (**B**) Immunoblotting detection of the expressed GTPBP8-GFP in U2OS cells using an anti-GFP antibody. Cells expressed only the GFP-vector were used as a negative control. (**C**) Immunoblotting analysis of subcellular localization of GTPBP8-myc. After transiently expressing GTPBP8-myc in U2OS cells for 48 h, cells were fractionated and the expressed GTPBP8-myc was detected in the whole-cell lysate (WCL), cytoplasm (CYT) and isolated mitochondria (MIT) with anti-myc and anti-GTPBP8 antibodies. Antibodies against mitochondrial proteins TOM40 and Cytochrome C, and GAPDH were used as controls. (**D**) Immunoblotting analysis of submitochondrial localization of GTPBP8-myc using proteinase K digestion of isolated mitochondria from GTPBP8-myc expressed cells. Mitochondria were subjected to hypotonic swelling or permeabilization with 1% NP-40, treated with 100 μg/mL proteinase K as indicated.

**Figure S3. Effects of GTPBP8 depletion on protein abundance of the oxidative phosphorylation complexes and functional networks by mass spectroscopy analysis.** (**A**) Main functionally affected biological processes by GTPBP8 depletion. (**B**) Steady-state protein levels of oxidative phosphorylation complexes after depletion of GTPBP8.

**Figure S4. Depletion of GTPBP8 impaired mitochondrial translation.** (**A**) Metabolic labeling of mitochondrial translation products with [$^{35}$S]-methionine/cysteine for indicated time in U2OS cells. GTPBP8 was depleted using siRNA #2 for 5 days. (**B**) The heatmap plot depicts the quantified $^{35}$S intensity in panel A for the mitochondrially-translated 13 proteins relative to the corresponding intensity in Coomassie staining. The color bar represents the normalized $^{35}$S intensity value between 0.32 and 1.

**Figure S5. Loss of GTPBP8 caused reduction of mitoribosomal protein abundance.** (**A**) Effects of GTPBP8 depletion on the steady-state levels of mitoribosomal proteins and mitoribosomal assembly related factors by label-free quantitative mass spectrometry (GTPBP8 silencing versus WT). Proteins with significant changes are labeled in yellow, and those without significant changes are indicated in cyan. U2OS cells were treated with control or GTPBP8 siRNA for 5 days. The silencing efficiency of GTPBP8 is shown in panel B. (**C**) Effects of GTPBP8 loss on the steady-state levels of mitoribosomal proteins.

**Figure S6. Loss of GTPBP8 caused defects in mitochondrial translation.** (**A**) Effects of GTPBP8 loss on the steady state levels of mitoribosomal proteins. GTPBP8 was depleted using siRNA #2. (**B**) Metabolic labeling of mitochondrial translation products with [$^{35}$S]-methionine/cysteine in U2OS cells treated with GTPBP8 siRNA #2 or expression of GTPBP8-myc. (**C**) Quantification of $^{35}$S intensity in panel B for all the mtDNA-encoded translation products relative to the corresponding intensity in coomassie staining.

**Supplemental Table S1.** High confidence interactors (HCIs) of GTPBP8 screened by BioID proximity protein purification and mass spectroscopy analysis. Interactions are assigned as HCIs based on peptide-spectrum match (PSM) of control samples and further refined by using CRAPome contaminant repository data (frequency cut-off ≥ 20% and average PSM change ≤ 3).

**Supplemental Table S2.** Label-free quantification (LFQ) intensities derived from MaxQuant for U2OS cells treated with control siRNA or GTPBP8 siRNA. N/A is short for non-applicable.

### Figure S1

A

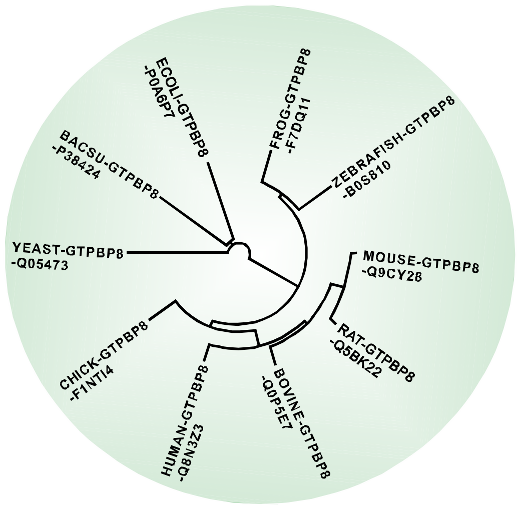

C

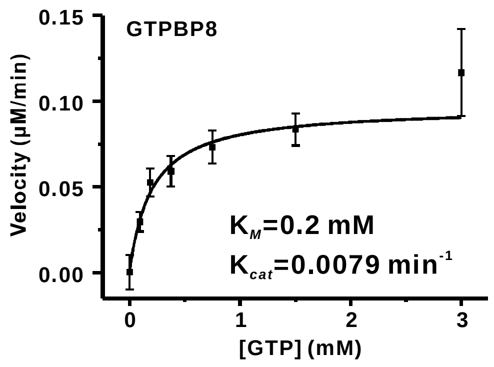

B

```
                                        1-46aa
GTPBP8_Human      MAAPGLRLGAGRLFEMPAVLERLSRYNSTSQAFAEVLRLPKQQLRKLLYPLQEVERFLAPYGRQDLHLRIFDPSPEDIAR 80
Ysxc_B.subtilis   -------------------------------------------------------------------------------- 0
Yiha_E.coli       -------------------------------------------------------------------------------- 0

                                                                   G1                           G2
GTPBP8_Human      ADNIFTATERNRIDYVSSAVRIDHAPDLPRPEVCFIGRSNVGKSSLIKALFSLAPEVEVRVSKKPGHTKKMNFFKVGKHF 160
Ysxc_B.subtilis   -------MKVTKSEIVISAVKPEQYPEGGLPEIALAGRSNVGKSSFINSLINR--KNLARTSSKPGKTQTLNFYIINDEL 71
Yiha_E.coli       ----MTNLNYQQTHFVMSAPDIRHLPSDTGIEVAFAGRSNAGKSSALNTLTNQ--KSLARTSKTPGRTQLINLFEVADGK 74
                         :  : . * **     : * .     *:.: ****.**** ::* .    :   .*.*..**:*: :*:: : .

                     G3                                                                   G4
GTPBP8_Human      TVVDMPGYGFRAP-----EDFVDMVETYLKERRNLKRTFLLVDSVVGIQKTDNIAIEMCEEFALPYVIVLTKIDKSSKGH 235
Ysxc_B.subtilis   HFVDVPGYGFAKVSKSEREAWGRMIETYITTREELKAVVQIVDLRHAPSNDDVQMYEFLKYYGIPVIVIATKADKIPKGK 151
Yiha_E.coli       RLVDLPGYGYAEVPEEMKRKWQRALGEYLEKRQSLQGLVVLMDIRHPLKDLDQQMIEWAVDSNIAVLVLLTKADKLASGA 154
                   .**:****:          . :    :   *:   *..*:  . ::*       .. *     *        :   ::: ** **   .*

                                            G5
GTPBP8_Human      LLKQVLQIQKFV-NMKTQGCFPQLFPVSAVTFSGIHLLRCFIASVTGSLD---------  284
Ysxc_B.subtilis   WDKHAKVVRQTL-NIDPE---DELILFSSETKKGKDEAWGAIKKMINR-----------  195
Yiha_E.coli       RKAQLNMVREAVLAFNGD---VQVETFSSLKKQGVDKLRQKLDTWFSEMQPVEETQDGE  210
                      :   ::: :  :. :     ::  .*: .  .* .      :  .   .
```

#### Figure S2

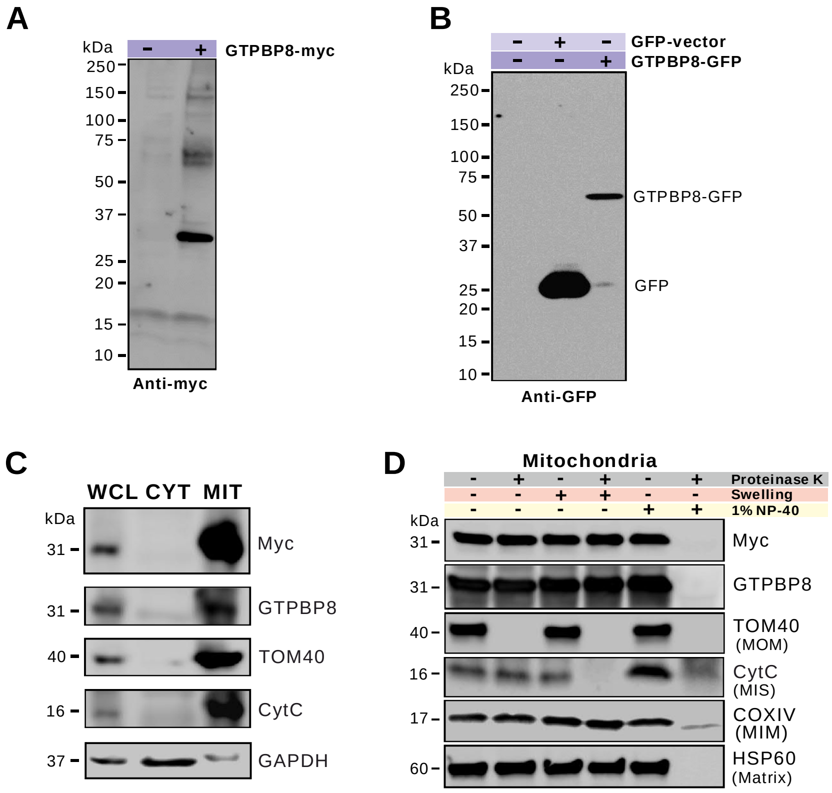
A
kDa
GTPBP8-myc
250
150
100
75
50
37
25
20
15
10
Anti-myc
B
GFP-vector
GTPBP8-GFP
kDa
250
150
100
75
50
37
25
20
15
10
GTPBP8-GFP
GFP
Anti-GFP
C
WCL CYT MIT
kDa
31
Myc
31
GTPBP8
40
TOM40
16
CytC
37
GAPDH
D
Mitochondria
Proteinase K
Swelling
1% NP-40
kDa
31
Myc
31
GTPBP8
40
TOM40
(MOM)
16
CytC
(MIS)
17
COXIV
(MIM)
60
HSP60
(Matrix)

### Figure S3

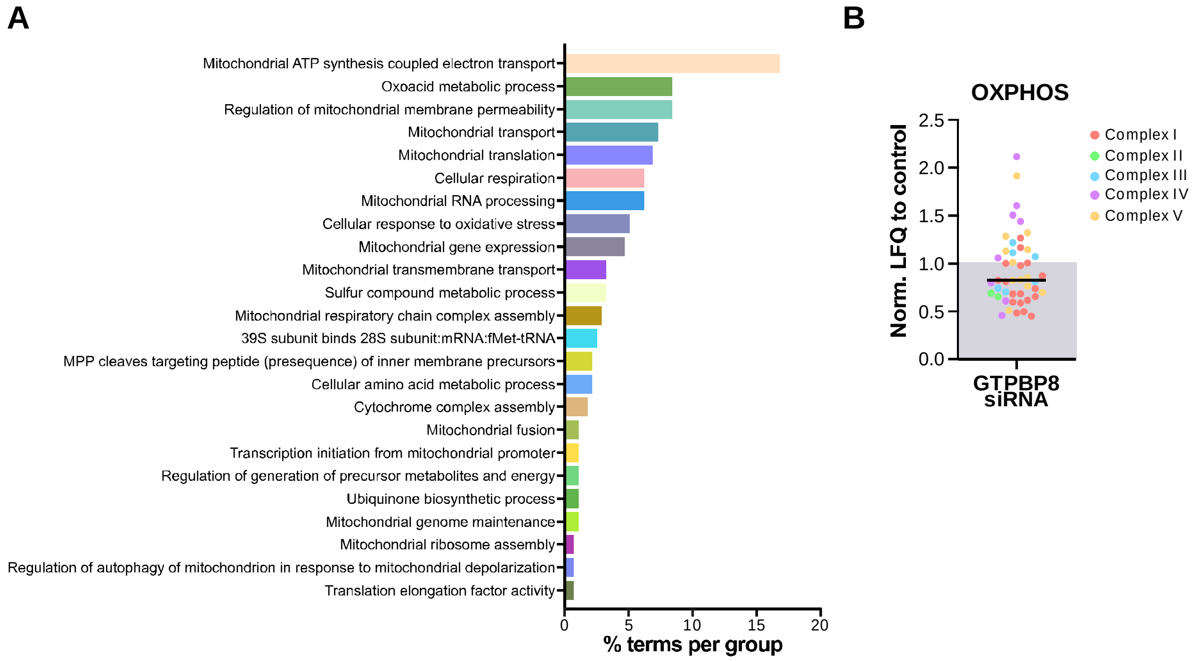
A
Mitochondrial ATP synthesis coupled electron transport
Oxoacid metabolic process
Regulation of mitochondrial membrane permeability
Mitochondrial transport
Mitochondrial translation
Cellular respiration
Mitochondrial RNA processing
Cellular response to oxidative stress
Mitochondrial gene expression
Mitochondrial transmembrane transport
Sulfur compound metabolic process
Mitochondrial respiratory chain complex assembly
39S subunit binds 28S subunit:mRNA:fMet-tRNA
MPP cleaves targeting peptide (presequence) of inner membrane precursors
Cellular amino acid metabolic process
Cytochrome complex assembly
Mitochondrial fusion
Transcription initiation from mitochondrial promoter
Regulation of generation of precursor metabolites and energy
Ubiquinone biosynthetic process
Mitochondrial genome maintenance
Mitochondrial ribosome assembly
Regulation of autophagy of mitochondrion in response to mitochondrial depolarization
Translation elongation factor activity
0
5
10
15
20
% terms per group
B
OXPHOS
Norm. LFQ to control
2.5
2.0
1.5
1.0
0.5
0.0
GTPBP8
siRNA
Complex I
Complex II
Complex III
Complex IV
Complex V

#### Figure S4

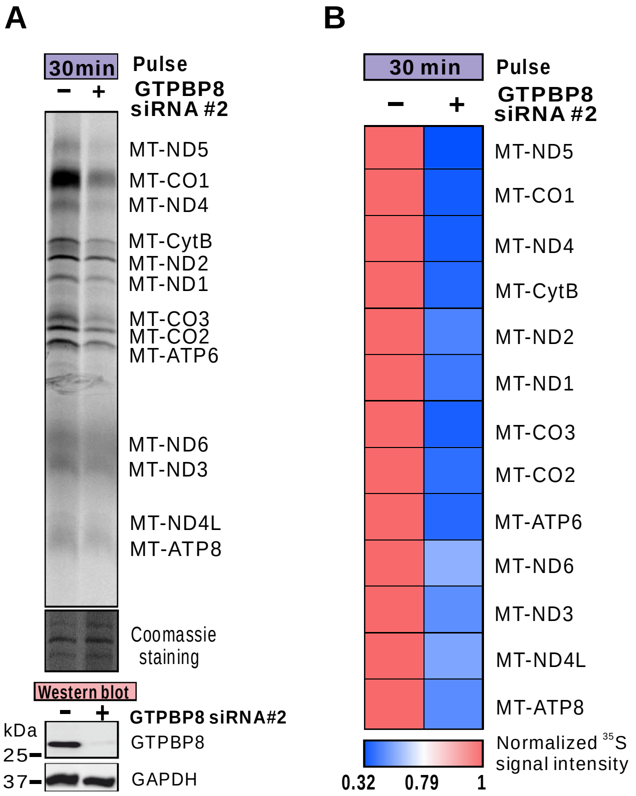

A
30min
Pulse
− +
GTPBP8 siRNA #2
MT-ND5
MT-CO1
MT-ND4
MT-CytB
MT-ND2
MT-ND1
MT-CO3
MT-CO2
MT-ATP6
MT-ND6
MT-ND3
MT-ND4L
MT-ATP8
Coomassie staining
Western blot
− +
GTPBP8 siRNA#2
kDa
25
37
GTPBP8
GAPDH
B
30 min
Pulse
− +
GTPBP8 siRNA #2
MT-ND5
MT-CO1
MT-ND4
MT-CytB
MT-ND2
MT-ND1
MT-CO3
MT-CO2
MT-ATP6
MT-ND6
MT-ND3
MT-ND4L
MT-ATP8
Normalized 35S signal intensity
0.32
0.79
1

### Figure S5

A

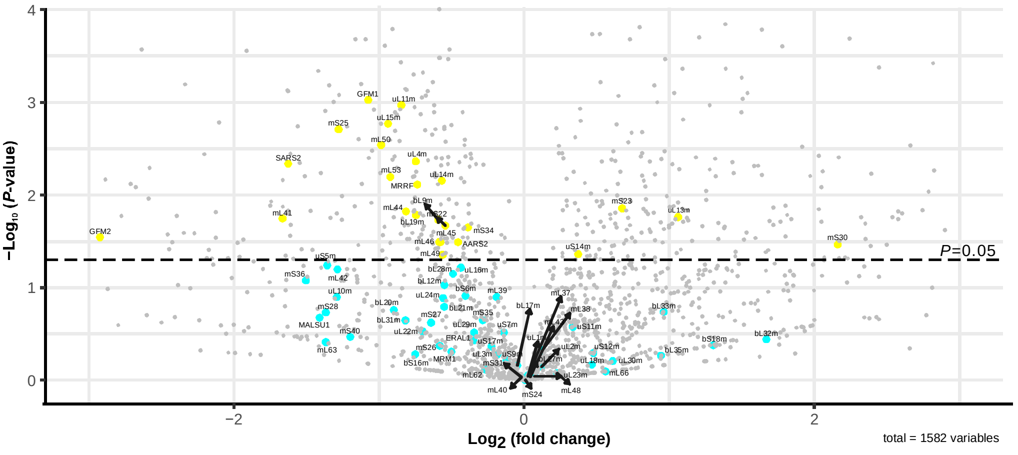

B

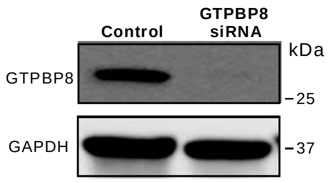

C

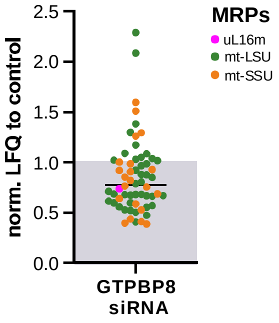

### Figure S6

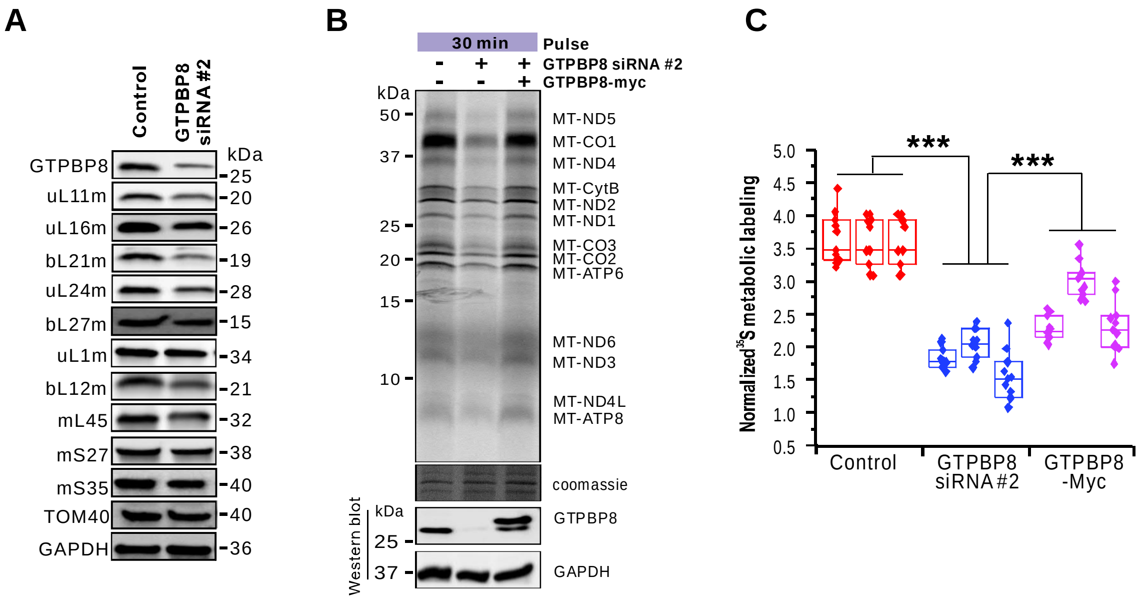
A
Control
GTPBP8 siRNA #2
kDa
GTPBP8 25
uL11m 20
uL16m 26
bL21m 19
uL24m 28
bL27m 15
uL1m 34
bL12m 21
mL45 32
mS27 38
mS35 40
TOM40 40
GAPDH 36
B
30 min
Pulse
- + + GTPBP8 siRNA #2
- - + GTPBP8-myc
kDa
50
37
25
20
15
10
MT-ND5
MT-CO1
MT-ND4
MT-CytB
MT-ND2
MT-ND1
MT-CO3
MT-CO2
MT-ATP6
MT-ND6
MT-ND3
MT-ND4L
MT-ATP8
coomassie
Western blot
kDa
25
GTPBP8
37
GAPDH
C
Normalized 35S metabolic labeling
5.0
4.5
4.0
3.5
3.0
2.5
2.0
1.5
1.0
0.5
***
***
Control
GTPBP8 siRNA #2
GTPBP8 -Myc

**Table S1: The high confidence interactions obtained from GTPBP8 BioID purification. The raw data was analyzed by Proteome Discoverer and the peptide-spectrum matching (PSM) vaules were used for identification of high confidence interactions (HCIs)**

| Bait_ID | Prey_ID | Gene names (primary ) | New MRP Name | Functional Group | Entry name | GTPBP8_Biological replicate 1 | GTPBP8_Biological replicate 2 | GTPBP8_Biological replicate 3 | GTPBP8_Biological replicate 4 | GTPBP8_psm_sum | GTPBP8_Frequency | Control_Biological replicate 1 | Control_Biological replicate 2 | Control_Biological replicate 3 | Control_Biological replicate 4 | Control_psm_sum | Control_Frequency |
|---|---|---|---|---|---|---|---|---|---|---|---|---|---|---|---|---|---|
| Q8N3Z3 | O95573 | ACSL3 | | Fatty acid metabolism | ACSL3_HUMAN | **7** | **2** | **3** | **3** | **15** | **4** | 0 | 0 | 0 | 0 | 0 | 0 |
| Q8N3Z3 | P30084 | ECHS1 | | Fatty acid metabolism | ECHM_HUMAN | **3** | **3** | **4** | **2** | **12** | **4** | 0 | 0 | 0 | 0 | 0 | 0 |
| Q8N3Z3 | P40939 | HADHA | | Fatty acid metabolism | ECHA_HUMAN | **10** | **12** | **13** | **10** | **45** | **4** | 0 | 0 | 1 | 1 | 2 | 2 |
| Q8N3Z3 | P49748 | ACADVL | | Fatty acid metabolism | ACADV_HUMAN | **8** | **8** | **9** | **7** | **32** | **4** | 0 | 1 | 1 | 0 | 2 | 2 |
| Q8N3Z3 | P49753 | ACOT2 | | Fatty acid metabolism | ACOT2_HUMAN | **57** | **53** | **62** | **55** | **227** | **4** | 12 | 0 | 0 | 0 | 12 | 1 |
| Q8N3Z3 | P55084 | HADHB | | Fatty acid metabolism | ECHB_HUMAN | **6** | **7** | **5** | **7** | **25** | **4** | 0 | 0 | 0 | 0 | 0 | 0 |
| Q8N3Z3 | Q7Z2W9 | MRPL21 | bL21m | Mitochondrial translation | RM21_HUMAN | **2** | **2** | **3** | **3** | **10** | **4** | 0 | 0 | 0 | 0 | 0 | 0 |
| Q8N3Z3 | Q7Z7F7 | MRPL55 | bL31m | Mitochondrial translation | RM55_HUMAN | **2** | **3** | **2** | **2** | **9** | **4** | 0 | 0 | 0 | 0 | 0 | 0 |
| Q8N3Z3 | Q8N983 | MRPL43 | mL43 | Mitochondrial translation | RM43_HUMAN | **3** | **3** | **2** | **3** | **11** | **4** | 0 | 0 | 0 | 0 | 0 | 0 |
| Q8N3Z3 | Q9BRJ2 | MRPL45 | mL45 | Mitochondrial translation | RM45_HUMAN | **2** | **3** | **2** | **2** | **9** | **4** | 0 | 0 | 0 | 0 | 0 | 0 |
| Q8N3Z3 | Q9BYD6 | MRPL1 | uL1m | Mitochondrial translation | RM01_HUMAN | **2** | **2** | **3** | **3** | **10** | **4** | 0 | 0 | 0 | 0 | 0 | 0 |
| Q8N3Z3 | Q9H2W6 | MRPL46 | mL46 | Mitochondrial translation | RM46_HUMAN | **5** | **5** | **5** | **4** | **19** | **4** | 0 | 0 | 0 | 0 | 0 | 0 |
| Q8N3Z3 | Q9H9J2 | MRPL44 | mL44 | Mitochondrial translation | RM44_HUMAN | **2** | **4** | **2** | **4** | **12** | **4** | 0 | 0 | 0 | 0 | 0 | 0 |
| Q8N3Z3 | Q9NQ50 | MRPL40 | mL40 | Mitochondrial translation | RM40_HUMAN | **4** | **2** | **4** | **2** | **12** | **4** | 1 | 0 | 1 | 0 | 2 | 2 |
| Q8N3Z3 | Q9NSE4 | IARS2 | | Mitochondrial translation | SYIM_HUMAN | **6** | **5** | **6** | **4** | **21** | **4** | 0 | 0 | 0 | 0 | 0 | 0 |
| Q8N3Z3 | Q9Y2Z4 | YARS2 | | Mitochondrial translation | SYYM_HUMAN | **1** | **3** | **2** | **2** | **8** | **4** | 0 | 0 | 0 | 0 | 0 | 0 |
| Q8N3Z3 | O75489 | NDUFS3 | | Respiratory electron transport and the citric acid (TCA) cycle | NDUS3_HUMAN | **5** | **5** | **3** | **3** | **16** | **4** | 0 | 0 | 0 | 0 | 0 | 0 |
| Q8N3Z3 | P07954 | FH | | Respiratory electron transport and the citric acid (TCA) cycle | FUMH_HUMAN | **6** | **2** | **5** | **2** | **15** | **4** | 0 | 0 | 0 | 0 | 0 | 0 |
| Q8N3Z3 | P10515 | DLAT | | Respiratory electron transport and the citric acid (TCA) cycle | ODP2_HUMAN | **15** | **17** | **16** | **16** | **64** | **4** | 2 | 2 | 4 | 3 | 11 | 4 |
| Q8N3Z3 | P10606 | COX5B | | Respiratory electron transport and the citric acid (TCA) cycle | COX5B_HUMAN | **2** | **2** | **2** | **2** | **8** | **4** | 0 | 0 | 0 | 0 | 0 | 0 |
| Q8N3Z3 | P13073 | COX4I1 | | Respiratory electron transport and the citric acid (TCA) cycle | COX41_HUMAN | **2** | **3** | **2** | **3** | **10** | **4** | 0 | 0 | 0 | 0 | 0 | 0 |
| Q8N3Z3 | P48047 | ATP5PO | | Respiratory electron transport and the citric acid (TCA) cycle | ATPO_HUMAN | **3** | **3** | **3** | **2** | **11** | **4** | 0 | 0 | 0 | 0 | 0 | 0 |
| Q8N3Z3 | P56134 | ATP5MF | | Respiratory electron transport and the citric acid (TCA) cycle | ATPK_HUMAN | **2** | **2** | **2** | **2** | **8** | **4** | 0 | 0 | 0 | 0 | 0 | 0 |
| Q8N3Z3 | Q16718 | NDUFA5 | | Respiratory electron transport and the citric acid (TCA) cycle | NDUA5_HUMAN | **2** | **3** | **1** | **2** | **8** | **4** | 0 | 0 | 0 | 0 | 0 | 0 |
| Q8N3Z3 | Q9BSH4 | TACO1 | | Respiratory electron transport and the citric acid (TCA) cycle | TACO1_HUMAN | **7** | **4** | **4** | **5** | **20** | **4** | 0 | 0 | 0 | 0 | 0 | 0 |
| Q8N3Z3 | O00429 | DNM1L | | | DNM1L_HUMAN | **15** | **10** | **14** | **12** | **51** | **4** | 2 | 3 | 3 | 4 | 12 | 4 |
| Q8N3Z3 | O43615 | TIMM44 | | | TIM44_HUMAN | **3** | **3** | **4** | **3** | **13** | **4** | 0 | 0 | 0 | 0 | 0 | 0 |
| Q8N3Z3 | O76031 | CLPX | | | CLPX_HUMAN | **4** | **2** | **2** | **3** | **11** | **4** | 0 | 0 | 0 | 0 | 0 | 0 |
| Q8N3Z3 | O94925 | GLS | | | GLSK_HUMAN | **10** | **11** | **11** | **10** | **42** | **4** | 0 | 1 | 0 | 0 | 1 | 1 |
| Q8N3Z3 | O95202 | LETM1 | | | LETM1_HUMAN | **2** | **4** | **4** | **2** | **12** | **4** | 0 | 0 | 0 | 0 | 0 | 0 |
| Q8N3Z3 | P32322 | PYCR1 | | | P5CR1_HUMAN | **20** | **17** | **19** | **18** | **74** | **4** | 1 | 0 | 2 | 1 | 4 | 3 |
| Q8N3Z3 | Q12849 | GRSF1 | | | GRSF1_HUMAN | **15** | **14** | **12** | **15** | **56** | **4** | 1 | 1 | 0 | 2 | 4 | 3 |
| Q8N3Z3 | Q8N3Z3 | GTPBP8 | | | GTPB8_HUMAN | **59** | **57** | **59** | **59** | **234** | **4** | 0 | 0 | 0 | 0 | 0 | 0 |
| Q8N3Z3 | Q96C36 | PYCR2 | | | P5CR2_HUMAN | **4** | **0** | **4** | **5** | **13** | **3** | 0 | 0 | 0 | 0 | 0 | 0 |
| Q8N3Z3 | Q9BVS5 | TRMT61B | | | TR61B_HUMAN | **3** | **5** | **9** | **6** | **23** | **4** | 0 | 0 | 0 | 0 | 0 | 0 |
| Q8N3Z3 | Q9BX68 | HINT2 | | | HINT2_HUMAN | **3** | **4** | **4** | **3** | **14** | **4** | 0 | 0 | 0 | 0 | 0 | 0 |
| Q8N3Z3 | Q9HAV7 | GRPEL1 | | | GRPE1_HUMAN | **3** | **3** | **5** | **2** | **13** | **4** | 0 | 1 | 1 | 1 | 3 | 3 |
| Q8N3Z3 | Q9Y4W6 | AFG3L2 | | | AFG32_HUMAN | **11** | **8** | **9** | **8** | **36** | **4** | 0 | 0 | 0 | 0 | 0 | 0 |

**Table S2: Proteomics analysis of siRNA-mediated depletion of GTPBP8. The data were processed by Maxquant using relative label-free quantification (LFQ)**

| Majority protein ID | Gene names | New MRP Name | LFQ intensity Control 1 | LFQ intensity Control 2 | LFQ intensity Control 3 | LFQ intensity siRNA-GTPBP8_1 | LFQ intensity siRNA-GTPBP8_2 | LFQ intensity siRNA-GTPBP8_3 | Average_Control | Average_siRNA-GTPBP8 | p-value | Fold_change (siRNA-GTPBP8/Control) | Log2(fold change) |
|---|---|---|---|---|---|---|---|---|---|---|---|---|---|
| Q9BYD3 | MRPL4 | uL4m | 1221400000 | 1244000000 | 1170600000 | 566980000 | 759910000 | 840670000 | 1212000000 | 722520000 | 0.004328022 | 0.596138614 | -0.74628027 |
| Q9BYD2 | MRPL9 | bL9m | 301680000 | 301850000 | 231920000 | 196820000 | 186180000 | 171250000 | 278483333.3 | 184750000 | 0.018520594 | 0.663414926 | -0.592016623 |
| Q9Y3B7 | MRPL11 | uL11m | 1237000000 | 1100700000 | 1170400000 | 573140000 | 735480000 | 642800000 | 1169366667 | 650473333.3 | 0.001068258 | 0.556261224 | -0.846165553 |
| Q9BYD1 | MRPL13 | uL13m | 203760000 | 254360000 | 331790000 | 454250000 | 525980000 | 668370000 | 263303333.3 | 549533333.3 | 0.017320783 | 2.087073211 | 1.061481209 |
| Q6P1L8 | MRPL14 | uL14m | 569000000 | 577150000 | 597550000 | 435170000 | 421140000 | 320560000 | 581233333.3 | 392290000 | 0.00700306 | 0.67492688 | -0.567196883 |
| Q9P015 | MRPL15 | uL15m | 2056900000 | 1809800000 | 1907100000 | 833930000 | 1000400000 | 1179100000 | 1924600000 | 1004476667 | 0.00170071 | 0.52191451 | -0.938114582 |
| P49406 | MRPL19 | bL19m | 690070000 | 945940000 | 771960000 | 419680000 | 495270000 | 521960000 | 802656666.7 | 478970000 | 0.01646398 | 0.596730856 | -0.744847716 |
| Q8IXM3 | MRPL41 | mL41 | 389490000 | 376700000 | 298830000 | 164320000 | 171610000 | 0 | 355006666.7 | 111976666.7 | 0.017978949 | 0.315421307 | -1.664647978 |
| Q9H9J2 | MRPL44 | mL44 | 914900000 | 761090000 | 724160000 | 333450000 | 507420000 | 524270000 | 800050000 | 455046666.7 | 0.015031472 | 0.568772785 | -0.814075659 |
| Q9BRJ2 | MRPL45 | mL45 | 340260000 | 384600000 | 389830000 | 193930000 | 236380000 | 313460000 | 371563333.3 | 247923333.3 | 0.032173898 | 0.667243808 | -0.583714083 |
| Q9H2W6 | MRPL46 | mL46 | 764050000 | 544010000 | 647540000 | 463300000 | 396750000 | 451140000 | 651866666.7 | 437063333.3 | 0.03236917 | 0.670479648 | -0.576734554 |
| Q13405 | MRPL49 | mL49 | 1614300000 | 1919300000 | 2357700000 | 1290200000 | 1397600000 | 1305600000 | 1963766667 | 1331133333 | 0.044229836 | 0.677846995 | -0.560968433 |
| Q8N5N7 | MRPL50 | mL50 | 840890000 | 787160000 | 707220000 | 368180000 | 331300000 | 480320000 | 778423333.3 | 393266666.7 | 0.002897387 | 0.505209248 | -0.985047047 |
| Q96EL3 | MRPL53 | mL53 | 490670000 | 474820000 | 414310000 | 213580000 | 311340000 | 203200000 | 459933333.3 | 242706666.7 | 0.006402104 | 0.527699667 | -0.922211023 |
| Q6P161 | MRPL54 | mL54 | 48283000 | 102660000 | 99356000 | 0 | 0 | 0 | 83433000 | 0 | 0.009035338 | 0 | N/A |
| O60783 | MRPS14 | uS14m | 231980000 | 321570000 | 311630000 | 385400000 | 373210000 | 361700000 | 288393333.3 | 373436666.7 | 0.043424842 | 1.294886613 | 0.372825774 |
| P82650 | MRPS22 | mS22 | 682610000 | 649730000 | 763700000 | 480900000 | 564240000 | 393620000 | 698680000 | 479586666.7 | 0.021482744 | 0.686418198 | -0.542840293 |
| Q9Y3D9 | MRPS23 | mS23 | 818430000 | 778220000 | 643480000 | 1090400000 | 1109100000 | 1377200000 | 746710000 | 1192233333 | 0.013961503 | 1.596648409 | 0.675046658 |
| P82663 | MRPS25 | mS25 | 557260000 | 456520000 | 452480000 | 165350000 | 203650000 | 235440000 | 488753333.3 | 201480000 | 0.001956502 | 0.412232483 | -1.278469906 |
| Q9NP92 | MRPS30 | mS30 | 0 | 82371000 | 0 | 115500000 | 105660000 | 147240000 | 27457000 | 122800000 | 0.034237845 | 4.472447828 | 2.161064653 |
| P82930 | MRPS34 | mS34 | 582910000 | 633270000 | 560950000 | 446590000 | 401260000 | 511570000 | 592376666.7 | 453140000 | 0.022444316 | 0.76495248 | -0.386557968 |
| Q96E11 | MRRF | | 194170000 | 182330000 | 222610000 | 136520000 | 123440000 | 99606000 | 199703333.3 | 119855333.3 | 0.007730743 | 0.600166914 | -0.736564306 |
| Q5JTZ9 | AARS2 | | 317730000 | 380700000 | 366390000 | 216830000 | 295750000 | 264240000 | 354940000 | 258940000 | 0.032303242 | 0.729531752 | -0.454957324 |
| Q96RP9 | GFM1 | | 848400000 | 796420000 | 810510000 | 459330000 | 405400000 | 300520000 | 818443333.3 | 388416666.7 | 0.000939749 | 0.474579792 | -1.075277425 |
| Q969S9 | GFM2 | | 169210000 | 103840000 | 81366000 | 46736000 | 0 | 0 | 118138666.7 | 15578666.67 | 0.028548997 | 0.131867636 | -2.922837567 |
| Q9NP81 | SARS2 | | 902320000 | 694570000 | 629120000 | 187300000 | 244000000 | 290300000 | 742003333.3 | 240533333.3 | 0.004604467 | 0.324167457 | -1.62518883 |
| Q9BYD6 | MRPL1 | uL1m | 1073000000 | 929590000 | 1040400000 | 864540000 | 931340000 | 1303300000 | 1014330000 | 1033060000 | 0.902267104 | 1.018465391 | 0.026396956 |
| Q5T653 | MRPL2 | uL2m | 293480000 | 301280000 | 224920000 | 395420000 | 287810000 | 207310000 | 273226666.7 | 296846666.7 | 0.712317747 | 1.08644837 | 0.119619617 |
| P09001 | MRPL3 | uL3m | 671180000 | 614510000 | 723970000 | 737100000 | 660110000 | 372780000 | 669886666.7 | 589996666.7 | 0.526425981 | 0.880741021 | -0.183210233 |
| Q7Z7H8 | MRPL10 | uL10m | 552970000 | 186290000 | 362910000 | 212690000 | 158650000 | 79051000 | 367390000 | 150130333.3 | 0.126300114 | 0.408640228 | -1.29109686 |
| P52815 | MRPL12 | bL12m | 5001100000 | 3828300000 | 4883400000 | 3073300000 | 4093000000 | 2206900000 | 4570933333 | 3124400000 | 0.093652099 | 0.68353655 | -0.548909612 |
| Q9NX20 | MRPL16 | uL16m | 134310000 | 169170000 | 135220000 | 126120000 | 95485000 | 102670000 | 146233333.3 | 108091666.7 | 0.060790379 | 0.739172555 | -0.436016903 |
| Q9NRX2 | MRPL17 | bL17m | 433920000 | 370280000 | 369210000 | 340760000 | 402690000 | 393060000 | 391136666.7 | 378836666.7 | 0.691034817 | 0.968553191 | -0.046096814 |
| Q9H0U6 | MRPL18 | uL18m | 154500000 | 0 | 173910000 | 244190000 | 209860000 | 0 | 109470000 | 151350000 | 0.679257366 | 1.382570567 | 0.467353118 |
| Q9BYC9 | MRPL20 | bL20m | 200570000 | 164530000 | 192990000 | 130900000 | 168480000 | 0 | 186030000 | 99793333.33 | 0.174047848 | 0.536436775 | -0.89851995 |
| Q7Z2W9 | MRPL21 | bL21m | 359590000 | 275590000 | 204880000 | 138910000 | 215450000 | 218840000 | 280020000 | 191066666.7 | 0.160905635 | 0.682332214 | -0.551453764 |
| Q9NWU5 | MRPL22 | uL22m | 84671000 | 109920000 | 115630000 | 97380000 | 93406000 | 0 | 103407000 | 63595333.33 | 0.296770179 | 0.615000274 | -0.701341042 |
| Q16540 | MRPL23 | uL23m | 445490000 | 535820000 | 511940000 | 272900000 | 387810000 | 909430000 | 497750000 | 523380000 | 0.903113241 | 1.051491713 | 0.07243748 |
| Q96A35 | MRPL24 | uL24m | 279200000 | 351710000 | 481510000 | 230140000 | 292180000 | 233200000 | 370806666.7 | 251840000 | 0.129839726 | 0.67916794 | -0.558159736 |
| Q9P0M9 | MRPL27 | bL27m | 211720000 | 163500000 | 222360000 | 287220000 | 206180000 | 121650000 | 199193333.3 | 205016666.7 | 0.914785425 | 1.029234579 | 0.041571834 |
| Q13084 | MRPL28 | bL28m | 1272900000 | 1029200000 | 914890000 | 773910000 | 878590000 | 638110000 | 1072330000 | 763536666.7 | 0.071068177 | 0.712035163 | -0.489979606 |
| Q8TCC3 | MRPL30 | uL30m | 0 | 33038000 | 49729000 | 77887000 | 48520000 | 0 | 27589000 | 42135666.67 | 0.618686343 | 1.527263281 | 0.610948786 |
| Q9BYC8 | MRPL32 | bL32m | 65201000 | 0 | 0 | 140120000 | 67181000 | 0 | 21733666.67 | 69100333.33 | 0.360665045 | 3.179414426 | 1.668761079 |
| O75394 | MRPL33 | bL33m | 73163000 | 0 | 107990000 | 142180000 | 122410000 | 88178000 | 60384333.33 | 117589333.3 | 0.182549826 | 1.947348374 | 0.961511 |
| Q9NZE8 | MRPL35 | bL35m | 0 | 112440000 | 0 | 111650000 | 104540000 | 0 | 37480000 | 72063333.33 | 0.54262392 | 1.922714337 | 0.943144433 |
| Q9BZE1 | MRPL37 | mL37 | 660990000 | 579870000 | 857640000 | 704220000 | 658610000 | 780970000 | 699500000 | 714600000 | 0.874708227 | 1.021586848 | 0.030811857 |
| Q96DV4 | MRPL38 | mL38 | 250170000 | 198760000 | 199310000 | 250240000 | 199850000 | 257010000 | 216080000 | 235700000 | 0.473350749 | 1.090799704 | 0.125386214 |
| Q9NYK5 | MRPL39 | mL39 | 464970000 | 508850000 | 437570000 | 369110000 | 430020000 | 438280000 | 470463333.3 | 412470000 | 0.126395232 | 0.876731449 | -0.189793095 |
| Q9NQ50 | MRPL40 | mL40 | 525010000 | 446470000 | 514070000 | 381520000 | 553520000 | 531930000 | 495183333.3 | 488990000 | 0.921990659 | 0.987492848 | -0.018157796 |
| Q9Y6G3 | MRPL42 | mL42 | 630490000 | 448430000 | 527370000 | 289180000 | 369760000 | 0 | 535430000 | 219646666.7 | 0.063557463 | 0.410224804 | -1.28551337 |
| Q8N983 | MRPL43 | mL43 | 744820000 | 943170000 | 1148900000 | 1076800000 | 1149000000 | 717720000 | 945630000 | 981173333.3 | 0.850798074 | 1.037586935 | 0.053232219 |

| | | | | | | | | | | | | | |
|---|---|---|---|---|---|---|---|---|---|---|---|---|---|
| Q9HD33 | MRPL47 | uL29m | 355490000 | 367860000 | 426450000 | 215330000 | 259810000 | 430380000 | 383266666.7 | 301840000 | 0.303940263 | 0.78754566 | -0.344564525 |
| Q96GC5 | MRPL48 | mL48 | 254610000 | 0 | 291870000 | 304510000 | 335450000 | 0 | 182160000 | 213320000 | 0.835865285 | 1.17105841 | 0.227813037 |
| Q7Z7F7 | MRPL55 | bL31m | 183800000 | 180090000 | 131530000 | 0 | 148370000 | 132730000 | 165140000 | 93700000 | 0.226176647 | 0.56739736 | -0.817568657 |
| Q9BQC6 | MRPL57 | mL63 | 349220000 | 341170000 | 0 | 0 | 0 | 267660000 | 230130000 | 89220000 | 0.388016587 | 0.387693912 | -1.367010014 |
| Q14197 | MRPL58 | mL62 | 0 | 611560000 | 425640000 | 272490000 | 300100000 | 289530000 | 345733333.3 | 287373333.3 | 0.763507207 | 0.831199383 | -0.266733512 |
| Q9Y399 | MRPS2 | uS2m | 51786000 | 0 | 16587000 | 0 | 0 | 0 | 22791000 | 0 | 0.209789008 | 0 | N/A |
| P82675 | MRPS5 | uS5m | 118960000 | 158080000 | 167400000 | 0 | 105150000 | 68430000 | 148146666.7 | 57860000 | 0.057574024 | 0.390558906 | -1.356387937 |
| P82932 | MRPS6 | bS6m | 310710000 | 326440000 | 374640000 | 228520000 | 206880000 | 328900000 | 337263333.3 | 254766666.7 | 0.122421705 | 0.755393906 | -0.40469895 |
| Q9Y2R9 | MRPS7 | uS7m | 968460000 | 1213900000 | 1097600000 | 1055200000 | 1023800000 | 899960000 | 1093320000 | 992986666.7 | 0.304554415 | 0.908230588 | -0.138869469 |
| P82933 | MRPS9 | uS9m | 896350000 | 886050000 | 721050000 | 971240000 | 727430000 | 603990000 | 834483333.3 | 767553333.3 | 0.612267228 | 0.919794683 | -0.120616236 |
| P82664 | MRPS10 | uS10m | 0 | 0 | 0 | 55998000 | 53002000 | 0 | 0 | 36333333.33 | 0.116416934 | N/A | N/A |
| P82912 | MRPS11 | uS11m | 180180000 | 215760000 | 226160000 | 293300000 | 308860000 | 182920000 | 207366666.7 | 261693333.3 | 0.265628181 | 1.261983604 | 0.335693167 |
| O15235 | MRPS12 | uS12m | 0 | 89686000 | 137070000 | 95115000 | 118090000 | 102330000 | 75585333.33 | 105178333.3 | 0.508027898 | 1.391517755 | 0.476659317 |
| Q9Y3D3 | MRPS16 | bS16m | 313750000 | 329380000 | 0 | 199350000 | 182970000 | 0 | 214376666.7 | 127440000 | 0.524625625 | 0.594467681 | -0.750329716 |
| Q9Y2R5 | MRPS17 | uS17m | 858950000 | 673840000 | 748840000 | 477180000 | 611910000 | 863280000 | 760543333.3 | 650790000 | 0.430390566 | 0.855690888 | -0.224838368 |
| Q9NVS2 | MRPS18A | mL66 | 0 | 0 | 37370000 | 55158000 | 0 | 0 | 12456666.67 | 18386000 | 0.802679574 | 1.475996789 | 0.561689583 |
| Q9Y676 | MRPS18B | mS40 | 581250000 | 246670000 | 95498000 | 89624000 | 38419000 | 274280000 | 307806000 | 134107666.7 | 0.339762362 | 0.43568893 | -1.198629639 |
| Q9Y3D5 | MRPS18C | bS18m | 101910000 | 0 | 0 | 0 | 147490000 | 104060000 | 33970000 | 83850000 | 0.418813081 | 2.46835443 | 1.303549566 |
| Q96EL2 | MRPS24 | mS24 | 27924000 | 34500000 | 35927000 | 51289000 | 47624000 | 0 | 32783666.67 | 32971000 | 0.991588176 | 1.005714228 | 0.008220423 |
| Q9BYN8 | MRPS26 | mS26 | 81306000 | 74888000 | 124140000 | 97460000 | 89872000 | 0 | 93444666.67 | 62444000 | 0.424685468 | 0.668245735 | -0.581549369 |
| Q92552 | MRPS27 | mS27 | 348430000 | 171940000 | 458840000 | 184090000 | 210750000 | 232910000 | 326403333.3 | 209250000 | 0.238963169 | 0.641078012 | -0.641428168 |
| Q9Y2Q9 | MRPS28 | mS28 | 518380000 | 546800000 | 173090000 | 0 | 129320000 | 350840000 | 412756666.7 | 160053333.3 | 0.184681158 | 0.387766804 | -1.366738794 |
| Q92665 | MRPS31 | mS31 | 517530000 | 800300000 | 659730000 | 703360000 | 662420000 | 583800000 | 659186666.7 | 649860000 | 0.921450257 | 0.985851251 | -0.020558111 |
| P82673 | MRPS35 | mS35 | 406820000 | 504580000 | 380240000 | 387470000 | 395710000 | 276860000 | 430546666.7 | 353346666.7 | 0.224815408 | 0.820693072 | -0.28508532 |
| P82909 | MRPS36 | mS36 | 220070000 | 240470000 | 422160000 | 171170000 | 139980000 | 0 | 294233333.3 | 103716666.7 | 0.083473865 | 0.352498017 | -1.504312951 |
| A4D1E9 | GTPBP10 | | 0 | 0 | 0 | 70904000 | 47299000 | 0 | 0 | 39401000 | 0.131731208 | N/A | N/A |
| Q96EH3 | MALSU1 | | 272760000 | 96853000 | 124990000 | 142910000 | 43343000 | 0 | 164867666.7 | 62084333.33 | 0.21075968 | 0.376570704 | -1.409007327 |
| Q6IN84 | MRM1 | | 214640000 | 397100000 | 313280000 | 70764000 | 154390000 | 428130000 | 308340000 | 217761333.3 | 0.492740697 | 0.706237703 | -0.501774253 |
| Q9HC36 | MRM3 | | 115680000 | 48854000 | 153570000 | 0 | 0 | 0 | 106034666.7 | 0 | 0.025725708 | 0 | N/A |
| O75616 | ERAL1 | | 185710000 | 139490000 | 160130000 | 104990000 | 86479000 | 189790000 | 161776666.7 | 127086333.3 | 0.371528871 | 0.785566522 | -0.348194647 |
| Q9NYY8 | FASTKD2 | | 58120000 | 0 | 92643000 | 115940000 | 54180000 | 0 | 50254333.33 | 56706666.67 | 0.888087921 | 1.128393571 | 0.174270352 |
| Q96E29 | MTERF3 | | 46495000 | 141980000 | 0 | 0 | 0 | 0 | 62825000 | 0 | 0.207191311 | 0 | N/A |
| Q9BSH4 | TACO1 | | 786100000 | 726350000 | 709350000 | 552650000 | 672970000 | 578670000 | 740600000 | 601430000 | 0.032533069 | 0.812084796 | -0.300297717 |
| Q9BVS5 | TRMT61B | | 147300000 | 106460000 | 145920000 | 54082000 | 0 | 0 | 133226666.7 | 18027333.33 | 0.006838137 | 0.135313251 | -2.885624972 |
| Q9HA77 | CARS2 | | 0 | 63087000 | 0 | 0 | 56116000 | 0 | 21029000 | 18705333.33 | 0.938166089 | 0.889501799 | -0.168930571 |
| Q96SZ6 | CDK5RAP1 | | 181680000 | 122240000 | 0 | 0 | 0 | 0 | 101306666.7 | 0 | 0.131103414 | 0 | N/A |
| Q6PI48 | DARS2 | | 683390000 | 715360000 | 718970000 | 547800000 | 565840000 | 760410000 | 705906666.7 | 624683333.3 | 0.304370283 | 0.884937574 | -0.176352407 |
| Q5JPH6 | EARS2 | | 0 | 0 | 40572000 | 0 | 0 | 48503000 | 13524000 | 16167666.67 | 0.906240785 | 1.195479641 | 0.257589561 |
| Q969Y2 | GTPBP3 | | 38668000 | 23395000 | 0 | 0 | 9575500 | 56273000 | 20687666.67 | 21949500 | 0.954322619 | 1.060994473 | 0.085417141 |
| Q8N442 | GUF1 | | 58279000 | 173100000 | 65877000 | 142170000 | 127920000 | 0 | 99085333.33 | 90030000 | 0.884404525 | 0.90861076 | -0.138265705 |
| P49590 | HARS2 | | 308180000 | 156050000 | 272450000 | 248860000 | 151750000 | 160430000 | 245560000 | 187013333.3 | 0.350395676 | 0.761578976 | -0.392934443 |
| Q15031 | LARS2 | | 345630000 | 280610000 | 755310000 | 253450000 | 262150000 | 0 | 460516666.7 | 171866666.7 | 0.167964721 | 0.37320401 | -1.421963607 |
| Q96GW9 | MARS2 | | 318540000 | 247860000 | 217380000 | 135440000 | 242130000 | 281550000 | 261260000 | 219706666.7 | 0.476365879 | 0.840950267 | -0.249907612 |
| P46199 | MTIF2 | | 109670000 | 93887000 | 98694000 | 0 | 0 | 0 | 100750333.3 | 0 | 2.73217E-05 | 0 | N/A |
| Q96I59 | NARS2 | | 148470000 | 127430000 | 83803000 | 66713000 | 101720000 | 120580000 | 119901000 | 96337666.67 | 0.394681192 | 0.803476757 | -0.315671804 |
| Q7L3T8 | PARS2 | | 0 | 0 | 0 | 20620000 | 17433000 | 0 | 0 | 12684333.33 | 0.118899056 | N/A | N/A |
| Q9H0R6 | QRSL1 | | 69998000 | 63972000 | 0 | 0 | 41116000 | 0 | 44656666.67 | 13705333.33 | 0.303811006 | 0.306904531 | -1.70413815 |
| Q8N0V3 | RBFA | | 77521000 | 0 | 149450000 | 60846000 | 12877000 | 84770000 | 75657000 | 52831000 | 0.659537691 | 0.698296258 | -0.518088853 |
| Q9BW92 | TARS2 | | 316610000 | 258310000 | 298330000 | 346570000 | 613260000 | 530680000 | 291083333.3 | 496836666.7 | 0.063294759 | 1.706853707 | 0.771339412 |
| Q7L0Y3 | TRMT10C | | 609870000 | 571420000 | 574930000 | 360900000 | 496680000 | 531180000 | 585406666.7 | 462920000 | 0.083521412 | 0.790766533 | -0.338676281 |
| P43897 | TSFM | | 864190000 | 747260000 | 753540000 | 632830000 | 563110000 | 914380000 | 788330000 | 703440000 | 0.497464716 | 0.892316669 | -0.164372304 |
| P49411 | TUFM | | 8798400000 | 9533400000 | 7836700000 | 10559000000 | 9557400000 | 7295000000 | 8722833333 | 9137133333 | 0.721557611 | 1.047496035 | 0.066944783 |
| Q5ST30 | VARS2 | | 208500000 | 179640000 | 234170000 | 207300000 | 242440000 | 141210000 | 207436666.7 | 196983333.3 | 0.771224442 | 0.949607109 | -0.074597359 |
| Q9Y2Z4 | YARS2 | | 1439900000 | 1854300000 | 1423500000 | 1094600000 | 1248300000 | 1670800000 | 1572566667 | 1337900000 | 0.351207963 | 0.850774742 | -0.233150892 |
| Q9UDR5 | AASS | | 0 | 0 | 0 | 56920000 | 101220000 | 0 | 0 | 52713333.33 | 0.146344196 | N/A | N/A |
| P80404 | ABAT | | 0 | 0 | 122840000 | 55841000 | 82771000 | 85445000 | 40946666.67 | 74685666.67 | 0.467052392 | 1.823974275 | 0.867085383 |
| Q9NRK6 | ABCB10 | | 108600000 | 45671000 | 48813000 | 53176000 | 38319000 | 0 | 67694666.67 | 30498333.33 | 0.224094027 | 0.45052786 | -1.150311771 |

| | | | | | | | | | | | | | |
|---|---|---|---|---|---|---|---|---|---|---|---|---|---|
| O75027 | ABCB7 | | 135340000 | 140360000 | 134300000 | 167340000 | 181670000 | 47261000 | 136666666.7 | 132090333.3 | 0.919730696 | 0.966514634 | -0.049136518 |
| Q9NUJ1 | ABHD10 | | 130020000 | 156930000 | 194110000 | 142060000 | 127430000 | 141600000 | 160353333.3 | 137030000 | 0.291059736 | 0.854550368 | -0.226762567 |
| P42765 | ACAA2 | | 2054300000 | 2106600000 | 2005000000 | 2069700000 | 1711300000 | 1660200000 | 2055300000 | 1813733333 | 0.14151012 | 0.882466469 | -0.180386634 |
| Q9H845 | ACAD9 | | 703850000 | 637860000 | 744580000 | 634920000 | 553530000 | 605910000 | 695430000 | 598120000 | 0.067881742 | 0.860072186 | -0.217470345 |
| P11310 | ACADM | | 2851600000 | 3254600000 | 4051800000 | 3464000000 | 4167500000 | 2954100000 | 3386000000 | 3528533333 | 0.788964094 | 1.042094901 | 0.059486666 |
| P16219 | ACADS | | 491180000 | 240450000 | 381450000 | 152530000 | 260980000 | 172840000 | 371026666.7 | 195450000 | 0.092735203 | 0.526781543 | -0.924723296 |
| P45954 | ACADSB | | 225330000 | 161080000 | 188090000 | 105870000 | 186900000 | 0 | 191500000 | 97590000 | 0.176144311 | 0.509608355 | -0.972539164 |
| P49748 | ACADVL | | 4016100000 | 3359300000 | 3560300000 | 2943500000 | 3111900000 | 3592200000 | 3645233333 | 3215866667 | 0.19322764 | 0.882211473 | -0.180803573 |
| P24752 | ACAT1 | | 4453400000 | 4306100000 | 4534000000 | 3698100000 | 3553400000 | 2944100000 | 4431166667 | 3398533333 | 0.01270285 | 0.766961297 | -0.382774318 |
| Q9Y305 | ACOT9 | | 61697000 | 75601000 | 73970000 | 0 | 93678000 | 0 | 70422666.67 | 31226000 | 0.28173652 | 0.443408372 | -1.173292084 |
| Q96CM8 | ACSF2 | | 269830000 | 277490000 | 232090000 | 111850000 | 166570000 | 146060000 | 259803333.3 | 141493333.3 | 0.005099772 | 0.544617082 | -0.876685861 |
| Q4G176 | ACSF3 | | 0 | 0 | 0 | 14880000 | 0 | 89700000 | 0 | 34860000 | 0.277467671 | N/A | N/A |
| Q9H6R3 | ACSS3 | | 0 | 0 | 0 | 29632000 | 67900000 | 0 | 0 | 32510666.67 | 0.173436757 | N/A | N/A |
| Q53H12 | AGK | | 116130000 | 82546000 | 0 | 102500000 | 25704000 | 108150000 | 66225333.33 | 78784666.67 | 0.787423943 | 1.189645453 | 0.250531675 |
| P54819 | AK2 | | 1775200000 | 1662600000 | 1114700000 | 1390300000 | 1196700000 | 1614500000 | 1517500000 | 1400500000 | 0.647479149 | 0.922899506 | -0.115754533 |
| Q9UIJ7 | AK3 | | 661070000 | 498690000 | 441250000 | 296060000 | 455790000 | 437570000 | 533670000 | 396473333.3 | 0.173484489 | 0.742918533 | -0.428724079 |
| P27144 | AK4 | | 808650000 | 721600000 | 997570000 | 618380000 | 703200000 | 624440000 | 842606666.7 | 648673333.3 | 0.086940402 | 0.769841207 | -0.377367199 |
| Q92667 | AKAP1 | | 138150000 | 125370000 | 0 | 172920000 | 186280000 | 108160000 | 87840000 | 155786666.7 | 0.247672379 | 1.773527626 | 0.826621803 |
| P30837 | ALDH1B1 | | 3289500000 | 3096600000 | 3384800000 | 1513900000 | 1670900000 | 1844800000 | 3256966667 | 1676533333 | 0.000245321 | 0.514752991 | -0.958047787 |
| P05091 | ALDH2 | | 1676800000 | 1489600000 | 1688600000 | 0 | 0 | 39348000 | 1618333333 | 13116000 | 1.67295E-05 | 0.008104634 | -6.947037178 |
| P30038 | ALDH4A1 | | 228930000 | 176370000 | 201460000 | 206950000 | 205230000 | 175090000 | 202253333.3 | 195756666.7 | 0.741426849 | 0.967878568 | -0.047102039 |
| P51649 | ALDH5A1 | | 220860000 | 204430000 | 229100000 | 236330000 | 259370000 | 203800000 | 218130000 | 233166666.7 | 0.442855915 | 1.068934427 | 0.096173355 |
| Q02252 | ALDH6A1 | | 1037100000 | 1059900000 | 1107100000 | 938210000 | 1032900000 | 1100300000 | 1068033333 | 1023803333 | 0.437473954 | 0.958587435 | -0.061018066 |
| P27695 | APEX1 | | 180130000 | 233280000 | 287680000 | 291120000 | 333020000 | 453300000 | 233696666.7 | 359146666.7 | 0.095265323 | 1.536806972 | 0.619935969 |
| Q9BUR5 | APOO | | 0 | 0 | 0 | 165260000 | 134520000 | 361110000 | 0 | 220296666.7 | 0.036068439 | N/A | N/A |
| Q6UXV4 | APOOL | | 0 | 112210000 | 0 | 146130000 | 111630000 | 84773000 | 37403333.33 | 114177666.7 | 0.137313536 | 3.05260672 | 1.610041733 |
| P78540 | ARG2 | | 643860000 | 614840000 | 730100000 | 0 | 141700000 | 0 | 662933333.3 | 47233333.33 | 0.000462664 | 0.071248994 | -3.810986538 |
| P25705 | ATP5F1A | | 32993000000 | 34874000000 | 30987000000 | 28358000000 | 23819000000 | 23702000000 | 32951333333 | 25293000000 | 0.015718586 | 0.767586542 | -0.381598678 |
| P06576 | ATP5F1B | | 46924000000 | 40118000000 | 44169000000 | 30941000000 | 38099000000 | 43170000000 | 43737000000 | 37403333333 | 0.193841627 | 0.855187446 | -0.22568742 |
| P36542 | ATP5F1C | | 2703800000 | 2394700000 | 2016800000 | 2297200000 | 2249600000 | 1376600000 | 2371766667 | 1974466667 | 0.330719709 | 0.832487738 | -0.264499073 |
| P30049 | ATP5F1D | | 812530000 | 1007300000 | 393590000 | 1116400000 | 847060000 | 877340000 | 737806666.7 | 946933333.3 | 0.354960865 | 1.283443721 | 0.360020035 |
| P56381 | ATP5F1E | | 42197000 | 0 | 33080000 | 0 | 0 | 0 | 25092333.33 | 0 | 0.12191815 | 0 | N/A |
| Q9UII2 | ATP5IF1 | | 41159000 | 0 | 61692000 | 36766000 | 10773000 | 16171000 | 34283666.67 | 21236666.67 | 0.545767587 | 0.619439772 | -0.690964078 |
| Q96IX5 | ATP5MD | | 232050000 | 348280000 | 408850000 | 422980000 | 335810000 | 244630000 | 329726666.7 | 334473333.3 | 0.951335382 | 1.014395762 | 0.020620623 |
| P56385 | ATP5ME | | 770700000 | 852540000 | 1115500000 | 794180000 | 913160000 | 550550000 | 912913333.3 | 752630000 | 0.342667291 | 0.824426561 | -0.278537109 |
| P56134 | ATP5MF | | 286510000 | 364110000 | 441740000 | 576390000 | 563640000 | 954640000 | 364120000 | 698223333.3 | 0.069758443 | 1.917563807 | 0.939274584 |
| O75964 | ATP5MG | | 1174300000 | 1201400000 | 1110300000 | 1267600000 | 1432200000 | 1306100000 | 1162000000 | 1335300000 | 0.037534692 | 1.149139415 | 0.200553838 |
| P24539 | ATP5PB | | 1348000000 | 1655200000 | 1597800000 | 2802100000 | 2003600000 | 1288900000 | 1533666667 | 2031533333 | 0.32787766 | 1.324625082 | 0.405584081 |
| O75947 | ATP5PD | | 3511300000 | 3727700000 | 3620300000 | 1264100000 | 1500100000 | 2811200000 | 3619766667 | 1858466667 | 0.022166482 | 0.513421675 | -0.961783891 |
| P18859 | ATP5PF | | 1580300000 | 2210500000 | 2457200000 | 2577300000 | 2268200000 | 2220200000 | 2082666667 | 2355233333 | 0.391612143 | 1.13087388 | 0.177438042 |
| P48047 | ATP5PO | | 7121400000 | 7438800000 | 7656800000 | 5207100000 | 4999500000 | 5337700000 | 7405666667 | 5181433333 | 0.000268662 | 0.69965792 | -0.51527837 |
| Q8N5M1 | ATPAF2 | | 48377000 | 32677000 | 58180000 | 0 | 31319000 | 0 | 46411333.33 | 10439666.67 | 0.048434756 | 0.224937874 | -2.152401497 |
| Q13825 | AUH | | 0 | 105720000 | 92271000 | 0 | 0 | 0 | 65997000 | 0 | 0.117948586 | 0 | N/A |
| P12694 | BCKDHA | | 0 | 0 | 190940000 | 170670000 | 0 | 0 | 63646666.67 | 56890000 | 0.940175373 | 0.893840997 | -0.161909878 |
| P21953 | BCKDHB | | 31200000 | 0 | 114330000 | 170850000 | 149290000 | 187020000 | 48510000 | 169053333.3 | 0.028186244 | 3.484917199 | 1.801124378 |
| Q9Y276 | BCS1L | | 98517000 | 95582000 | 35866000 | 67697000 | 18780000 | 53585000 | 76655000 | 46687333.33 | 0.297785851 | 0.6090579 | -0.71534871 |
| Q07021 | C1QBP | | 4989500000 | 4441300000 | 5013900000 | 2529100000 | 3575000000 | 3875000000 | 4814900000 | 3326366667 | 0.029453689 | 0.690848547 | -0.533558629 |
| Q96ER9 | CCDC51 | | 529290000 | 515960000 | 547240000 | 414290000 | 562250000 | 468770000 | 530830000 | 481770000 | 0.328712126 | 0.907578698 | -0.139905348 |
| Q96BP2 | CHCHD1 | | 120890000 | 119330000 | 0 | 128570000 | 135990000 | 144100000 | 80073333.33 | 136220000 | 0.235884567 | 1.701190575 | 0.766544767 |
| Q9NX63 | CHCHD3 | | 288490000 | 342740000 | 432500000 | 546740000 | 293560000 | 124380000 | 354576666.7 | 321560000 | 0.811628537 | 0.906884266 | -0.141009645 |
| Q9BRQ6 | CHCHD6 | | 159380000 | 175230000 | 174270000 | 372830000 | 265840000 | 168440000 | 169626666.7 | 269036666.7 | 0.168669222 | 1.586051721 | 0.665439818 |
| P0C7P0 | CISD3 | | 48513000 | 48666000 | 0 | 0 | 0 | 0 | 32393000 | 0 | 0.116117509 | 0 | N/A |
| P12532 | CKMT1A | | 865540000 | 762420000 | 987800000 | 351670000 | 321890000 | 337860000 | 871920000 | 337140000 | 0.001239851 | 0.386663914 | -1.370847965 |
| Q16740 | CLPP | | 314990000 | 284030000 | 0 | 228630000 | 221320000 | 0 | 199673333.3 | 149983333.3 | 0.711720351 | 0.751143534 | -0.412839479 |
| O76031 | CLPX | | 971810000 | 973110000 | 1044200000 | 654820000 | 627440000 | 561380000 | 996373333.3 | 614546666.7 | 0.00047803 | 0.616783535 | -0.697163842 |
| Q7Z7K0 | CMC1 | | 28531000 | 31789000 | 35186000 | 0 | 0 | 0 | 31835333.33 | 0 | 7.76958E-05 | 0 | N/A |
| Q96BR5 | COA7 | | 134680000 | 138210000 | 142260000 | 151640000 | 119430000 | 127460000 | 138383333.3 | 132843333.3 | 0.606469062 | 0.959966277 | -0.058944369 |

| | | | | | | | | | | | | | |
|---|---|---|---|---|---|---|---|---|---|---|---|---|---|
| Q9Y2Z9 | COQ6 | | 119330000 | 149530000 | 122870000 | 147500000 | 127040000 | 153950000 | 130576666.7 | 142830000 | 0.383004629 | 1.093840145 | 0.129401916 |
| Q96D53 | COQ8B | | 161060000 | 116270000 | 98535000 | 93092000 | 115520000 | 152380000 | 125288333.3 | 120330666.7 | 0.854725066 | 0.960429942 | -0.058247714 |
| O75208 | COQ9 | | 138260000 | 130200000 | 187090000 | 289300000 | 235070000 | 187150000 | 151850000 | 237173333.3 | 0.0684303 | 1.561892218 | 0.643294901 |
| Q14061 | COX17 | | 45114000 | 51556000 | 0 | 0 | 0 | 0 | 32223333.33 | 0 | 0.11787987 | 0 | N/A |
| Q5RI15 | COX20 | | 934260000 | 1208300000 | 1007700000 | 670050000 | 521640000 | 725440000 | 1050086667 | 639043333.3 | 0.015747989 | 0.608562468 | -0.716522734 |
| P13073 | COX4I1 | | 617810000 | 1010000000 | 1851700000 | 1473900000 | 823400000 | 492340000 | 1159836667 | 929880000 | 0.646404731 | 0.801733577 | -0.318805198 |
| P20674 | COX5A | | 4302600000 | 5017700000 | 5169100000 | 2277300000 | 2217400000 | 2149900000 | 4829800000 | 2214866667 | 0.00063354 | 0.458583516 | -1.124743597 |
| P10606 | COX5B | | 1934600000 | 2614000000 | 3079700000 | 3659700000 | 2808400000 | 1621700000 | 2542766667 | 2696600000 | 0.831649215 | 1.060498407 | 0.084742454 |
| P12074 | COX6A1 | | 646860000 | 613950000 | 919130000 | 2015500000 | 1417000000 | 1177300000 | 726646666.7 | 1536600000 | 0.038798037 | 2.114645357 | 1.080415732 |
| P14854 | COX6B1 | | 951720000 | 1202000000 | 1961400000 | 1750100000 | 1822800000 | 2617500000 | 1371706667 | 2063466667 | 0.16804491 | 1.504306071 | 0.589098133 |
| P09669 | COX6C | | 56552000 | 43309000 | 85903000 | 74069000 | 32642000 | 15229000 | 61921333.33 | 40646666.67 | 0.378737672 | 0.65642428 | -0.607299492 |
| P14406 | COX7A2 | | 503740000 | 0 | 0 | 650760000 | 714770000 | 0 | 167913333.3 | 455176666.7 | 0.368134193 | 2.710783341 | 1.43870981 |
| O14548 | COX7A2L | | 166720000 | 136150000 | 541470000 | 469460000 | 346740000 | 400390000 | 281446666.7 | 405530000 | 0.410240719 | 1.440876898 | 0.526947083 |
| P15954 | COX7C | | 193780000 | 139450000 | 39677000 | 152170000 | 171110000 | 275330000 | 124302333.3 | 199536666.7 | 0.272531356 | 1.605252784 | 0.682800501 |
| P36551 | CPOX | | 314900000 | 319870000 | 357270000 | 289270000 | 355470000 | 177800000 | 330680000 | 274180000 | 0.350798727 | 0.829139954 | -0.270312454 |
| P31327 | CPS1 | | 163910000 | 0 | 68440000 | 0 | 55999000 | 0 | 77450000 | 18666333.33 | 0.313795635 | 0.241011405 | -2.052826675 |
| P23786 | CPT2 | | 98522000 | 83377000 | 110030000 | 0 | 0 | 0 | 97309666.67 | 0 | 0.000227786 | 0 | N/A |
| O75390 | CS | | 2720500000 | 2756000000 | 3017000000 | 672950000 | 792790000 | 1273800000 | 2831166667 | 913180000 | 0.000740923 | 0.322545476 | -1.632425513 |
| P00167 | CYB5A | | 410020000 | 488810000 | 642960000 | 592880000 | 562850000 | 341030000 | 513930000 | 498920000 | 0.893056689 | 0.970793688 | -0.042763367 |
| P00387 | CYB5R3 | | 795800000 | 597130000 | 797210000 | 1151700000 | 1134700000 | 1099700000 | 730046666.7 | 1128700000 | 0.004271531 | 1.546065548 | 0.628601486 |
| P08574 | CYC1 | | 3224800000 | 3103900000 | 3098100000 | 3660700000 | 4160600000 | 3736700000 | 3142266667 | 3852666667 | 0.011557231 | 1.226078839 | 0.29405175 |
| Q8N465 | D2HGDH | | 0 | 0 | 0 | 0 | 48341000 | 72377000 | 0 | 40239333.33 | 0.131640555 | N/A | N/A |
| P51398 | DAP3 | | 1271900000 | 1349600000 | 1386200000 | 951530000 | 910720000 | 926530000 | 1335900000 | 929593333.3 | 0.000341106 | 0.695855478 | -0.52314039 |
| P11182 | DBT | | 178110000 | 215290000 | 170770000 | 255060000 | 287710000 | 176280000 | 188056666.7 | 239683333.3 | 0.223047405 | 1.274527182 | 0.349962142 |
| Q16698 | DECR1 | | 476300000 | 456850000 | 376810000 | 0 | 185620000 | 243690000 | 436653333.3 | 143103333.3 | 0.021012433 | 0.327727564 | -1.609431076 |
| Q7Z3D6 | DGLUCY | | 64381000 | 73662000 | 0 | 0 | 0 | 0 | 46014333.33 | 0 | 0.11791137 | 0 | N/A |
| Q02127 | DHODH | | 237920000 | 244130000 | 222270000 | 81057000 | 68338000 | 37789000 | 234773333.3 | 62394666.67 | 0.000278495 | 0.265765561 | -1.911773928 |
| Q13268 | DHRS2 | | 15551000000 | 14376000000 | 13670000000 | 6334100000 | 6090600000 | 7069700000 | 14532333333 | 6498133333 | 0.000207919 | 0.447150033 | -1.161169112 |
| Q9NR28 | DIABLO | | 211010000 | 200920000 | 184750000 | 173310000 | 237300000 | 0 | 198893333.3 | 136870000 | 0.433421022 | 0.688157807 | -0.539188657 |
| P10515 | DLAT | | 1526600000 | 1231800000 | 1471800000 | 715830000 | 934530000 | 1093000000 | 1410066667 | 914453333.3 | 0.025090958 | 0.648517801 | -0.624781921 |
| P09622 | DLD | | 2317400000 | 2642500000 | 2926000000 | 281060000 | 186340000 | 207230000 | 2628633333 | 224876666.7 | 0.000174605 | 0.085548891 | -3.547107036 |
| P36957 | DLST | | 177150000 | 163220000 | 0 | 0 | 0 | 0 | 113456666.7 | 0 | 0.116782308 | 0 | N/A |
| Q96EY1 | DNAJA3 | | 454020000 | 402090000 | 489070000 | 207050000 | 313860000 | 277740000 | 448393333.3 | 266216666.7 | 0.01063356 | 0.593712366 | -0.752163934 |
| Q96DA6 | DNAJC19 | | 3077100000 | 1628100000 | 1382100000 | 369740000 | 679080000 | 623400000 | 2029100000 | 557406666.7 | 0.051953783 | 0.274706356 | -1.864037805 |
| O00429 | DNM1L | | 216550000 | 224950000 | 217320000 | 327450000 | 236240000 | 432740000 | 219606666.7 | 332143333.3 | 0.118800391 | 1.512446495 | 0.596884106 |
| P33316 | DUT | | 215130000 | 0 | 228180000 | 122880000 | 160150000 | 171140000 | 147770000 | 151390000 | 0.964013418 | 1.02449753 | 0.034916506 |
| Q13011 | ECH1 | | 29478000 | 15506000 | 40334000 | 50669000 | 21115000 | 0 | 28439333.33 | 23928000 | 0.796364781 | 0.841369934 | -0.249187831 |
| P30084 | ECHS1 | | 5409500000 | 6963400000 | 5671600000 | 4442900000 | 3127700000 | 2805400000 | 6014833333 | 3458666667 | 0.021134464 | 0.57502286 | -0.798308783 |
| P42126 | ECI1 | | 0 | 0 | 200510000 | 0 | 54165000 | 0 | 66836666.67 | 18055000 | 0.519916165 | 0.270136153 | -1.888241364 |
| Q9BQ95 | ECSIT | | 206650000 | 296940000 | 107510000 | 69316000 | 69085000 | 0 | 203700000 | 46133666.67 | 0.056740933 | 0.226478481 | -2.142554114 |
| P13804 | ETFA | | 5520600000 | 5047100000 | 4964200000 | 2674500000 | 3145400000 | 4480300000 | 5177300000 | 3433400000 | 0.037265208 | 0.663164198 | -0.592561973 |
| O95571 | ETHE1 | | 511710000 | 917850000 | 1119800000 | 933530000 | 868900000 | 883540000 | 849786666.7 | 895323333.3 | 0.812626288 | 1.053585998 | 0.075308078 |
| Q6P587 | FAHD1 | | 205270000 | 217250000 | 136540000 | 144340000 | 134450000 | 175770000 | 186353333.3 | 151520000 | 0.282313363 | 0.813079097 | -0.298532389 |
| Q6P4F2 | FDX2 | | 43486000 | 0 | 42470000 | 0 | 0 | 0 | 28652000 | 0 | 0.116172091 | 0 | N/A |
| P22570 | FDXR | | 2955200000 | 3080300000 | 3640800000 | 4840600000 | 4387800000 | 3793400000 | 3225433333 | 4340600000 | 0.039174255 | 1.345741657 | 0.428401482 |
| P22830 | FECH | | 218270000 | 120520000 | 226990000 | 0 | 0 | 0 | 188593333.3 | 0 | 0.005238875 | 0 | N/A |
| P07954 | FH | | 3031700000 | 3177600000 | 3079100000 | 2179600000 | 2280400000 | 2762100000 | 3096133333 | 2407366667 | 0.020349355 | 0.777539727 | -0.363011706 |
| Q9Y3D6 | FIS1 | | 0 | 0 | 103880000 | 0 | 66598000 | 84507000 | 34626666.67 | 50368333.33 | 0.733589004 | 1.45461109 | 0.54063348 |
| Q8TAE8 | GADD45GIP1 | | 571460000 | 606160000 | 602300000 | 338860000 | 337310000 | 355450000 | 593306666.7 | 343873333.3 | 3.62851E-05 | 0.579587847 | -0.786900751 |
| P0DPI2 | GATD3A | | 507990000 | 452020000 | 461230000 | 562510000 | 442720000 | 239000000 | 473746666.7 | 414743333.3 | 0.572076143 | 0.875453829 | -0.191897001 |
| Q86SX6 | GLRX5 | | 2000800000 | 1982500000 | 1778100000 | 266080000 | 375950000 | 320600000 | 1920466667 | 320876666.7 | 3.35767E-05 | 0.167082654 | -2.581366134 |
| O94925 | GLS | | 5598300000 | 4943700000 | 5403300000 | 3923200000 | 3934000000 | 4207300000 | 5315100000 | 4021500000 | 0.0038535 | 0.756617938 | -0.402363115 |
| P00367 | GLUD1 | | 5338600000 | 6295300000 | 5904600000 | 3854100000 | 3676400000 | 3404700000 | 5846166667 | 3645066667 | 0.002001769 | 0.623496878 | -0.681545758 |
| Q9H4A6 | GOLPH3 | | 208470000 | 163930000 | 258140000 | 153650000 | 208750000 | 151130000 | 210180000 | 171176666.7 | 0.303628324 | 0.814428902 | -0.296139335 |
| P43304 | GPD2 | | 2162100000 | 2332500000 | 1930700000 | 1216300000 | 1196200000 | 1090300000 | 2141766667 | 1167600000 | 0.001367738 | 0.545157425 | -0.875255199 |
| Q9HAV7 | GRPEL1 | | 2299400000 | 2131200000 | 2458600000 | 2009600000 | 1768400000 | 1242300000 | 2296400000 | 1673433333 | 0.064121874 | 0.728720316 | -0.456562883 |
| Q8TAA5 | GRPEL2 | | 193630000 | 260350000 | 244390000 | 0 | 0 | 0 | 232790000 | 0 | 0.000318448 | 0 | N/A |

| | | | | | | | | | | | | | |
|---|---|---|---|---|---|---|---|---|---|---|---|---|---|
| Q16836 | HADH | | 809530000 | 411850000 | 667440000 | 409230000 | 420960000 | 405180000 | 629606666.7 | 411790000 | 0.134743726 | 0.654043265 | -0.612542021 |
| P40939 | HADHA | | 10847000000 | 11373000000 | 10748000000 | 10512000000 | 12413000000 | 12978000000 | 10989333333 | 11967666667 | 0.273119478 | 1.089025722 | 0.12303803 |
| P55084 | HADHB | | 5136200000 | 5497000000 | 4821100000 | 6723100000 | 6626500000 | 6878200000 | 5151433333 | 6742600000 | 0.001585398 | 1.308878435 | 0.38833111 |
| P31937 | HIBADH | | 1693000000 | 1461400000 | 1314200000 | 992100000 | 1209500000 | 1444200000 | 1489533333 | 1215266667 | 0.183737688 | 0.815870743 | -0.293587489 |
| Q6NVY1 | HIBCH | | 201910000 | 290380000 | 209490000 | 116610000 | 96819000 | 39691000 | 233926666.7 | 84373333.33 | 0.014904942 | 0.360682835 | -1.471197328 |
| Q9BW72 | HIGD2A | | 1095600000 | 708180000 | 497930000 | 52448000 | 146000000 | 462610000 | 767236666.7 | 220352666.7 | 0.06339088 | 0.287202993 | -1.799857313 |
| Q9BX68 | HINT2 | | 1457100000 | 1063800000 | 1671800000 | 868120000 | 1093500000 | 845850000 | 1397566667 | 935823333.3 | 0.076795049 | 0.669609082 | -0.578608998 |
| P35914 | HMGCL | | 46890000 | 60343000 | 0 | 92188000 | 107190000 | 54825000 | 35744333.33 | 84734333.33 | 0.110985381 | 2.370566896 | 1.245232106 |
| Q99714 | HSD17B10 | | 9046600000 | 8625900000 | 8663400000 | 7041200000 | 8580600000 | 9874300000 | 8778633333 | 8498700000 | 0.752820482 | 0.968111969 | -0.04675418 |
| P38646 | HSPA9 | | 31444000000 | 28741000000 | 28631000000 | 18474000000 | 20944000000 | 21299000000 | 29605333333 | 20239000000 | 0.001849406 | 0.683626824 | -0.548719089 |
| P10809 | HSPD1 | | 87661000000 | 79291000000 | 71589000000 | 55788000000 | 58320000000 | 65331000000 | 79513666667 | 59813000000 | 0.022438752 | 0.752235465 | -0.41074377 |
| P61604 | HSPE1 | | 21309000000 | 20659000000 | 23798000000 | 14659000000 | 14743000000 | 12960000000 | 21922000000 | 14120666667 | 0.00222654 | 0.644132226 | -0.634571222 |
| O43464 | HTRA2 | | 0 | 0 | 183180000 | 75231000 | 94193000 | 139740000 | 61060000 | 103054666.7 | 0.547486315 | 1.687760673 | 0.755110342 |
| Q9NSE4 | IARS2 | | 2087500000 | 1933500000 | 2092500000 | 3078600000 | 3016400000 | 2278700000 | 2037833333 | 2791233333 | 0.045285115 | 1.369706388 | 0.453866668 |
| Q5T440 | IBA57 | | 684310000 | 682030000 | 620270000 | 491550000 | 522640000 | 482810000 | 662203333.3 | 499000000 | 0.002523641 | 0.753544984 | -0.408234457 |
| P48735 | IDH2 | | 293000000 | 235770000 | 277650000 | 213580000 | 149900000 | 242510000 | 268806666.7 | 201996666.7 | 0.107115799 | 0.751457057 | -0.412237433 |
| P50213 | IDH3A | | 857290000 | 732990000 | 704290000 | 633620000 | 670070000 | 789920000 | 764856666.7 | 697870000 | 0.371322614 | 0.91241932 | -0.132231099 |
| O43837 | IDH3B | | 146900000 | 100170000 | 143980000 | 156870000 | 192470000 | 144390000 | 130350000 | 164576666.7 | 0.176486764 | 1.262575118 | 0.336369225 |
| P51553 | IDH3G | | 0 | 85738000 | 0 | 90118000 | 80299000 | 0 | 28579333.33 | 56805666.67 | 0.523171446 | 1.987648417 | 0.99106259 |
| Q16891 | IMMT | | 4099900000 | 4769900000 | 4897300000 | 4681600000 | 4462200000 | 3329700000 | 4589033333 | 4157833333 | 0.425463496 | 0.906036856 | -0.142358357 |
| Q86U28 | ISCA2 | | 365330000 | 373800000 | 343410000 | 156790000 | 205000000 | 277580000 | 360846666.7 | 213123333.3 | 0.015164257 | 0.590620208 | -0.759697376 |
| Q9H1K1 | ISCU | | 0 | 72101000 | 0 | 0 | 0 | 64127000 | 24033666.67 | 21375666.67 | 0.93810919 | 0.88940514 | -0.169087353 |
| P26440 | IVD | | 95438000 | 181480000 | 129270000 | 290320000 | 187430000 | 105700000 | 135396000 | 194483333.3 | 0.373150808 | 1.436403833 | 0.522461408 |
| Q9H9P8 | L2HGDH | | 200550000 | 169740000 | 123380000 | 156870000 | 81821000 | 196330000 | 164556666.7 | 145007000 | 0.653628405 | 0.881197966 | -0.182461929 |
| P83111 | LACTB | | 1270100000 | 824260000 | 639460000 | 714050000 | 758510000 | 642630000 | 911273333.3 | 705063333.3 | 0.33925473 | 0.773712241 | -0.370130995 |
| O95202 | LETM1 | | 1551800000 | 1448500000 | 1837700000 | 1115000000 | 1150700000 | 1184300000 | 1612666667 | 1150000000 | 0.017286875 | 0.713104589 | -0.487814407 |
| P36776 | LONP1 | | 2559400000 | 2373700000 | 2193600000 | 1816600000 | 2135800000 | 2333900000 | 2375566667 | 2095433333 | 0.202569189 | 0.882077259 | -0.181023072 |
| P42704 | LRPPRC | | 15714000000 | 14870000000 | 14931000000 | 8381500000 | 8612700000 | 9205800000 | 15171666667 | 8733333333 | 6.14638E-05 | 0.575634406 | -0.796775269 |
| Q8WWC4 | MAIP1 | | 0 | 215060000 | 175930000 | 0 | 0 | 82820000 | 130330000 | 27606666.67 | 0.225053411 | 0.211821274 | -2.239080602 |
| Q9NX47 | MARCHF5 | | 0 | 0 | 0 | 55381000 | 64631000 | 0 | 0 | 40004000 | 0.118474101 | N/A | N/A |
| Q8IVS2 | MCAT | | 156610000 | 227400000 | 144950000 | 103110000 | 69796000 | 173310000 | 176320000 | 115405333.3 | 0.201812577 | 0.654522081 | -0.611486231 |
| Q96RQ3 | MCCC1 | | 124670000 | 0 | 88954000 | 40777000 | 102780000 | 0 | 71208000 | 47852333.33 | 0.64943396 | 0.672007827 | -0.573450059 |
| Q9HCC0 | MCCC2 | | 692490000 | 528710000 | 867820000 | 1010200000 | 940390000 | 889690000 | 696340000 | 946760000 | 0.07364385 | 1.359623173 | 0.443206856 |
| Q96PE7 | MCEE | | 0 | 0 | 0 | 118650000 | 74729000 | 0 | 0 | 64459666.67 | 0.136216592 | N/A | N/A |
| Q8NE86 | MCU | | 186680000 | 226180000 | 187200000 | 81767000 | 149240000 | 166380000 | 200020000 | 132462333.3 | 0.079935523 | 0.662245442 | -0.594562086 |
| P40926 | MDH2 | | 34905000000 | 31124000000 | 34717000000 | 24425000000 | 22772000000 | 24022000000 | 33582000000 | 23739666667 | 0.001763612 | 0.706916404 | -0.500388476 |
| P23368 | ME2 | | 2125400000 | 1865400000 | 1255100000 | 698040000 | 840650000 | 1122200000 | 1748633333 | 886963333.3 | 0.039626163 | 0.507232315 | -0.979281435 |
| Q9GZY8 | MFF | | 97963000 | 0 | 124790000 | 0 | 0 | 0 | 74251000 | 0 | 0.121853662 | 0 | N/A |
| Q8IWA4 | MFN1 | | 302740000 | 275140000 | 179030000 | 142990000 | 147740000 | 337110000 | 252303333.3 | 209280000 | 0.592704095 | 0.829477745 | -0.269724821 |
| O95140 | MFN2 | | 297850000 | 261470000 | 258230000 | 414710000 | 362410000 | 170500000 | 272516666.7 | 315873333.3 | 0.595681922 | 1.159097303 | 0.213001682 |
| Q9BQP7 | MGME1 | | 330980000 | 322380000 | 297960000 | 175860000 | 169890000 | 161060000 | 317106666.7 | 168936666.7 | 0.000162567 | 0.532743977 | -0.908485718 |
| Q5XKP0 | MICOS13 | | 0 | 78087000 | 53012000 | 41516000 | 0 | 0 | 43699666.67 | 13838666.67 | 0.328527839 | 0.31667671 | -1.658917327 |
| Q9BPX6 | MICU1 | | 128730000 | 90958000 | 116780000 | 0 | 69637000 | 0 | 112156000 | 23212333.33 | 0.02595588 | 0.206964704 | -2.272543346 |
| Q8IYU8 | MICU2 | | 26694000 | 115140000 | 195820000 | 0 | 0 | 0 | 112551333.3 | 0 | 0.082528835 | 0 | N/A |
| Q99797 | MIPEP | | 259080000 | 198180000 | 131800000 | 199120000 | 65527000 | 150100000 | 196353333.3 | 138249000 | 0.339339033 | 0.704082776 | -0.506183045 |
| P22033 | MMUT | | 373570000 | 408860000 | 242860000 | 118600000 | 141870000 | 307370000 | 341763333.3 | 189280000 | 0.122191015 | 0.55383355 | -0.852475645 |
| Q9NZJ7 | MTCH1 | | 0 | 0 | 74418000 | 49459000 | 0 | 0 | 24806000 | 16486333.33 | 0.793842814 | 0.664610712 | -0.589418548 |
| Q9Y6C9 | MTCH2 | | 784000000 | 437690000 | 558160000 | 2775600000 | 2825000000 | 1810500000 | 593283333.3 | 2470366667 | 0.00556829 | 4.163890215 | 2.057932031 |
| P00403 | MT | | 1326200000 | 1216400000 | 1243500000 | 489730000 | 651530000 | 1165800000 | 1262033333 | 769020000 | 0.075347951 | 0.609349991 | -0.714656992 |
| Q9H019 | MTFR1L | | 172090000 | 235130000 | 62838000 | 139870000 | 167560000 | 162980000 | 156686000 | 156803333.3 | 0.998276321 | 1.000748844 | 0.001079949 |
| Q6UB35 | MTHFD1L | | 5160500000 | 5017000000 | 4084800000 | 2497800000 | 2914800000 | 3757300000 | 4754100000 | 3056633333 | 0.027564816 | 0.64294679 | -0.63722875 |
| P13995 | MTHFD2 | | 2774000000 | 2167900000 | 2936700000 | 2254100000 | 2352500000 | 3074200000 | 2626200000 | 2560266667 | 0.859212749 | 0.974894017 | -0.036682707 |
| Q9NVV4 | MTPAP | | 249810000 | 0 | 187620000 | 0 | 72173000 | 0 | 145810000 | 24057666.67 | 0.197412162 | 0.164993256 | -2.599521038 |
| Q13505 | MTX1 | | 239000000 | 240480000 | 342850000 | 328470000 | 333890000 | 306510000 | 274110000 | 322956666.7 | 0.239492652 | 1.178200966 | 0.23658564 |
| O75431 | MTX2 | | 1009500000 | 1573400000 | 1599800000 | 768840000 | 848580000 | 1627900000 | 1394233333 | 1081773333 | 0.403636734 | 0.775891171 | -0.366073786 |
| Q4G0N4 | NADK2 | | 185730000 | 164220000 | 181110000 | 136020000 | 143120000 | 0 | 177020000 | 93046666.67 | 0.148691209 | 0.525627989 | -0.927885995 |
| O95299 | NDUFA10 | | 349660000 | 210120000 | 222480000 | 135140000 | 124610000 | 0 | 260753333.3 | 86583333.33 | 0.048864729 | 0.332050725 | -1.590524447 |

| | | | | | | | | | | | | | |
|---|---|---|---|---|---|---|---|---|---|---|---|---|---|
| Q9UI09 | NDUFA12 | | 871680000 | 710700000 | 910580000 | 390720000 | 460590000 | 355090000 | 830986666.7 | 402133333.3 | 0.003335042 | 0.483922726 | -1.047151401 |
| Q9P0J0 | NDUFA13 | | 289180000 | 315900000 | 292470000 | 155470000 | 160540000 | 219610000 | 299183333.3 | 178540000 | 0.005599333 | 0.596757841 | -0.744782478 |
| O43678 | NDUFA2 | | 571340000 | 489940000 | 703680000 | 336990000 | 376790000 | 372190000 | 588320000 | 361990000 | 0.023542019 | 0.615294398 | -0.700651238 |
| Q16718 | NDUFA5 | | 1648800000 | 1556100000 | 1285400000 | 908640000 | 939330000 | 791800000 | 1496766667 | 879923333.3 | 0.006379077 | 0.58788277 | -0.766399601 |
| P56556 | NDUFA6 | | 139690000 | 185610000 | 174240000 | 276860000 | 212290000 | 94542000 | 166513333.3 | 194564000 | 0.637659802 | 1.168458982 | 0.22460709 |
| O95182 | NDUFA7 | | 125070000 | 174020000 | 263460000 | 401420000 | 200490000 | 109580000 | 187516666.7 | 237163333.3 | 0.629786262 | 1.264758688 | 0.33886215 |
| P51970 | NDUFA8 | | 208420000 | 0 | 180330000 | 93810000 | 98069000 | 0 | 129583333.3 | 63959666.67 | 0.417853647 | 0.493579421 | -1.018645849 |
| Q16795 | NDUFA9 | | 909280000 | 953840000 | 593260000 | 873220000 | 752670000 | 845860000 | 818793333.3 | 823916666.7 | 0.967781816 | 1.006257175 | 0.008999071 |
| Q8N183 | NDUFAF2 | | 901490000 | 709090000 | 868810000 | 645540000 | 852460000 | 932850000 | 826463333.3 | 810283333.3 | 0.884117939 | 0.980422604 | -0.028524349 |
| Q7L592 | NDUFAF7 | | 71094000 | 0 | 96771000 | 57833000 | 110760000 | 0 | 55955000 | 56197666.67 | 0.995780759 | 1.004336818 | 0.006243178 |
| O96000 | NDUFB10 | | 311550000 | 0 | 341060000 | 222120000 | 317950000 | 245310000 | 217536666.7 | 261793333.3 | 0.714942213 | 1.20344463 | 0.267169766 |
| O95168 | NDUFB4 | | 679690000 | 541040000 | 711280000 | 454090000 | 485390000 | 628480000 | 644003333.3 | 522653333.3 | 0.180665464 | 0.811569298 | -0.301213806 |
| O43674 | NDUFB5 | | 32282000 | 0 | 167200000 | 0 | 0 | 0 | 66494000 | 0 | 0.263916355 | 0 | N/A |
| O95169 | NDUFB8 | | 0 | 0 | 191770000 | 180930000 | 217500000 | 0 | 63923333.33 | 132810000 | 0.499014417 | 2.077645096 | 1.054949233 |
| Q9Y6M9 | NDUFB9 | | 183590000 | 201000000 | 0 | 123570000 | 225700000 | 279000000 | 128196666.7 | 209423333.3 | 0.361015627 | 1.633609818 | 0.708063442 |
| P28331 | NDUFS1 | | 2900400000 | 2953400000 | 3188100000 | 1886400000 | 2057300000 | 2228100000 | 3013966667 | 2057266667 | 0.001948849 | 0.682577777 | -0.550934651 |
| O75306 | NDUFS2 | | 1157100000 | 1044100000 | 1117400000 | 893350000 | 926050000 | 911540000 | 1106200000 | 910313333.3 | 0.004709851 | 0.822919303 | -0.28117713 |
| O75489 | NDUFS3 | | 1866800000 | 1781800000 | 1679300000 | 972910000 | 1114500000 | 1572100000 | 1775966667 | 1219836667 | 0.042128624 | 0.686857861 | -0.541916517 |
| O43181 | NDUFS4 | | 72510000 | 145140000 | 183240000 | 106880000 | 0 | 0 | 133630000 | 35626666.67 | 0.111846818 | 0.2666068 | -1.907214518 |
| O43920 | NDUFS5 | | 125210000 | 228490000 | 194600000 | 208010000 | 178990000 | 92683000 | 182766666.7 | 159894333.3 | 0.645682007 | 0.874855006 | -0.192884162 |
| O75380 | NDUFS6 | | 126940000 | 228920000 | 214770000 | 279660000 | 183800000 | 109150000 | 190210000 | 190870000 | 0.99157615 | 1.003469849 | 0.004997269 |
| O75251 | NDUFS7 | | 559210000 | 0 | 0 | 468150000 | 425970000 | 602280000 | 186403333.3 | 498800000 | 0.182325216 | 2.675917813 | 1.420033806 |
| O00217 | NDUFS8 | | 565440000 | 353090000 | 449080000 | 151760000 | 160800000 | 306880000 | 455870000 | 206480000 | 0.034747391 | 0.452936144 | -1.142620425 |
| P49821 | NDUFV1 | | 670690000 | 929210000 | 1041300000 | 627450000 | 582550000 | 520670000 | 880400000 | 576890000 | 0.056273887 | 0.655258973 | -0.60986289 |
| P19404 | NDUFV2 | | 758260000 | 931380000 | 1011100000 | 563040000 | 562360000 | 868010000 | 900246666.7 | 664470000 | 0.135103566 | 0.738097707 | -0.438116288 |
| Q9Y697 | NFS1 | | 36675000 | 53341000 | 31650000 | 57937000 | 42004000 | 0 | 40555333.33 | 33313666.67 | 0.715165718 | 0.821437378 | -0.283777499 |
| Q9UMS0 | NFU1 | | 470180000 | 495800000 | 516630000 | 247860000 | 232330000 | 0 | 494203333.3 | 160063333.3 | 0.014717311 | 0.323881533 | -1.626461883 |
| Q9BYT8 | NLN | | 211460000 | 170190000 | 179050000 | 136310000 | 191620000 | 114520000 | 186900000 | 147483333.3 | 0.206232343 | 0.789102907 | -0.34171464 |
| Q13423 | NNT | | 2331900000 | 2307800000 | 2693300000 | 2698000000 | 2309200000 | 2075600000 | 2444333333 | 2360933333 | 0.724141145 | 0.965880267 | -0.050083735 |
| P80303 | NUCB2 | | 927150000 | 753910000 | 578060000 | 503050000 | 466020000 | 218010000 | 753040000 | 395693333.3 | 0.056889404 | 0.525461242 | -0.928343741 |
| Q9BW91 | NUDT9 | | 166210000 | 204610000 | 213790000 | 191720000 | 183270000 | 234480000 | 194870000 | 203156666.7 | 0.719928311 | 1.042524076 | 0.060080701 |
| P04181 | OAT | | 5759900000 | 6858700000 | 5632400000 | 4048200000 | 4016200000 | 4009500000 | 6083666667 | 4024633333 | 0.006140475 | 0.661547312 | -0.596083756 |
| Q02218 | OGDH | | 1251800000 | 1225900000 | 1348400000 | 1113900000 | 976890000 | 1322400000 | 1275366667 | 1137730000 | 0.268273991 | 0.892080709 | -0.164753855 |
| O60313 | OPA1 | | 1585100000 | 1491900000 | 1814500000 | 1445400000 | 1524000000 | 1622000000 | 1630500000 | 1530466667 | 0.409175811 | 0.938648676 | -0.091342818 |
| Q15070 | OXA1L | | 41913000 | 40111000 | 37959000 | 0 | 0 | 0 | 39994333.33 | 0 | 3.97966E-06 | 0 | N/A |
| P55809 | OXCT1 | | 1449800000 | 1329900000 | 855740000 | 670530000 | 1218200000 | 1408200000 | 1211813333 | 1098976667 | 0.713317278 | 0.906886099 | -0.141006729 |
| Q9Y3D7 | PAM16 | | 396170000 | 476070000 | 385260000 | 385850000 | 411970000 | 146370000 | 419166666.7 | 314730000 | 0.306819689 | 0.750846918 | -0.413409292 |
| P11498 | PC | | 136780000 | 354520000 | 168640000 | 168110000 | 214610000 | 120340000 | 219980000 | 167686666.7 | 0.514172606 | 0.76228142 | -0.391604384 |
| P05166 | PCCB | | 143650000 | 157320000 | 144910000 | 0 | 0 | 0 | 148626666.7 | 0 | 4.42549E-06 | 0 | N/A |
| Q16822 | PCK2 | | 961930000 | 839060000 | 1017400000 | 563830000 | 670400000 | 859560000 | 939463333.3 | 697930000 | 0.075559365 | 0.742902863 | -0.428754509 |
| Q6L8Q7 | PDE12 | | 218360000 | 151670000 | 156430000 | 103600000 | 0 | 0 | 175486666.7 | 34533333.33 | 0.025680149 | 0.196786081 | -2.345299918 |
| Q9HBH1 | PDF | | 0 | 334750000 | 73277000 | 93268000 | 111990000 | 205540000 | 136009000 | 136932666.7 | 0.993547996 | 1.006791217 | 0.009764536 |
| P08559 | PDHA1 | | 2535700000 | 2469900000 | 2376000000 | 2180600000 | 2510300000 | 2636700000 | 2460533333 | 2442533333 | 0.906317882 | 0.992684513 | -0.01059281 |
| P11177 | PDHB | | 2124500000 | 2086000000 | 2040700000 | 1532000000 | 1310600000 | 2130600000 | 2083733333 | 1657733333 | 0.158514492 | 0.795559253 | -0.32995871 |
| O00330 | PDHX | | 313890000 | 336600000 | 370230000 | 135160000 | 169790000 | 0 | 340240000 | 101650000 | 0.011763398 | 0.298759699 | -1.742942545 |
| Q9P0J1 | PDP1 | | 0 | 0 | 65031000 | 112830000 | 92688000 | 0 | 21677000 | 68506000 | 0.316601224 | 3.160308161 | 1.660065242 |
| Q8NCN5 | PDPR | | 137520000 | 77590000 | 126640000 | 77807000 | 148570000 | 93228000 | 113916666.7 | 106535000 | 0.807147763 | 0.93520117 | -0.096651359 |
| Q96HS1 | PGAM5 | | 810690000 | 1000200000 | 1334400000 | 1272100000 | 1297000000 | 784870000 | 1048430000 | 1117990000 | 0.773935281 | 1.066346823 | 0.092676743 |
| Q9UG56 | PISD | | 213200000 | 153670000 | 215190000 | 113660000 | 97938000 | 0 | 194020000 | 70532666.67 | 0.039152599 | 0.363532969 | -1.459841886 |
| Q5JRX3 | PITRM1 | | 178910000 | 172190000 | 172860000 | 183650000 | 114750000 | 180100000 | 174653333.3 | 159500000 | 0.537555306 | 0.913237652 | -0.130937753 |
| Q10713 | PMPCA | | 965430000 | 835310000 | 999760000 | 957040000 | 892790000 | 778840000 | 933500000 | 876223333.3 | 0.472419847 | 0.938643099 | -0.091351389 |
| O75439 | PMPCB | | 448500000 | 280630000 | 284880000 | 404690000 | 383130000 | 567690000 | 338003333.3 | 451836666.7 | 0.229280473 | 1.336781689 | 0.418763876 |
| Q8TCS8 | PNPT1 | | 978560000 | 905990000 | 1113900000 | 865320000 | 732360000 | 643620000 | 999483333.3 | 747100000 | 0.04656459 | 0.747486201 | -0.419881148 |
| P54098 | POLG | | 0 | 0 | 0 | 0 | 52754000 | 62324000 | 0 | 38359333.33 | 0.118860052 | N/A | N/A |
| Q9UHN1 | POLG2 | | 0 | 0 | 0 | 0 | 35192000 | 31430000 | 0 | 22207333.33 | 0.117383265 | N/A | N/A |
| P30405 | PPIF | | 2773900000 | 2123400000 | 2105700000 | 1534200000 | 1634000000 | 1624100000 | 2334333333 | 1597433333 | 0.02944634 | 0.684321005 | -0.547254863 |
| P30048 | PRDX3 | | 6423700000 | 4899600000 | 5564000000 | 6029400000 | 7551500000 | 7647300000 | 5629100000 | 7076066667 | 0.102231231 | 1.257051157 | 0.330043363 |

| | | | | | | | | | | | | | |
|---|---|---|---|---|---|---|---|---|---|---|---|---|---|
| P30044 | PRDX5 | | 8044200000 | 6384100000 | 5384200000 | 3330100000 | 3571000000 | 5045900000 | 6604166667 | 3982333333 | 0.049812317 | 0.603003155 | -0.729762545 |
| Q96EY7 | PTCD3 | | 1133800000 | 750950000 | 883300000 | 831650000 | 920520000 | 854190000 | 922683333.3 | 868786666.7 | 0.664735315 | 0.941587038 | -0.086833635 |
| Q8WUK0 | PTPMT1 | | 247360000 | 218060000 | 162040000 | 32995000 | 43746000 | 0 | 209153333.3 | 25580333.33 | 0.002903918 | 0.122304211 | -3.031454022 |
| Q9Y3E5 | PTRH2 | | 826720000 | 774010000 | 1052600000 | 837070000 | 927270000 | 760850000 | 884443333.3 | 841730000 | 0.685588152 | 0.951705969 | -0.071412175 |
| P32322 | PYCR1 | | 3330900000 | 2975500000 | 3142400000 | 2980600000 | 2477200000 | 1974700000 | 3149600000 | 2477500000 | 0.094510973 | 0.786607823 | -0.34628356 |
| Q8IXI2 | RHOT1 | | 112610000 | 105590000 | 0 | 65571000 | 80818000 | 72555000 | 72733333.33 | 72981333.33 | 0.994930365 | 1.003409716 | 0.004910813 |
| Q8IXI1 | RHOT2 | | 596280000 | 508800000 | 448280000 | 316020000 | 463230000 | 424540000 | 517786666.7 | 401263333.3 | 0.131214531 | 0.774958799 | -0.367808484 |
| P60602 | ROMO1 | | 0 | 0 | 0 | 148870000 | 0 | 132800000 | 0 | 93890000 | 0.117409604 | N/A | N/A |
| O75880 | SCO1 | | 0 | 0 | 492750000 | 282580000 | 0 | 199340000 | 164250000 | 160640000 | 0.985319154 | 0.978021309 | -0.032062196 |
| P31040 | SDHA | | 3914900000 | 4472100000 | 5007000000 | 3531500000 | 3197400000 | 2546200000 | 4464666667 | 3091700000 | 0.032631906 | 0.692481708 | -0.530152132 |
| P21912 | SDHB | | 2278600000 | 2495300000 | 2508700000 | 1485500000 | 1415100000 | 1870700000 | 2427533333 | 1590433333 | 0.006380237 | 0.655164364 | -0.610071207 |
| P34897 | SHMT2 | | 9112100000 | 7110000000 | 7494600000 | 3447200000 | 3313400000 | 3416800000 | 7905566667 | 3392466667 | 0.001832749 | 0.429123782 | -1.220534238 |
| P53007 | SLC25A1 | | 503270000 | 496590000 | 458120000 | 329350000 | 388920000 | 352210000 | 485993333.3 | 356826666.7 | 0.00444377 | 0.73422132 | -0.445713088 |
| Q9UBX3 | SLC25A10 | | 1253500000 | 526080000 | 746910000 | 72481000 | 100320000 | 291010000 | 842163333.3 | 154603666.7 | 0.038319813 | 0.183579195 | -2.445525528 |
| Q02978 | SLC25A11 | | 79011000 | 124830000 | 95207000 | 0 | 0 | 0 | 99682666.67 | 0 | 0.001751096 | 0 | N/A |
| O75746 | SLC25A12 | | 193100000 | 194340000 | 169700000 | 64857000 | 159980000 | 255830000 | 185713333.3 | 160222333.3 | 0.671008798 | 0.862740065 | -0.213002139 |
| Q9UJS0 | SLC25A13 | | 2132200000 | 2104000000 | 1677300000 | 1692500000 | 2281300000 | 2305400000 | 1971166667 | 2093066667 | 0.649609759 | 1.061841549 | 0.086568499 |
| Q9H936 | SLC25A22 | | 324210000 | 78720000 | 171970000 | 133250000 | 73552000 | 272970000 | 191633333.3 | 159924000 | 0.749759745 | 0.834531223 | -0.260962068 |
| Q6NUK1 | SLC25A24 | | 1004900000 | 941210000 | 981840000 | 529420000 | 523900000 | 579960000 | 975983333.3 | 544426666.7 | 7.46658E-05 | 0.557823733 | -0.842118778 |
| Q00325 | SLC25A3 | | 385920000 | 540890000 | 412450000 | 822170000 | 713320000 | 342370000 | 446420000 | 625953333.3 | 0.305487542 | 1.402162388 | 0.487653441 |
| Q9GZT3 | SLIRP | | 3246000000 | 3306800000 | 3778400000 | 3255800000 | 3215300000 | 2342700000 | 3443733333 | 2937933333 | 0.213330312 | 0.853124516 | -0.229171772 |
| Q04837 | SSBP1 | | 206800000 | 464180000 | 491440000 | 345990000 | 491970000 | 521070000 | 387473333.3 | 453010000 | 0.568552603 | 1.169138521 | 0.225445873 |
| Q9UJZ1 | STOML2 | | 4685800000 | 5196300000 | 4594300000 | 4317500000 | 4277600000 | 5797600000 | 4825466667 | 4797566667 | 0.960841701 | 0.994218176 | -0.008365617 |
| Q9P2R7 | SUCLA2 | | 260370000 | 233320000 | 272340000 | 286040000 | 343360000 | 173410000 | 255343333.3 | 267603333.3 | 0.822644205 | 1.048013785 | 0.067657694 |
| P53597 | SUCLG1 | | 1123900000 | 1129900000 | 1225800000 | 984570000 | 1208700000 | 805890000 | 1159866667 | 999720000 | 0.256613935 | 0.861926658 | -0.21436298 |
| Q96I99 | SUCLG2 | | 2956500000 | 2986600000 | 2715100000 | 1923200000 | 2325100000 | 2159100000 | 2886066667 | 2135800000 | 0.00660663 | 0.740038345 | -0.434328069 |
| P51687 | SUOX | | 112630000 | 81026000 | 21531000 | 0 | 0 | 0 | 71729000 | 0 | 0.054890784 | 0 | N/A |
| Q8IYB8 | SUPV3L1 | | 123550000 | 135740000 | 198850000 | 185080000 | 192650000 | 202480000 | 152713333.3 | 193403333.3 | 0.163497955 | 1.266446938 | 0.340786631 |
| P57105 | SYNJ2BP | | 1047800000 | 848020000 | 786900000 | 1426600000 | 1135200000 | 755480000 | 894240000 | 1105760000 | 0.370105859 | 1.236536053 | 0.306304304 |
| Q00059 | TFAM | | 1229400000 | 1970300000 | 1807200000 | 1281500000 | 1107900000 | 673050000 | 1668966667 | 1020816667 | 0.088021404 | 0.611645929 | -0.709231352 |
| P62072 | TIMM10 | | 46375000 | 57105000 | 0 | 70209000 | 42744000 | 0 | 34493333.33 | 37651000 | 0.912254489 | 1.09154426 | 0.12637063 |
| Q9Y5J6 | TIMM10B | | 192560000 | 0 | 138320000 | 0 | 0 | 0 | 110293333.3 | 0 | 0.126693793 | 0 | N/A |
| Q9Y5L4 | TIMM13 | | 989060000 | 0 | 1126600000 | 441460000 | 412340000 | 439790000 | 705220000 | 431196666.7 | 0.483205258 | 0.611435675 | -0.709727365 |
| Q99595 | TIMM17A | | 42813000 | 0 | 177010000 | 51239000 | 125480000 | 0 | 73274333.33 | 58906333.33 | 0.834821148 | 0.803914968 | -0.314885183 |
| O60830 | TIMM17B | | 123340000 | 0 | 0 | 130500000 | 140990000 | 240250000 | 41113333.33 | 170580000 | 0.074454443 | 4.149018972 | 2.052770254 |
| Q9BVV7 | TIMM21 | | 122620000 | 110680000 | 0 | 113500000 | 154610000 | 122410000 | 77766666.67 | 130173333.3 | 0.270149654 | 1.673896271 | 0.743210129 |
| Q9Y584 | TIMM22 | | 238210000 | 307340000 | 233950000 | 152610000 | 259660000 | 261340000 | 259833333.3 | 224536666.7 | 0.458994285 | 0.864156511 | -0.210635467 |
| O14925 | TIMM23 | | 113980000 | 0 | 80075000 | 0 | 0 | 0 | 64685000 | 0 | 0.128114686 | 0 | N/A |
| O43615 | TIMM44 | | 1567800000 | 1569500000 | 1521500000 | 1036500000 | 851720000 | 679320000 | 1552933333 | 855846666.7 | 0.002607872 | 0.551116167 | -0.859571645 |
| Q3ZCQ8 | TIMM50 | | 1052800000 | 1310000000 | 1072600000 | 774190000 | 650150000 | 826030000 | 1145133333 | 750123333.3 | 0.015578508 | 0.655053269 | -0.610315864 |
| O60220 | TIMM8A | | 728590000 | 674920000 | 693020000 | 321450000 | 363570000 | 446210000 | 698843333.3 | 377076666.7 | 0.001282939 | 0.539572532 | -0.890111189 |
| Q9Y5J7 | TIMM9 | | 384450000 | 469510000 | 619780000 | 220630000 | 280580000 | 249160000 | 491246666.7 | 250123333.3 | 0.027306366 | 0.509160368 | -0.973807969 |
| Q9NS69 | TOMM22 | | 533310000 | 461330000 | 590370000 | 191120000 | 276410000 | 561930000 | 528336666.7 | 343153333.3 | 0.192181109 | 0.64949748 | -0.622604168 |
| Q15785 | TOMM34 | | 74216000 | 91449000 | 180040000 | 355860000 | 489250000 | 586130000 | 115235000 | 477080000 | 0.008245289 | 4.140061613 | 2.049652238 |
| O96008 | TOMM40 | | 1219200000 | 1025500000 | 959110000 | 900690000 | 638980000 | 720390000 | 1067936667 | 753353333.3 | 0.045754495 | 0.705428849 | -0.503427518 |
| Q96B49 | TOMM6 | | 3757400000 | 2276100000 | 3312000000 | 1037600000 | 213830000 | 2292600000 | 3115166667 | 1181343333 | 0.060732419 | 0.379223156 | -1.398881036 |
| O94826 | TOMM70 | | 3418200000 | 3339900000 | 3659900000 | 2893400000 | 3569100000 | 3403300000 | 3472666667 | 3288600000 | 0.459191191 | 0.946995585 | -0.078570396 |
| Q12931 | TRAP1 | | 5529400000 | 5321800000 | 5169700000 | 3499100000 | 3996400000 | 5561100000 | 5340300000 | 4352200000 | 0.191832692 | 0.814972942 | -0.295175935 |
| Q96Q11 | TRNT1 | | 118460000 | 0 | 119590000 | 0 | 0 | 82242000 | 79350000 | 27414000 | 0.342128892 | 0.345482042 | -1.533317375 |
| P30536 | TSPO | | 0 | 0 | 29531000 | 29892000 | 82218000 | 277100000 | 9843666.667 | 129736666.7 | 0.189135998 | 13.17970946 | 3.720246662 |
| Q99757 | TXN2 | | 177180000 | 0 | 0 | 138570000 | 122170000 | 117470000 | 59060000 | 126070000 | 0.322391585 | 2.134608872 | 1.093971747 |
| Q9BRT2 | UQCC2 | | 0 | 152500000 | 176970000 | 126550000 | 130130000 | 0 | 109823333.3 | 85560000 | 0.746254765 | 0.779069415 | -0.360176218 |
| Q9UDW1 | UQCR10 | | 113580000 | 121780000 | 0 | 523730000 | 553390000 | 583480000 | 78453333.33 | 553533333.3 | 0.000378726 | 7.055574439 | 2.818763547 |
| P14927 | UQCRB | | 2008000000 | 1994200000 | 2763900000 | 1730000000 | 1803500000 | 1496800000 | 2255366667 | 1676766667 | 0.099271229 | 0.743456349 | -0.427680057 |
| P31930 | UQCRC1 | | 8950100000 | 9843900000 | 11414000000 | 7925300000 | 8430100000 | 8246200000 | 10069333333 | 8200533333 | 0.063827335 | 0.81440678 | -0.296178523 |
| P22695 | UQCRC2 | | 7888900000 | 7270500000 | 8851800000 | 9476400000 | 6727300000 | 9557500000 | 8003733333 | 8587066667 | 0.604041053 | 1.072882655 | 0.101492292 |
| P47985 | UQCRFS1 | | 4146200000 | 3054600000 | 2944000000 | 2110200000 | 2160300000 | 2877500000 | 3381600000 | 2382666667 | 0.093988526 | 0.704597429 | -0.505128884 |

| | | | | | | | | | | | | | |
|---|---|---|---|---|---|---|---|---|---|---|---|---|---|
| P07919 | UQCRH | | 926520000 | 1128600000 | 749430000 | 812370000 | 1107200000 | 1203100000 | 934850000 | 1040890000 | 0.545366973 | 1.113429962 | 0.155010811 |
| O14949 | UQCRQ | | 606230000 | 0 | 564550000 | 701760000 | 812360000 | 623510000 | 390260000 | 712543333.3 | 0.187622705 | 1.825816977 | 0.868542154 |
| P21796 | VDAC1 | | 630650000 | 731850000 | 896710000 | 8791700000 | 10817000000 | 8131800000 | 753070000 | 9246833333 | 0.000471072 | 12.27884969 | 3.618103507 |
| P45880 | VDAC2 | | 3636900000 | 3500300000 | 3084400000 | 10293000000 | 10448000000 | 8295500000 | 3407200000 | 9678833333 | 0.000920183 | 2.840700086 | 1.506246523 |
| Q9Y277 | VDAC3 | | 1869000000 | 1741900000 | 1917800000 | 2317500000 | 2292600000 | 1475700000 | 1842900000 | 2028600000 | 0.545465037 | 1.100765098 | 0.138506633 |
| Q9NRG9 | AAAS | | 114770000 | 76845000 | 133410000 | 98505000 | 105080000 | 63526000 | 108341666.7 | 89037000 | 0.411068854 | 0.821816783 | -0.283111301 |
| P49588 | AARS1 | | 84513000 | 13493000 | 0 | 91121000 | 85120000 | 0 | 32668666.67 | 58747000 | 0.544300286 | 1.798267453 | 0.846607606 |
| P08183 | ABCB1 | | 61641000 | 61136000 | 68210000 | 0 | 61112000 | 118650000 | 63662333.33 | 59920666.67 | 0.918463438 | 0.941226366 | -0.08738636 |
| P33527 | ABCC1 | | 326490000 | 143880000 | 176050000 | 141160000 | 152970000 | 0 | 215473333.3 | 98043333.33 | 0.191112196 | 0.455013768 | -1.136017895 |
| P28288 | ABCD3 | | 208830000 | 208230000 | 311800000 | 75476000 | 116010000 | 113530000 | 242953333.3 | 101672000 | 0.018535967 | 0.418483659 | -1.256756804 |
| P61221 | ABCE1 | | 0 | 0 | 0 | 188320000 | 134350000 | 70689000 | 0 | 131119666.7 | 0.018194198 | N/A | N/A |
| Q8NE71 | ABCF1 | | 80245000 | 54679000 | 53487000 | 186140000 | 155170000 | 191090000 | 62803666.67 | 177466666.7 | 0.001288138 | 2.825737351 | 1.498627375 |
| Q8NFV4 | ABHD11 | | 97896000 | 96465000 | 110040000 | 114570000 | 56915000 | 153090000 | 101467000 | 108191666.7 | 0.823702666 | 1.066274421 | 0.092578784 |
| Q8N2K0 | ABHD12 | | 0 | 0 | 0 | 0 | 68185000 | 55772000 | 0 | 41319000 | 0.120089929 | N/A | N/A |
| Q8IZP0 | ABI1 | | 153440000 | 190700000 | 271100000 | 172250000 | 121010000 | 159870000 | 205080000 | 151043333.3 | 0.228035573 | 0.73650933 | -0.441224294 |
| O14639 | ABLIM1 | | 149490000 | 136930000 | 298350000 | 129200000 | 82065000 | 61293000 | 194923333.3 | 90852666.67 | 0.134537734 | 0.466094362 | -1.101306033 |
| P09110 | ACAA1 | | 248060000 | 371410000 | 164210000 | 541870000 | 498640000 | 466170000 | 261226666.7 | 502226666.7 | 0.019723841 | 1.922570437 | 0.943036455 |
| Q6JQN1 | ACAD10 | | 48028000 | 45393000 | 34264000 | 0 | 0 | 37831000 | 42561666.67 | 12610333.33 | 0.087405092 | 0.296283823 | -1.754948235 |
| P53396 | ACLY | | 487840000 | 286040000 | 363620000 | 150450000 | 149820000 | 150920000 | 379166666.7 | 150396666.7 | 0.017653031 | 0.396650549 | -1.334059547 |
| Q86TX2 | ACOT1 | | 1457200000 | 1418900000 | 1367500000 | 1034700000 | 1004400000 | 1124800000 | 1414533333 | 1054633333 | 0.001273026 | 0.745569799 | -0.423584673 |
| O14734 | ACOT8 | | 0 | 157370000 | 0 | 25925000 | 41157000 | 141660000 | 52456666.67 | 69580666.67 | 0.801651255 | 1.326440872 | 0.407560367 |
| Q15067 | ACOX1 | | 412590000 | 240220000 | 255130000 | 0 | 123550000 | 113730000 | 302646666.7 | 79093333.33 | 0.030167731 | 0.261338855 | -1.936006459 |
| P33121 | ACSL1 | | 336140000 | 142870000 | 153120000 | 237090000 | 234450000 | 235130000 | 210710000 | 235556666.7 | 0.712515793 | 1.117918783 | 0.160815379 |
| O95573 | ACSL3 | | 1228900000 | 1672800000 | 1352200000 | 1331200000 | 1758000000 | 1099900000 | 1417966667 | 1396366667 | 0.930828183 | 0.98476692 | -0.022145795 |
| O60488 | ACSL4 | | 163190000 | 125990000 | 188740000 | 120370000 | 46249000 | 0 | 159306666.7 | 55539666.67 | 0.058401605 | 0.348633663 | -1.520216218 |
| P68133 | ACTA1 | | 1961500000 | 1803800000 | 2847500000 | 2534500000 | 4074500000 | 6439700000 | 2204266667 | 4349566667 | 0.143516705 | 1.973248851 | 0.980572909 |
| P63261 | ACTG1 | | 60745000000 | 59727000000 | 61586000000 | 55073000000 | 56735000000 | 57747000000 | 60686000000 | 56518333333 | 0.011674917 | 0.931324084 | -0.102644809 |
| O96019 | ACTL6A | | 62657000 | 4571500 | 0 | 0 | 0 | 0 | 22409500 | 0 | 0.32877159 | 0 | N/A |
| P12814 | ACTN1 | | 1398600000 | 1313600000 | 1306000000 | 1133500000 | 1151800000 | 947860000 | 1339400000 | 1077720000 | 0.021668351 | 0.804628938 | -0.31360447 |
| O43707 | ACTN4 | | 359500000 | 175350000 | 274360000 | 171960000 | 115110000 | 0 | 269736666.7 | 95690000 | 0.076760193 | 0.354753401 | -1.495111578 |
| P61163 | ACTR1A | | 270510000 | 220870000 | 211840000 | 151340000 | 233900000 | 207090000 | 234406666.7 | 197443333.3 | 0.290820884 | 0.842311083 | -0.247574945 |
| P61160 | ACTR2 | | 183820000 | 212480000 | 188580000 | 197640000 | 183780000 | 84266000 | 194960000 | 155228666.7 | 0.340941724 | 0.796207769 | -0.328783146 |
| P61158 | ACTR3 | | 1508100000 | 1308400000 | 1408900000 | 1179500000 | 1201200000 | 1305700000 | 1408466667 | 1228800000 | 0.061181848 | 0.872438112 | -0.1968753 |
| O14672 | ADAM10 | | 124710000 | 139840000 | 161620000 | 229760000 | 170570000 | 102610000 | 142056666.7 | 167646666.7 | 0.540258394 | 1.180139381 | 0.23895726 |
| P78536 | ADAM17 | | 155280000 | 140600000 | 178780000 | 221460000 | 186140000 | 138190000 | 158220000 | 181930000 | 0.422610309 | 1.149854633 | 0.201451484 |
| P35611 | ADD1 | | 117810000 | 161080000 | 203290000 | 105960000 | 87884000 | 96451000 | 160726666.7 | 96765000 | 0.064262116 | 0.602046953 | -0.732052088 |
| Q9UEY8 | ADD3 | | 139940000 | 174850000 | 157110000 | 0 | 0 | 0 | 157300000 | 0 | 9.83916E-05 | 0 | N/A |
| Q9Y653 | ADGRG1 | | 64646000 | 63026000 | 82617000 | 0 | 0 | 0 | 70096333.33 | 0 | 0.000366202 | 0 | N/A |
| Q9NX46 | ADPRS | | 0 | 0 | 25691000 | 30491000 | 0 | 0 | 8563666.667 | 10163666.67 | 0.909981305 | 1.186835857 | 0.24712042 |
| P30566 | ADSL | | 125140000 | 75570000 | 82114000 | 146850000 | 186470000 | 147040000 | 94274666.67 | 160120000 | 0.031945933 | 1.698441434 | 0.764211471 |
| P55196 | AFDN | | 281010000 | 264180000 | 216440000 | 0 | 144810000 | 0 | 253876666.7 | 48270000 | 0.016761776 | 0.190131691 | -2.394929074 |
| Q9Y4W6 | AFG3L2 | | 1070900000 | 927940000 | 819540000 | 777900000 | 831130000 | 1023800000 | 939460000 | 877610000 | 0.585064006 | 0.934164307 | -0.098251772 |
| Q99943 | AGPAT1 | | 0 | 50157000 | 55878000 | 0 | 35754000 | 0 | 35345000 | 11918000 | 0.334726328 | 0.33719055 | -1.568363989 |
| O00116 | AGPS | | 2298500000 | 2183600000 | 2493300000 | 2764100000 | 3578400000 | 3344100000 | 2325133333 | 3228866667 | 0.024940402 | 1.388680219 | 0.473714418 |
| O00468 | AGRN | | 164770000 | 69243000 | 98543000 | 634000000 | 383410000 | 359870000 | 110852000 | 459093333.3 | 0.019455034 | 4.141497973 | 2.050152683 |
| Q6RW13 | AGTRAP | | 0 | 0 | 198150000 | 155190000 | 229420000 | 299260000 | 66050000 | 227956666.7 | 0.106718409 | 3.451274287 | 1.787129135 |
| P23526 | AHCY | | 0 | 0 | 0 | 0 | 137560000 | 187520000 | 0 | 108360000 | 0.125425537 | N/A | N/A |
| Q09666 | AHNAK | | 36344000000 | 34276000000 | 36873000000 | 42093000000 | 43447000000 | 41361000000 | 35831000000 | 42300333333 | 0.00294764 | 1.180551292 | 0.239460724 |
| Q8IVF2 | AHNAK2 | | 237810000 | 177580000 | 231380000 | 251070000 | 340570000 | 206110000 | 215590000 | 265916666.7 | 0.315456151 | 1.233436925 | 0.302683942 |
| Q96BJ3 | AIDA | | 0 | 29413000 | 0 | 0 | 0 | 44946000 | 9804333.333 | 14982000 | 0.786815241 | 1.52809982 | 0.611738787 |
| Q12904 | AIMP1 | | 477800000 | 480900000 | 370590000 | 377150000 | 463930000 | 389090000 | 443096666.7 | 410056666.7 | 0.506241381 | 0.925433878 | -0.111798181 |
| Q13155 | AIMP2 | | 141590000 | 294590000 | 184540000 | 192940000 | 256130000 | 172930000 | 206906666.7 | 207333333.3 | 0.993846766 | 1.002062121 | 0.002971949 |
| Q02952 | AKAP12 | | 0 | 0 | 40796000 | 683350000 | 290600000 | 112530000 | 13598666.67 | 362160000 | 0.108399061 | 26.63202275 | 4.735090103 |
| Q9Y2D5 | AKAP2 | | 69367000 | 75767000 | 46319000 | 224130000 | 158270000 | 59082000 | 63817666.67 | 147160666.7 | 0.162812909 | 2.305954986 | 1.205364351 |
| Q13740 | ALCAM | | 157950000 | 0 | 224430000 | 119100000 | 145200000 | 152160000 | 127460000 | 138820000 | 0.874172975 | 1.089126 | 0.123170868 |
| Q8IZ83 | ALDH16A1 | | 154540000 | 125350000 | 160160000 | 101750000 | 105180000 | 173810000 | 146683333.3 | 126913333.3 | 0.486713379 | 0.865219861 | -0.208861312 |
| P54886 | ALDH18A1 | | 6735600000 | 6554600000 | 5844500000 | 7914000000 | 8737000000 | 6674200000 | 6378233333 | 7775066667 | 0.101130584 | 1.219000037 | 0.285698169 |

| | | | | | | | | | | | | | |
|---|---|---|---|---|---|---|---|---|---|---|---|---|---|
| P49419 | ALDH7A1 | | 2293000000 | 1743500000 | 1976400000 | 1608500000 | 2076300000 | 1790300000 | 2004300000 | 1825033333 | 0.440423397 | 0.910558965 | -0.13517565 |
| P49189 | ALDH9A1 | | 329200000 | 312170000 | 241230000 | 342360000 | 344450000 | 294020000 | 294200000 | 326943333.3 | 0.358306143 | 1.11129617 | 0.152243359 |
| P04075 | ALDOA | | 11132000000 | 11189000000 | 9771400000 | 8710900000 | 8583700000 | 6652400000 | 10697466667 | 7982333333 | 0.028657518 | 0.746189129 | -0.422386753 |
| P09972 | ALDOC | | 0 | 0 | 0 | 70976000 | 76114000 | 79093000 | 0 | 75394333.33 | 5.82646E-06 | N/A | N/A |
| Q86V81 | ALYREF | | 263930000 | 235170000 | 140050000 | 159300000 | 168420000 | 220670000 | 213050000 | 182796666.7 | 0.511481304 | 0.857998905 | -0.220952289 |
| Q9UKV5 | AMFR | | 0 | 56197000 | 0 | 68593000 | 0 | 0 | 18732333.33 | 22864333.33 | 0.895580119 | 1.22058117 | 0.287568239 |
| Q86SJ2 | AMIGO2 | | 38662000 | 0 | 117480000 | 103380000 | 306050000 | 364520000 | 52047333.33 | 257983333.3 | 0.075572983 | 4.956706075 | 2.309381712 |
| Q01433 | AMPD2 | | 0 | 0 | 0 | 28078000 | 0 | 44567000 | 0 | 24215000 | 0.136191764 | N/A | N/A |
| Q9NQW6 | ANLN | | 104700000 | 79646000 | 0 | 0 | 0 | 0 | 61448666.67 | 0 | 0.123411276 | 0 | N/A |
| P39687 | ANP32A | | 65311000 | 54388000 | 49291000 | 79306000 | 63219000 | 55635000 | 56330000 | 66053333.33 | 0.312867154 | 1.172613764 | 0.229727896 |
| P04083 | ANXA1 | | 246630000 | 425500000 | 493600000 | 860250000 | 848960000 | 1323200000 | 388576666.7 | 1010803333 | 0.02270725 | 2.601297041 | 1.37923115 |
| P07355 | ANXA2 | | 44032000000 | 40393000000 | 41499000000 | 54556000000 | 59182000000 | 68571000000 | 41974666667 | 60769666667 | 0.011594908 | 1.447770084 | 0.53383251 |
| P08758 | ANXA5 | | 203200000 | 167250000 | 191720000 | 352680000 | 738400000 | 605180000 | 187390000 | 565420000 | 0.029170762 | 3.017343508 | 1.593278949 |
| P08133 | ANXA6 | | 80342000 | 74680000 | 82139000 | 105420000 | 137080000 | 127420000 | 79053666.67 | 123306666.7 | 0.0100802 | 1.559784282 | 0.641346518 |
| P20073 | ANXA7 | | 0 | 147160000 | 0 | 155280000 | 193550000 | 162250000 | 49053333.33 | 170360000 | 0.073976783 | 3.472954607 | 1.796163554 |
| O43747 | AP1G1 | | 0 | 0 | 0 | 17731000 | 65733000 | 0 | 0 | 27821333.33 | 0.229457434 | N/A | N/A |
| Q9BXS5 | AP1M1 | | 0 | 0 | 0 | 59813000 | 0 | 100290000 | 0 | 53367666.67 | 0.140892071 | N/A | N/A |
| O95782 | AP2A1 | | 1500700000 | 1962700000 | 1305500000 | 1242000000 | 1681200000 | 1733400000 | 1589633333 | 1552200000 | 0.888002013 | 0.976451593 | -0.03437957 |
| P63010 | AP2B1 | | 501280000 | 526860000 | 619100000 | 784920000 | 716430000 | 572170000 | 549080000 | 691173333.3 | 0.120405284 | 1.258784391 | 0.332031194 |
| Q96CW1 | AP2M1 | | 216430000 | 175450000 | 268430000 | 186370000 | 281540000 | 172600000 | 220103333.3 | 213503333.3 | 0.886885793 | 0.970014084 | -0.0439224 |
| O14617 | AP3D1 | | 0 | 0 | 0 | 32186000 | 25920000 | 0 | 0 | 19368666.67 | 0.120721442 | N/A | N/A |
| O43299 | AP5Z1 | | 0 | 0 | 0 | 35262000 | 29594000 | 0 | 0 | 21618666.67 | 0.119145596 | N/A | N/A |
| Q9BZZ5 | API5 | | 105780000 | 79681000 | 0 | 76229000 | 82239000 | 0 | 61820333.33 | 52822666.67 | 0.838528195 | 0.854454575 | -0.226924298 |
| Q06481 | APLP2 | | 33293000 | 24524000 | 52715000 | 106230000 | 89643000 | 47246000 | 36844000 | 81039666.67 | 0.085373378 | 2.199534976 | 1.137198543 |
| Q9HDC9 | APMAP | | 1028800000 | 1200100000 | 1334000000 | 476100000 | 788010000 | 825360000 | 1187633333 | 696490000 | 0.025637611 | 0.586452047 | -0.769914947 |
| Q9BQE5 | APOL2 | | 64418000 | 115310000 | 118500000 | 107740000 | 153780000 | 88194000 | 99409333.33 | 116571333.3 | 0.547774274 | 1.172639725 | 0.229759837 |
| P05067 | APP | | 112150000 | 76825000 | 70931000 | 100420000 | 149080000 | 150400000 | 86635333.33 | 133300000 | 0.089147531 | 1.538633198 | 0.621649342 |
| P48444 | ARCN1 | | 53069000 | 0 | 0 | 50848000 | 0 | 70260000 | 17689666.67 | 40369333.33 | 0.45462542 | 2.282085587 | 1.190352899 |
| P84077 | ARF1 | | 0 | 0 | 0 | 40219000 | 0 | 76793000 | 0 | 39004000 | 0.153432533 | N/A | N/A |
| P18085 | ARF4 | | 0 | 0 | 0 | 0 | 25194000 | 66714000 | 0 | 30636000 | 0.190349565 | N/A | N/A |
| P62330 | ARF6 | | 43398000 | 136340000 | 122530000 | 109910000 | 185070000 | 135160000 | 100756000 | 143380000 | 0.306769653 | 1.423041804 | 0.508978044 |
| Q8N6T3 | ARFGAP1 | | 208540000 | 198410000 | 264260000 | 443710000 | 554510000 | 630610000 | 223736666.7 | 542943333.3 | 0.005314267 | 2.426706992 | 1.278999924 |
| P53365 | ARFIP2 | | 0 | 0 | 0 | 74475000 | 0 | 66451000 | 0 | 46975333.33 | 0.117404407 | N/A | N/A |
| Q07960 | ARHGAP1 | | 0 | 57269000 | 77505000 | 48798000 | 58569000 | 220840000 | 44924666.67 | 109402333.3 | 0.346065433 | 2.435239735 | 1.284063804 |
| P52565 | ARHGDIA | | 0 | 0 | 164110000 | 326280000 | 580140000 | 553890000 | 54703333.33 | 486770000 | 0.011373942 | 8.898360856 | 3.153539605 |
| Q9NXU5 | ARL15 | | 102890000 | 123140000 | 288190000 | 0 | 0 | 0 | 171406666.7 | 0 | 0.04320614 | 0 | N/A |
| Q15041 | ARL6IP1 | | 149980000 | 153590000 | 214290000 | 207510000 | 108560000 | 0 | 172620000 | 105356666.7 | 0.34887869 | 0.610338702 | -0.71231802 |
| O75915 | ARL6IP5 | | 250780000 | 295210000 | 278650000 | 261920000 | 368710000 | 523900000 | 274880000 | 384843333.3 | 0.227211851 | 1.40004123 | 0.485469314 |
| Q96BM9 | ARL8A | | 0 | 0 | 0 | 0 | 60125000 | 67539000 | 0 | 42554666.67 | 0.117456274 | N/A | N/A |
| Q9NVJ2 | ARL8B | | 183930000 | 230360000 | 223360000 | 755590000 | 780010000 | 651660000 | 212550000 | 729086666.7 | 0.000249227 | 3.430188975 | 1.778288059 |
| Q8N2F6 | ARMC10 | | 161520000 | 0 | 0 | 0 | 0 | 109160000 | 53840000 | 36386666.67 | 0.801532507 | 0.675829619 | -0.565268516 |
| Q9UH62 | ARMCX3 | | 0 | 0 | 0 | 40080000 | 0 | 37575000 | 0 | 25885000 | 0.116530251 | N/A | N/A |
| Q92747 | ARPC1A | | 0 | 0 | 106880000 | 0 | 97833000 | 0 | 35626666.67 | 32611000 | 0.953209334 | 0.915353668 | -0.127598826 |
| O15143 | ARPC1B | | 161490000 | 209070000 | 172370000 | 227750000 | 210850000 | 161890000 | 180976666.7 | 200163333.3 | 0.476268868 | 1.10601735 | 0.145374018 |
| O15144 | ARPC2 | | 100140000 | 137080000 | 142480000 | 223050000 | 174340000 | 89850000 | 126566666.7 | 162413333.3 | 0.432571815 | 1.283223598 | 0.359772577 |
| P59998 | ARPC4 | | 702390000 | 861960000 | 799580000 | 369650000 | 527940000 | 695370000 | 787976666.7 | 530986666.7 | 0.070413725 | 0.673860901 | -0.569477275 |
| Q9BPX5 | ARPC5L | | 64694000 | 196020000 | 112020000 | 108120000 | 122910000 | 0 | 124244666.7 | 77010000 | 0.43537469 | 0.619825398 | -0.690066222 |
| O00192 | ARVCF | | 291980000 | 332070000 | 443550000 | 210610000 | 204210000 | 202440000 | 355866666.7 | 205753333.3 | 0.029775686 | 0.578175347 | -0.790421001 |
| Q12797 | ASPH | | 1247800000 | 1174000000 | 1224400000 | 518900000 | 707750000 | 910720000 | 1215400000 | 712456666.7 | 0.012010606 | 0.586191103 | -0.770557023 |
| Q8NBU5 | ATAD1 | | 205650000 | 275930000 | 226200000 | 292980000 | 217200000 | 219190000 | 235926666.7 | 243123333.3 | 0.835656016 | 1.030503829 | 0.043349865 |
| Q9NVI7 | ATAD3A | | 412710000 | 479820000 | 625140000 | 492140000 | 475380000 | 708290000 | 505890000 | 558603333.3 | 0.618323935 | 1.1041992 | 0.143000461 |
| Q5T9A4 | ATAD3B | | 800600000 | 1103200000 | 838300000 | 1233300000 | 1005100000 | 867030000 | 914033333.3 | 1035143333 | 0.444946792 | 1.132500638 | 0.179511863 |
| P31939 | ATIC | | 597850000 | 373970000 | 418140000 | 676290000 | 721940000 | 864120000 | 463320000 | 754116666.7 | 0.030655525 | 1.627636767 | 0.702778775 |
| Q8NHH9 | ATL2 | | 277000000 | 176080000 | 168510000 | 123120000 | 46671000 | 143760000 | 207196666.7 | 104517000 | 0.088292684 | 0.504433791 | -0.987263173 |
| Q6DD88 | ATL3 | | 115180000 | 142010000 | 132970000 | 0 | 81937000 | 127310000 | 130053333.3 | 69749000 | 0.188429056 | 0.536310744 | -0.898858939 |
| P05023 | ATP1A1 | | 7437900000 | 7194300000 | 8869300000 | 8236800000 | 7418500000 | 5223700000 | 7833833333 | 6959666667 | 0.448025952 | 0.888411378 | -0.170700225 |
| P05026 | ATP1B1 | | 603250000 | 637250000 | 887890000 | 458030000 | 743930000 | 988420000 | 709463333.3 | 730126666.7 | 0.912990035 | 1.029125301 | 0.041418648 |

| | | | | | | | | | | | | | |
|---|---|---|---|---|---|---|---|---|---|---|---|---|---|
| P54709 | ATP1B3 | | 125620000 | 206160000 | 296010000 | 483640000 | 388420000 | 400450000 | 209263333.3 | 424170000 | 0.020281606 | 2.026967617 | 1.01932304 |
| P16615 | ATP2A2 | | 1668400000 | 1597600000 | 1723300000 | 382660000 | 468200000 | 494770000 | 1663100000 | 448543333.3 | 1.66051E-05 | 0.269703165 | -1.890555644 |
| P20020 | ATP2B1 | | 0 | 86537000 | 66203000 | 104960000 | 99581000 | 127200000 | 50913333.33 | 110580333.3 | 0.095489224 | 2.171932696 | 1.118979398 |
| P23634 | ATP2B4 | | 312830000 | 389830000 | 292590000 | 227620000 | 203100000 | 241510000 | 331750000 | 224076666.7 | 0.027304615 | 0.675438332 | -0.566104038 |
| Q93050 | ATP6V0A1 | | 179120000 | 180340000 | 135550000 | 205410000 | 266490000 | 382140000 | 165003333.3 | 284680000 | 0.090473402 | 1.725298479 | 0.786845972 |
| P61421 | ATP6V0D1 | | 340770000 | 343600000 | 382120000 | 420240000 | 672250000 | 688730000 | 355496666.7 | 593740000 | 0.053512294 | 1.670170372 | 0.739995278 |
| P38606 | ATP6V1A | | 579460000 | 670580000 | 773530000 | 903960000 | 837850000 | 1100900000 | 674523333.3 | 947570000 | 0.047894825 | 1.404799439 | 0.490364173 |
| P21281 | ATP6V1B2 | | 660190000 | 637090000 | 576360000 | 780830000 | 955770000 | 805800000 | 624546666.7 | 847466666.7 | 0.020644391 | 1.356930894 | 0.440347249 |
| P21283 | ATP6V1C1 | | 0 | 153830000 | 0 | 96230000 | 113950000 | 211110000 | 51276666.67 | 140430000 | 0.226800658 | 2.738672561 | 1.453476786 |
| Q9Y5K8 | ATP6V1D | | 124180000 | 149100000 | 0 | 135690000 | 93485000 | 0 | 91093333.33 | 76391666.67 | 0.821691071 | 0.838608753 | -0.253930207 |
| P36543 | ATP6V1E1 | | 209400000 | 177200000 | 148270000 | 350700000 | 221250000 | 105990000 | 178290000 | 225980000 | 0.548451558 | 1.267485557 | 0.341969308 |
| O75348 | ATP6V1G1 | | 555990000 | 446810000 | 474910000 | 514110000 | 401410000 | 332220000 | 492570000 | 415913333.3 | 0.2859219 | 0.844374065 | -0.244045828 |
| Q9UI12 | ATP6V1H | | 0 | 0 | 0 | 100280000 | 123320000 | 85981000 | 0 | 103193666.7 | 0.000688733 | N/A | N/A |
| Q9Y679 | AUP1 | | 360920000 | 289530000 | 240200000 | 163470000 | 210740000 | 0 | 296883333.3 | 124736666.7 | 0.077347627 | 0.42015382 | -1.251010493 |
| P30530 | AXL | | 0 | 36575000 | 59550000 | 0 | 0 | 0 | 32041666.67 | 0 | 0.138323882 | 0 | N/A |
| O94766 | B3GAT3 | | 0 | 18675000 | 47285000 | 80226000 | 83421000 | 0 | 21986666.67 | 54549000 | 0.346661017 | 2.481003639 | 1.310923851 |
| Q92934 | BAD | | 41784000 | 0 | 30356000 | 0 | 0 | 0 | 24046666.67 | 0 | 0.126001418 | 0 | N/A |
| O95816 | BAG2 | | 1188500000 | 959790000 | 1020700000 | 782150000 | 1094600000 | 805730000 | 1056330000 | 894160000 | 0.252931185 | 0.8464779 | -0.240455693 |
| Q9UL15 | BAG5 | | 0 | 0 | 0 | 37396000 | 0 | 56205000 | 0 | 31200333.33 | 0.131924025 | N/A | N/A |
| Q9UQB8 | BAIAP2 | | 486230000 | 509420000 | 636510000 | 277160000 | 241410000 | 258740000 | 544053333.3 | 259103333.3 | 0.00398706 | 0.476246201 | -1.070220509 |
| Q9UHR4 | BAIAP2L1 | | 250840000 | 285570000 | 322350000 | 98048000 | 183080000 | 0 | 286253333.3 | 93709333.33 | 0.027505726 | 0.327365038 | -1.611027842 |
| Q16611 | BAK1 | | 138360000 | 122520000 | 0 | 0 | 0 | 0 | 86960000 | 0 | 0.117580833 | 0 | N/A |
| O75531 | BANF1 | | 0 | 0 | 104080000 | 0 | 0 | 151170000 | 34693333.33 | 50390000 | 0.810164789 | 1.45244043 | 0.538478995 |
| P50895 | BCAM | | 470050000 | 542200000 | 817510000 | 306120000 | 0 | 279650000 | 609920000 | 195256666.7 | 0.045222509 | 0.320134881 | -1.643248217 |
| P51572 | BCAP31 | | 471370000 | 412470000 | 269760000 | 458610000 | 581120000 | 223450000 | 384533333.3 | 421060000 | 0.777459539 | 1.094989598 | 0.130917165 |
| O75934 | BCAS2 | | 267360000 | 189220000 | 0 | 0 | 22808000 | 138650000 | 152193333.3 | 53819333.33 | 0.336889319 | 0.353624776 | -1.499708739 |
| Q9BXK5 | BCL2L13 | | 0 | 0 | 0 | 168520000 | 142660000 | 194290000 | 0 | 168490000 | 0.000348963 | N/A | N/A |
| P55957 | BID | | 0 | 0 | 184440000 | 0 | 216770000 | 0 | 61480000 | 72256666.67 | 0.915035068 | 1.175287356 | 0.233013537 |
| Q6QNY1 | BLOC1S2 | | 247010000 | 126850000 | 219460000 | 191700000 | 0 | 228830000 | 197773333.3 | 140176666.7 | 0.509750325 | 0.708774354 | -0.496601691 |
| O60238 | BNIP3L | | 0 | 0 | 0 | 224270000 | 135080000 | 73906000 | 0 | 144418666.7 | 0.029708674 | N/A | N/A |
| Q8TDN6 | BRIX1 | | 0 | 0 | 0 | 56534000 | 54223000 | 0 | 0 | 36919000 | 0.116289662 | N/A | N/A |
| P35613 | BSG | | 0 | 373860000 | 273230000 | 186470000 | 207930000 | 226020000 | 215696666.7 | 206806666.7 | 0.940692282 | 0.958784713 | -0.060721188 |
| Q7L1Q6 | BZW1 | | 255040000 | 38725000 | 0 | 167410000 | 45038000 | 279850000 | 97921666.67 | 164099333.3 | 0.560500782 | 1.675822511 | 0.744869359 |
| Q9Y6E2 | BZW2 | | 0 | 0 | 0 | 58618000 | 52323000 | 0 | 0 | 36980333.33 | 0.117395575 | N/A | N/A |
| Q99622 | C12orf57 | | 38447000 | 72558000 | 77149000 | 0 | 0 | 0 | 62718000 | 0 | 0.006802596 | 0 | N/A |
| Q9UFG5 | C19orf25 | | 61095000 | 0 | 49852000 | 0 | 0 | 47662000 | 36982333.33 | 15887333.33 | 0.439395014 | 0.429592508 | -1.218959261 |
| Q6P1X6 | C8orf82 | | 1259300000 | 889050000 | 596730000 | 615710000 | 750000000 | 1110800000 | 915026666.7 | 825503333.3 | 0.730273227 | 0.902163143 | -0.148539748 |
| Q9HB71 | CACYBP | | 584310000 | 569130000 | 0 | 513290000 | 397050000 | 474970000 | 384480000 | 461770000 | 0.712500051 | 1.201024761 | 0.264265894 |
| P27708 | CAD | | 1298300000 | 1276600000 | 1535400000 | 1397700000 | 1396800000 | 1514600000 | 1370100000 | 1436366667 | 0.509689247 | 1.048366299 | 0.068142883 |
| Q9BY67 | CADM1 | | 0 | 0 | 110510000 | 74723000 | 102700000 | 0 | 36836666.67 | 59141000 | 0.665838735 | 1.605492716 | 0.683016119 |
| Q8N126 | CADM3 | | 0 | 95821000 | 132330000 | 0 | 48382000 | 109180000 | 76050333.33 | 52520666.67 | 0.665765737 | 0.69060403 | -0.534069342 |
| Q05682 | CALD1 | | 700310000 | 852520000 | 1052500000 | 305320000 | 350810000 | 166330000 | 868443333.3 | 274153333.3 | 0.006891598 | 0.315683618 | -1.6634487 |
| P0DP25 | CALM3 | | 9408800000 | 8111100000 | 7881100000 | 5606400000 | 5717700000 | 5360100000 | 8467000000 | 5561400000 | 0.00396798 | 0.656832408 | -0.606402783 |
| P27797 | CALR | | 10851000000 | 11167000000 | 12957000000 | 15350000000 | 15040000000 | 14214000000 | 11658333333 | 14868000000 | 0.012174747 | 1.275310936 | 0.350849037 |
| O43852 | CALU | | 1434000000 | 883550000 | 1346600000 | 1519800000 | 1427600000 | 1576100000 | 1221383333 | 1507833333 | 0.179318345 | 1.234529154 | 0.303960907 |
| Q86VP6 | CAND1 | | 0 | 0 | 0 | 242700000 | 289260000 | 0 | 0 | 177320000 | 0.11915471 | N/A | N/A |
| Q8WVQ1 | CANT1 | | 158200000 | 153820000 | 138510000 | 169880000 | 250360000 | 204710000 | 150176666.7 | 208316666.7 | 0.073000646 | 1.38714403 | 0.472117594 |
| P27824 | CANX | | 1852300000 | 1973600000 | 2374800000 | 2976400000 | 3334000000 | 3507800000 | 2066900000 | 3272733333 | 0.00559674 | 1.583401874 | 0.663027464 |
| Q01518 | CAP1 | | 329840000 | 209060000 | 72134000 | 1175400000 | 1117800000 | 1039500000 | 203678000 | 1110900000 | 0.000421074 | 5.454197311 | 2.447366892 |
| P17655 | CAPN2 | | 28001000 | 167220000 | 21667000 | 181370000 | 172980000 | 182640000 | 72296000 | 178996666.7 | 0.088435133 | 2.475886172 | 1.307944988 |
| P04632 | CAPNS1 | | 92508000 | 88663000 | 99455000 | 82177000 | 126890000 | 142970000 | 93542000 | 117345666.7 | 0.266717393 | 1.254470363 | 0.327078387 |
| Q14444 | CAPRIN1 | | 388700000 | 155620000 | 319610000 | 295250000 | 204380000 | 266120000 | 287976666.7 | 255250000 | 0.681676624 | 0.886356534 | -0.174040961 |
| P52907 | CAPZA1 | | 872530000 | 827390000 | 950760000 | 0 | 0 | 0 | 883560000 | 0 | 1.64235E-05 | 0 | N/A |
| P47755 | CAPZA2 | | 0 | 0 | 141490000 | 0 | 226050000 | 0 | 47163333.33 | 75350000 | 0.767039866 | 1.597639409 | 0.675941825 |
| P47756 | CAPZB | | 792440000 | 632170000 | 1120700000 | 311540000 | 360400000 | 667580000 | 848436666.7 | 446506666.7 | 0.091671832 | 0.526269885 | -0.926125255 |
| Q86X55 | CARM1 | | 0 | 118920000 | 0 | 98481000 | 169220000 | 148990000 | 39640000 | 138897000 | 0.091438844 | 3.503960646 | 1.808986572 |
| O14936 | CASK | | 0 | 129050000 | 125350000 | 116270000 | 109800000 | 96294000 | 84800000 | 107454666.7 | 0.624755167 | 1.267154088 | 0.341591969 |

| | | | | | | | | | | | | | |
|---|---|---|---|---|---|---|---|---|---|---|---|---|---|
| Q03135 | CAV1 | | 513840000 | 413410000 | 561220000 | 185600000 | 153210000 | 165830000 | 496156666.7 | 168213333.3 | 0.001819462 | 0.339032698 | -1.560503674 |
| Q6NZI2 | CAVIN1 | | 2750200000 | 2036900000 | 2109500000 | 529200000 | 635860000 | 442120000 | 2298866667 | 535726666.7 | 0.001647093 | 0.233039469 | -2.101353777 |
| Q13185 | CBX3 | | 0 | 0 | 45273000 | 0 | 29897000 | 0 | 15091000 | 9965666.667 | 0.79092686 | 0.660371524 | -0.598650184 |
| Q8N163 | CCAR2 | | 64615000 | 107010000 | 35646000 | 135780000 | 102270000 | 150490000 | 69090333.33 | 129513333.3 | 0.074226104 | 1.874550709 | 0.906544853 |
| Q96CT7 | CCDC124 | | 0 | 69210000 | 0 | 91782000 | 93953000 | 0 | 23070000 | 61911666.67 | 0.371354145 | 2.683643982 | 1.424193293 |
| O60826 | CCDC22 | | 84315000 | 0 | 45875000 | 75459000 | 68484000 | 88739000 | 43396666.67 | 77560666.67 | 0.24487192 | 1.787249405 | 0.837740972 |
| Q96A33 | CCDC47 | | 163460000 | 236180000 | 73948000 | 677860000 | 507170000 | 600620000 | 157862666.7 | 595216666.7 | 0.003020338 | 3.77047138 | 1.914744899 |
| O00622 | CCN1 | | 0 | 0 | 26928000 | 0 | 41570000 | 0 | 8976000 | 13856666.67 | 0.782230349 | 1.543746286 | 0.626435666 |
| P78371 | CCT2 | | 13528000000 | 12360000000 | 15091000000 | 6434500000 | 8258500000 | 6884300000 | 13659666667 | 7192433333 | 0.002556887 | 0.526545304 | -0.92537043 |
| P49368 | CCT3 | | 10931000000 | 10844000000 | 11538000000 | 6966200000 | 6515300000 | 5407100000 | 11104333333 | 6296200000 | 0.000716825 | 0.567003872 | -0.818569507 |
| P50991 | CCT4 | | 9957500000 | 8574200000 | 8716900000 | 5256500000 | 5456500000 | 5438400000 | 9082866667 | 5383800000 | 0.001132999 | 0.592742379 | -0.754522885 |
| P48643 | CCT5 | | 8639800000 | 10344000000 | 7542400000 | 4398200000 | 4585100000 | 4355100000 | 8842066667 | 4446133333 | 0.005794395 | 0.502838703 | -0.991832398 |
| P40227 | CCT6A | | 10429000000 | 9973700000 | 8549500000 | 4555700000 | 4602700000 | 5459500000 | 9650733333 | 4872633333 | 0.001697723 | 0.504897728 | -0.98593691 |
| Q99832 | CCT7 | | 8728500000 | 8787200000 | 9474900000 | 5528200000 | 4770600000 | 4688600000 | 8996866667 | 4995800000 | 0.000368631 | 0.555282209 | -0.848706921 |
| P50990 | CCT8 | | 11255000000 | 11516000000 | 11369000000 | 7941300000 | 8038400000 | 6375000000 | 11380000000 | 7451566667 | 0.001954051 | 0.654794962 | -0.610884873 |
| P86791 | CCZ1 | | 0 | 0 | 0 | 291740000 | 341700000 | 486090000 | 0 | 373176666.7 | 0.003053341 | N/A | N/A |
| Q6YHK3 | CD109 | | 253320000 | 86990000 | 170500000 | 190380000 | 209760000 | 474060000 | 170270000 | 291400000 | 0.306161023 | 1.711399542 | 0.77517661 |
| Q9Y5K6 | CD2AP | | 0 | 0 | 103490000 | 204650000 | 173620000 | 149790000 | 34496666.67 | 176020000 | 0.020358098 | 5.102521983 | 2.351210493 |
| P25942 | CD40 | | 203050000 | 138150000 | 185360000 | 0 | 197370000 | 198920000 | 175520000 | 132096666.7 | 0.562379573 | 0.752601793 | -0.410041369 |
| P16070 | CD44 | | 919510000 | 1003400000 | 846140000 | 2155700000 | 1909800000 | 1339600000 | 923016666.7 | 1801700000 | 0.023318503 | 1.951969087 | 0.964930205 |
| P13987 | CD59 | | 211350000 | 391900000 | 574510000 | 1332200000 | 825350000 | 361100000 | 392586666.7 | 839550000 | 0.209734286 | 2.138508694 | 1.096605073 |
| P21926 | CD9 | | 379800000 | 695540000 | 372900000 | 219850000 | 434340000 | 476590000 | 482746666.7 | 376926666.7 | 0.470229392 | 0.780796001 | -0.356982432 |
| Q16543 | CDC37 | | 118510000 | 90227000 | 25673000 | 117670000 | 34527000 | 102470000 | 78136666.67 | 84889000 | 0.865945078 | 1.086416962 | 0.119577909 |
| P60953 | CDC42 | | 644460000 | 877150000 | 694880000 | 197950000 | 410640000 | 320820000 | 738830000 | 309803333.3 | 0.010224146 | 0.419316126 | -1.253889781 |
| P55290 | CDH13 | | 97299000 | 109340000 | 94174000 | 231850000 | 171570000 | 207730000 | 100271000 | 203716666.7 | 0.004651974 | 2.031660866 | 1.022659601 |
| P19022 | CDH2 | | 1413800000 | 1249100000 | 1370300000 | 753340000 | 800660000 | 824680000 | 1344400000 | 792893333.3 | 0.000501318 | 0.589774869 | -0.761763747 |
| Q96JB5 | CDK5RAP3 | | 99626000 | 116600000 | 110000000 | 173770000 | 190720000 | 131920000 | 108742000 | 165470000 | 0.03539073 | 1.521675158 | 0.60566041 |
| Q5VV42 | CDKAL1 | | 76083000 | 23027000 | 85689000 | 35347000 | 0 | 0 | 61599666.67 | 11782333.33 | 0.093917902 | 0.19127268 | -2.38629727 |
| Q9UKY7 | CDV3 | | 0 | 0 | 37758000 | 91396000 | 41462000 | 43788000 | 12586000 | 58882000 | 0.087587531 | 4.678372795 | 2.226006827 |
| P23528 | CFL1 | | 4199600000 | 3579200000 | 2971000000 | 3956000000 | 2681900000 | 3145900000 | 3583266667 | 3261266667 | 0.565131008 | 0.910137863 | -0.135843001 |
| Q9Y6H1 | CHCHD2 | | 426420000 | 428030000 | 622300000 | 1597300000 | 1357700000 | 892240000 | 492250000 | 1282413333 | 0.021926034 | 2.605207381 | 1.381398219 |
| Q9BWS9 | CHID1 | | 0 | 0 | 0 | 68746000 | 46328000 | 0 | 0 | 38358000 | 0.130988369 | N/A | N/A |
| Q9UHD1 | CHORDC1 | | 179030000 | 162430000 | 144830000 | 231520000 | 249860000 | 333450000 | 162096666.7 | 271610000 | 0.029106762 | 1.675605092 | 0.744682173 |
| Q99653 | CHP1 | | 213340000 | 203280000 | 172210000 | 309480000 | 305200000 | 309790000 | 196276666.7 | 308156666.7 | 0.000853254 | 1.570011718 | 0.650775327 |
| Q14011 | CIRBP | | 109070000 | 51742000 | 0 | 0 | 0 | 0 | 53604000 | 0 | 0.164020527 | 0 | N/A |
| Q9NZ45 | CISD1 | | 486430000 | 489980000 | 555330000 | 230240000 | 475990000 | 388890000 | 510580000 | 365040000 | 0.125571864 | 0.714951624 | -0.484082468 |
| Q8N5K1 | CISD2 | | 489450000 | 414100000 | 443260000 | 440560000 | 0 | 403990000 | 448936666.7 | 281516666.7 | 0.306233098 | 0.627074346 | -0.673291596 |
| O14578 | CIT | | 86598000 | 41257000 | 0 | 0 | 0 | 0 | 42618333.33 | 0 | 0.163549804 | 0 | N/A |
| Q07065 | CKAP4 | | 14254000000 | 13203000000 | 14246000000 | 13147000000 | 12289000000 | 12124000000 | 13901000000 | 12520000000 | 0.042871387 | 0.900654629 | -0.150954108 |
| Q14008 | CKAP5 | | 500230000 | 466430000 | 528720000 | 300990000 | 395320000 | 422420000 | 498460000 | 372910000 | 0.037492502 | 0.748124223 | -0.418650252 |
| Q7Z460 | CLASP1 | | 0 | 37939000 | 0 | 44726000 | 88263000 | 0 | 12646333.33 | 44329666.67 | 0.327772006 | 3.505337515 | 1.809553362 |
| P51797 | CLCN6 | | 0 | 0 | 0 | 42452000 | 48783000 | 29901000 | 0 | 40378666.67 | 0.001894321 | N/A | N/A |
| P51798 | CLCN7 | | 0 | 0 | 0 | 127960000 | 133240000 | 136690000 | 0 | 132630000 | 8.03246E-07 | N/A | N/A |
| Q9Y240 | CLEC11A | | 0 | 76789000 | 0 | 123020000 | 120670000 | 0 | 25596333.33 | 81230000 | 0.311030911 | 3.173501413 | 1.666075485 |
| O00299 | CLIC1 | | 218100000 | 294340000 | 265350000 | 615890000 | 722600000 | 698930000 | 259263333.3 | 679140000 | 0.000432599 | 2.619498836 | 1.389290821 |
| Q9Y696 | CLIC4 | | 174320000 | 0 | 112270000 | 103340000 | 112520000 | 139050000 | 95530000 | 118303333.3 | 0.684733284 | 1.238389337 | 0.308464954 |
| Q9H078 | CLPB | | 586530000 | 610180000 | 649000000 | 329290000 | 394330000 | 328860000 | 615236666.7 | 350826666.7 | 0.000737479 | 0.570230426 | -0.810383075 |
| O96005 | CLPTM1 | | 105550000 | 75658000 | 112130000 | 0 | 0 | 0 | 97779333.33 | 0 | 0.000955708 | 0 | N/A |
| P09496 | CLTA | | 179330000 | 224350000 | 204910000 | 199620000 | 233720000 | 148910000 | 202863333.3 | 194083333.3 | 0.768524202 | 0.956719631 | -0.063831894 |
| Q00610 | CLTC | | 5097400000 | 5528800000 | 5337200000 | 6828400000 | 7109000000 | 6314500000 | 5321133333 | 6750633333 | 0.005635356 | 1.268645777 | 0.343289306 |
| P10909 | CLU | | 1127000000 | 1141100000 | 1014800000 | 2731800000 | 2433100000 | 2416500000 | 1094300000 | 2527133333 | 0.000200208 | 2.309360626 | 1.20749348 |
| P62633 | CNBP | | 49557000 | 0 | 84429000 | 152150000 | 115600000 | 46088000 | 44662000 | 104612666.7 | 0.204548016 | 2.342319347 | 1.227937783 |
| Q99439 | CNN2 | | 321810000 | 263080000 | 248900000 | 209280000 | 195650000 | 155280000 | 277930000 | 186736666.7 | 0.02977135 | 0.671883808 | -0.573716333 |
| Q15417 | CNN3 | | 0 | 142260000 | 0 | 101700000 | 193120000 | 273440000 | 47420000 | 189420000 | 0.107340678 | 3.994517081 | 1.998021099 |
| Q6P4Q7 | CNNM4 | | 0 | 0 | 0 | 54653000 | 0 | 65450000 | 0 | 40034333.33 | 0.119321132 | N/A | N/A |
| P09543 | CNP | | 691450000 | 665900000 | 762310000 | 506440000 | 523170000 | 443470000 | 706553333.3 | 491026666.7 | 0.004626027 | 0.694960513 | -0.524997088 |
| Q9Y2B0 | CNPY2 | | 833670000 | 617790000 | 714050000 | 1041800000 | 1207500000 | 1349300000 | 721836666.7 | 1199533333 | 0.011703942 | 1.661779442 | 0.732728915 |

| | | | | | | | | | | | | | |
|---|---|---|---|---|---|---|---|---|---|---|---|---|---|
| Q9BT09 | CNPY3 | | 258790000 | 335210000 | 380280000 | 273090000 | 300170000 | 252240000 | 324760000 | 275166666.7 | 0.262684258 | 0.847292359 | -0.239068235 |
| Q96JB2 | COG3 | | 0 | 0 | 0 | 0 | 90361000 | 264880000 | 0 | 118413666.7 | 0.202377261 | N/A | N/A |
| P08572 | COL4A2 | | 0 | 0 | 0 | 264840000 | 209270000 | 0 | 0 | 158036666.7 | 0.121552159 | N/A | N/A |
| P12109 | COL6A1 | | 286710000 | 353940000 | 292410000 | 1787000000 | 1752400000 | 1911900000 | 311020000 | 1817100000 | 9.13104E-06 | 5.842389557 | 2.546558557 |
| P12110 | COL6A2 | | 106240000 | 111650000 | 0 | 0 | 198840000 | 219740000 | 72630000 | 139526666.7 | 0.444244485 | 1.921061086 | 0.941903395 |
| Q8NBJ5 | COLGALT1 | | 859620000 | 995360000 | 845020000 | 516860000 | 621550000 | 836360000 | 900000000 | 658256666.7 | 0.083769347 | 0.731396296 | -0.451274774 |
| Q9Y6G5 | COMMD10 | | 107270000 | 70201000 | 87346000 | 62418000 | 0 | 124390000 | 88272333.33 | 62269333.33 | 0.525915471 | 0.70542299 | -0.5034395 |
| Q9H0A8 | COMMD4 | | 41628000 | 0 | 0 | 0 | 37286000 | 0 | 13876000 | 12428666.67 | 0.941801744 | 0.895695205 | -0.158920212 |
| Q9P000 | COMMD9 | | 145020000 | 229090000 | 63331000 | 22604000 | 259670000 | 135060000 | 145813666.7 | 139111333.3 | 0.939901724 | 0.954034944 | -0.067885985 |
| P21964 | COMT | | 977710000 | 880010000 | 892150000 | 599230000 | 622580000 | 788120000 | 916623333.3 | 669976666.7 | 0.021111863 | 0.730918189 | -0.452218159 |
| Q86VU5 | COMTD1 | | 371860000 | 226400000 | 177700000 | 41819000 | 102630000 | 0 | 258653333.3 | 48149666.67 | 0.03246091 | 0.186155214 | -2.425422068 |
| P53621 | COPA | | 250140000 | 226280000 | 318910000 | 424480000 | 419220000 | 357850000 | 265110000 | 400516666.7 | 0.018096387 | 1.510756541 | 0.595271189 |
| P53618 | COPB1 | | 98067000 | 0 | 100030000 | 87007000 | 100560000 | 0 | 66032333.33 | 62522333.33 | 0.942390555 | 0.946844223 | -0.078801006 |
| O14579 | COPE | | 0 | 0 | 80519000 | 0 | 0 | 96772000 | 26839666.67 | 32257333.33 | 0.903505701 | 1.201852979 | 0.265260424 |
| Q9Y678 | COPG1 | | 259380000 | 232630000 | 185000000 | 169210000 | 237760000 | 225320000 | 225670000 | 210763333.3 | 0.648451975 | 0.933944846 | -0.098590741 |
| P61923 | COPZ1 | | 178200000 | 0 | 0 | 88914000 | 140290000 | 170770000 | 59400000 | 133324666.7 | 0.312511002 | 2.244523008 | 1.166408885 |
| Q9BR76 | CORO1B | | 352340000 | 372300000 | 468970000 | 183820000 | 159070000 | 327820000 | 397870000 | 223570000 | 0.052239037 | 0.561917209 | -0.83157051 |
| Q9ULV4 | CORO1C | | 1280200000 | 1574800000 | 1470300000 | 815410000 | 710360000 | 812200000 | 1441766667 | 779323333.3 | 0.002043079 | 0.540533605 | -0.887543783 |
| Q9UI42 | CPA4 | | 54888000 | 44285000 | 0 | 300390000 | 324350000 | 325380000 | 33057666.67 | 316706666.7 | 0.000109839 | 9.580430157 | 3.260090434 |
| O75976 | CPD | | 284890000 | 218860000 | 246680000 | 197850000 | 188670000 | 313910000 | 250143333.3 | 233476666.7 | 0.727698222 | 0.933371534 | -0.099476627 |
| O75131 | CPNE3 | | 34823000 | 0 | 0 | 0 | 48826000 | 0 | 11607666.67 | 16275333.33 | 0.826841237 | 1.402119289 | 0.487609096 |
| Q16630 | CPSF6 | | 0 | 0 | 102040000 | 93419000 | 161600000 | 0 | 34013333.33 | 85006333.33 | 0.428111746 | 2.499206194 | 1.321469934 |
| Q6UXH1 | CRELD2 | | 63730000 | 111350000 | 86239000 | 120270000 | 92771000 | 0 | 87106333.33 | 71013666.67 | 0.700289873 | 0.815252622 | -0.294680918 |
| O75718 | CRTAP | | 323110000 | 337390000 | 253160000 | 330600000 | 378620000 | 332120000 | 304553333.3 | 347113333.3 | 0.234417391 | 1.139745638 | 0.188711888 |
| Q08257 | CRYZ | | 643630000 | 471010000 | 489170000 | 487930000 | 460930000 | 630700000 | 534603333.3 | 526520000 | 0.92039921 | 0.984879755 | -0.0219805 |
| O75534 | CSDE1 | | 118400000 | 161640000 | 188980000 | 414910000 | 92536000 | 224580000 | 156340000 | 244008666.7 | 0.411887277 | 1.560756471 | 0.642245447 |
| P55060 | CSE1L | | 377750000 | 385560000 | 538510000 | 719050000 | 782990000 | 1195800000 | 433940000 | 899280000 | 0.042409725 | 2.072360234 | 1.051274806 |
| P67870 | CSNK2B | | 0 | 0 | 0 | 80187000 | 0 | 46227000 | 0 | 42138000 | 0.143991357 | N/A | N/A |
| Q6UVK1 | CSPG4 | | 2205600000 | 2339800000 | 2611400000 | 2386600000 | 2525700000 | 2146800000 | 2385600000 | 2353033333 | 0.851176184 | 0.986348647 | -0.019830405 |
| P21291 | CSRP1 | | 250510000 | 226290000 | 368820000 | 342150000 | 310730000 | 270630000 | 281873333.3 | 307836666.7 | 0.621867283 | 1.092109931 | 0.127118084 |
| Q16527 | CSRP2 | | 0 | 108970000 | 124790000 | 0 | 0 | 62731000 | 77920000 | 20910333.33 | 0.268944689 | 0.268356434 | -1.897777617 |
| Q8NHU0 | CT45A3 | | 52593000 | 0 | 55481000 | 0 | 0 | 77834000 | 36024666.67 | 25944666.67 | 0.765665969 | 0.72019172 | -0.473547081 |
| P35221 | CTNNA1 | | 3765900000 | 4255800000 | 4028100000 | 2260400000 | 2562700000 | 2573000000 | 4016600000 | 2465366667 | 0.00089009 | 0.61379442 | -0.704172565 |
| P35222 | CTNNB1 | | 2526900000 | 2555800000 | 2098000000 | 1647700000 | 1490400000 | 1549500000 | 2393566667 | 1562533333 | 0.005835594 | 0.652805437 | -0.615275022 |
| O60716 | CTNND1 | | 1880900000 | 1697800000 | 2106800000 | 1492100000 | 1468100000 | 1300500000 | 1895166667 | 1420233333 | 0.023217222 | 0.74939759 | -0.416196756 |
| P17812 | CTPS1 | | 0 | 0 | 0 | 57427000 | 45836000 | 96634000 | 0 | 66632333.33 | 0.012297665 | N/A | N/A |
| P10619 | CTSA | | 209020000 | 143960000 | 156620000 | 324490000 | 308070000 | 457230000 | 169866666.7 | 363263333.3 | 0.019542947 | 2.138520408 | 1.096612975 |
| P07858 | CTSB | | 1316000000 | 1020100000 | 1171300000 | 2628200000 | 2634300000 | 3475700000 | 1169133333 | 2912733333 | 0.004059842 | 2.491361122 | 1.316934155 |
| P53634 | CTSC | | 581840000 | 534550000 | 522080000 | 342220000 | 413740000 | 385220000 | 546156666.7 | 380393333.3 | 0.003883501 | 0.696491239 | -0.52182289 |
| P07339 | CTSD | | 843690000 | 1152100000 | 924550000 | 1213000000 | 1153100000 | 992520000 | 973446666.7 | 1119540000 | 0.267065463 | 1.150078416 | 0.201732231 |
| P07711 | CTSL | | 0 | 0 | 167670000 | 251580000 | 180430000 | 322710000 | 55890000 | 251573333.3 | 0.04776428 | 4.50122264 | 2.170316925 |
| Q9UBR2 | CTSZ | | 353450000 | 284510000 | 350880000 | 613780000 | 517330000 | 258270000 | 329613333.3 | 463126666.7 | 0.285975003 | 1.405060475 | 0.490632227 |
| Q14247 | CTTN | | 70908000 | 74320000 | 85563000 | 99367000 | 62288000 | 57974000 | 76930333.33 | 73209666.67 | 0.801683752 | 0.951635896 | -0.071518404 |
| Q13948 | CUX1 | | 98489000 | 220980000 | 215960000 | 63601000 | 92418000 | 301780000 | 178476333.3 | 152599666.7 | 0.776121734 | 0.855013456 | -0.225980969 |
| P78310 | CXADR | | 0 | 0 | 45240000 | 114480000 | 0 | 0 | 15080000 | 38160000 | 0.603809024 | 2.530503979 | 1.339424743 |
| Q7L576 | CYFIP1 | | 537910000 | 563480000 | 746560000 | 343860000 | 294170000 | 405040000 | 615983333.3 | 347690000 | 0.021396346 | 0.56444709 | -0.825089742 |
| Q96F07 | CYFIP2 | | 0 | 70760000 | 24400000 | 36013000 | 37160000 | 0 | 31720000 | 24391000 | 0.775963041 | 0.768947037 | -0.379043863 |
| Q9NUQ9 | CYRIB | | 381780000 | 416320000 | 475960000 | 445980000 | 563810000 | 392490000 | 424686666.7 | 467426666.7 | 0.499269895 | 1.100638902 | 0.138341227 |
| Q14118 | DAG1 | | 620740000 | 593670000 | 563850000 | 303640000 | 257830000 | 0 | 592753333.3 | 187156666.7 | 0.013388832 | 0.31574123 | -1.663185432 |
| Q8NCG7 | DAGLB | | 0 | 0 | 0 | 43084000 | 55781000 | 58226000 | 0 | 52363666.67 | 0.000367286 | N/A | N/A |
| P14868 | DARS1 | | 265660000 | 222010000 | 237810000 | 232730000 | 276870000 | 333340000 | 241826666.7 | 280980000 | 0.285537602 | 1.1619066 | 0.216494102 |
| Q16643 | DBN1 | | 2033800000 | 1694500000 | 2050000000 | 748510000 | 890230000 | 748010000 | 1926100000 | 795583333.3 | 0.00083281 | 0.413054012 | -1.275597649 |
| Q8N8Z6 | DCBLD1 | | 0 | 69008000 | 73952000 | 148920000 | 106150000 | 0 | 47653333.33 | 85023333.33 | 0.498727186 | 1.784205372 | 0.835281687 |
| Q96PD2 | DCBLD2 | | 74204000 | 0 | 111400000 | 0 | 0 | 212910000 | 61868000 | 70970000 | 0.912905551 | 1.147119674 | 0.198015909 |
| Q14203 | DCTN1 | | 0 | 0 | 90054000 | 139220000 | 114830000 | 71257000 | 30018000 | 108435666.7 | 0.094947358 | 3.612354809 | 1.852939603 |
| Q13561 | DCTN2 | | 366240000 | 250210000 | 368770000 | 217510000 | 297460000 | 448200000 | 328406666.7 | 321056666.7 | 0.929568206 | 0.977619212 | -0.032655458 |
| Q7Z4W1 | DCXR | | 648380000 | 514530000 | 308820000 | 155190000 | 201570000 | 297280000 | 490576666.7 | 218013333.3 | 0.063882152 | 0.444402166 | -1.170062246 |

| | | | | | | | | | | | | | |
|---|---|---|---|---|---|---|---|---|---|---|---|---|---|
| P39656 | DDOST | | 2285200000 | 1530600000 | 1639300000 | 1080900000 | 1204100000 | 1665800000 | 1818366667 | 1316933333 | 0.164652051 | 0.724239702 | -0.465460828 |
| Q96HY6 | DDRGK1 | | 437220000 | 357710000 | 321930000 | 236480000 | 352000000 | 351200000 | 372286666.7 | 313226666.7 | 0.313888645 | 0.84135881 | -0.249206905 |
| Q92499 | DDX1 | | 1244200000 | 872800000 | 943800000 | 555700000 | 704630000 | 839570000 | 1020266667 | 699966666.7 | 0.084472727 | 0.686062467 | -0.543588152 |
| Q92841 | DDX17 | | 170610000 | 161960000 | 199600000 | 147070000 | 180700000 | 166600000 | 177390000 | 164790000 | 0.447832188 | 0.928970066 | -0.106295985 |
| Q9NVP1 | DDX18 | | 0 | 0 | 0 | 50001000 | 27783000 | 0 | 0 | 25928000 | 0.147501418 | N/A | N/A |
| Q9NUU7 | DDX19A | | 0 | 0 | 89652000 | 0 | 88213000 | 90093000 | 29884000 | 59435333.33 | 0.521867922 | 1.988868068 | 0.991947578 |
| Q9NR30 | DDX21 | | 135580000 | 272820000 | 251390000 | 111290000 | 139350000 | 196380000 | 219930000 | 149006666.7 | 0.224685102 | 0.677518604 | -0.561667532 |
| Q13838 | DDX39B | | 285050000 | 205840000 | 226420000 | 239070000 | 317330000 | 322100000 | 239103333.3 | 292833333.3 | 0.208648123 | 1.224714559 | 0.292445543 |
| O00571 | DDX3X | | 903460000 | 710640000 | 672490000 | 625160000 | 548490000 | 846810000 | 762196666.7 | 673486666.7 | 0.481730661 | 0.883612716 | -0.178513915 |
| Q9H0S4 | DDX47 | | 97992000 | 108200000 | 121740000 | 55036000 | 42424000 | 0 | 109310666.7 | 32486666.67 | 0.012998401 | 0.29719576 | -1.750514561 |
| Q9Y6V7 | DDX49 | | 232770000 | 168870000 | 82746000 | 0 | 20270000 | 0 | 161462000 | 6756666.667 | 0.024517156 | 0.041846792 | -4.578739174 |
| P17844 | DDX5 | | 1517800000 | 1268300000 | 1332000000 | 834610000 | 1066200000 | 1231100000 | 1372700000 | 1043970000 | 0.07468922 | 0.760523057 | -0.394936108 |
| Q9BUN8 | DERL1 | | 207940000 | 143230000 | 301570000 | 110700000 | 139630000 | 392640000 | 217580000 | 214323333.3 | 0.975739297 | 0.985032325 | -0.021757025 |
| Q9GZP9 | DERL2 | | 0 | 0 | 126690000 | 55814000 | 112910000 | 0 | 42230000 | 56241333.33 | 0.80579355 | 1.33178625 | 0.41336255 |
| Q9BTZ2 | DHRS4 | | 111720000 | 125700000 | 75679000 | 404770000 | 317300000 | 146530000 | 104366333.3 | 289533333.3 | 0.074673776 | 2.774202409 | 1.472073052 |
| Q9Y394 | DHRS7 | | 373350000 | 301740000 | 443360000 | 477620000 | 649250000 | 1038100000 | 372816666.7 | 721656666.7 | 0.110588298 | 1.935687782 | 0.952846271 |
| Q7Z478 | DHX29 | | 0 | 0 | 0 | 10960000 | 8630000 | 0 | 0 | 6530000 | 0.121712833 | N/A | N/A |
| Q7L2E3 | DHX30 | | 883400000 | 1015800000 | 969240000 | 849550000 | 747490000 | 1024300000 | 956146666.7 | 873780000 | 0.410174738 | 0.913855615 | -0.129961851 |
| Q08211 | DHX9 | | 501360000 | 424650000 | 471080000 | 406750000 | 287430000 | 252450000 | 465696666.7 | 315543333.3 | 0.044079324 | 0.677572669 | -0.561552413 |
| O60610 | DIAPH1 | | 0 | 0 | 53242000 | 0 | 37281000 | 142890000 | 17747333.33 | 60057000 | 0.41275445 | 3.384001352 | 1.758730145 |
| Q12959 | DLG1 | | 483820000 | 502410000 | 449430000 | 298900000 | 304730000 | 323910000 | 478553333.3 | 309180000 | 0.000604611 | 0.64607219 | -0.630232719 |
| Q9NW81 | DMAC2 | | 0 | 0 | 52595000 | 52835000 | 0 | 0 | 17531666.67 | 17611666.67 | 0.997585533 | 1.004563171 | 0.00656829 |
| P31689 | DNAJA1 | | 494330000 | 599490000 | 477390000 | 677400000 | 677510000 | 492330000 | 523736666.7 | 615746666.7 | 0.273621027 | 1.17567989 | 0.233495302 |
| O60884 | DNAJA2 | | 134300000 | 194690000 | 139130000 | 180990000 | 312050000 | 217130000 | 156040000 | 236723333.3 | 0.138044534 | 1.517068273 | 0.601286013 |
| P25685 | DNAJB1 | | 26327000 | 75412000 | 0 | 188530000 | 208880000 | 246460000 | 33913000 | 214623333.3 | 0.002912772 | 6.328644866 | 2.661896613 |
| Q9UBS4 | DNAJB11 | | 365830000 | 456850000 | 439520000 | 533710000 | 671920000 | 434150000 | 420733333.3 | 546593333.3 | 0.165848418 | 1.299144351 | 0.377561741 |
| Q9NXW2 | DNAJB12 | | 164890000 | 175900000 | 160200000 | 0 | 0 | 0 | 166996666.7 | 0 | 3.59754E-06 | 0 | N/A |
| P25686 | DNAJB2 | | 115360000 | 70766000 | 84433000 | 77227000 | 70716000 | 84219000 | 90186333.33 | 77387333.33 | 0.404779623 | 0.85808271 | -0.22081138 |
| Q96KC8 | DNAJC1 | | 0 | 0 | 0 | 0 | 54134000 | 45029000 | 0 | 33054333.33 | 0.119459081 | N/A | N/A |
| Q8IXB1 | DNAJC10 | | 161560000 | 311060000 | 316810000 | 554920000 | 588010000 | 854250000 | 263143333.3 | 665726666.7 | 0.020042434 | 2.529901321 | 1.339081114 |
| Q9NVH1 | DNAJC11 | | 689190000 | 558270000 | 552990000 | 320630000 | 458880000 | 334820000 | 600150000 | 371443333.3 | 0.021648563 | 0.618917493 | -0.692180997 |
| O75165 | DNAJC13 | | 73946000 | 0 | 177780000 | 0 | 0 | 0 | 83908666.67 | 0 | 0.178994196 | 0 | N/A |
| Q13217 | DNAJC3 | | 165370000 | 109950000 | 155660000 | 155450000 | 188360000 | 126020000 | 143660000 | 156610000 | 0.629426952 | 1.090143394 | 0.124517915 |
| O75937 | DNAJC8 | | 76497000 | 0 | 0 | 86232000 | 86441000 | 0 | 25499000 | 57557666.67 | 0.45130565 | 2.257251918 | 1.174567438 |
| Q8WXX5 | DNAJC9 | | 738370000 | 627430000 | 547190000 | 432850000 | 550720000 | 558240000 | 637663333.3 | 513936666.7 | 0.14609867 | 0.805968667 | -0.311204342 |
| O60762 | DPM1 | | 0 | 0 | 63905000 | 57785000 | 41511000 | 96258000 | 21301666.67 | 65184666.67 | 0.176641256 | 3.060073547 | 1.613566327 |
| Q9C005 | DPY30 | | 106020000 | 42420000 | 64570000 | 0 | 0 | 0 | 71003333.33 | 0 | 0.018951274 | 0 | N/A |
| Q16555 | DPYSL2 | | 0 | 0 | 0 | 338200000 | 346460000 | 349100000 | 0 | 344586666.7 | 4.94066E-08 | N/A | N/A |
| Q9Y295 | DRG1 | | 0 | 0 | 75398000 | 0 | 0 | 128450000 | 25132666.67 | 42816666.67 | 0.739692446 | 1.703626091 | 0.76860873 |
| Q14126 | DSG2 | | 460930000 | 462840000 | 546170000 | 298350000 | 339710000 | 407310000 | 489980000 | 348456666.7 | 0.028903391 | 0.711165082 | -0.491743606 |
| Q03001 | DST | | 265580000 | 151200000 | 159130000 | 274250000 | 286890000 | 204440000 | 191970000 | 255193333.3 | 0.231979673 | 1.329339654 | 0.410709768 |
| P60981 | DSTN | | 0 | 0 | 0 | 423320000 | 281380000 | 152420000 | 0 | 285706666.7 | 0.021732251 | N/A | N/A |
| P23919 | DTYMK | | 67216000 | 89419000 | 0 | 109510000 | 79432000 | 0 | 52211666.67 | 62980666.67 | 0.811612699 | 1.206256584 | 0.270536817 |
| Q14204 | DYNC1H1 | | 2033900000 | 1764700000 | 1428300000 | 2902800000 | 2890600000 | 4328500000 | 1742300000 | 3373966667 | 0.032609642 | 1.936501559 | 0.953452663 |
| Q13409 | DYNC1I2 | | 73431000 | 94017000 | 78670000 | 0 | 0 | 0 | 82039333.33 | 0 | 0.000185734 | 0 | N/A |
| Q9Y6G9 | DYNC1LI1 | | 0 | 96491000 | 0 | 142250000 | 0 | 0 | 32163666.67 | 47416666.67 | 0.803233842 | 1.474230757 | 0.559962363 |
| O43237 | DYNC1LI2 | | 0 | 0 | 0 | 40756000 | 78174000 | 0 | 0 | 39643333.33 | 0.153903374 | N/A | N/A |
| P63167 | DYNLL1 | | 105990000 | 151360000 | 252470000 | 275080000 | 304280000 | 0 | 169940000 | 193120000 | 0.837839149 | 1.136401083 | 0.184472111 |
| Q15125 | EBP | | 321580000 | 412800000 | 345490000 | 0 | 0 | 164760000 | 359956666.7 | 54920000 | 0.007633518 | 0.152573921 | -2.712419708 |
| P42892 | ECE1 | | 0 | 0 | 90362000 | 111310000 | 106420000 | 71962000 | 30120666.67 | 96564000 | 0.110912078 | 3.205905137 | 1.680731737 |
| Q9NTX5 | ECHDC1 | | 224640000 | 331950000 | 94511000 | 0 | 132270000 | 0 | 217033666.7 | 44090000 | 0.101372223 | 0.203148206 | -2.299395472 |
| O75521 | ECI2 | | 985210000 | 1025000000 | 976560000 | 548570000 | 654010000 | 776680000 | 995590000 | 659753333.3 | 0.007653917 | 0.662675733 | -0.593625005 |
| Q6P2E9 | EDC4 | | 113140000 | 124480000 | 76502000 | 56670000 | 111210000 | 196840000 | 104707333.3 | 121573333.3 | 0.716663697 | 1.161077543 | 0.215464327 |
| O60869 | EDF1 | | 184810000 | 0 | 215100000 | 225620000 | 238250000 | 284070000 | 133303333.3 | 249313333.3 | 0.170535967 | 1.870270811 | 0.903247184 |
| O43854 | EDIL3 | | 391010000 | 362100000 | 413430000 | 392600000 | 509890000 | 435180000 | 388846666.7 | 445890000 | 0.201512136 | 1.146698784 | 0.197486473 |
| Q15075 | EEA1 | | 276960000 | 303290000 | 193400000 | 553890000 | 633830000 | 228200000 | 257883333.3 | 471973333.3 | 0.170764384 | 1.830181607 | 0.871986812 |
| P68104 | EEF1A1 | | 3664500000 | 3643100000 | 3891800000 | 7753900000 | 7463500000 | 6728000000 | 3733133333 | 7315133333 | 0.000343177 | 1.959515688 | 0.970497124 |

| | | | | | | | | | | | | | |
|---|---|---|---|---|---|---|---|---|---|---|---|---|---|
| Q05639 | EEF1A2 | | 40388000 | 0 | 72716000 | 132790000 | 137560000 | 239840000 | 37701333.33 | 170063333.3 | 0.031458894 | 4.510804216 | 2.173384669 |
| P24534 | EEF1B2 | | 720730000 | 493260000 | 417690000 | 469200000 | 153800000 | 627160000 | 543893333.3 | 416720000 | 0.487053751 | 0.766179643 | -0.3842454 |
| P29692 | EEF1D | | 1360600000 | 1251400000 | 1485200000 | 1754900000 | 2144300000 | 1637100000 | 1365733333 | 1845433333 | 0.045731779 | 1.351239871 | 0.434283804 |
| O43324 | EEF1E1 | | 374950000 | 0 | 310760000 | 150510000 | 218690000 | 357740000 | 228570000 | 242313333.3 | 0.921409184 | 1.060127459 | 0.08423773 |
| P26641 | EEF1G | | 2492200000 | 2575900000 | 2479200000 | 3635500000 | 3731600000 | 3469900000 | 2515766667 | 3612333333 | 0.0001827 | 1.435877731 | 0.521932905 |
| P13639 | EEF2 | | 3921100000 | 2866000000 | 3173700000 | 3526900000 | 3608100000 | 3729800000 | 3320266667 | 3621600000 | 0.398036019 | 1.090755763 | 0.125328096 |
| Q96C19 | EFHD2 | | 87050000 | 80074000 | 75830000 | 135550000 | 90512000 | 83233000 | 80984666.67 | 103098333.3 | 0.255684629 | 1.273059921 | 0.348300326 |
| Q15029 | EFTUD2 | | 0 | 52807000 | 0 | 30418000 | 52826000 | 33447000 | 17602333.33 | 38897000 | 0.324011502 | 2.209763857 | 1.143892207 |
| Q8N3D4 | EHBP1L1 | | 224410000 | 78651000 | 169980000 | 89272000 | 104900000 | 92626000 | 157680333.3 | 95599333.33 | 0.220433466 | 0.606285713 | -0.721930269 |
| Q9H4M9 | EHD1 | | 270540000 | 264880000 | 431590000 | 494660000 | 576770000 | 445300000 | 322336666.7 | 505576666.7 | 0.0516521 | 1.568473956 | 0.649361574 |
| Q9NZN4 | EHD2 | | 35818000 | 139110000 | 0 | 0 | 0 | 0 | 58309333.33 | 0 | 0.234593252 | 0 | N/A |
| Q9H223 | EHD4 | | 226220000 | 208830000 | 200470000 | 0 | 0 | 282280000 | 211840000 | 94093333.33 | 0.280316994 | 0.444171702 | -1.170810613 |
| P41567 | EIF1 | | 0 | 150240000 | 51906000 | 90909000 | 58750000 | 58543000 | 67382000 | 69400666.67 | 0.966628297 | 1.029958545 | 0.042586271 |
| Q9BY44 | EIF2A | | 0 | 87976000 | 73043000 | 77555000 | 0 | 63314000 | 53673000 | 46956333.33 | 0.861647602 | 0.874859489 | -0.192876771 |
| P19525 | EIF2AK2 | | 156020000 | 96374000 | 69343000 | 84561000 | 109250000 | 221890000 | 107245666.7 | 138567000 | 0.560638418 | 1.292052204 | 0.369664362 |
| P05198 | EIF2S1 | | 211050000 | 139310000 | 155490000 | 186800000 | 188250000 | 135800000 | 168616666.7 | 170283333.3 | 0.954969816 | 1.009884353 | 0.014190092 |
| P20042 | EIF2S2 | | 0 | 0 | 187420000 | 142200000 | 249500000 | 0 | 62473333.33 | 130566666.7 | 0.515310322 | 2.089958382 | 1.063474214 |
| P41091 | EIF2S3 | | 581530000 | 581490000 | 565110000 | 585750000 | 540710000 | 485300000 | 576043333.3 | 537253333.3 | 0.259655931 | 0.932661316 | -0.100574815 |
| Q14152 | EIF3A | | 263670000 | 281360000 | 190670000 | 171270000 | 175730000 | 197820000 | 245233333.3 | 181606666.7 | 0.092830134 | 0.740546418 | -0.433337927 |
| Q99613 | EIF3C | | 288500000 | 350220000 | 231440000 | 169160000 | 227650000 | 312280000 | 290053333.3 | 236363333.3 | 0.375358273 | 0.814896111 | -0.295311949 |
| P60228 | EIF3E | | 52775000 | 46960000 | 0 | 0 | 0 | 0 | 33245000 | 0 | 0.117466902 | 0 | N/A |
| O00303 | EIF3F | | 200420000 | 159870000 | 166170000 | 0 | 0 | 168500000 | 175486666.7 | 56166666.67 | 0.106880436 | 0.320062303 | -1.643575329 |
| O75821 | EIF3G | | 0 | 94012000 | 80742000 | 94235000 | 0 | 73071000 | 58251333.33 | 55768666.67 | 0.954577391 | 0.957380089 | -0.062836293 |
| Q13347 | EIF3I | | 0 | 146420000 | 0 | 165520000 | 165140000 | 102330000 | 48806666.67 | 144330000 | 0.146609896 | 2.957177981 | 1.564221076 |
| O75822 | EIF3J | | 533440000 | 456940000 | 724380000 | 317280000 | 365260000 | 816220000 | 571586666.7 | 499586666.7 | 0.706093791 | 0.87403485 | -0.194237289 |
| Q9Y262 | EIF3L | | 59067000 | 56019000 | 91854000 | 0 | 0 | 0 | 68980000 | 0 | 0.003850605 | 0 | N/A |
| Q7L2H7 | EIF3M | | 135850000 | 134210000 | 43114000 | 0 | 0 | 184710000 | 104391333.3 | 61570000 | 0.567258484 | 0.58979992 | -0.761702469 |
| P60842 | EIF4A1 | | 749660000 | 507880000 | 656570000 | 1062700000 | 921050000 | 1074500000 | 638036666.7 | 1019416667 | 0.01135967 | 1.597739942 | 0.676032605 |
| Q14240 | EIF4A2 | | 85179000 | 0 | 0 | 0 | 203080000 | 0 | 28393000 | 67693333.33 | 0.62076864 | 2.384155719 | 1.253478467 |
| P38919 | EIF4A3 | | 0 | 0 | 0 | 0 | 38704000 | 38990000 | 0 | 25898000 | 0.116121913 | N/A | N/A |
| P23588 | EIF4B | | 149810000 | 41626000 | 114870000 | 95418000 | 77785000 | 209360000 | 102102000 | 127521000 | 0.65127888 | 1.248956925 | 0.320723722 |
| P06730 | EIF4E | | 168060000 | 150690000 | 147430000 | 271980000 | 246400000 | 139380000 | 155393333.3 | 219253333.3 | 0.195336414 | 1.410957141 | 0.496674166 |
| Q04637 | EIF4G1 | | 194720000 | 219210000 | 267310000 | 348800000 | 308150000 | 403410000 | 227080000 | 353453333.3 | 0.022284195 | 1.556514591 | 0.638319101 |
| Q15056 | EIF4H | | 333110000 | 252630000 | 282190000 | 250440000 | 156350000 | 162220000 | 289310000 | 189670000 | 0.060601345 | 0.655594345 | -0.609124684 |
| P55010 | EIF5 | | 82158000 | 0 | 27667000 | 68222000 | 88473000 | 89366000 | 36608333.33 | 82020333.33 | 0.144707548 | 2.240482586 | 1.163809513 |
| P63241 | EIF5A | | 998740000 | 609140000 | 620750000 | 783700000 | 769320000 | 1076500000 | 742876666.7 | 876506666.7 | 0.456983768 | 1.179881811 | 0.238642352 |
| O60841 | EIF5B | | 0 | 0 | 94701000 | 163670000 | 100530000 | 132880000 | 31567000 | 132360000 | 0.050585848 | 4.192986347 | 2.067978132 |
| Q9BQ52 | ELAC2 | | 207720000 | 133440000 | 129750000 | 236530000 | 89835000 | 118450000 | 156970000 | 148271666.7 | 0.874268946 | 0.944586014 | -0.08224592 |
| Q15717 | ELAVL1 | | 0 | 0 | 0 | 0 | 81737000 | 52171000 | 0 | 44636000 | 0.135134262 | N/A | N/A |
| P0C7U0 | ELFN1 | | 87265000 | 0 | 0 | 0 | 65043000 | 0 | 29088333.33 | 21681000 | 0.848184664 | 0.74535037 | -0.424009337 |
| Q8IZ81 | ELMOD2 | | 0 | 0 | 0 | 31953000 | 29887000 | 0 | 0 | 20613333.33 | 0.116560282 | N/A | N/A |
| Q8N766 | EMC1 | | 38541000 | 66166000 | 0 | 83096000 | 222480000 | 185720000 | 34902333.33 | 163765333.3 | 0.048468387 | 4.692102725 | 2.230234599 |
| Q5UCC4 | EMC10 | | 104690000 | 130210000 | 156050000 | 119940000 | 116340000 | 0 | 130316666.7 | 78760000 | 0.287820656 | 0.604373961 | -0.726486591 |
| Q15006 | EMC2 | | 67227000 | 111570000 | 151010000 | 117020000 | 136280000 | 0 | 109935666.7 | 84433333.33 | 0.63007328 | 0.768024936 | -0.380774943 |
| Q5J8M3 | EMC4 | | 173320000 | 171690000 | 0 | 0 | 0 | 185020000 | 115003333.3 | 61673333.33 | 0.561439119 | 0.536274311 | -0.898956949 |
| Q9NPA0 | EMC7 | | 358890000 | 0 | 277750000 | 296140000 | 306920000 | 269560000 | 212213333.3 | 290873333.3 | 0.511267371 | 1.37066474 | 0.454875736 |
| O43402 | EMC8 | | 60862000 | 76083000 | 44909000 | 84360000 | 92622000 | 0 | 60618000 | 58994000 | 0.960645334 | 0.973209278 | -0.039178021 |
| P50402 | EMD | | 189970000 | 197470000 | 297720000 | 246210000 | 240480000 | 244420000 | 228386666.7 | 243703333.3 | 0.682370277 | 1.067064627 | 0.093647556 |
| Q8N8S7 | ENAH | | 85003000 | 136930000 | 172680000 | 227000000 | 165670000 | 106330000 | 131537666.7 | 166333333.3 | 0.465174259 | 1.264530059 | 0.338601331 |
| P06733 | ENO1 | | 1786500000 | 1891500000 | 1757200000 | 3450800000 | 3754500000 | 3507300000 | 1811733333 | 3570866667 | 6.57542E-05 | 1.97096703 | 0.978903643 |
| P11171 | EPB41 | | 85876000 | 0 | 69576000 | 0 | 78131000 | 52239000 | 51817333.33 | 43456666.67 | 0.822685646 | 0.838651159 | -0.253857255 |
| O43491 | EPB41L2 | | 1548800000 | 1489100000 | 1669000000 | 725840000 | 573490000 | 460810000 | 1568966667 | 586713333.3 | 0.000459432 | 0.373948884 | -1.419087019 |
| Q9Y2J2 | EPB41L3 | | 2218600000 | 2032100000 | 2109800000 | 2050700000 | 2042900000 | 1773000000 | 2120166667 | 1955533333 | 0.195731306 | 0.922348872 | -0.116615552 |
| Q9HCM4 | EPB41L5 | | 117200000 | 53170000 | 0 | 0 | 0 | 0 | 56790000 | 0 | 0.169015652 | 0 | N/A |
| P16422 | EPCAM | | 0 | 0 | 53345000 | 0 | 0 | 54518000 | 17781666.67 | 18172666.67 | 0.98846667 | 1.02198894 | 0.031379583 |
| P29317 | EPHA2 | | 192880000 | 252240000 | 379850000 | 117590000 | 67863000 | 186460000 | 274990000 | 123971000 | 0.080833095 | 0.45082003 | -1.149376479 |
| P29323 | EPHB2 | | 208080000 | 391760000 | 338220000 | 241640000 | 291940000 | 356010000 | 312686666.7 | 296530000 | 0.812552283 | 0.948329531 | -0.076539632 |

| | | | | | | | | | | | | | |
|---|---|---|---|---|---|---|---|---|---|---|---|---|---|
| P54760 | EPHB4 | | 111690000 | 107230000 | 94145000 | 0 | 0 | 0 | 104355000 | 0 | 3.82216E-05 | 0 | N/A |
| Q9Y6I3 | EPN1 | | 78452000 | 75484000 | 122390000 | 56399000 | 64309000 | 84899000 | 92108666.67 | 68535666.67 | 0.246542204 | 0.744074029 | -0.426481932 |
| P07814 | EPRS1 | | 1290400000 | 1144500000 | 1015100000 | 1064800000 | 1262100000 | 1408500000 | 1150000000 | 1245133333 | 0.49683604 | 1.082724638 | 0.114666378 |
| Q9UBC2 | EPS15L1 | | 111030000 | 56987000 | 161580000 | 109960000 | 226970000 | 159440000 | 109865666.7 | 165456666.7 | 0.288038615 | 1.505990649 | 0.590712812 |
| Q12929 | EPS8 | | 65563000 | 92220000 | 90168000 | 0 | 0 | 0 | 82650333.33 | 0 | 0.000644839 | 0 | N/A |
| Q9H6S3 | EPS8L2 | | 0 | 0 | 0 | 70438000 | 0 | 101480000 | 0 | 57306000 | 0.128920027 | N/A | N/A |
| Q96RT1 | ERBIN | | 497250000 | 216560000 | 322620000 | 0 | 11774000 | 0 | 345476666.7 | 3924666.667 | 0.01404118 | 0.01136015 | -6.459874338 |
| Q8IUD2 | ERC1 | | 159570000 | 139930000 | 195350000 | 104070000 | 108100000 | 0 | 164950000 | 70723333.33 | 0.072695422 | 0.428756189 | -1.221770599 |
| Q96RQ1 | ERGIC2 | | 113620000 | 131190000 | 117620000 | 155860000 | 114790000 | 116270000 | 120810000 | 128973333.3 | 0.602607098 | 1.067571669 | 0.094332925 |
| P84090 | ERH | | 178310000 | 278410000 | 278810000 | 407890000 | 373490000 | 217370000 | 245176666.7 | 332916666.7 | 0.263405514 | 1.357864397 | 0.441339412 |
| Q96DZ1 | ERLEC1 | | 202990000 | 241060000 | 199280000 | 245580000 | 261910000 | 265000000 | 214443333.3 | 257496666.7 | 0.04242088 | 1.20076788 | 0.263957291 |
| O75477 | ERLIN1 | | 942920000 | 913520000 | 869700000 | 507330000 | 422490000 | 576850000 | 908713333.3 | 502223333.3 | 0.001193209 | 0.552675211 | -0.855496189 |
| O94905 | ERLIN2 | | 47985000 | 0 | 0 | 0 | 35481000 | 12076000 | 15995000 | 15852333.33 | 0.994394156 | 0.991080546 | -0.012925784 |
| Q96HE7 | ERO1A | | 528770000 | 669760000 | 792220000 | 652760000 | 785220000 | 997850000 | 663583333.3 | 811943333.3 | 0.304548403 | 1.22357403 | 0.291101391 |
| P30040 | ERP29 | | 4697900000 | 4721100000 | 6592300000 | 4641300000 | 5216300000 | 5707100000 | 5337100000 | 5188233333 | 0.84179251 | 0.972107199 | -0.040812679 |
| Q9BS26 | ERP44 | | 432710000 | 432960000 | 172650000 | 247370000 | 352500000 | 767120000 | 346106666.7 | 455663333.3 | 0.577278664 | 1.31654018 | 0.396751553 |
| Q14674 | ESPL1 | | 0 | 17830000 | 0 | 5065300 | 0 | 0 | 5943333.333 | 1688433.333 | 0.528898796 | 0.284088615 | -1.815587081 |
| Q9BSJ8 | ESYT1 | | 4580100000 | 4205800000 | 4393500000 | 2857900000 | 2885700000 | 3071000000 | 4393133333 | 2938200000 | 0.0003321 | 0.668816486 | -0.580317685 |
| P62495 | ETF1 | | 617590000 | 537940000 | 470280000 | 376680000 | 452140000 | 531840000 | 541936666.7 | 453553333.3 | 0.22589377 | 0.836912062 | -0.256852054 |
| P38117 | ETFB | | 3551500000 | 3424400000 | 3427400000 | 2698600000 | 2893500000 | 2428100000 | 3467766667 | 2673400000 | 0.004920627 | 0.770928455 | -0.375331115 |
| Q01844 | EWSR1 | | 108830000 | 146020000 | 119220000 | 91101000 | 0 | 112600000 | 124690000 | 67900333.33 | 0.19224307 | 0.544553158 | -0.876855205 |
| Q9NVH0 | EXD2 | | 55175000 | 82999000 | 52949000 | 110690000 | 154290000 | 134030000 | 63707666.67 | 133003333.3 | 0.012023437 | 2.087713148 | 1.061923499 |
| Q9NV70 | EXOC1 | | 0 | 148680000 | 222790000 | 143170000 | 0 | 182780000 | 123823333.3 | 108650000 | 0.868320684 | 0.877459822 | -0.188595028 |
| Q96KP1 | EXOC2 | | 0 | 41836000 | 37752000 | 31135000 | 38944000 | 0 | 26529333.33 | 23359666.67 | 0.867733283 | 0.880522189 | -0.183568734 |
| O00471 | EXOC5 | | 97304000 | 378840000 | 101600000 | 57353000 | 129340000 | 0 | 192581333.3 | 62231000 | 0.263866338 | 0.323141391 | -1.629762537 |
| Q9Y2D4 | EXOC6B | | 116640000 | 173550000 | 128490000 | 81950000 | 103950000 | 122820000 | 139560000 | 102906666.7 | 0.155491674 | 0.737365052 | -0.439549054 |
| Q9UPT5 | EXOC7 | | 344750000 | 270100000 | 196640000 | 201460000 | 229350000 | 348540000 | 270496666.7 | 259783333.3 | 0.871505581 | 0.960393843 | -0.05830194 |
| Q8IYI6 | EXOC8 | | 80797000 | 86832000 | 0 | 42139000 | 0 | 112940000 | 55876333.33 | 51693000 | 0.927574301 | 0.925132286 | -0.112268421 |
| P15311 | EZR | | 2006800000 | 1810800000 | 1817600000 | 2554100000 | 3427000000 | 2723700000 | 1878400000 | 2901600000 | 0.020421519 | 1.54471891 | 0.627344337 |
| O95864 | FADS2 | | 469750000 | 487980000 | 416080000 | 229460000 | 0 | 214070000 | 457936666.7 | 147843333.3 | 0.015862753 | 0.322846682 | -1.631078895 |
| Q96CS3 | FAF2 | | 787810000 | 754500000 | 707720000 | 561180000 | 697160000 | 672170000 | 750010000 | 643503333.3 | 0.089844413 | 0.857993005 | -0.22096221 |
| Q96GK7 | FAHD2A | | 178800000 | 316560000 | 255400000 | 174060000 | 211640000 | 265880000 | 250253333.3 | 217193333.3 | 0.528372593 | 0.867893868 | -0.204409465 |
| Q9NZB2 | FAM120A | | 0 | 0 | 0 | 59297000 | 43356000 | 0 | 0 | 34217666.67 | 0.1256191 | N/A | N/A |
| Q96C01 | FAM136A | | 160980000 | 154090000 | 243400000 | 0 | 101700000 | 0 | 186156666.7 | 33900000 | 0.026576298 | 0.182104679 | -2.457160105 |
| Q96A26 | FAM162A | | 136910000 | 168960000 | 221530000 | 509780000 | 363700000 | 205680000 | 175800000 | 359720000 | 0.1139506 | 2.046188851 | 1.032939303 |
| O75063 | FAM20B | | 97198000 | 58705000 | 18988000 | 23944000 | 0 | 0 | 58297000 | 7981333.333 | 0.103525503 | 0.136908131 | -2.86871996 |
| Q92520 | FAM3C | | 172780000 | 32102000 | 61241000 | 71332000 | 59534000 | 108030000 | 88707666.67 | 79632000 | 0.850945331 | 0.897690166 | -0.155710504 |
| Q8NCA5 | FAM98A | | 143880000 | 123290000 | 81223000 | 79368000 | 87952000 | 86866000 | 116131000 | 84728666.67 | 0.167241459 | 0.7295956 | -0.454831065 |
| Q52LJ0 | FAM98B | | 39055000 | 0 | 37199000 | 0 | 0 | 0 | 25418000 | 0 | 0.116352105 | 0 | N/A |
| Q8WVX9 | FAR1 | | 0 | 308110000 | 239880000 | 0 | 0 | 0 | 182663333.3 | 0 | 0.122246158 | 0 | N/A |
| Q9Y285 | FARSA | | 208420000 | 318370000 | 0 | 242820000 | 178740000 | 0 | 175596666.7 | 140520000 | 0.781599642 | 0.800242981 | -0.321489977 |
| Q9NSD9 | FARSB | | 245050000 | 268050000 | 268510000 | 391870000 | 457410000 | 548080000 | 260536666.7 | 465786666.7 | 0.011098093 | 1.787796983 | 0.838182918 |
| P25445 | FAS | | 0 | 0 | 42953000 | 0 | 73550000 | 65174000 | 14317666.67 | 46241333.33 | 0.30721771 | 3.229669639 | 1.6913866 |
| P49327 | FASN | | 1493600000 | 1399800000 | 1405100000 | 955850000 | 962050000 | 950540000 | 1432833333 | 956146666.7 | 9.91839E-05 | 0.667311853 | -0.583566965 |
| P62861 | FAU | | 0 | 0 | 0 | 247810000 | 199850000 | 247540000 | 0 | 231733333.3 | 0.000130249 | N/A | N/A |
| P35556 | FBN2 | | 348780000 | 293920000 | 414520000 | 0 | 0 | 0 | 352406666.7 | 0 | 0.000538939 | 0 | N/A |
| P37268 | FDFT1 | | 0 | 0 | 235190000 | 138400000 | 146960000 | 0 | 78396666.67 | 95120000 | 0.864203447 | 1.213316893 | 0.278956401 |
| P39748 | FEN1 | | 1194100000 | 1040200000 | 1463400000 | 551230000 | 522500000 | 326490000 | 1232566667 | 466740000 | 0.005778056 | 0.378673229 | -1.400974663 |
| Q96AC1 | FERMT2 | | 170490000 | 257520000 | 129460000 | 183200000 | 125290000 | 142220000 | 185823333.3 | 150236666.7 | 0.439340464 | 0.808491937 | -0.306694709 |
| Q9Y613 | FHOD1 | | 0 | 0 | 0 | 56592000 | 66578000 | 0 | 0 | 41056666.67 | 0.11872448 | N/A | N/A |
| Q96AY3 | FKBP10 | | 242280000 | 165750000 | 719410000 | 989410000 | 930760000 | 933190000 | 375813333.3 | 951120000 | 0.029898313 | 2.530830909 | 1.339611121 |
| Q9NWM8 | FKBP14 | | 133260000 | 113180000 | 96950000 | 253170000 | 201740000 | 170180000 | 114463333.3 | 208363333.3 | 0.02355776 | 1.820350039 | 0.864215896 |
| P26885 | FKBP2 | | 1197700000 | 1331500000 | 1237500000 | 817170000 | 751070000 | 880100000 | 1255566667 | 816113333.3 | 0.001277262 | 0.649996018 | -0.621497216 |
| Q00688 | FKBP3 | | 520560000 | 672680000 | 244570000 | 1591700000 | 1509300000 | 699910000 | 479270000 | 1266970000 | 0.064412826 | 2.643541219 | 1.402471822 |
| Q02790 | FKBP4 | | 0 | 0 | 0 | 219320000 | 303880000 | 321160000 | 0 | 281453333.3 | 0.000863911 | N/A | N/A |
| Q14318 | FKBP8 | | 151770000 | 72109000 | 115410000 | 180050000 | 189180000 | 0 | 113096333.3 | 123076666.7 | 0.886712208 | 1.088246303 | 0.122005119 |
| O95302 | FKBP9 | | 179100000 | 54742000 | 54194000 | 407730000 | 428120000 | 341450000 | 96012000 | 392433333.3 | 0.003794587 | 4.087336305 | 2.031160953 |

| | | | | | | | | | | | | | |
|---|---|---|---|---|---|---|---|---|---|---|---|---|---|
| Q13045 | FLII | | 76950000 | 62930000 | 43784000 | 45577000 | 21378000 | 79927000 | 61221333.33 | 48960666.67 | 0.563958924 | 0.79973212 | -0.322411263 |
| P21333 | FLNA | | 23205000000 | 24304000000 | 25038000000 | 23510000000 | 22755000000 | 19664000000 | 24182333333 | 21976333333 | 0.162798673 | 0.908776379 | -0.138002758 |
| O75369 | FLNB | | 3374900000 | 3346200000 | 3915500000 | 2435400000 | 2506200000 | 2212600000 | 3545533333 | 2384733333 | 0.004813641 | 0.672602147 | -0.57217471 |
| Q14315 | FLNC | | 3112100000 | 3386300000 | 3385100000 | 6422600000 | 6195500000 | 6106900000 | 3294500000 | 6241666667 | 2.31028E-05 | 1.894571761 | 0.921871786 |
| O75955 | FLOT1 | | 790510000 | 768990000 | 730630000 | 1276700000 | 912840000 | 745510000 | 763376666.7 | 978350000 | 0.244702646 | 1.281608468 | 0.357955585 |
| Q14254 | FLOT2 | | 864120000 | 877930000 | 581090000 | 709560000 | 1022500000 | 1365200000 | 774380000 | 1032420000 | 0.291630061 | 1.333221416 | 0.414916397 |
| P02751 | FN1 | | 0 | 0 | 0 | 339940000 | 250550000 | 204600000 | 0 | 265030000 | 0.002625493 | N/A | N/A |
| Q5T0N5 | FNBP1L | | 26072000 | 15273000 | 57422000 | 54958000 | 45760000 | 181500000 | 32922333.33 | 94072666.67 | 0.25084286 | 2.857411889 | 1.514709012 |
| Q53EP0 | FNDC3B | | 181050000 | 129090000 | 123410000 | 147680000 | 193910000 | 186610000 | 144516666.7 | 176066666.7 | 0.246924878 | 1.21831392 | 0.284885917 |
| Q9BZ67 | FRMD8 | | 74701000 | 100880000 | 77200000 | 87429000 | 116910000 | 116510000 | 84260333.33 | 106949666.7 | 0.15194221 | 1.269276567 | 0.344006458 |
| Q16658 | FSCN1 | | 1539000000 | 1575600000 | 1640400000 | 969470000 | 842380000 | 1090700000 | 1585000000 | 967516666.7 | 0.001349726 | 0.61042061 | -0.712124422 |
| Q12841 | FSTL1 | | 390870000 | 299120000 | 356750000 | 59341000 | 174740000 | 132930000 | 348913333.3 | 122337000 | 0.006249493 | 0.350622886 | -1.512007926 |
| Q8IY81 | FTSJ3 | | 0 | 0 | 0 | 0 | 29076000 | 41497000 | 0 | 23524333.33 | 0.12828942 | N/A | N/A |
| Q96AE4 | FUBP1 | | 76857000 | 81959000 | 0 | 160350000 | 159110000 | 98150000 | 52938666.67 | 139203333.3 | 0.061798629 | 2.629520955 | 1.394799994 |
| Q9BWH2 | FUNDC2 | | 100800000 | 164430000 | 131620000 | 0 | 221970000 | 0 | 132283333.3 | 73990000 | 0.487106727 | 0.559329722 | -0.838229101 |
| P35637 | FUS | | 320320000 | 298230000 | 243150000 | 214120000 | 261170000 | 173570000 | 287233333.3 | 216286666.7 | 0.106422886 | 0.752999884 | -0.409278452 |
| P51114 | FXR1 | | 317830000 | 375820000 | 406560000 | 534090000 | 728540000 | 432860000 | 366736666.7 | 565163333.3 | 0.093611807 | 1.541060343 | 0.623923354 |
| Q13283 | G3BP1 | | 407360000 | 447710000 | 448830000 | 543160000 | 643080000 | 686990000 | 434633333.3 | 624410000 | 0.013193974 | 1.436636245 | 0.522694819 |
| P11413 | G6PD | | 708660000 | 722100000 | 745630000 | 1128900000 | 901850000 | 822870000 | 725463333.3 | 951206666.7 | 0.07088821 | 1.311171251 | 0.390856128 |
| P10253 | GAA | | 0 | 108070000 | 148940000 | 242610000 | 207080000 | 0 | 85670000 | 149896666.7 | 0.504693338 | 1.749698455 | 0.807106308 |
| P54803 | GALC | | 0 | 0 | 47270000 | 0 | 0 | 65124000 | 15756666.67 | 21708000 | 0.835283264 | 1.37770256 | 0.46226445 |
| Q8N428 | GALNT16 | | 0 | 38771000 | 0 | 28687000 | 75299000 | 0 | 12923666.67 | 34662000 | 0.441394278 | 2.682056176 | 1.423339455 |
| Q10471 | GALNT2 | | 886480000 | 1568400000 | 1360200000 | 1449200000 | 1790200000 | 1883400000 | 1271693333 | 1707600000 | 0.144877145 | 1.34277656 | 0.425219259 |
| Q14697 | GANAB | | 7768200000 | 7761400000 | 7877300000 | 9166300000 | 9853700000 | 9052800000 | 7802300000 | 9357600000 | 0.003551747 | 1.199338657 | 0.262239089 |
| P04406 | GAPDH | | 67507000000 | 61777000000 | 58236000000 | 66194000000 | 69991000000 | 86972000000 | 62506666667 | 74385666667 | 0.161912204 | 1.190043729 | 0.251014587 |
| P41250 | GARS1 | | 494250000 | 323890000 | 434080000 | 247940000 | 187150000 | 469190000 | 417406666.7 | 301426666.7 | 0.307078937 | 0.722141477 | -0.469646588 |
| P22102 | GART | | 207460000 | 249120000 | 60947000 | 169260000 | 171360000 | 371080000 | 172509000 | 237233333.3 | 0.502603421 | 1.375193951 | 0.459635104 |
| P04062 | GBA | | 919020000 | 858010000 | 670040000 | 703370000 | 550080000 | 765480000 | 815690000 | 672976666.7 | 0.221130128 | 0.825039741 | -0.277464481 |
| Q92538 | GBF1 | | 68211000 | 76616000 | 104470000 | 80430000 | 64843000 | 65604000 | 83099000 | 70292333.33 | 0.34867189 | 0.845886633 | -0.24146377 |
| Q8IWJ2 | GCC2 | | 0 | 0 | 0 | 48540000 | 77489000 | 63914000 | 0 | 63314333.33 | 0.001631333 | N/A | N/A |
| Q92616 | GCN1 | | 326920000 | 217320000 | 235850000 | 308780000 | 324180000 | 348500000 | 260030000 | 327153333.3 | 0.133973067 | 1.258136882 | 0.331288892 |
| P50395 | GDI2 | | 87568000 | 66722000 | 0 | 211360000 | 317970000 | 280020000 | 51430000 | 269783333.3 | 0.00591772 | 5.245641325 | 2.391119165 |
| Q8TEQ6 | GEMIN5 | | 39821000 | 0 | 0 | 0 | 0 | 24787000 | 13273666.67 | 8262333.333 | 0.764621283 | 0.622460511 | -0.68394578 |
| O43681 | GET3 | | 514560000 | 224510000 | 301120000 | 175500000 | 329280000 | 675100000 | 346730000 | 393293333.3 | 0.799249404 | 1.134292773 | 0.181793064 |
| Q9UJY4 | GGA2 | | 144580000 | 34172000 | 0 | 0 | 16894000 | 0 | 59584000 | 5631333.333 | 0.287277256 | 0.094510831 | -3.403376523 |
| P38435 | GGCX | | 0 | 0 | 0 | 0 | 123650000 | 133290000 | 0 | 85646666.67 | 0.116676112 | N/A | N/A |
| Q8N2G8 | GHDC | | 0 | 30345000 | 0 | 0 | 22885000 | 0 | 10115000 | 7628333.333 | 0.853960489 | 0.754160488 | -0.407056529 |
| Q6Y7W6 | GIGYF2 | | 61247000 | 139050000 | 0 | 0 | 0 | 206060000 | 66765666.67 | 68686666.67 | 0.981903112 | 1.028772273 | 0.040923666 |
| Q9Y2X7 | GIT1 | | 0 | 59153000 | 68448000 | 63406000 | 74498000 | 50022000 | 42533666.67 | 62642000 | 0.423371967 | 1.472762753 | 0.558525045 |
| P06280 | GLA | | 215640000 | 257490000 | 301000000 | 0 | 244870000 | 341670000 | 258043333.3 | 195513333.3 | 0.582226747 | 0.757676359 | -0.400346361 |
| P16278 | GLB1 | | 228450000 | 295740000 | 367600000 | 348020000 | 439870000 | 411100000 | 297263333.3 | 399663333.3 | 0.10221274 | 1.344475717 | 0.427043699 |
| Q92896 | GLG1 | | 394520000 | 438180000 | 381610000 | 826620000 | 742060000 | 578770000 | 404770000 | 715816666.7 | 0.014117834 | 1.768452866 | 0.822487768 |
| Q9H4G4 | GLIPR2 | | 213110000 | 233920000 | 331630000 | 242180000 | 205660000 | 149400000 | 259553333.3 | 199080000 | 0.25386352 | 0.767009992 | -0.382682724 |
| O76003 | GLRX3 | | 251130000 | 269030000 | 0 | 951690000 | 1131600000 | 0 | 173386666.7 | 694430000 | 0.223102737 | 4.005094586 | 2.001836314 |
| P17900 | GM2A | | 148010000 | 104290000 | 38530000 | 119730000 | 261420000 | 181250000 | 96943333.33 | 187466666.7 | 0.156153413 | 1.933775745 | 0.951420499 |
| P29992 | GNA11 | | 158890000 | 114790000 | 156490000 | 241000000 | 242240000 | 272320000 | 143390000 | 251853333.3 | 0.003520381 | 1.75642188 | 0.812639411 |
| Q03113 | GNA12 | | 0 | 0 | 50827000 | 0 | 68075000 | 55425000 | 16942333.33 | 41166666.67 | 0.418883384 | 2.429810927 | 1.280844057 |
| Q14344 | GNA13 | | 344880000 | 404080000 | 416090000 | 215510000 | 166760000 | 239560000 | 388350000 | 207276666.7 | 0.004137928 | 0.533736749 | -0.905799746 |
| P63096 | GNAI1 | | 248000000 | 206570000 | 252860000 | 0 | 125040000 | 161470000 | 235810000 | 95503333.33 | 0.051470021 | 0.405001202 | -1.304001907 |
| P04899 | GNAI2 | | 2924700000 | 2510800000 | 2491500000 | 2622700000 | 2975400000 | 2969500000 | 2642333333 | 2855866667 | 0.308525912 | 1.080812413 | 0.112116149 |
| P08754 | GNAI3 | | 1650300000 | 1659200000 | 1614500000 | 1376300000 | 1169000000 | 1283800000 | 1641333333 | 1276366667 | 0.004040571 | 0.77764013 | -0.362825424 |
| P09471 | GNAO1 | | 234690000 | 258740000 | 251690000 | 136270000 | 183430000 | 0 | 248373333.3 | 106566666.7 | 0.062831581 | 0.429058407 | -1.220754043 |
| P63092 | GNAS | | 544260000 | 506670000 | 492080000 | 614850000 | 661980000 | 612740000 | 514336666.7 | 629856666.7 | 0.006667919 | 1.22459997 | 0.292310553 |
| P62873 | GNB1 | | 3016200000 | 3173400000 | 2269100000 | 2063300000 | 2138900000 | 2706600000 | 2819566667 | 2302933333 | 0.208619357 | 0.816768534 | -0.292000807 |
| P62879 | GNB2 | | 4002000000 | 3404100000 | 4732000000 | 4805400000 | 4209900000 | 3711600000 | 4046033333 | 4242300000 | 0.713274877 | 1.048508416 | 0.068338441 |
| Q9HAV0 | GNB4 | | 233430000 | 175910000 | 181490000 | 0 | 0 | 0 | 196943333.3 | 0 | 0.000423952 | 0 | N/A |
| P50151 | GNG10 | | 0 | 0 | 0 | 0 | 116940000 | 97480000 | 0 | 71473333.33 | 0.119382445 | N/A | N/A |

| | | | | | | | | | | | | | |
|---|---|---|---|---|---|---|---|---|---|---|---|---|---|
| Q9UBI6 | GNG12 | | 316710000 | 677200000 | 397960000 | 519460000 | 464440000 | 173760000 | 463956666.7 | 385886666.7 | 0.636813091 | 0.831729975 | -0.265812869 |
| P63218 | GNG5 | | 16108000 | 37589000 | 29592000 | 0 | 18251000 | 8489100 | 27763000 | 8913366.667 | 0.082823808 | 0.321052 | -1.63912111 |
| P15586 | GNS | | 795300000 | 848890000 | 774600000 | 1381400000 | 1788400000 | 1905400000 | 806263333.3 | 1691733333 | 0.005249785 | 2.098239202 | 1.069179157 |
| Q08379 | GOLGA2 | | 227300000 | 235150000 | 213280000 | 386140000 | 312530000 | 159710000 | 225243333.3 | 286126666.7 | 0.414844638 | 1.270300268 | 0.345169555 |
| Q13439 | GOLGA4 | | 79030000 | 72993000 | 73473000 | 296780000 | 229930000 | 44528000 | 75165333.33 | 190412666.7 | 0.20148909 | 2.533251144 | 1.340990112 |
| Q8TBA6 | GOLGA5 | | 0 | 0 | 101680000 | 107400000 | 88684000 | 0 | 33893333.33 | 65361333.33 | 0.543009353 | 1.928442172 | 0.947435884 |
| Q14789 | GOLGB1 | | 1930500000 | 1831600000 | 1942800000 | 1688500000 | 1804900000 | 1709500000 | 1901633333 | 1734300000 | 0.02904352 | 0.912005469 | -0.132885619 |
| O00461 | GOLIM4 | | 543520000 | 720110000 | 651720000 | 1543400000 | 1915000000 | 561380000 | 638450000 | 1339926667 | 0.15992191 | 2.09871825 | 1.0695085 |
| O95249 | GOSR1 | | 0 | 382440000 | 399000000 | 491050000 | 503930000 | 635680000 | 260480000 | 543553333.3 | 0.11004921 | 2.086737305 | 1.061248994 |
| O14653 | GOSR2 | | 47976000 | 177110000 | 0 | 158590000 | 133350000 | 0 | 75028666.67 | 97313333.33 | 0.773091276 | 1.297015363 | 0.375195568 |
| O43292 | GPAA1 | | 0 | 0 | 50996000 | 0 | 0 | 43215000 | 16998666.67 | 14405000 | 0.912941638 | 0.847419405 | -0.238851929 |
| P35052 | GPC1 | | 1950400000 | 1913300000 | 2487800000 | 1831900000 | 1950100000 | 1981600000 | 2117166667 | 1921200000 | 0.363188886 | 0.907439188 | -0.140127131 |
| O75487 | GPC4 | | 142760000 | 199180000 | 249210000 | 248640000 | 150390000 | 65254000 | 197050000 | 154761333.3 | 0.527960051 | 0.785391187 | -0.348516686 |
| Q9Y625 | GPC6 | | 85696000 | 0 | 0 | 107720000 | 123180000 | 80294000 | 28565333.33 | 103731333.3 | 0.073599807 | 3.631371359 | 1.860514474 |
| Q8NFJ5 | GPRC5A | | 220650000 | 217540000 | 301150000 | 259400000 | 262650000 | 230320000 | 246446666.7 | 250790000 | 0.889076386 | 1.017623827 | 0.025204355 |
| Q8TD30 | GPT2 | | 127800000 | 169800000 | 178090000 | 163090000 | 166090000 | 148130000 | 158563333.3 | 159103333.3 | 0.975501639 | 1.003405579 | 0.004904865 |
| P07203 | GPX1 | | 190450000 | 219930000 | 152410000 | 661850000 | 581570000 | 444290000 | 187596666.7 | 562570000 | 0.004857771 | 2.998827271 | 1.584398427 |
| Q8TED1 | GPX8 | | 202130000 | 280140000 | 278310000 | 0 | 0 | 0 | 253526666.7 | 0 | 0.000592726 | 0 | N/A |
| Q9UBQ7 | GRHPR | | 654690000 | 590040000 | 665930000 | 681160000 | 649630000 | 583050000 | 636886666.7 | 637946666.7 | 0.978721588 | 1.001664346 | 0.002399148 |
| P28799 | GRN | | 196670000 | 0 | 0 | 423720000 | 348460000 | 0 | 65556666.67 | 257393333.3 | 0.25930654 | 3.926272436 | 1.973160282 |
| Q12849 | GRSF1 | | 2641200000 | 2451200000 | 2305700000 | 1250000000 | 774870000 | 1334000000 | 2466033333 | 1119623333 | 0.002505444 | 0.454017924 | -1.139178842 |
| P49841 | GSK3B | | 108340000 | 122580000 | 77936000 | 77775000 | 104190000 | 97463000 | 102952000 | 93142666.67 | 0.557981787 | 0.904719351 | -0.144457765 |
| Q9Y2Q3 | GSTK1 | | 651360000 | 509020000 | 669270000 | 827340000 | 667490000 | 615950000 | 609883333.3 | 703593333.3 | 0.31357322 | 1.153652338 | 0.206208522 |
| P09211 | GSTP1 | | 178780000 | 218420000 | 154920000 | 992960000 | 858650000 | 630050000 | 184040000 | 827220000 | 0.003928517 | 4.494783743 | 2.168251705 |
| O43708 | GSTZ1 | | 291320000 | 247550000 | 291920000 | 110580000 | 69811000 | 0 | 276930000 | 60130333.33 | 0.003627954 | 0.217131887 | -2.203356489 |
| Q4G148 | GXYLT1 | | 0 | 76115000 | 91737000 | 155760000 | 190570000 | 92718000 | 55950666.67 | 146349333.3 | 0.088240911 | 2.615685247 | 1.387188947 |
| P07305 | H1 | | 62797000 | 0 | 0 | 116720000 | 132420000 | 0 | 20932333.33 | 83046666.67 | 0.254448764 | 3.967386977 | 1.988189124 |
| Q02539 | H1 | | 296000000 | 189980000 | 503600000 | 366860000 | 373930000 | 194230000 | 329860000 | 311673333.3 | 0.875863929 | 0.944865498 | -0.081819118 |
| Q92522 | H1 | | 176420000 | 290160000 | 346150000 | 0 | 0 | 0 | 270910000 | 0 | 0.005596763 | 0 | N/A |
| P16403 | H1 | | 74152000 | 0 | 266910000 | 540580000 | 276080000 | 211560000 | 113687333.3 | 342740000 | 0.148752195 | 3.01475978 | 1.592043051 |
| P10412 | H1 | | 364260000 | 530160000 | 554210000 | 1522900000 | 1011400000 | 584510000 | 482876666.7 | 1039603333 | 0.115533946 | 2.152937603 | 1.106306508 |
| Q99878 | H2AC14 | | 8009900000 | 6980600000 | 7842200000 | 15283000000 | 10428000000 | 7749300000 | 7610900000 | 11153433333 | 0.18700198 | 1.465455246 | 0.55134891 |
| Q16777 | H2AC20 | | 178270000 | 0 | 469450000 | 0 | 202780000 | 373630000 | 215906666.7 | 192136666.7 | 0.898114676 | 0.889906132 | -0.168274927 |
| Q93077 | H2AC6 | | 161320000 | 221100000 | 111410000 | 148390000 | 0 | 0 | 164610000 | 49463333.33 | 0.121579284 | 0.300488022 | -1.73462061 |
| P16104 | H2AX | | 392400000 | 304200000 | 228130000 | 275600000 | 232450000 | 267150000 | 308243333.3 | 258400000 | 0.368896494 | 0.838298747 | -0.254463622 |
| Q71UI9 | H2AZ2 | | 620000000 | 487870000 | 1002400000 | 1359900000 | 1012400000 | 1576800000 | 703423333.3 | 1316366667 | 0.053045243 | 1.87137191 | 0.904096303 |
| Q99879 | H2BC14 | | 4078600000 | 5418000000 | 6006700000 | 15245000000 | 12217000000 | 6230500000 | 5167766667 | 11230833333 | 0.088832024 | 2.173246986 | 1.119852144 |
| Q16778 | H2BC21 | | 0 | 216740000 | 325500000 | 714670000 | 605520000 | 401200000 | 180746666.7 | 573796666.7 | 0.041418907 | 3.174590587 | 1.666570546 |
| P57053 | H2BS1 | | 0 | 0 | 0 | 64296000 | 49088000 | 0 | 0 | 37794666.67 | 0.123222872 | N/A | N/A |
| P84243 | H3 | | 63926000 | 55263000 | 0 | 360010000 | 593040000 | 218330000 | 39729666.67 | 390460000 | 0.034244591 | 9.827920362 | 3.296886167 |
| P68431 | H3C1 | | 233470000 | 294680000 | 184830000 | 545410000 | 301510000 | 154460000 | 237660000 | 333793333.3 | 0.46221517 | 1.404499425 | 0.490056034 |
| Q71DI3 | H3C15 | | 0 | 0 | 0 | 207240000 | 170610000 | 252140000 | 0 | 209996666.7 | 0.000878099 | N/A | N/A |
| P62805 | H4C1 | | 2300200000 | 2448800000 | 2638300000 | 4382700000 | 4475600000 | 3044300000 | 2462433333 | 3967533333 | 0.03339112 | 1.611224669 | 0.688157678 |
| P12081 | HARS1 | | 0 | 0 | 0 | 75451000 | 72003000 | 0 | 0 | 49151333.33 | 0.116333964 | N/A | N/A |
| O00165 | HAX1 | | 335600000 | 264030000 | 327390000 | 162870000 | 224250000 | 161700000 | 309006666.7 | 182940000 | 0.014659757 | 0.592026062 | -0.756267408 |
| Q9BXW7 | HDHD5 | | 0 | 232740000 | 206150000 | 208820000 | 273150000 | 274420000 | 146296666.7 | 252130000 | 0.239607696 | 1.72341589 | 0.785270891 |
| Q00341 | HDLBP | | 512670000 | 423830000 | 453530000 | 999230000 | 1118900000 | 1254200000 | 463343333.3 | 1124110000 | 0.001071607 | 2.426084329 | 1.278629698 |
| Q9H583 | HEATR1 | | 52100000 | 0 | 0 | 0 | 31162000 | 22421000 | 17366666.67 | 17861000 | 0.981173747 | 1.028464491 | 0.040491984 |
| Q9NRV9 | HEBP1 | | 279200000 | 276430000 | 372760000 | 163520000 | 127190000 | 137280000 | 309463333.3 | 142663333.3 | 0.007569166 | 0.46100238 | -1.117153895 |
| P06865 | HEXA | | 133150000 | 0 | 132460000 | 84726000 | 232550000 | 368880000 | 88536666.67 | 228718666.7 | 0.20711752 | 2.583321411 | 1.369227152 |
| P07686 | HEXB | | 283400000 | 199480000 | 255990000 | 205180000 | 250290000 | 362040000 | 246290000 | 272503333.3 | 0.645400437 | 1.106432796 | 0.145915825 |
| O14964 | HGS | | 0 | 0 | 40903000 | 194440000 | 134860000 | 0 | 13634333.33 | 109766666.7 | 0.179202158 | 8.050754223 | 3.009123946 |
| O00291 | HIP1 | | 156470000 | 124070000 | 166590000 | 102720000 | 0 | 72684000 | 149043333.3 | 58468000 | 0.0520069 | 0.392288596 | -1.350012698 |
| O75146 | HIP1R | | 146090000 | 262620000 | 110290000 | 204090000 | 380820000 | 191780000 | 173000000 | 258896666.7 | 0.32403483 | 1.496512524 | 0.581604353 |
| P19367 | HK1 | | 2923100000 | 2781600000 | 2869100000 | 2395700000 | 2765900000 | 2219900000 | 2857933333 | 2460500000 | 0.074956274 | 0.860936807 | -0.216020747 |
| P10321 | HLA | | 0 | 0 | 0 | 17978000 | 7965300 | 0 | 0 | 8647766.667 | 0.171704695 | N/A | N/A |
| P17096 | HMGA1 | | 0 | 0 | 0 | 229440000 | 0 | 158070000 | 0 | 129170000 | 0.129430428 | N/A | N/A |

| | | | | | | | | | | | | | |
|---|---|---|---|---|---|---|---|---|---|---|---|---|---|
| P09429 | HMGB1 | | 729480000 | 737720000 | 645190000 | 1019100000 | 1075400000 | 474020000 | 704130000 | 856173333.3 | 0.477075461 | 1.21593077 | 0.282061091 |
| P26583 | HMGB2 | | 194910000 | 312970000 | 290180000 | 364350000 | 277950000 | 100640000 | 266020000 | 247646666.7 | 0.840597636 | 0.930932511 | -0.103251513 |
| P30519 | HMOX2 | | 563890000 | 491420000 | 460390000 | 444720000 | 513630000 | 584880000 | 505233333.3 | 514410000 | 0.865352953 | 1.018163225 | 0.025968863 |
| Q13151 | HNRNPA0 | | 171240000 | 117560000 | 123160000 | 94482000 | 131270000 | 138950000 | 137320000 | 121567333.3 | 0.511308443 | 0.885284979 | -0.175786152 |
| P09651 | HNRNPA1 | | 683670000 | 319200000 | 182830000 | 434390000 | 579830000 | 540130000 | 395233333.3 | 518116666.7 | 0.474041182 | 1.310913384 | 0.390572366 |
| P22626 | HNRNPA2B1 | | 1630700000 | 1832100000 | 1380000000 | 2686000000 | 1479600000 | 1520100000 | 1614266667 | 1895233333 | 0.537020337 | 1.174052201 | 0.231496556 |
| P51991 | HNRNPA3 | | 295790000 | 260360000 | 334100000 | 483610000 | 351060000 | 299150000 | 296750000 | 377940000 | 0.240157832 | 1.273597304 | 0.348909187 |
| Q99729 | HNRNPAB | | 0 | 57520000 | 0 | 129700000 | 169440000 | 0 | 19173333.33 | 99713333.33 | 0.214448539 | 5.200625869 | 2.378685255 |
| P07910 | HNRNPC | | 428980000 | 293540000 | 365020000 | 467700000 | 581290000 | 506960000 | 362513333.3 | 518650000 | 0.038439056 | 1.430705997 | 0.516727236 |
| Q14103 | HNRNPD | | 1024200000 | 1040800000 | 1254700000 | 886900000 | 959590000 | 976360000 | 1106566667 | 940950000 | 0.104500553 | 0.850332861 | -0.233900402 |
| O14979 | HNRNPDL | | 286180000 | 261130000 | 361800000 | 512700000 | 553010000 | 531450000 | 303036666.7 | 532386666.7 | 0.002107534 | 1.756839106 | 0.812982073 |
| P52597 | HNRNPF | | 308170000 | 95338000 | 197140000 | 121020000 | 0 | 356030000 | 200216000 | 159016666.7 | 0.751105451 | 0.79422557 | -0.332379286 |
| P31943 | HNRNPH1 | | 771840000 | 420580000 | 546270000 | 524800000 | 387720000 | 580330000 | 579563333.3 | 497616666.7 | 0.52436538 | 0.858606192 | -0.219931518 |
| P31942 | HNRNPH3 | | 0 | 169070000 | 132220000 | 192710000 | 140010000 | 169760000 | 100430000 | 167493333.3 | 0.27865562 | 1.667761957 | 0.737913385 |
| P61978 | HNRNPK | | 2058000000 | 1712400000 | 1679000000 | 2773700000 | 2378100000 | 1987000000 | 1816466667 | 2379600000 | 0.093921791 | 1.310015782 | 0.389584192 |
| P14866 | HNRNPL | | 269550000 | 288840000 | 335090000 | 374930000 | 379450000 | 403930000 | 297826666.7 | 386103333.3 | 0.014623568 | 1.296402829 | 0.374514074 |
| P52272 | HNRNPM | | 994060000 | 955410000 | 824800000 | 1021900000 | 956880000 | 1021900000 | 924756666.7 | 1000226667 | 0.246240974 | 1.081610658 | 0.113181273 |
| O43390 | HNRNPR | | 0 | 205440000 | 187790000 | 0 | 249840000 | 365890000 | 131076666.7 | 205243333.3 | 0.588857055 | 1.565826615 | 0.646924471 |
| Q00839 | HNRNPU | | 1149900000 | 1009900000 | 1112600000 | 1336200000 | 1409600000 | 1269400000 | 1090800000 | 1338400000 | 0.013140109 | 1.226989366 | 0.295122745 |
| Q9BUJ2 | HNRNPUL1 | | 152920000 | 262370000 | 212300000 | 70228000 | 73688000 | 0 | 209196666.7 | 47972000 | 0.01535008 | 0.229315317 | -2.124595371 |
| Q9Y251 | HPSE | | 83044000 | 0 | 66318000 | 46617000 | 51019000 | 69221000 | 49787333.33 | 55619000 | 0.835284516 | 1.117131533 | 0.159799061 |
| P01112 | HRAS | | 318710000 | 345230000 | 284940000 | 370870000 | 367190000 | 445300000 | 316293333.3 | 394453333.3 | 0.064431861 | 1.247112385 | 0.318591481 |
| Q7LGA3 | HS2ST1 | | 99232000 | 0 | 115880000 | 0 | 0 | 0 | 71704000 | 0 | 0.118493466 | 0 | N/A |
| Q53GQ0 | HSD17B12 | | 0 | 0 | 0 | 157620000 | 177300000 | 166500000 | 0 | 167140000 | 7.9978E-06 | N/A | N/A |
| P51659 | HSD17B4 | | 3149600000 | 2453700000 | 2829100000 | 1755900000 | 1895800000 | 2012700000 | 2810800000 | 1888133333 | 0.012602887 | 0.671742327 | -0.574020157 |
| Q3SXM5 | HSDL1 | | 62992000 | 37203000 | 46521000 | 77971000 | 117210000 | 78745000 | 48905333.33 | 91308666.67 | 0.047377768 | 1.867049265 | 0.900759996 |
| Q6YN16 | HSDL2 | | 629580000 | 597490000 | 404870000 | 443440000 | 410000000 | 419520000 | 543980000 | 424320000 | 0.166599995 | 0.780028678 | -0.35840093 |
| P07900 | HSP90AA1 | | 637180000 | 767500000 | 782020000 | 1131600000 | 1310200000 | 1211400000 | 728900000 | 1217733333 | 0.002118722 | 1.670645265 | 0.740405432 |
| P08238 | HSP90AB1 | | 4074400000 | 4272500000 | 4519400000 | 7117200000 | 7766100000 | 7473500000 | 4288766667 | 7452266667 | 0.00015517 | 1.737624647 | 0.797116472 |
| Q58FF8 | HSP90AB2P | | 183710000 | 151990000 | 139320000 | 235790000 | 297440000 | 0 | 158340000 | 177743333.3 | 0.842585106 | 1.122542209 | 0.166769693 |
| P14625 | HSP90B1 | | 25351000000 | 24275000000 | 26088000000 | 18632000000 | 20451000000 | 21094000000 | 25238000000 | 20059000000 | 0.004631323 | 0.794793565 | -0.331347902 |
| P48723 | HSPA13 | | 312320000 | 289240000 | 329790000 | 272310000 | 0 | 342310000 | 310450000 | 204873333.3 | 0.371831754 | 0.659923767 | -0.599628719 |
| P0DMV9 | HSPA1B | | 1916700000 | 1879200000 | 1900700000 | 2349400000 | 2585100000 | 2516100000 | 1898866667 | 2483533333 | 0.001173158 | 1.30790296 | 0.387255504 |
| P34932 | HSPA4 | | 76828000 | 203170000 | 332650000 | 641360000 | 774260000 | 1089000000 | 204216000 | 834873333.3 | 0.014236825 | 4.088187671 | 2.031461426 |
| P11021 | HSPA5 | | 31020000000 | 29863000000 | 31673000000 | 36355000000 | 36370000000 | 31847000000 | 30852000000 | 34857333333 | 0.066029248 | 1.129824106 | 0.176098188 |
| P17066 | HSPA6 | | 720650000 | 1368300000 | 827290000 | 962500000 | 481350000 | 214340000 | 972080000 | 552730000 | 0.23060885 | 0.568605465 | -0.814500131 |
| P11142 | HSPA8 | | 12572000000 | 12891000000 | 12854000000 | 12778000000 | 13346000000 | 12861000000 | 12772333333 | 12995000000 | 0.335906299 | 1.017433515 | 0.024934524 |
| P04792 | HSPB1 | | 1609700000 | 1903500000 | 1383800000 | 3470300000 | 3631200000 | 3336700000 | 1632333333 | 3479400000 | 0.000434654 | 2.131549929 | 1.091902849 |
| Q92598 | HSPH1 | | 140520000 | 197590000 | 196010000 | 326000000 | 374710000 | 298360000 | 178040000 | 333023333.3 | 0.006025477 | 1.870497267 | 0.903421858 |
| Q9BUP3 | HTATIP2 | | 0 | 114990000 | 80903000 | 147280000 | 218670000 | 262350000 | 65297666.67 | 209433333.3 | 0.039409764 | 3.207363203 | 1.681387735 |
| Q7Z6Z7 | HUWE1 | | 37693000 | 150730000 | 76462000 | 121490000 | 31408000 | 109500000 | 88295000 | 87466000 | 0.985727347 | 0.99061102 | -0.013609425 |
| Q9Y4L1 | HYOU1 | | 2382100000 | 2533700000 | 2637600000 | 2794000000 | 2699800000 | 3851900000 | 2517800000 | 3115233333 | 0.187942767 | 1.237283872 | 0.307176538 |
| P41252 | IARS1 | | 506540000 | 444130000 | 436920000 | 364250000 | 400200000 | 490630000 | 462530000 | 418360000 | 0.368441171 | 0.904503492 | -0.144802023 |
| Q9Y6M1 | IGF2BP2 | | 649920000 | 725340000 | 596210000 | 1021700000 | 1032000000 | 862840000 | 657156666.7 | 972180000 | 0.008977507 | 1.47937326 | 0.564986104 |
| O00425 | IGF2BP3 | | 0 | 135220000 | 72919000 | 122610000 | 47893000 | 164440000 | 69379666.67 | 111647666.7 | 0.460743955 | 1.609227487 | 0.686368286 |
| P11717 | IGF2R | | 160320000 | 165820000 | 154750000 | 194610000 | 190170000 | 182780000 | 160296666.7 | 189186666.7 | 0.003560522 | 1.180228327 | 0.23906599 |
| Q969P0 | IGSF8 | | 190580000 | 160160000 | 125660000 | 163330000 | 168620000 | 82506000 | 158800000 | 138152000 | 0.571968421 | 0.869974811 | -0.200954465 |
| Q14116 | IL18 | | 0 | 0 | 0 | 51104000 | 60484000 | 24999000 | 0 | 45529000 | 0.01275968 | N/A | N/A |
| Q12905 | ILF2 | | 209980000 | 148470000 | 0 | 0 | 127500000 | 0 | 119483333.3 | 42500000 | 0.365185399 | 0.355698145 | -1.491274645 |
| Q12906 | ILF3 | | 79724000 | 114900000 | 129110000 | 187930000 | 126350000 | 56777000 | 107911333.3 | 123685666.7 | 0.717602444 | 1.146178653 | 0.196831932 |
| Q13418 | ILK | | 119810000 | 108240000 | 117500000 | 51435000 | 77917000 | 56870000 | 115183333.3 | 62074000 | 0.003824494 | 0.538914774 | -0.891870958 |
| A1L0T0 | ILVBL | | 386790000 | 197160000 | 311210000 | 291530000 | 339880000 | 386600000 | 298386666.7 | 339336666.7 | 0.54238213 | 1.137238036 | 0.185534257 |
| P12268 | IMPDH2 | | 1347800000 | 1570100000 | 1467300000 | 1561900000 | 1865800000 | 1510100000 | 1461733333 | 1645933333 | 0.224123847 | 1.126014777 | 0.17122576 |
| Q8TEX9 | IPO4 | | 0 | 0 | 56645000 | 84128000 | 115160000 | 129730000 | 18881666.67 | 109672666.7 | 0.017296039 | 5.808420867 | 2.538145992 |
| O00410 | IPO5 | | 662510000 | 718360000 | 847660000 | 773640000 | 1158900000 | 664500000 | 742843333.3 | 865680000 | 0.484609694 | 1.165360125 | 0.220775852 |
| Q96P70 | IPO9 | | 263440000 | 253470000 | 120050000 | 304490000 | 256370000 | 328400000 | 212320000 | 296420000 | 0.173470705 | 1.396100226 | 0.481402516 |
| P46940 | IQGAP1 | | 1211400000 | 1285300000 | 1208100000 | 362520000 | 116370000 | 117570000 | 1234933333 | 198820000 | 0.000267765 | 0.160996545 | -2.634898366 |

| | | | | | | | | | | | | | |
|---|---|---|---|---|---|---|---|---|---|---|---|---|---|
| Q96AB3 | ISOC2 | | 0 | 0 | 0 | 49591000 | 106890000 | 0 | 0 | 52160333.33 | 0.16649698 | N/A | N/A |
| P17301 | ITGA2 | | 212870000 | 202700000 | 49129000 | 315810000 | 220200000 | 379750000 | 154899666.7 | 305253333.3 | 0.09953165 | 1.97065197 | 0.97867301 |
| P26006 | ITGA3 | | 178240000 | 220080000 | 295480000 | 524640000 | 466620000 | 662870000 | 231266666.7 | 551376666.7 | 0.009051492 | 2.3841597 | 1.253480876 |
| P08648 | ITGA5 | | 305660000 | 326500000 | 411510000 | 406900000 | 497320000 | 446030000 | 347890000 | 450083333.3 | 0.070105413 | 1.293751856 | 0.371560933 |
| P23229 | ITGA6 | | 264650000 | 337860000 | 261600000 | 546230000 | 542380000 | 522700000 | 288036666.7 | 537103333.3 | 0.000660644 | 1.864704725 | 0.898947199 |
| P06756 | ITGAV | | 1385900000 | 1539900000 | 1329400000 | 1384200000 | 1476600000 | 1179400000 | 1418400000 | 1346733333 | 0.543319157 | 0.949473585 | -0.07480023 |
| P05556 | ITGB1 | | 4702100000 | 4787400000 | 4391500000 | 4004000000 | 3939100000 | 3092900000 | 4627000000 | 3678666667 | 0.04034244 | 0.795043585 | -0.330894143 |
| P05106 | ITGB3 | | 0 | 0 | 0 | 62849000 | 0 | 27503000 | 0 | 30117333.33 | 0.173122163 | N/A | N/A |
| P16144 | ITGB4 | | 138310000 | 118350000 | 113810000 | 0 | 0 | 0 | 123490000 | 0 | 8.07192E-05 | 0 | N/A |
| P18084 | ITGB5 | | 610530000 | 499960000 | 666510000 | 799620000 | 852430000 | 932520000 | 592333333.3 | 861523333.3 | 0.012469514 | 1.45445695 | 0.540480595 |
| Q9NZM3 | ITSN2 | | 0 | 0 | 31443000 | 0 | 0 | 96849000 | 10481000 | 32283000 | 0.555619875 | 3.080145024 | 1.62299828 |
| P14923 | JUP | | 324950000 | 303250000 | 315050000 | 108820000 | 166480000 | 247580000 | 314416666.7 | 174293333.3 | 0.026291526 | 0.554338723 | -0.851160306 |
| Q63ZY3 | KANK2 | | 50441000 | 37828000 | 32642000 | 0 | 0 | 0 | 40303666.67 | 0 | 0.00158774 | 0 | N/A |
| Q15046 | KARS1 | | 1000300000 | 1063200000 | 1018900000 | 1290300000 | 1142500000 | 1276600000 | 1027466667 | 1236466667 | 0.014591242 | 1.203412925 | 0.267131757 |
| Q06136 | KDSR | | 0 | 31513000 | 0 | 0 | 58104000 | 0 | 10504333.33 | 19368000 | 0.708042758 | 1.843810491 | 0.882690381 |
| Q07666 | KHDRBS1 | | 0 | 0 | 79621000 | 67417000 | 0 | 61681000 | 26540333.33 | 43032666.67 | 0.654900851 | 1.621406413 | 0.697245754 |
| Q92945 | KHSRP | | 136840000 | 116550000 | 0 | 120380000 | 164420000 | 166710000 | 84463333.33 | 150503333.3 | 0.21798658 | 1.781877738 | 0.833398351 |
| Q9P206 | KIAA1522 | | 62486000 | 0 | 63528000 | 0 | 0 | 0 | 42004666.67 | 0 | 0.116143719 | 0 | N/A |
| Q8IYS2 | KIAA2013 | | 141560000 | 150850000 | 165530000 | 173270000 | 204390000 | 148480000 | 152646666.7 | 175380000 | 0.266402791 | 1.148927807 | 0.200288149 |
| Q9ULH0 | KIDINS220 | | 354210000 | 157710000 | 302480000 | 242810000 | 161190000 | 314450000 | 271466666.7 | 239483333.3 | 0.686335771 | 0.882183202 | -0.180849805 |
| P52732 | KIF11 | | 76781000 | 112610000 | 101680000 | 0 | 0 | 0 | 97023666.67 | 0 | 0.000791309 | 0 | N/A |
| Q02241 | KIF23 | | 121710000 | 166610000 | 119250000 | 99177000 | 0 | 0 | 135856666.7 | 33059000 | 0.047880573 | 0.243337341 | -2.038970372 |
| O00139 | KIF2A | | 0 | 54735000 | 0 | 0 | 0 | 44697000 | 18245000 | 14899000 | 0.893909722 | 0.81660729 | -0.292285649 |
| O95239 | KIF4A | | 0 | 59023000 | 0 | 0 | 43992000 | 0 | 19674333.33 | 14664000 | 0.848175588 | 0.745336564 | -0.42403606 |
| P33176 | KIF5B | | 402860000 | 391000000 | 290330000 | 487960000 | 415670000 | 363830000 | 361396666.7 | 422486666.7 | 0.294599978 | 1.169038637 | 0.225322612 |
| P52292 | KPNA2 | | 244320000 | 230780000 | 284920000 | 176730000 | 167460000 | 186310000 | 253340000 | 176833333.3 | 0.011157445 | 0.698007947 | -0.518684633 |
| O60684 | KPNA6 | | 56420000 | 0 | 40215000 | 0 | 0 | 0 | 32211666.67 | 0 | 0.127179877 | 0 | N/A |
| Q14974 | KPNB1 | | 700410000 | 676080000 | 727520000 | 628380000 | 655700000 | 603760000 | 701336666.7 | 629280000 | 0.026953226 | 0.897258093 | -0.156405064 |
| Q13601 | KRR1 | | 0 | 0 | 0 | 0 | 108000000 | 108100000 | 0 | 72033333.33 | 0.116116609 | N/A | N/A |
| Q86UP2 | KTN1 | | 6112300000 | 5782400000 | 5907500000 | 6820900000 | 6742000000 | 5441300000 | 5934066667 | 6334733333 | 0.430608483 | 1.067519745 | 0.094262754 |
| Q53H82 | LACTB2 | | 0 | 0 | 91079000 | 84754000 | 0 | 0 | 30359666.67 | 28251333.33 | 0.9618915 | 0.930554793 | -0.103836993 |
| P55268 | LAMB2 | | 126030000 | 146440000 | 154890000 | 0 | 0 | 0 | 142453333.3 | 0 | 7.66017E-05 | 0 | N/A |
| Q13751 | LAMB3 | | 555750000 | 441340000 | 319990000 | 217430000 | 288130000 | 174890000 | 439026666.7 | 226816666.7 | 0.048567333 | 0.516635284 | -0.952781918 |
| P11047 | LAMC1 | | 97529000 | 112190000 | 63170000 | 135050000 | 150240000 | 128110000 | 90963000 | 137800000 | 0.042372401 | 1.514901663 | 0.599224147 |
| Q13753 | LAMC2 | | 0 | 0 | 0 | 101660000 | 0 | 30234000 | 0 | 43964666.67 | 0.218397792 | N/A | N/A |
| P11279 | LAMP1 | | 881170000 | 683740000 | 883850000 | 378550000 | 736600000 | 832380000 | 816253333.3 | 649176666.7 | 0.336701139 | 0.79531273 | -0.330405832 |
| Q6IAA8 | LAMTOR1 | | 976440000 | 900760000 | 945440000 | 1687700000 | 1818300000 | 2506400000 | 940880000 | 2004133333 | 0.01401614 | 2.130062636 | 1.090895855 |
| Q9Y2Q5 | LAMTOR2 | | 264900000 | 231750000 | 69135000 | 68719000 | 404860000 | 385580000 | 188595000 | 286386333.3 | 0.47655292 | 1.518525588 | 0.602671219 |
| Q9UHA4 | LAMTOR3 | | 740660000 | 506560000 | 0 | 1159300000 | 1335300000 | 1530200000 | 415740000 | 1341600000 | 0.019044044 | 3.227016886 | 1.690201127 |
| Q0VGL1 | LAMTOR4 | | 229120000 | 374320000 | 330970000 | 143920000 | 142680000 | 692400000 | 311470000 | 326333333.3 | 0.940789654 | 1.047719952 | 0.067253146 |
| O43504 | LAMTOR5 | | 641170000 | 804170000 | 618150000 | 1346800000 | 986320000 | 720950000 | 687830000 | 1018023333 | 0.158202375 | 1.480050788 | 0.565646683 |
| P28838 | LAP3 | | 995950000 | 927560000 | 884410000 | 899630000 | 1041400000 | 1239400000 | 935973333.3 | 1060143333 | 0.297430786 | 1.132664036 | 0.179720001 |
| Q6PKG0 | LARP1 | | 0 | 0 | 0 | 61530000 | 36666000 | 0 | 0 | 32732000 | 0.140966353 | N/A | N/A |
| Q9P2J5 | LARS1 | | 249920000 | 275730000 | 305370000 | 228280000 | 303110000 | 329750000 | 277006666.7 | 287046666.7 | 0.784534304 | 1.036244615 | 0.051364605 |
| Q14847 | LASP1 | | 60503000 | 47106000 | 45619000 | 76207000 | 35039000 | 16383000 | 51076000 | 42543000 | 0.665196053 | 0.832935234 | -0.263723774 |
| Q9GZY6 | LAT2 | | 0 | 0 | 0 | 69732000 | 61547000 | 0 | 0 | 43759666.67 | 0.117660419 | N/A | N/A |
| Q6UX15 | LAYN | | 132190000 | 106940000 | 0 | 0 | 45116000 | 0 | 79710000 | 15038666.67 | 0.208878931 | 0.188667252 | -2.406084066 |
| Q14739 | LBR | | 368250000 | 0 | 314640000 | 216360000 | 148860000 | 245620000 | 227630000 | 203613333.3 | 0.849134396 | 0.894492524 | -0.16085867 |
| Q6UWP7 | LCLAT1 | | 67350000 | 64334000 | 51221000 | 86755000 | 0 | 89612000 | 60968333.33 | 58789000 | 0.945248548 | 0.964254668 | -0.05251387 |
| P00338 | LDHA | | 141290000 | 130080000 | 0 | 271960000 | 230210000 | 181330000 | 90456666.67 | 227833333.3 | 0.058583699 | 2.518701404 | 1.332680098 |
| P07195 | LDHB | | 61391000 | 0 | 36481000 | 210970000 | 162060000 | 80650000 | 32624000 | 151226666.7 | 0.047578637 | 4.63544221 | 2.212706975 |
| P01130 | LDLR | | 77860000 | 49478000 | 48395000 | 0 | 0 | 0 | 58577666.67 | 0 | 0.003714953 | 0 | N/A |
| P09382 | LGALS1 | | 7576900000 | 7532500000 | 7382800000 | 9115300000 | 9426000000 | 8565200000 | 7497400000 | 9035500000 | 0.003999347 | 1.205151119 | 0.269214064 |
| Q99538 | LGMN | | 161430000 | 109840000 | 139440000 | 274670000 | 168610000 | 104340000 | 136903333.3 | 182540000 | 0.42856666 | 1.333349565 | 0.415055063 |
| Q9UHB6 | LIMA1 | | 1350600000 | 1183500000 | 1099200000 | 1254300000 | 1012000000 | 746890000 | 1211100000 | 1004396667 | 0.276280725 | 0.829325957 | -0.269988847 |
| P48059 | LIMS1 | | 0 | 84311000 | 75813000 | 45477000 | 91114000 | 60748000 | 53374666.67 | 65779666.67 | 0.700156183 | 1.232413629 | 0.301486543 |
| Q9NUP9 | LIN7C | | 498470000 | 737350000 | 962680000 | 496130000 | 504450000 | 449880000 | 732833333.3 | 483486666.7 | 0.138677832 | 0.659749829 | -0.600009022 |

| | | | | | | | | | | | | | |
|---|---|---|---|---|---|---|---|---|---|---|---|---|---|
| Q15334 | LLGL1 | | 90625000 | 102140000 | 124240000 | 132600000 | 126780000 | 118890000 | 105668333.3 | 126090000 | 0.127182535 | 1.193261936 | 0.254910767 |
| P49257 | LMAN1 | | 951020000 | 999060000 | 756390000 | 635850000 | 665500000 | 450150000 | 902156666.7 | 583833333.3 | 0.033662622 | 0.647152934 | -0.627821408 |
| Q9BU23 | LMF2 | | 65924000 | 50605000 | 38505000 | 53671000 | 56055000 | 0 | 51678000 | 36575333.33 | 0.491100383 | 0.707754428 | -0.498679225 |
| P02545 | LMNA | | 442060000 | 444120000 | 447550000 | 933370000 | 698300000 | 749260000 | 444576666.7 | 793643333.3 | 0.008113843 | 1.785166413 | 0.836058568 |
| P20700 | LMNB1 | | 335000000 | 301990000 | 564190000 | 695650000 | 491240000 | 372970000 | 400393333.3 | 519953333.3 | 0.393762398 | 1.29860637 | 0.376964192 |
| Q03252 | LMNB2 | | 62296000 | 79799000 | 63649000 | 65571000 | 0 | 93599000 | 68581333.33 | 53056666.67 | 0.612464536 | 0.773631309 | -0.370281913 |
| Q8NF37 | LPCAT1 | | 56822000 | 129900000 | 42576000 | 47608000 | 68909000 | 77306000 | 76432666.67 | 64607666.67 | 0.699057916 | 0.845288664 | -0.242483993 |
| Q96II8 | LRCH3 | | 69293000 | 37981000 | 76133000 | 0 | 0 | 0 | 61135666.67 | 0 | 0.00649175 | 0 | N/A |
| O75427 | LRCH4 | | 0 | 0 | 0 | 40209000 | 16552000 | 0 | 0 | 18920333.33 | 0.180204077 | N/A | N/A |
| Q07954 | LRP1 | | 487020000 | 465600000 | 493180000 | 657660000 | 551630000 | 500640000 | 481933333.3 | 569976666.7 | 0.134297878 | 1.182687785 | 0.242069271 |
| P30533 | LRPAP1 | | 607650000 | 633270000 | 658870000 | 802700000 | 504120000 | 337870000 | 633263333.3 | 548230000 | 0.567849488 | 0.865722001 | -0.208024271 |
| Q9BTT6 | LRRC1 | | 142230000 | 74086000 | 52355000 | 172030000 | 216960000 | 144550000 | 89557000 | 177846666.7 | 0.061847676 | 1.985848863 | 0.989755828 |
| Q9H9A6 | LRRC40 | | 0 | 0 | 0 | 57598000 | 47281000 | 64720000 | 0 | 56533000 | 0.000365986 | N/A | N/A |
| Q8N1G4 | LRRC47 | | 0 | 45486000 | 105820000 | 40500000 | 26849000 | 74606000 | 50435333.33 | 47318333.33 | 0.930913561 | 0.938198089 | -0.092035533 |
| Q96AG4 | LRRC59 | | 12521000000 | 12004000000 | 11810000000 | 8734100000 | 9338700000 | 10088000000 | 12111666667 | 9386933333 | 0.00361503 | 0.775032338 | -0.367671587 |
| Q6NSJ5 | LRRC8E | | 0 | 0 | 0 | 37029000 | 22792000 | 62951000 | 0 | 40924000 | 0.025321847 | N/A | N/A |
| Q9H089 | LSG1 | | 197300000 | 208390000 | 202830000 | 235460000 | 224570000 | 270690000 | 202840000 | 243573333.3 | 0.046293554 | 1.200815092 | 0.264014015 |
| Q9Y4Y9 | LSM5 | | 0 | 0 | 0 | 0 | 30618000 | 163510000 | 0 | 64709333.33 | 0.266759151 | N/A | N/A |
| Q86X29 | LSR | | 184050000 | 139810000 | 198020000 | 272760000 | 263240000 | 228060000 | 173960000 | 254686666.7 | 0.022019863 | 1.464053039 | 0.54996782 |
| P48449 | LSS | | 358580000 | 401700000 | 330190000 | 309680000 | 368840000 | 393070000 | 363490000 | 357196666.7 | 0.855168617 | 0.982686365 | -0.025197057 |
| O94822 | LTN1 | | 0 | 0 | 0 | 33899000 | 59357000 | 0 | 0 | 31085333.33 | 0.144871965 | N/A | N/A |
| Q9NQ29 | LUC7L | | 0 | 0 | 0 | 0 | 19455000 | 18160000 | 0 | 12538333.33 | 0.116587752 | N/A | N/A |
| Q9Y383 | LUC7L2 | | 48445000 | 49105000 | 46670000 | 64141000 | 53976000 | 46654000 | 48073333.33 | 54923666.67 | 0.252073929 | 1.142497573 | 0.192191101 |
| Q9HD34 | LYRM4 | | 194730000 | 222850000 | 0 | 121120000 | 113700000 | 0 | 139193333.3 | 78273333.33 | 0.490233969 | 0.562335361 | -0.830497325 |
| Q5U5X0 | LYRM7 | | 356940000 | 435300000 | 383560000 | 550350000 | 480030000 | 212400000 | 391933333.3 | 414260000 | 0.842728845 | 1.05696547 | 0.079928247 |
| P20645 | M6PR | | 115910000 | 88076000 | 71309000 | 192080000 | 99320000 | 132890000 | 91765000 | 141430000 | 0.173994966 | 1.541219419 | 0.624072269 |
| Q9UPN3 | MACF1 | | 898860000 | 896760000 | 858390000 | 644870000 | 642420000 | 504210000 | 884670000 | 597166666.7 | 0.004000089 | 0.675016296 | -0.567005763 |
| O75367 | MACROH2A1 | | 139810000 | 0 | 125410000 | 119040000 | 132480000 | 0 | 88406666.67 | 83840000 | 0.944086723 | 0.94834477 | -0.076516449 |
| P43358 | MAGEA4 | | 278440000 | 251970000 | 280490000 | 672030000 | 786570000 | 833670000 | 270300000 | 764090000 | 0.000539766 | 2.82682205 | 1.499181067 |
| O15479 | MAGEB2 | | 96078000 | 94646000 | 104640000 | 0 | 0 | 0 | 98454666.67 | 0 | 6.01198E-06 | 0 | N/A |
| Q9UNF1 | MAGED2 | | 145150000 | 164570000 | 198480000 | 339270000 | 325080000 | 246250000 | 169400000 | 303533333.3 | 0.015078435 | 1.791814246 | 0.841421084 |
| Q9H0U3 | MAGT1 | | 41419000 | 83291000 | 40846000 | 0 | 0 | 18876000 | 55185333.33 | 6292000 | 0.033686313 | 0.114015801 | -3.132694315 |
| P33908 | MAN1A1 | | 0 | 64686000 | 0 | 8133800 | 5659900 | 0 | 21562000 | 4597900 | 0.47796652 | 0.213240887 | -2.229444008 |
| Q9UKM7 | MAN1B1 | | 0 | 0 | 0 | 100150000 | 77819000 | 73359000 | 0 | 83776000 | 0.000538983 | N/A | N/A |
| Q16706 | MAN2A1 | | 272640000 | 301310000 | 216600000 | 126460000 | 196040000 | 94522000 | 263516666.7 | 139007333.3 | 0.033002001 | 0.527508696 | -0.922733217 |
| O00754 | MAN2B1 | | 138020000 | 0 | 80376000 | 168790000 | 153370000 | 180850000 | 72798666.67 | 167670000 | 0.080694198 | 2.303201524 | 1.203640649 |
| Q9NQG1 | MANBAL | | 0 | 0 | 57286000 | 41895000 | 0 | 0 | 19095333.33 | 13965000 | 0.838926652 | 0.731330517 | -0.451404531 |
| P55145 | MANF | | 320600000 | 383930000 | 641580000 | 564670000 | 726350000 | 382700000 | 448703333.3 | 557906666.7 | 0.477769313 | 1.243375356 | 0.314261889 |
| Q02750 | MAP2K1 | | 0 | 0 | 0 | 106480000 | 80951000 | 0 | 0 | 62477000 | 0.123442993 | N/A | N/A |
| P27816 | MAP4 | | 333800000 | 132080000 | 235790000 | 281910000 | 379250000 | 697540000 | 233890000 | 452900000 | 0.188612651 | 1.93638035 | 0.953362359 |
| O95819 | MAP4K4 | | 0 | 0 | 0 | 65887000 | 63611000 | 0 | 0 | 43166000 | 0.116239374 | N/A | N/A |
| Q15691 | MAPRE1 | | 171330000 | 175240000 | 141030000 | 141090000 | 205610000 | 314270000 | 162533333.3 | 220323333.3 | 0.326046656 | 1.355557834 | 0.438886666 |
| P49006 | MARCKSL1 | | 847020000 | 807110000 | 648070000 | 532950000 | 403420000 | 297880000 | 767400000 | 411416666.7 | 0.017478383 | 0.536117627 | -0.899378526 |
| P56192 | MARS1 | | 79333000 | 0 | 89228000 | 154390000 | 209960000 | 238980000 | 56187000 | 201110000 | 0.018224754 | 3.579297702 | 1.839676543 |
| P43243 | MATR3 | | 271280000 | 277560000 | 240780000 | 306880000 | 343440000 | 365100000 | 263206666.7 | 338473333.3 | 0.02114954 | 1.285960335 | 0.362846144 |
| Q96N66 | MBOAT7 | | 231040000 | 205550000 | 252080000 | 347280000 | 244790000 | 258880000 | 229556666.7 | 283650000 | 0.194860886 | 1.235642615 | 0.305261533 |
| P43121 | MCAM | | 467090000 | 446160000 | 464980000 | 447590000 | 482480000 | 368600000 | 459410000 | 432890000 | 0.483005248 | 0.942273786 | -0.085781786 |
| P25205 | MCM3 | | 888360000 | 704960000 | 642700000 | 341370000 | 398630000 | 468320000 | 745340000 | 402773333.3 | 0.014155262 | 0.540388726 | -0.887930517 |
| P33992 | MCM5 | | 184920000 | 135150000 | 214830000 | 0 | 0 | 118680000 | 178300000 | 39560000 | 0.039011961 | 0.221873247 | -2.172192372 |
| P40925 | MDH1 | | 0 | 0 | 0 | 162070000 | 0 | 205110000 | 0 | 122393333.3 | 0.121552955 | N/A | N/A |
| Q6P9B6 | MEAK7 | | 27763000 | 0 | 72934000 | 69991000 | 23572000 | 81020000 | 33565666.67 | 58194333.33 | 0.422570805 | 1.733745792 | 0.793892381 |
| P08582 | MELTF | | 379570000 | 350650000 | 415930000 | 338050000 | 329240000 | 359260000 | 382050000 | 342183333.3 | 0.128865094 | 0.895650657 | -0.158991968 |
| Q14696 | MESD | | 242680000 | 0 | 278560000 | 357490000 | 199100000 | 359860000 | 173746666.7 | 305483333.3 | 0.267649705 | 1.758211189 | 0.814108371 |
| P53582 | METAP1 | | 0 | 0 | 0 | 100290000 | 122550000 | 107900000 | 0 | 110246666.7 | 7.22467E-05 | N/A | N/A |
| P50579 | METAP2 | | 93934000 | 0 | 79594000 | 125240000 | 104010000 | 91891000 | 57842666.67 | 107047000 | 0.185371153 | 1.850658107 | 0.888038394 |
| Q08431 | MFGE8 | | 0 | 0 | 33718000 | 35927000 | 78248000 | 0 | 11239333.33 | 38058333.33 | 0.34808678 | 3.386173557 | 1.75965592 |
| P26572 | MGAT1 | | 0 | 0 | 0 | 339680000 | 638960000 | 618020000 | 0 | 532220000 | 0.005267228 | N/A | N/A |

| | | | | | | | | | | | | | |
|---|---|---|---|---|---|---|---|---|---|---|---|---|---|
| Q10469 | MGAT2 | | 0 | 0 | 0 | 54821000 | 66035000 | 0 | 0 | 40285333.33 | 0.119529813 | N/A | N/A |
| O14880 | MGST3 | | 510630000 | 510790000 | 668890000 | 181160000 | 220040000 | 351300000 | 563436666.7 | 250833333.3 | 0.013237775 | 0.445184611 | -1.167524372 |
| Q5JRA6 | MIA3 | | 221640000 | 128350000 | 196290000 | 423480000 | 541240000 | 399380000 | 182093333.3 | 454700000 | 0.006297694 | 2.497071099 | 1.3202369 |
| O94851 | MICAL2 | | 0 | 39490000 | 0 | 0 | 38779000 | 0 | 13163333.33 | 12926333.33 | 0.990365632 | 0.981995442 | -0.026211767 |
| L0R8F8 | MIEF1 | | 47069000 | 52681000 | 70713000 | 64034000 | 49973000 | 0 | 56821000 | 38002333.33 | 0.414664061 | 0.668807894 | -0.58033622 |
| P14174 | MIF | | 0 | 0 | 0 | 401020000 | 0 | 584900000 | 0 | 328640000 | 0.129764646 | N/A | N/A |
| Q9UNW1 | MINPP1 | | 192700000 | 273600000 | 152500000 | 623800000 | 461820000 | 586430000 | 206266666.7 | 557350000 | 0.004398836 | 2.70208468 | 1.434072888 |
| Q8IVT2 | MISP | | 0 | 118880000 | 178920000 | 0 | 0 | 0 | 99266666.67 | 0 | 0.132026776 | 0 | N/A |
| Q14165 | MLEC | | 24192000 | 44355000 | 0 | 0 | 83098000 | 45273000 | 22849000 | 42790333.33 | 0.504541047 | 1.872744248 | 0.905153891 |
| Q8N4V1 | MMGT1 | | 139380000 | 160110000 | 174420000 | 0 | 161390000 | 141030000 | 157970000 | 100806666.7 | 0.331338921 | 0.638138043 | -0.648059551 |
| Q96T76 | MMS19 | | 33065000 | 34707000 | 0 | 0 | 57234000 | 22996000 | 22590666.67 | 26743333.33 | 0.846466019 | 1.183822227 | 0.24345245 |
| Q9H8S9 | MOB1A | | 0 | 0 | 20464000 | 0 | 0 | 53598000 | 6821333.333 | 17866000 | 0.594530994 | 2.619136044 | 1.389090998 |
| Q13724 | MOGS | | 1129400000 | 1127700000 | 1091400000 | 1261500000 | 1338700000 | 1303900000 | 1116166667 | 1301366667 | 0.001917619 | 1.165925041 | 0.221475039 |
| Q00013 | MPP1 | | 91856000 | 94973000 | 0 | 0 | 0 | 0 | 62276333.33 | 0 | 0.116227223 | 0 | N/A |
| Q14168 | MPP2 | | 0 | 22195000 | 217300000 | 0 | 30167000 | 61926000 | 79831666.67 | 30697666.67 | 0.528693905 | 0.384529948 | -1.378832131 |
| Q9NZW5 | MPP6 | | 332970000 | 307310000 | 309430000 | 236350000 | 0 | 190720000 | 316570000 | 142356666.7 | 0.075060268 | 0.449684641 | -1.153014487 |
| Q5T2T1 | MPP7 | | 59106000 | 68758000 | 51547000 | 0 | 0 | 0 | 59803666.67 | 0 | 0.000275774 | 0 | N/A |
| Q6WCQ1 | MPRIP | | 29283000 | 42084000 | 49032000 | 64686000 | 48396000 | 38820000 | 40133000 | 50634000 | 0.331504451 | 1.261654997 | 0.335317456 |
| P25325 | MPST | | 448490000 | 568170000 | 296290000 | 204500000 | 129740000 | 307040000 | 437650000 | 213760000 | 0.07577676 | 0.488426825 | -1.033785658 |
| O95297 | MPZL1 | | 0 | 0 | 0 | 143970000 | 97873000 | 0 | 0 | 80614333.33 | 0.130362484 | N/A | N/A |
| Q9UBG0 | MRC2 | | 332560000 | 386250000 | 431720000 | 460200000 | 361640000 | 369510000 | 383510000 | 397116666.7 | 0.765809499 | 1.035479301 | 0.050298714 |
| P52701 | MSH6 | | 0 | 0 | 0 | 0 | 56836000 | 140300000 | 0 | 65712000 | 0.182079398 | N/A | N/A |
| P26038 | MSN | | 3053100000 | 3212900000 | 3957000000 | 8208500000 | 7082000000 | 5133700000 | 3407666667 | 6808066667 | 0.022432175 | 1.997867554 | 0.998460944 |
| Q86UE4 | MTDH | | 422800000 | 541180000 | 444650000 | 428360000 | 317990000 | 265420000 | 469543333.3 | 337256666.7 | 0.093040365 | 0.718265265 | -0.477411346 |
| P11586 | MTHFD1 | | 1863800000 | 1814300000 | 1732600000 | 3682500000 | 3475100000 | 3062400000 | 1803566667 | 3406666667 | 0.001000731 | 1.888849872 | 0.917508039 |
| P42285 | MTREX | | 43040000 | 24967000 | 44387000 | 0 | 0 | 47517000 | 37464666.67 | 15839000 | 0.273018241 | 0.422771678 | -1.242049362 |
| Q14764 | MVP | | 350010000 | 341810000 | 311040000 | 285270000 | 351200000 | 389350000 | 334286666.7 | 341940000 | 0.826087345 | 1.022894522 | 0.032657386 |
| Q96S97 | MYADM | | 72523000 | 0 | 0 | 0 | 0 | 106610000 | 24174333.33 | 35536666.67 | 0.804561 | 1.470016409 | 0.555832259 |
| Q969H8 | MYDGF | | 1144200000 | 1028400000 | 1054800000 | 1917000000 | 1397300000 | 1176400000 | 1075800000 | 1496900000 | 0.131093887 | 1.391429634 | 0.476567952 |
| P35580 | MYH10 | | 1093100000 | 961680000 | 1235900000 | 1128900000 | 1112400000 | 926120000 | 1096893333 | 1055806667 | 0.708906694 | 0.962542696 | -0.055077558 |
| P35579 | MYH9 | | 17475000000 | 19843000000 | 17143000000 | 23593000000 | 21036000000 | 15050000000 | 18153666667 | 19893000000 | 0.550358011 | 1.095811682 | 0.131999888 |
| P05976 | MYL1 | | 0 | 209830000 | 272840000 | 0 | 163500000 | 0 | 160890000 | 54500000 | 0.342414358 | 0.338740755 | -1.561746524 |
| P19105 | MYL12A | | 1011400000 | 900190000 | 944340000 | 823190000 | 947580000 | 1388500000 | 951976666.7 | 1053090000 | 0.593410782 | 1.106214088 | 0.14563062 |
| P60660 | MYL6 | | 3373000000 | 3503900000 | 3701100000 | 2351600000 | 2684000000 | 2227900000 | 3526000000 | 2421166667 | 0.002661464 | 0.686660995 | -0.542330081 |
| Q92614 | MYO18A | | 41426000 | 43597000 | 85647000 | 127940000 | 73588000 | 88992000 | 56890000 | 96840000 | 0.138744814 | 1.702232378 | 0.767427998 |
| Q96H55 | MYO19 | | 0 | 0 | 88314000 | 0 | 36479000 | 0 | 29438000 | 12159666.67 | 0.616296534 | 0.413060217 | -1.275575978 |
| O43795 | MYO1B | | 873430000 | 1063300000 | 1208600000 | 1253500000 | 1360200000 | 1692000000 | 1048443333 | 1435233333 | 0.077612725 | 1.368918365 | 0.453036414 |
| O00159 | MYO1C | | 4283400000 | 3713300000 | 3341600000 | 2482500000 | 2523100000 | 3032600000 | 3779433333 | 2679400000 | 0.02795518 | 0.708942258 | -0.496259968 |
| O94832 | MYO1D | | 106140000 | 99867000 | 86156000 | 126080000 | 121350000 | 0 | 97387666.67 | 82476666.67 | 0.73861382 | 0.846890263 | -0.239753052 |
| Q9NZM1 | MYOF | | 509430000 | 433140000 | 446230000 | 460920000 | 485560000 | 578240000 | 462933333.3 | 508240000 | 0.34931366 | 1.097868664 | 0.134705477 |
| Q13765 | NACA | | 0 | 0 | 0 | 240720000 | 231280000 | 209090000 | 0 | 227030000 | 1.72483E-05 | N/A | N/A |
| P17050 | NAGA | | 0 | 0 | 56971000 | 0 | 34116000 | 0 | 18990333.33 | 11372000 | 0.748045657 | 0.598830984 | -0.739779225 |
| P43490 | NAMPT | | 134390000 | 135160000 | 100830000 | 138880000 | 131770000 | 193350000 | 123460000 | 154666666.7 | 0.237790847 | 1.252767428 | 0.325118608 |
| P55209 | NAP1L1 | | 611060000 | 656470000 | 513240000 | 691660000 | 766190000 | 1215400000 | 593590000 | 891083333.3 | 0.153067217 | 1.501176457 | 0.58609357 |
| Q99733 | NAP1L4 | | 0 | 0 | 0 | 41951000 | 27203000 | 24889000 | 0 | 31347666.67 | 0.004215885 | N/A | N/A |
| P54920 | NAPA | | 177590000 | 235290000 | 271740000 | 242550000 | 309150000 | 208460000 | 228206666.7 | 253386666.7 | 0.56612306 | 1.110338582 | 0.150999673 |
| Q99747 | NAPG | | 163270000 | 122060000 | 0 | 125460000 | 151840000 | 0 | 95110000 | 92433333.33 | 0.970400892 | 0.971857148 | -0.041183826 |
| Q69YL0 | NCBP2AS2 | | 215000000 | 212130000 | 241100000 | 345680000 | 209460000 | 360010000 | 222743333.3 | 305050000 | 0.167298849 | 1.369513491 | 0.453663478 |
| Q6PIU2 | NCEH1 | | 153100000 | 181480000 | 196600000 | 94688000 | 11466000 | 124560000 | 177060000 | 76904666.67 | 0.050339062 | 0.434342407 | -1.203095276 |
| Q9Y2A7 | NCKAP1 | | 124370000 | 0 | 40338000 | 0 | 35948000 | 119780000 | 54902666.67 | 51909333.33 | 0.956014648 | 0.945479272 | -0.080882264 |
| P19338 | NCL | | 2742300000 | 2778300000 | 2917100000 | 2539400000 | 2567100000 | 2765700000 | 2812566667 | 2624066667 | 0.101538687 | 0.932979366 | -0.10008292 |
| Q969V3 | NCLN | | 367360000 | 327960000 | 310250000 | 434020000 | 390860000 | 467530000 | 335190000 | 430803333.3 | 0.026549536 | 1.285251151 | 0.362050304 |
| P62166 | NCS1 | | 218420000 | 43646000 | 93573000 | 0 | 0 | 0 | 118546333.3 | 0 | 0.08470911 | 0 | N/A |
| Q9UMX5 | NENF | | 411400000 | 382170000 | 411590000 | 238920000 | 418990000 | 494660000 | 401720000 | 384190000 | 0.829939323 | 0.956362641 | -0.064370321 |
| P48681 | NES | | 962160000 | 1066400000 | 950320000 | 450660000 | 501070000 | 445120000 | 992960000 | 465616666.7 | 0.000209651 | 0.468917848 | -1.092592902 |
| Q99519 | NEU1 | | 70710000 | 89745000 | 0 | 353860000 | 154030000 | 311290000 | 53485000 | 273060000 | 0.030049712 | 5.105356642 | 2.352011746 |
| Q0ZGT2 | NEXN | | 77395000 | 77088000 | 48616000 | 69361000 | 79033000 | 49662000 | 67699666.67 | 66018666.67 | 0.902414991 | 0.975169745 | -0.036274729 |

| | | | | | | | | | | | | | |
|---|---|---|---|---|---|---|---|---|---|---|---|---|---|
| P35240 | NF2 | | 475590000 | 225910000 | 375310000 | 268180000 | 228060000 | 276010000 | 358936666.7 | 257416666.7 | 0.242245304 | 0.717164588 | -0.479623842 |
| Q6ZNB6 | NFXL1 | | 38626000 | 33264000 | 48004000 | 85700000 | 53489000 | 0 | 39964666.67 | 46396333.33 | 0.81230145 | 1.160933825 | 0.215285739 |
| Q8NEJ9 | NGDN | | 0 | 0 | 133490000 | 0 | 183320000 | 0 | 44496666.67 | 61106666.67 | 0.836835344 | 1.373286388 | 0.45763252 |
| Q5JS37 | NHLRC3 | | 0 | 0 | 0 | 51268000 | 54461000 | 0 | 0 | 35243000 | 0.116479157 | N/A | N/A |
| Q96TA1 | NIBAN2 | | 105650000 | 163980000 | 103160000 | 106440000 | 0 | 95921000 | 124263333.3 | 67453666.67 | 0.221473749 | 0.542828402 | -0.881431887 |
| P14543 | NID1 | | 0 | 0 | 0 | 117600000 | 72857000 | 107380000 | 0 | 99279000 | 0.001839673 | N/A | N/A |
| Q9BPW8 | NIPSNAP1 | | 797830000 | 1055300000 | 806820000 | 1242400000 | 1172700000 | 1490000000 | 886650000 | 1301700000 | 0.031604166 | 1.468110303 | 0.553960366 |
| O75323 | NIPSNAP2 | | 367470000 | 258500000 | 312490000 | 0 | 253550000 | 137940000 | 312820000 | 130496666.7 | 0.084221141 | 0.417162159 | -1.261319798 |
| P15531 | NME1 | | 0 | 0 | 0 | 42771000 | 34191000 | 75765000 | 0 | 50909000 | 0.015899833 | N/A | N/A |
| P22392 | NME2 | | 1182400000 | 1362300000 | 1673000000 | 1046700000 | 1072900000 | 743550000 | 1405900000 | 954383333.3 | 0.064262217 | 0.678841549 | -0.558853227 |
| Q13232 | NME3 | | 65943000 | 0 | 103040000 | 0 | 0 | 0 | 56327666.67 | 0 | 0.134921769 | 0 | N/A |
| P30419 | NMT1 | | 0 | 0 | 0 | 0 | 154550000 | 165930000 | 0 | 106826666.7 | 0.116617808 | N/A | N/A |
| P40261 | NNMT | | 24758000 | 68996000 | 18439000 | 114240000 | 155730000 | 99060000 | 37397666.67 | 123010000 | 0.021113456 | 3.289242644 | 1.717755438 |
| Q8NC60 | NOA1 | | 475690000 | 537590000 | 429960000 | 119420000 | 45413000 | 120800000 | 481080000 | 95211000 | 0.000640213 | 0.19791095 | -2.337076656 |
| Q14978 | NOLC1 | | 54078000 | 24397000 | 34467000 | 0 | 36060000 | 0 | 37647333.33 | 12020000 | 0.159401097 | 0.319278922 | -1.647110782 |
| Q5JPE7 | NOMO2 | | 366980000 | 439310000 | 468930000 | 500520000 | 455940000 | 466430000 | 425073333.3 | 474296666.7 | 0.21158129 | 1.115799627 | 0.158077974 |
| Q15233 | NONO | | 269780000 | 151050000 | 338960000 | 499830000 | 504070000 | 213370000 | 253263333.3 | 405756666.7 | 0.240573678 | 1.602113742 | 0.679976575 |
| P46087 | NOP2 | | 0 | 40229000 | 0 | 0 | 0 | 55871000 | 13409666.67 | 18623666.67 | 0.831408969 | 1.388823983 | 0.473863766 |
| Q8TAT6 | NPLOC4 | | 88367000 | 66780000 | 126370000 | 64942000 | 114030000 | 145830000 | 93839000 | 108267333.3 | 0.647928883 | 1.153756256 | 0.206338471 |
| P06748 | NPM1 | | 526980000 | 365710000 | 515730000 | 482280000 | 431520000 | 599690000 | 469473333.3 | 504496666.7 | 0.652059082 | 1.074601326 | 0.103801524 |
| O14786 | NRP1 | | 0 | 53020000 | 0 | 109230000 | 130160000 | 83153000 | 17673333.33 | 107514333.3 | 0.015746269 | 6.08342135 | 2.604882932 |
| Q15738 | NSDHL | | 244680000 | 301690000 | 247420000 | 222510000 | 314320000 | 398740000 | 264596666.7 | 311856666.7 | 0.432196826 | 1.178611472 | 0.237088213 |
| P46459 | NSF | | 752020000 | 867040000 | 806370000 | 876900000 | 851250000 | 1135500000 | 808476666.7 | 954550000 | 0.205283137 | 1.180677241 | 0.239614632 |
| Q08J23 | NSUN2 | | 0 | 0 | 0 | 121540000 | 83713000 | 0 | 0 | 68417666.67 | 0.129447345 | N/A | N/A |
| Q96CB9 | NSUN4 | | 0 | 0 | 71117000 | 74800000 | 101680000 | 0 | 23705666.67 | 58826666.67 | 0.413979499 | 2.481544497 | 1.311238324 |
| Q8TCD5 | NT5C | | 46508000 | 0 | 15531000 | 0 | 0 | 0 | 20679666.67 | 0 | 0.204891442 | 0 | N/A |
| Q9BSD7 | NTPCR | | 258760000 | 74341000 | 114270000 | 293940000 | 259020000 | 408060000 | 149123666.7 | 320340000 | 0.075759238 | 2.148149969 | 1.103094716 |
| Q02818 | NUCB1 | | 281470000 | 151140000 | 146750000 | 91814000 | 142570000 | 0 | 193120000 | 78128000 | 0.131436858 | 0.404556752 | -1.305585993 |
| Q9Y266 | NUDC | | 0 | 0 | 0 | 369740000 | 194230000 | 0 | 0 | 187990000 | 0.153122041 | N/A | N/A |
| O43809 | NUDT21 | | 0 | 0 | 0 | 63808000 | 54485000 | 67050000 | 0 | 61781000 | 8.08412E-05 | N/A | N/A |
| P49757 | NUMB | | 148840000 | 177310000 | 163740000 | 217090000 | 203190000 | 102300000 | 163296666.7 | 174193333.3 | 0.783546402 | 1.066729266 | 0.093194069 |
| P35658 | NUP214 | | 0 | 21887000 | 0 | 48255000 | 88379000 | 76579000 | 7295666.667 | 71071000 | 0.010281264 | 9.741536072 | 3.284149278 |
| Q9NX40 | OCIAD1 | | 707970000 | 864310000 | 834470000 | 800640000 | 719780000 | 432430000 | 802250000 | 650950000 | 0.281233559 | 0.811405422 | -0.301505151 |
| Q56VL3 | OCIAD2 | | 310550000 | 277370000 | 226100000 | 737640000 | 666920000 | 410020000 | 271340000 | 604860000 | 0.031282684 | 2.229158989 | 1.156499517 |
| Q16625 | OCLN | | 0 | 0 | 136180000 | 0 | 0 | 64697000 | 45393333.33 | 21565666.67 | 0.660130069 | 0.475084447 | -1.073744117 |
| Q5SWX8 | ODR4 | | 0 | 34467000 | 0 | 71730000 | 64967000 | 120720000 | 11489000 | 85805666.67 | 0.023999376 | 7.468506107 | 2.900819696 |
| Q9NTK5 | OLA1 | | 0 | 0 | 0 | 0 | 75815000 | 56267000 | 0 | 44027333.33 | 0.124755306 | N/A | N/A |
| Q9H6K4 | OPA3 | | 0 | 72989000 | 0 | 0 | 65389000 | 0 | 24329666.67 | 21796333.33 | 0.941906827 | 0.895874721 | -0.158631095 |
| Q13438 | OS9 | | 0 | 0 | 0 | 0 | 47172000 | 68698000 | 0 | 38623333.33 | 0.12965968 | N/A | N/A |
| P22059 | OSBP | | 0 | 0 | 0 | 74921000 | 67743000 | 105620000 | 0 | 82761333.33 | 0.002051308 | N/A | N/A |
| Q9BXB5 | OSBPL10 | | 127650000 | 0 | 0 | 54478000 | 82087000 | 0 | 42550000 | 45521666.67 | 0.954465588 | 1.069839405 | 0.097394248 |
| Q9BZF1 | OSBPL8 | | 62298000 | 78348000 | 247760000 | 81710000 | 113530000 | 84808000 | 129468666.7 | 93349333.33 | 0.580760305 | 0.721018728 | -0.471891362 |
| Q96SU4 | OSBPL9 | | 0 | 0 | 70612000 | 58427000 | 43411000 | 60253000 | 23537333.33 | 54030333.33 | 0.275046797 | 2.295516343 | 1.198818703 |
| Q32P28 | P3H1 | | 463290000 | 424930000 | 239220000 | 362350000 | 296090000 | 618330000 | 375813333.3 | 425590000 | 0.699955487 | 1.132450507 | 0.179448 |
| Q8IVL5 | P3H2 | | 0 | 103710000 | 180140000 | 0 | 0 | 0 | 94616666.67 | 0 | 0.144116301 | 0 | N/A |
| Q8IVL6 | P3H3 | | 1069600000 | 916320000 | 1060800000 | 866870000 | 896510000 | 879080000 | 1015573333 | 880820000 | 0.055686212 | 0.867313045 | -0.205375286 |
| Q92791 | P3H4 | | 0 | 0 | 73289000 | 86816000 | 96453000 | 77947000 | 24429666.67 | 87072000 | 0.066413475 | 3.564191079 | 1.833574683 |
| P13674 | P4HA1 | | 1317400000 | 1378000000 | 1668900000 | 2018000000 | 2401600000 | 2688300000 | 1454766667 | 2369300000 | 0.014711881 | 1.62864606 | 0.703673109 |
| O15460 | P4HA2 | | 1842400000 | 1955300000 | 2244300000 | 2952800000 | 3237900000 | 3209900000 | 2014000000 | 3133533333 | 0.001729963 | 1.555875538 | 0.637726657 |
| P07237 | P4HB | | 11222000000 | 13184000000 | 12684000000 | 24548000000 | 23774000000 | 19267000000 | 12363333333 | 22529666667 | 0.004356418 | 1.822297115 | 0.865758201 |
| Q9UQ80 | PA2G4 | | 548220000 | 623530000 | 616640000 | 1440400000 | 1245600000 | 891640000 | 596130000 | 1192546667 | 0.02134712 | 2.000480879 | 1.00034684 |
| P11940 | PABPC1 | | 2855200000 | 2804700000 | 2248400000 | 3303000000 | 3108700000 | 3932200000 | 2636100000 | 3447966667 | 0.061778943 | 1.307980223 | 0.387340727 |
| Q13310 | PABPC4 | | 275420000 | 305270000 | 252950000 | 235670000 | 269560000 | 218800000 | 277880000 | 241343333.3 | 0.160963974 | 0.868516386 | -0.203375027 |
| Q86U42 | PABPN1 | | 0 | 0 | 0 | 0 | 504100000 | 489080000 | 0 | 331060000 | 0.116207484 | N/A | N/A |
| Q9UNF0 | PACSIN2 | | 186270000 | 147120000 | 199560000 | 0 | 0 | 0 | 177650000 | 0 | 0.000351145 | 0 | N/A |
| Q9UKS6 | PACSIN3 | | 772380000 | 778700000 | 741270000 | 289770000 | 449510000 | 379420000 | 764116666.7 | 372900000 | 0.001199716 | 0.488014483 | -1.035004131 |
| Q15102 | PAFAH1B3 | | 0 | 74575000 | 0 | 0 | 108780000 | 137740000 | 24858333.33 | 82173333.33 | 0.304862405 | 3.305665437 | 1.724940719 |

| | | | | | | | | | | | | | |
|---|---|---|---|---|---|---|---|---|---|---|---|---|---|
| P22234 | PAICS | | 1021700000 | 1249100000 | 1376400000 | 1896500000 | 1810700000 | 1831000000 | 1215733333 | 1846066667 | 0.004141392 | 1.51847993 | 0.60262784 |
| Q13177 | PAK2 | | 0 | 0 | 71449000 | 97206000 | 94526000 | 66721000 | 23816333.33 | 86151000 | 0.072578845 | 3.61730745 | 1.854916224 |
| Q8WX93 | PALLD | | 189110000 | 150310000 | 122480000 | 0 | 88233000 | 71317000 | 153966666.7 | 53183333.33 | 0.03867217 | 0.345421087 | -1.533571938 |
| Q8IXS6 | PALM2 | | 19996000 | 20436000 | 0 | 0 | 0 | 0 | 13477333.33 | 0 | 0.116163626 | 0 | N/A |
| A6NDB9 | PALM3 | | 124420000 | 74273000 | 67248000 | 0 | 65780000 | 0 | 88647000 | 21926666.67 | 0.078356988 | 0.247348096 | -2.015385301 |
| O95340 | PAPSS2 | | 0 | 14728000 | 0 | 76159000 | 62243000 | 153300000 | 4909333.333 | 97234000 | 0.032520348 | 19.80594785 | 4.307861841 |
| Q8TEW0 | PARD3 | | 252540000 | 236390000 | 0 | 75453000 | 94682000 | 129150000 | 162976666.7 | 99761666.67 | 0.489298446 | 0.61212239 | -0.708107956 |
| Q99497 | PARK7 | | 76024000 | 78954000 | 84367000 | 76568000 | 69842000 | 0 | 79781666.67 | 48803333.33 | 0.276410704 | 0.611711128 | -0.709077573 |
| P09874 | PARP1 | | 881140000 | 653700000 | 752070000 | 703170000 | 771710000 | 588620000 | 762303333.3 | 687833333.3 | 0.429376165 | 0.902309229 | -0.148306152 |
| Q460N5 | PARP14 | | 0 | 0 | 24540000 | 0 | 23835000 | 38646000 | 8180000 | 20827000 | 0.414832338 | 2.54608802 | 1.348282295 |
| Q8NI35 | PATJ | | 71791000 | 73821000 | 0 | 0 | 0 | 0 | 48537333.33 | 0 | 0.116193822 | 0 | N/A |
| Q96AQ6 | PBXIP1 | | 128230000 | 206870000 | 47568000 | 122830000 | 261900000 | 265990000 | 127556000 | 216906666.7 | 0.246005123 | 1.70048188 | 0.765943633 |
| Q15365 | PCBP1 | | 2226400000 | 2398400000 | 2802400000 | 2788200000 | 2990700000 | 3624900000 | 2475733333 | 3134600000 | 0.096403744 | 1.266129901 | 0.340425428 |
| Q15366 | PCBP2 | | 1069900000 | 1007900000 | 1711500000 | 1231900000 | 1659500000 | 2164200000 | 1263100000 | 1685200000 | 0.295430696 | 1.334177816 | 0.415950959 |
| Q9Y5E4 | PCDHB5 | | 0 | 0 | 94717000 | 0 | 68650000 | 86974000 | 31572333.33 | 51874666.67 | 0.648005691 | 1.643041904 | 0.716369275 |
| P12004 | PCNA | | 73782000 | 79410000 | 65008000 | 0 | 109070000 | 78190000 | 72733333.33 | 62420000 | 0.768419827 | 0.858203483 | -0.220608339 |
| Q9UHG3 | PCYOX1 | | 82089000 | 122830000 | 0 | 0 | 0 | 0 | 68306333.33 | 0 | 0.131595561 | 0 | N/A |
| Q13442 | PDAP1 | | 0 | 0 | 0 | 111690000 | 116510000 | 166270000 | 0 | 131490000 | 0.001659588 | N/A | N/A |
| Q9BUL8 | PDCD10 | | 109230000 | 0 | 133890000 | 173810000 | 184390000 | 0 | 81040000 | 119400000 | 0.625043725 | 1.473346496 | 0.559096757 |
| O14737 | PDCD5 | | 0 | 0 | 59457000 | 0 | 96770000 | 0 | 19819000 | 32256666.67 | 0.758991687 | 1.627562776 | 0.70271319 |
| O75340 | PDCD6 | | 0 | 0 | 0 | 42930000 | 230890000 | 278680000 | 0 | 184166666.7 | 0.062669151 | N/A | N/A |
| Q8WUM4 | PDCD6IP | | 538200000 | 406950000 | 435380000 | 962920000 | 810060000 | 646100000 | 460176666.7 | 806360000 | 0.025600989 | 1.752283543 | 0.809236241 |
| P30101 | PDIA3 | | 21597000000 | 18101000000 | 24087000000 | 27458000000 | 25500000000 | 25220000000 | 21261666667 | 26059333333 | 0.062578154 | 1.225648663 | 0.293545485 |
| P13667 | PDIA4 | | 2305400000 | 3081900000 | 3348300000 | 3087400000 | 3029200000 | 1868500000 | 2911866667 | 2661700000 | 0.646571177 | 0.914087183 | -0.129596322 |
| Q14554 | PDIA5 | | 205920000 | 156590000 | 120360000 | 237750000 | 283640000 | 278990000 | 160956666.7 | 266793333.3 | 0.021218664 | 1.65754758 | 0.729050284 |
| Q15084 | PDIA6 | | 19962000000 | 17826000000 | 17259000000 | 22397000000 | 21143000000 | 19699000000 | 18349000000 | 21079666667 | 0.073630958 | 1.148818283 | 0.200150614 |
| Q96JY6 | PDLIM2 | | 0 | 100710000 | 96821000 | 125080000 | 179330000 | 191040000 | 65843666.67 | 165150000 | 0.062251773 | 2.508213901 | 1.326660386 |
| P50479 | PDLIM4 | | 303480000 | 258070000 | 274000000 | 925530000 | 672740000 | 260910000 | 278516666.7 | 619726666.7 | 0.153652883 | 2.225097241 | 1.153868386 |
| Q9NR12 | PDLIM7 | | 487530000 | 564530000 | 676450000 | 248610000 | 225380000 | 359100000 | 576170000 | 277696666.7 | 0.012164755 | 0.48197002 | -1.052984684 |
| Q15121 | PEA15 | | 0 | 0 | 75077000 | 104670000 | 0 | 150450000 | 25025666.67 | 85040000 | 0.305174642 | 3.398111272 | 1.764733095 |
| Q8IZL8 | PELP1 | | 0 | 0 | 0 | 0 | 117430000 | 162720000 | 0 | 93383333.33 | 0.126406793 | N/A | N/A |
| O96011 | PEX11B | | 0 | 0 | 0 | 241280000 | 248420000 | 238290000 | 0 | 242663333.3 | 1.40944E-07 | N/A | N/A |
| O75381 | PEX14 | | 160080000 | 0 | 163470000 | 379140000 | 308510000 | 375440000 | 107850000 | 354363333.3 | 0.013633341 | 3.285705455 | 1.716203157 |
| Q9UHV9 | PFDN2 | | 139860000 | 0 | 99894000 | 186480000 | 0 | 0 | 79918000 | 62160000 | 0.823984758 | 0.777797242 | -0.362533975 |
| P17858 | PFKL | | 0 | 55150000 | 0 | 0 | 0 | 95021000 | 18383333.33 | 31673666.67 | 0.73503902 | 1.722955576 | 0.784885504 |
| P18669 | PGAM1 | | 354400000 | 338780000 | 434540000 | 684410000 | 738980000 | 574130000 | 375906666.7 | 665840000 | 0.006977813 | 1.771290746 | 0.824801041 |
| P52209 | PGD | | 950410000 | 662840000 | 880310000 | 863240000 | 1125600000 | 1117400000 | 831186666.7 | 1035413333 | 0.169749614 | 1.245704936 | 0.316962385 |
| P00558 | PGK1 | | 1540500000 | 1324300000 | 1414100000 | 933200000 | 1008400000 | 1157700000 | 1426300000 | 1033100000 | 0.012448908 | 0.724321671 | -0.465297554 |
| O00264 | PGRMC1 | | 1386600000 | 1515300000 | 1589100000 | 1198400000 | 1126700000 | 1044600000 | 1497000000 | 1123233333 | 0.007224451 | 0.750322868 | -0.414416566 |
| O15173 | PGRMC2 | | 429480000 | 453370000 | 427490000 | 261750000 | 330510000 | 419810000 | 436780000 | 337356666.7 | 0.099324199 | 0.772372056 | -0.372632125 |
| P35232 | PHB | | 9313400000 | 8992100000 | 7043800000 | 10256000000 | 10447000000 | 15956000000 | 8449766667 | 12219666667 | 0.132372349 | 1.446154332 | 0.532221523 |
| Q99623 | PHB2 | | 2655500000 | 2478100000 | 2679400000 | 4452500000 | 4410300000 | 4083000000 | 2604333333 | 4315266667 | 0.000210017 | 1.656956355 | 0.728535602 |
| O43175 | PHGDH | | 2512100000 | 2725500000 | 2727400000 | 2053400000 | 2083700000 | 1854800000 | 2655000000 | 1997300000 | 0.002900566 | 0.752278719 | -0.410660815 |
| Q9BTU6 | PI4K2A | | 488860000 | 427240000 | 410630000 | 513100000 | 479560000 | 528890000 | 442243333.3 | 507183333.3 | 0.080384234 | 1.146842236 | 0.197666943 |
| P42356 | PI4KA | | 0 | 43195000 | 0 | 0 | 58792000 | 0 | 14398333.33 | 19597333.33 | 0.841164871 | 1.361083459 | 0.444755533 |
| Q13492 | PICALM | | 321680000 | 442760000 | 381540000 | 287460000 | 295840000 | 213770000 | 381993333.3 | 265690000 | 0.055973961 | 0.695535699 | -0.523803531 |
| Q92643 | PIGK | | 105340000 | 0 | 0 | 53294000 | 69722000 | 0 | 35113333.33 | 41005333.33 | 0.892515637 | 1.167799506 | 0.223792606 |
| Q9H490 | PIGU | | 0 | 218450000 | 0 | 72612000 | 46901000 | 77803000 | 72816666.67 | 65772000 | 0.92819521 | 0.903254749 | -0.146795159 |
| P48739 | PITPNB | | 0 | 0 | 0 | 129980000 | 133890000 | 182620000 | 0 | 148830000 | 0.000924087 | N/A | N/A |
| P14618 | PKM | | 2048700000 | 2270000000 | 2177600000 | 5148200000 | 4849500000 | 5335300000 | 2165433333 | 5111000000 | 4.55631E-05 | 2.360266613 | 1.238949834 |
| Q16513 | PKN2 | | 257240000 | 229770000 | 207760000 | 165060000 | 261180000 | 172940000 | 231590000 | 199726666.7 | 0.401393372 | 0.8624149 | -0.213545991 |
| Q99959 | PKP2 | | 111070000 | 96622000 | 340280000 | 0 | 0 | 203440000 | 182657333.3 | 67813333.33 | 0.331655377 | 0.371259845 | -1.42949881 |
| Q01970 | PLCB3 | | 410620000 | 347230000 | 386010000 | 88918000 | 161050000 | 120520000 | 381286666.7 | 123496000 | 0.000758552 | 0.323892784 | -1.626411768 |
| P51178 | PLCD1 | | 30217000 | 28785000 | 0 | 0 | 33922000 | 0 | 19667333.33 | 11307333.33 | 0.606803716 | 0.574929663 | -0.798542627 |
| Q8N3E9 | PLCD3 | | 201080000 | 199230000 | 233900000 | 184860000 | 369340000 | 292160000 | 211403333.3 | 282120000 | 0.265424052 | 1.334510651 | 0.41631082 |
| Q8IV08 | PLD3 | | 0 | 204720000 | 172550000 | 315250000 | 374520000 | 402190000 | 125756666.7 | 363986666.7 | 0.025447822 | 2.89437273 | 1.533250721 |
| Q15149 | PLEC | | 49503000000 | 50244000000 | 48949000000 | 51986000000 | 53915000000 | 50177000000 | 49565333333 | 52026000000 | 0.097588051 | 1.049644913 | 0.069901358 |

| | | | | | | | | | | | | | |
|---|---|---|---|---|---|---|---|---|---|---|---|---|---|
| Q9HAU0 | PLEKHA5 | | 0 | 0 | 179380000 | 0 | 77221000 | 73729000 | 59793333.33 | 50316666.67 | 0.890933063 | 0.841509644 | -0.248948289 |
| Q99541 | PLIN2 | | 107210000 | 131140000 | 166330000 | 248060000 | 222790000 | 190020000 | 134893333.3 | 220290000 | 0.023694214 | 1.633068103 | 0.707584956 |
| O60664 | PLIN3 | | 762970000 | 678110000 | 933440000 | 1994500000 | 2164200000 | 2050900000 | 791506666.7 | 2069866667 | 0.000143564 | 2.615096946 | 1.386864431 |
| Q02809 | PLOD1 | | 2306200000 | 2998600000 | 2894600000 | 1428100000 | 1688500000 | 1953700000 | 2733133333 | 1690100000 | 0.016723817 | 0.618374515 | -0.693447232 |
| O00469 | PLOD2 | | 648400000 | 563820000 | 610160000 | 1408000000 | 1515400000 | 2034100000 | 607460000 | 1652500000 | 0.005832654 | 2.720343726 | 1.443788953 |
| O60568 | PLOD3 | | 1119300000 | 817570000 | 994980000 | 812070000 | 862810000 | 1011500000 | 977283333.3 | 895460000 | 0.483415752 | 0.916274707 | -0.126147899 |
| O94903 | PLPBP | | 0 | 67272000 | 0 | 0 | 90646000 | 0 | 22424000 | 30215333.33 | 0.846071693 | 1.347455108 | 0.430237208 |
| Q14651 | PLS1 | | 71779000 | 68166000 | 126750000 | 64543000 | 88894000 | 0 | 88898333.33 | 51145666.67 | 0.311255811 | 0.575327621 | -0.797544359 |
| P13797 | PLS3 | | 175870000 | 97346000 | 195490000 | 280310000 | 147900000 | 167360000 | 156235333.3 | 198523333.3 | 0.453769909 | 1.270668607 | 0.345587821 |
| Q9UIW2 | PLXNA1 | | 39499000 | 27247000 | 0 | 0 | 0 | 0 | 22248666.67 | 0 | 0.129342971 | 0 | N/A |
| O15031 | PLXNB2 | | 594680000 | 427840000 | 490050000 | 792110000 | 598200000 | 454320000 | 504190000 | 614876666.7 | 0.368509572 | 1.219533641 | 0.286329556 |
| Q8IY17 | PNPLA6 | | 61104000 | 0 | 0 | 30741000 | 6218900 | 66374000 | 20368000 | 34444633.33 | 0.627549643 | 1.691115148 | 0.757974896 |
| O00592 | PODXL | | 275960000 | 0 | 169340000 | 468410000 | 288210000 | 269600000 | 148433333.3 | 342073333.3 | 0.131442308 | 2.304558724 | 1.204490531 |
| Q9H488 | POFUT1 | | 213180000 | 189390000 | 128630000 | 163170000 | 195150000 | 0 | 177066666.7 | 119440000 | 0.428411962 | 0.674548193 | -0.568006575 |
| Q9Y2G5 | POFUT2 | | 0 | 0 | 0 | 62812000 | 50474000 | 0 | 0 | 37762000 | 0.120813096 | N/A | N/A |
| Q7Z4H8 | POGLUT3 | | 0 | 0 | 102110000 | 90100000 | 72630000 | 81067000 | 34036666.67 | 81265666.67 | 0.241806512 | 2.387591813 | 1.255556212 |
| Q9Y2S7 | POLDIP2 | | 285190000 | 354770000 | 359530000 | 358930000 | 346980000 | 219280000 | 333163333.3 | 308396666.7 | 0.651023297 | 0.925662088 | -0.11144246 |
| Q15165 | PON2 | | 372270000 | 335230000 | 457370000 | 521160000 | 422420000 | 459200000 | 388290000 | 467593333.3 | 0.161430925 | 1.204237383 | 0.268119808 |
| P16435 | POR | | 897100000 | 818220000 | 512920000 | 391420000 | 538380000 | 802260000 | 742746666.7 | 577353333.3 | 0.380196229 | 0.777322012 | -0.363415724 |
| Q6S8J3 | POTEE | | 439420000 | 0 | 359530000 | 393650000 | 498430000 | 728650000 | 266316666.7 | 540243333.3 | 0.177283981 | 2.028575005 | 1.020466646 |
| Q15181 | PPA1 | | 0 | 0 | 0 | 95460000 | 113770000 | 0 | 0 | 69743333.33 | 0.119153739 | N/A | N/A |
| Q86W92 | PPFIBP1 | | 96522000 | 118260000 | 130470000 | 171420000 | 81146000 | 0 | 115084000 | 84188666.67 | 0.573696149 | 0.731541019 | -0.450989333 |
| P62937 | PPIA | | 4855800000 | 3878300000 | 3694200000 | 3586000000 | 3930000000 | 4341500000 | 4142766667 | 3952500000 | 0.67505475 | 0.95407256 | -0.067829103 |
| P23284 | PPIB | | 3495600000 | 3771700000 | 3296800000 | 2102200000 | 2247900000 | 2256500000 | 3521366667 | 2202200000 | 0.000842501 | 0.625382191 | -0.67718996 |
| P45877 | PPIC | | 281360000 | 213110000 | 100610000 | 203880000 | 244340000 | 212660000 | 198360000 | 220293333.3 | 0.705967567 | 1.110573368 | 0.151304705 |
| Q9UNP9 | PPIE | | 278530000 | 293260000 | 182290000 | 282690000 | 0 | 179960000 | 251360000 | 154216666.7 | 0.339465164 | 0.613529069 | -0.704796396 |
| P50336 | PPOX | | 60358000 | 0 | 40174000 | 23597000 | 0 | 0 | 33510666.67 | 7865666.667 | 0.256834396 | 0.234721283 | -2.090979434 |
| P62136 | PPP1CA | | 201870000 | 712410000 | 110790000 | 158030000 | 196840000 | 842010000 | 341690000 | 398960000 | 0.853206572 | 1.167608066 | 0.223556082 |
| P36873 | PPP1CC | | 281560000 | 223060000 | 244330000 | 0 | 0 | 0 | 249650000 | 0 | 0.000127914 | 0 | N/A |
| O14974 | PPP1R12A | | 181580000 | 81714000 | 94745000 | 80310000 | 25329000 | 67519000 | 119346333.3 | 57719333.33 | 0.157334023 | 0.483628878 | -1.048027702 |
| P67775 | PPP2CA | | 168200000 | 149610000 | 138160000 | 181840000 | 183570000 | 183710000 | 151990000 | 183040000 | 0.024035362 | 1.204289756 | 0.268182551 |
| P30153 | PPP2R1A | | 697280000 | 520400000 | 783700000 | 678330000 | 580500000 | 772740000 | 667126666.7 | 677190000 | 0.920996847 | 1.015084592 | 0.021599959 |
| Q16537 | PPP2R5E | | 42633000 | 0 | 0 | 68637000 | 0 | 0 | 14211000 | 22879000 | 0.763696363 | 1.609950039 | 0.687015918 |
| P50897 | PPT1 | | 1902800000 | 1629700000 | 1760800000 | 1715500000 | 1556000000 | 1859600000 | 1764433333 | 1710366667 | 0.670409163 | 0.96935749 | -0.044899279 |
| O60831 | PRAF2 | | 647070000 | 265400000 | 343410000 | 341240000 | 252260000 | 436500000 | 418626666.7 | 343333333.3 | 0.587981654 | 0.820142052 | -0.286054283 |
| P42785 | PRCP | | 79112000 | 247390000 | 0 | 179690000 | 74056000 | 160400000 | 108834000 | 138048666.7 | 0.732985901 | 1.268433271 | 0.343047625 |
| Q06830 | PRDX1 | | 2553000000 | 2075800000 | 2273900000 | 5330400000 | 5323100000 | 5743300000 | 2300900000 | 5465600000 | 8.61818E-05 | 2.375418315 | 1.248181596 |
| P32119 | PRDX2 | | 0 | 0 | 35138000 | 68950000 | 63428000 | 0 | 11712666.67 | 44126000 | 0.265017174 | 3.767374353 | 1.913559397 |
| Q13162 | PRDX4 | | 151370000 | 207470000 | 263130000 | 269860000 | 257110000 | 196010000 | 207323333.3 | 240993333.3 | 0.442024624 | 1.162403331 | 0.217110743 |
| P30041 | PRDX6 | | 0 | 0 | 0 | 0 | 216350000 | 190790000 | 0 | 135713333.3 | 0.117681819 | N/A | N/A |
| Q9HCU5 | PREB | | 576710000 | 514190000 | 467380000 | 549530000 | 620040000 | 554190000 | 519426666.7 | 574586666.7 | 0.230192214 | 1.106194009 | 0.145604433 |
| P49643 | PRIM2 | | 26101000 | 35760000 | 0 | 0 | 0 | 0 | 20620333.33 | 0 | 0.125722427 | 0 | N/A |
| P17612 | PRKACA | | 0 | 0 | 51706000 | 0 | 0 | 33072000 | 17235333.33 | 11024000 | 0.776573625 | 0.639616292 | -0.644721408 |
| P10644 | PRKAR1A | | 0 | 0 | 0 | 40740000 | 46999000 | 0 | 0 | 29246333.33 | 0.118136737 | N/A | N/A |
| P13861 | PRKAR2A | | 248920000 | 199080000 | 355140000 | 167630000 | 152810000 | 324820000 | 267713333.3 | 215086666.7 | 0.50388034 | 0.80342157 | -0.315770899 |
| P14314 | PRKCSH | | 2110300000 | 2254000000 | 2299800000 | 2822500000 | 2081600000 | 1171300000 | 2221366667 | 2025133333 | 0.704129075 | 0.911660989 | -0.133430652 |
| P78527 | PRKDC | | 599350000 | 497130000 | 546310000 | 205880000 | 272640000 | 84368000 | 547596666.7 | 187629333.3 | 0.004512606 | 0.342641482 | -1.545228273 |
| O75569 | PRKRA | | 195580000 | 109440000 | 152330000 | 197100000 | 104540000 | 253600000 | 152450000 | 185080000 | 0.550077727 | 1.214037389 | 0.279812854 |
| Q9UNN8 | PROCR | | 88635000 | 91797000 | 102770000 | 90262000 | 91835000 | 94133000 | 94400666.67 | 92076666.67 | 0.627443309 | 0.97538153 | -0.035961442 |
| Q9UMS4 | PRPF19 | | 234390000 | 246990000 | 324770000 | 295520000 | 357380000 | 287270000 | 268716666.7 | 313390000 | 0.281191619 | 1.166246976 | 0.221873341 |
| Q6P2Q9 | PRPF8 | | 0 | 82592000 | 54608000 | 47052000 | 0 | 0 | 45733333.33 | 15684000 | 0.356887483 | 0.342944606 | -1.543952529 |
| P11908 | PRPS2 | | 108460000 | 0 | 0 | 118330000 | 137340000 | 0 | 36153333.33 | 85223333.33 | 0.431504375 | 2.357274571 | 1.237119811 |
| O60256 | PRPSAP2 | | 0 | 85796000 | 22864000 | 44475000 | 54213000 | 54564000 | 36220000 | 51084000 | 0.596269669 | 1.410381005 | 0.49608495 |
| Q96HE9 | PRR11 | | 0 | 149570000 | 159490000 | 0 | 0 | 0 | 103020000 | 0 | 0.11652614 | 0 | N/A |
| Q5THK1 | PRR14L | | 0 | 0 | 1791000000 | 1034200000 | 959050000 | 709340000 | 597000000 | 900863333.3 | 0.641889535 | 1.508983808 | 0.593577325 |
| P07602 | PSAP | | 439620000 | 363560000 | 289670000 | 548880000 | 342680000 | 559310000 | 364283333.3 | 483623333.3 | 0.222759192 | 1.327602141 | 0.408822861 |
| P28066 | PSMA5 | | 0 | 0 | 0 | 147020000 | 152520000 | 50681000 | 0 | 116740333.3 | 0.024223315 | N/A | N/A |

| | | | | | | | | | | | | | |
|---|---|---|---|---|---|---|---|---|---|---|---|---|---|
| P35998 | PSMC2 | | 0 | 0 | 0 | 0 | 165930000 | 158580000 | 0 | 108170000 | 0.116320529 | N/A | N/A |
| P17980 | PSMC3 | | 141360000 | 86047000 | 47138000 | 70897000 | 91870000 | 104560000 | 91515000 | 89109000 | 0.93796155 | 0.973709228 | -0.03843708 |
| P43686 | PSMC4 | | 127270000 | 80201000 | 63180000 | 227730000 | 114870000 | 62979000 | 90217000 | 135193000 | 0.438086133 | 1.498531319 | 0.583549236 |
| P62333 | PSMC6 | | 0 | 156610000 | 139140000 | 223540000 | 275920000 | 194880000 | 98583333.33 | 231446666.7 | 0.0728821 | 2.34772612 | 1.231264117 |
| Q99460 | PSMD1 | | 0 | 0 | 0 | 0 | 38793000 | 23934000 | 0 | 20909000 | 0.137939526 | N/A | N/A |
| O00231 | PSMD11 | | 296680000 | 327900000 | 224100000 | 213230000 | 361670000 | 425250000 | 282893333.3 | 333383333.3 | 0.510293374 | 1.178477165 | 0.236923804 |
| Q13200 | PSMD2 | | 0 | 130030000 | 137970000 | 291060000 | 264090000 | 211270000 | 89333333.33 | 255473333.3 | 0.030203925 | 2.859776119 | 1.515902209 |
| O43242 | PSMD3 | | 0 | 0 | 0 | 88303000 | 103210000 | 0 | 0 | 63837666.67 | 0.118520866 | N/A | N/A |
| P51665 | PSMD7 | | 7407900 | 6125100 | 0 | 0 | 0 | 49357000 | 4511000 | 16452333.33 | 0.511955069 | 3.647158797 | 1.866773016 |
| Q06323 | PSME1 | | 117970000 | 148530000 | 0 | 143310000 | 119290000 | 69666000 | 88833333.33 | 110755333.3 | 0.684918193 | 1.246776735 | 0.31820314 |
| Q9UL46 | PSME2 | | 295660000 | 317290000 | 204710000 | 98173000 | 110230000 | 344430000 | 272553333.3 | 184277666.7 | 0.368923459 | 0.67611599 | -0.564657329 |
| P26599 | PTBP1 | | 298080000 | 324040000 | 356760000 | 244800000 | 272400000 | 462520000 | 326293333.3 | 326573333.3 | 0.997021831 | 1.000858124 | 0.00123748 |
| Q9H7Z7 | PTGES2 | | 543450000 | 373680000 | 443370000 | 308750000 | 352760000 | 373900000 | 453500000 | 345136666.7 | 0.109759628 | 0.761051084 | -0.3939348 |
| Q9P2B2 | PTGFRN | | 159920000 | 154550000 | 0 | 0 | 310580000 | 159480000 | 104823333.3 | 156686666.7 | 0.643785012 | 1.494768976 | 0.579922526 |
| Q13308 | PTK7 | | 1134700000 | 1067600000 | 978250000 | 954900000 | 1007400000 | 1198800000 | 1060183333 | 1053700000 | 0.944091382 | 0.993884705 | -0.008849592 |
| P18031 | PTPN1 | | 215860000 | 281150000 | 206710000 | 326410000 | 366400000 | 321700000 | 234573333.3 | 338170000 | 0.019405734 | 1.44163872 | 0.527709665 |
| P10586 | PTPRF | | 499230000 | 449620000 | 414720000 | 306640000 | 317230000 | 449610000 | 454523333.3 | 357826666.7 | 0.137153338 | 0.78725698 | -0.345093452 |
| P53801 | PTTG1IP | | 0 | 0 | 0 | 127130000 | 136550000 | 183410000 | 0 | 149030000 | 0.001021269 | N/A | N/A |
| Q9Y606 | PUS1 | | 61648000 | 108650000 | 78758000 | 0 | 103030000 | 0 | 83018666.67 | 34343333.33 | 0.25853032 | 0.413682063 | -1.273405689 |
| Q92626 | PXDN | | 26915000 | 24012000 | 26424000 | 33209000 | 33208000 | 0 | 25783666.67 | 22139000 | 0.759238039 | 0.858644361 | -0.219867384 |
| Q96C36 | PYCR2 | | 981110000 | 848540000 | 679020000 | 555900000 | 528650000 | 351020000 | 836223333.3 | 478523333.3 | 0.030011044 | 0.57224346 | -0.805299026 |
| P47897 | QARS1 | | 899070000 | 679860000 | 522150000 | 917730000 | 580850000 | 1095500000 | 700360000 | 864693333.3 | 0.427619352 | 1.234641232 | 0.304091878 |
| P09417 | QDPR | | 48831000 | 58417000 | 0 | 78705000 | 66492000 | 0 | 35749333.33 | 48399000 | 0.698846663 | 1.353843428 | 0.437060901 |
| Q6ZRP7 | QSOX2 | | 96393000 | 417010000 | 423240000 | 101770000 | 175790000 | 411370000 | 312214333.3 | 229643333.3 | 0.5938318 | 0.73553104 | -0.443141869 |
| P61026 | RAB10 | | 352350000 | 435690000 | 192960000 | 279290000 | 270370000 | 241090000 | 327000000 | 263583333.3 | 0.428974625 | 0.80606524 | -0.311031486 |
| Q15907 | RAB11B | | 2381200000 | 2541600000 | 3093300000 | 1966000000 | 2186500000 | 1667700000 | 2672033333 | 1940066667 | 0.049596386 | 0.726063797 | -0.461831777 |
| Q6WKZ4 | RAB11FIP1 | | 0 | 0 | 154150000 | 0 | 0 | 96118000 | 51383333.33 | 32039333.33 | 0.765372395 | 0.623535517 | -0.681456355 |
| Q9BXF6 | RAB11FIP5 | | 0 | 0 | 0 | 50617000 | 58925000 | 0 | 0 | 36514000 | 0.118399465 | N/A | N/A |
| P51153 | RAB13 | | 0 | 0 | 16349000 | 0 | 68290000 | 43344000 | 5449666.667 | 37211333.33 | 0.199401284 | 6.82818521 | 2.771502191 |
| P61106 | RAB14 | | 621850000 | 480100000 | 469310000 | 287130000 | 288390000 | 386970000 | 523753333.3 | 320830000 | 0.026644568 | 0.612559347 | -0.70707847 |
| Q9NP72 | RAB18 | | 362980000 | 304070000 | 447800000 | 399810000 | 476410000 | 610550000 | 371616666.7 | 495590000 | 0.170912003 | 1.333605418 | 0.41533187 |
| P62820 | RAB1A | | 2047200000 | 1838400000 | 2191200000 | 1787900000 | 2222100000 | 1766700000 | 2025600000 | 1925566667 | 0.60858805 | 0.950615456 | -0.073066237 |
| Q9H0U4 | RAB1B | | 356050000 | 408850000 | 360980000 | 198470000 | 353710000 | 516840000 | 375293333.3 | 356340000 | 0.849167623 | 0.949497282 | -0.074764224 |
| Q9UL25 | RAB21 | | 100330000 | 206880000 | 161460000 | 124630000 | 128500000 | 156960000 | 156223333.3 | 136696666.7 | 0.580444769 | 0.875008001 | -0.192631885 |
| Q9ULC3 | RAB23 | | 102700000 | 133990000 | 132460000 | 0 | 61347000 | 0 | 123050000 | 20449000 | 0.010895709 | 0.166184478 | -2.589142459 |
| P61019 | RAB2A | | 1385700000 | 1391800000 | 1407300000 | 1399100000 | 1831400000 | 2113100000 | 1394933333 | 1781200000 | 0.136493786 | 1.276906901 | 0.352653342 |
| Q13636 | RAB31 | | 0 | 0 | 79364000 | 0 | 0 | 82932000 | 26454666.67 | 27644000 | 0.976692217 | 1.044957411 | 0.063444145 |
| Q9BZG1 | RAB34 | | 237780000 | 277140000 | 345980000 | 176960000 | 201460000 | 185700000 | 286966666.7 | 188040000 | 0.037980073 | 0.655267743 | -0.609843581 |
| Q15286 | RAB35 | | 114640000 | 79841000 | 58985000 | 154720000 | 158550000 | 89084000 | 84488666.67 | 134118000 | 0.148559255 | 1.587408173 | 0.666673139 |
| P51148 | RAB5C | | 140600000 | 0 | 0 | 106010000 | 149290000 | 0 | 46866666.67 | 85100000 | 0.585349271 | 1.815789474 | 0.860596943 |
| P51149 | RAB7A | | 1672300000 | 1260000000 | 1737600000 | 1415900000 | 1417300000 | 1977700000 | 1556633333 | 1603633333 | 0.85395553 | 1.030193366 | 0.042915155 |
| P61006 | RAB8A | | 70916000 | 137280000 | 172820000 | 209820000 | 171210000 | 145960000 | 127005333.3 | 175663333.3 | 0.238650996 | 1.383117769 | 0.467924004 |
| Q92930 | RAB8B | | 68297000 | 45425000 | 62307000 | 185630000 | 130370000 | 69495000 | 58676333.33 | 128498333.3 | 0.110972039 | 2.189951656 | 1.130899022 |
| Q9UI14 | RABAC1 | | 0 | 0 | 0 | 105410000 | 134440000 | 125410000 | 0 | 121753333.3 | 0.000143024 | N/A | N/A |
| Q5HYI8 | RABL3 | | 0 | 0 | 0 | 39182000 | 44060000 | 0 | 0 | 27747333.33 | 0.117480617 | N/A | N/A |
| P63000 | RAC1 | | 904900000 | 1270100000 | 1937400000 | 2768400000 | 1481000000 | 999250000 | 1370800000 | 1749550000 | 0.567360488 | 1.276298512 | 0.351965799 |
| Q9H0H5 | RACGAP1 | | 109130000 | 166220000 | 0 | 141210000 | 122910000 | 103490000 | 91783333.33 | 122536666.7 | 0.571502549 | 1.335064463 | 0.416909404 |
| P63244 | RACK1 | | 471170000 | 323500000 | 325730000 | 749370000 | 807470000 | 836520000 | 373466666.7 | 797786666.7 | 0.001536874 | 2.136165655 | 1.095023529 |
| Q9P0K7 | RAI14 | | 83664000 | 88241000 | 76961000 | 140600000 | 99326000 | 0 | 82955333.33 | 79975333.33 | 0.946655601 | 0.964077054 | -0.052779637 |
| P11233 | RALA | | 892870000 | 753700000 | 878650000 | 515880000 | 696060000 | 677570000 | 841740000 | 629836666.7 | 0.042803837 | 0.748255598 | -0.418396929 |
| P11234 | RALB | | 0 | 0 | 145400000 | 268600000 | 0 | 265460000 | 48466666.67 | 178020000 | 0.270302508 | 3.67303989 | 1.876974564 |
| Q9UKM9 | RALY | | 87632000 | 88061000 | 91666000 | 126550000 | 119040000 | 124340000 | 89119666.67 | 123310000 | 0.000184353 | 1.383645211 | 0.46847406 |
| P62826 | RAN | | 91476000 | 80033000 | 90325000 | 293130000 | 199460000 | 130370000 | 87278000 | 207653333.3 | 0.063657945 | 2.379217367 | 1.250487083 |
| P43487 | RANBP1 | | 0 | 0 | 0 | 367790000 | 474590000 | 441930000 | 0 | 428103333.3 | 0.000171744 | N/A | N/A |
| P46060 | RANGAP1 | | 0 | 0 | 84583000 | 165540000 | 103720000 | 209140000 | 28194333.33 | 159466666.7 | 0.034320154 | 5.655982881 | 2.499777753 |
| P61224 | RAP1B | | 1224600000 | 1369100000 | 785270000 | 1234600000 | 1515600000 | 693920000 | 1126323333 | 1148040000 | 0.945450814 | 1.019281024 | 0.027551868 |
| P61225 | RAP2B | | 38160000 | 212510000 | 99499000 | 98488000 | 176120000 | 207430000 | 116723000 | 160679333.3 | 0.507503771 | 1.376586734 | 0.461095512 |

| | | | | | | | | | | | | | |
|---|---|---|---|---|---|---|---|---|---|---|---|---|---|
| P54136 | RARS1 | | 486350000 | 633490000 | 750260000 | 546010000 | 570320000 | 717120000 | 623366666.7 | 611150000 | 0.902040688 | 0.980402118 | -0.028554495 |
| Q96PK6 | RBM14 | | 51324000 | 89768000 | 0 | 0 | 0 | 0 | 47030666.67 | 0 | 0.144766897 | 0 | N/A |
| P98179 | RBM3 | | 331280000 | 281750000 | 169440000 | 98947000 | 201290000 | 238470000 | 260823333.3 | 179569000 | 0.269871153 | 0.688469845 | -0.538534628 |
| Q14498 | RBM39 | | 204710000 | 208790000 | 180200000 | 0 | 0 | 152520000 | 197900000 | 50840000 | 0.046442381 | 0.256897423 | -1.960735677 |
| P38159 | RBMX | | 307610000 | 310960000 | 233360000 | 214930000 | 234260000 | 277070000 | 283976666.7 | 242086666.7 | 0.251562322 | 0.85248788 | -0.230248771 |
| P18754 | RCC1 | | 59736000 | 56072000 | 39603000 | 0 | 0 | 0 | 51803666.67 | 0 | 0.001115798 | 0 | N/A |
| Q96I51 | RCC1L | | 133900000 | 107770000 | 113450000 | 108300000 | 88931000 | 75964000 | 118373333.3 | 91065000 | 0.09054961 | 0.769303334 | -0.378375534 |
| Q9P258 | RCC2 | | 0 | 52376000 | 71158000 | 93002000 | 63483000 | 50440000 | 41178000 | 68975000 | 0.32397729 | 1.675044927 | 0.744199791 |
| Q15293 | RCN1 | | 2192800000 | 2095500000 | 2710700000 | 3901600000 | 4382700000 | 3227000000 | 2333000000 | 3837100000 | 0.01755277 | 1.644706387 | 0.717830057 |
| Q14257 | RCN2 | | 1265700000 | 1486000000 | 856500000 | 734620000 | 857600000 | 2034700000 | 1202733333 | 1208973333 | 0.989682376 | 1.005188182 | 0.007465615 |
| Q96D15 | RCN3 | | 583660000 | 711630000 | 653750000 | 941930000 | 1342200000 | 1338000000 | 649680000 | 1207376667 | 0.015508047 | 1.858417477 | 0.894074627 |
| Q8TC12 | RDH11 | | 319270000 | 423380000 | 478940000 | 445130000 | 394200000 | 358300000 | 407196666.7 | 399210000 | 0.887820392 | 0.980386218 | -0.028577892 |
| Q8NBN7 | RDH13 | | 37089000 | 0 | 13627000 | 0 | 0 | 0 | 16905333.33 | 0 | 0.193599109 | 0 | N/A |
| Q9HBH5 | RDH14 | | 128380000 | 202100000 | 109240000 | 202640000 | 285310000 | 204080000 | 146573333.3 | 230676666.7 | 0.099331844 | 1.573796962 | 0.654249428 |
| P35241 | RDX | | 341900000 | 346510000 | 327880000 | 280460000 | 265410000 | 319310000 | 338763333.3 | 288393333.3 | 0.041479263 | 0.851312125 | -0.232239916 |
| P46063 | RECQL | | 0 | 0 | 224940000 | 0 | 0 | 142630000 | 74980000 | 47543333.33 | 0.7727249 | 0.634080199 | -0.657262769 |
| Q00765 | REEP5 | | 119090000 | 131260000 | 164130000 | 546850000 | 319020000 | 191850000 | 138160000 | 352573333.3 | 0.109982365 | 2.551920479 | 1.351583373 |
| Q8IUW5 | RELL1 | | 13994000 | 29487000 | 28066000 | 59784000 | 36638000 | 7607700 | 23849000 | 34676566.67 | 0.532853562 | 1.45400506 | 0.54003229 |
| Q96D71 | REPS1 | | 0 | 0 | 0 | 44024000 | 68833000 | 0 | 0 | 37619000 | 0.134971476 | N/A | N/A |
| Q86VR2 | RETREG3 | | 0 | 0 | 0 | 95190000 | 85187000 | 0 | 0 | 60125666.67 | 0.117338331 | N/A | N/A |
| Q6NUM9 | RETSAT | | 0 | 0 | 0 | 104490000 | 0 | 70599000 | 0 | 58363000 | 0.130801148 | N/A | N/A |
| P35250 | RFC2 | | 49912000 | 62316000 | 0 | 59614000 | 63630000 | 0 | 37409333.33 | 41081333.33 | 0.902114427 | 1.098157323 | 0.13508475 |
| P35249 | RFC4 | | 68686000 | 0 | 0 | 56586000 | 0 | 87126000 | 22895333.33 | 47904000 | 0.506180414 | 2.092304109 | 1.065092557 |
| Q14699 | RFTN1 | | 135150000 | 76258000 | 90844000 | 372440000 | 275140000 | 276810000 | 100750666.7 | 308130000 | 0.004837327 | 3.058342046 | 1.612749767 |
| Q96CC6 | RHBDF1 | | 0 | 7850300 | 0 | 0 | 6374000 | 0 | 2616766.667 | 2124666.667 | 0.890988715 | 0.811943493 | -0.300548769 |
| P61586 | RHOA | | 1416200000 | 1073600000 | 785490000 | 1249800000 | 1647700000 | 1999200000 | 1091763333 | 1632233333 | 0.128782512 | 1.495043187 | 0.58018716 |
| P08134 | RHOC | | 0 | 0 | 0 | 0 | 250410000 | 93836000 | 0 | 114748666.7 | 0.19126538 | N/A | N/A |
| P84095 | RHOG | | 119830000 | 229940000 | 365040000 | 259650000 | 255020000 | 159340000 | 238270000 | 224670000 | 0.870187196 | 0.942921895 | -0.084789821 |
| P17081 | RHOQ | | 0 | 0 | 32978000 | 0 | 0 | 39029000 | 10992666.67 | 13009666.67 | 0.911440614 | 1.18348596 | 0.243042592 |
| Q96DB5 | RMDN1 | | 432860000 | 396490000 | 452450000 | 221100000 | 247870000 | 438780000 | 427266666.7 | 302583333.3 | 0.151557053 | 0.708183804 | -0.497804245 |
| Q96TC7 | RMDN3 | | 455530000 | 85778000 | 467630000 | 97973000 | 164720000 | 195590000 | 336312666.7 | 152761000 | 0.22661709 | 0.454223153 | -1.138526847 |
| Q9NWS8 | RMND1 | | 0 | 68440000 | 0 | 0 | 92846000 | 38696000 | 22813333.33 | 43847333.33 | 0.58325226 | 1.922004676 | 0.942611846 |
| O00584 | RNASET2 | | 115120000 | 0 | 0 | 0 | 75453000 | 0 | 38373333.33 | 25151000 | 0.787520295 | 0.655429117 | -0.609488329 |
| P13489 | RNH1 | | 113600000 | 128890000 | 206900000 | 274210000 | 292210000 | 322180000 | 149796666.7 | 296200000 | 0.010332335 | 1.97734707 | 0.98356612 |
| Q13464 | ROCK1 | | 0 | 0 | 0 | 96127000 | 0 | 74061000 | 0 | 56729333.33 | 0.122760012 | N/A | N/A |
| O75116 | ROCK2 | | 303240000 | 361630000 | 370930000 | 114780000 | 77838000 | 230810000 | 345266666.7 | 141142666.7 | 0.015805363 | 0.408793203 | -1.290556885 |
| P35244 | RPA3 | | 0 | 0 | 0 | 42385000 | 41522000 | 39435000 | 0 | 41114000 | 1.23104E-06 | N/A | N/A |
| P27635 | RPL10 | | 528080000 | 803060000 | 731070000 | 1382700000 | 1141400000 | 597430000 | 687403333.3 | 1040510000 | 0.225116797 | 1.513681924 | 0.598062078 |
| P62906 | RPL10A | | 2152700000 | 1967400000 | 1918900000 | 2535600000 | 2561400000 | 1665000000 | 2013000000 | 2254000000 | 0.471039481 | 1.119721808 | 0.163140343 |
| P62913 | RPL11 | | 0 | 0 | 1088200000 | 0 | 0 | 1148800000 | 362733333.3 | 382933333.3 | 0.971286182 | 1.055688293 | 0.078183921 |
| P30050 | RPL12 | | 2298600000 | 2149200000 | 2694700000 | 2604100000 | 2785100000 | 3014700000 | 2380833333 | 2801300000 | 0.105227025 | 1.17660483 | 0.234629864 |
| P26373 | RPL13 | | 931180000 | 1064400000 | 1105000000 | 2064800000 | 1629700000 | 703180000 | 1033526667 | 1465893333 | 0.345783035 | 1.418341085 | 0.504204516 |
| P40429 | RPL13A | | 418770000 | 757470000 | 707490000 | 1683400000 | 1052700000 | 378870000 | 627910000 | 1038323333 | 0.353292761 | 1.653618088 | 0.725626074 |
| P50914 | RPL14 | | 817260000 | 618510000 | 617370000 | 1020900000 | 1185000000 | 971480000 | 684380000 | 1059126667 | 0.015527452 | 1.547571038 | 0.630005634 |
| P61313 | RPL15 | | 273370000 | 385270000 | 606510000 | 1535500000 | 658640000 | 385700000 | 421716666.7 | 859946666.7 | 0.290842204 | 2.039157412 | 1.027973148 |
| P18621 | RPL17 | | 317230000 | 284870000 | 392910000 | 776190000 | 586340000 | 497680000 | 331670000 | 620070000 | 0.030757328 | 1.869539 | 0.902682567 |
| Q07020 | RPL18 | | 1349200000 | 1766600000 | 1815100000 | 4795600000 | 3274100000 | 1506300000 | 1643633333 | 3192000000 | 0.182736591 | 1.942038979 | 0.957572157 |
| Q02543 | RPL18A | | 162370000 | 210760000 | 322520000 | 539740000 | 641560000 | 299530000 | 231883333.3 | 493610000 | 0.07953581 | 2.128699777 | 1.089972493 |
| P84098 | RPL19 | | 598010000 | 1238100000 | 955340000 | 1921000000 | 1711000000 | 990640000 | 930483333.3 | 1540880000 | 0.144460987 | 1.65599957 | 0.727702298 |
| P46778 | RPL21 | | 794170000 | 670660000 | 523140000 | 1025400000 | 1287900000 | 492820000 | 662656666.7 | 935373333.3 | 0.330912661 | 1.411550476 | 0.49728072 |
| P35268 | RPL22 | | 1622600000 | 1845200000 | 1861500000 | 3083900000 | 2863700000 | 1707800000 | 1776433333 | 2551800000 | 0.148304207 | 1.436473833 | 0.522531713 |
| P62829 | RPL23 | | 1174500000 | 1137300000 | 1461700000 | 1295300000 | 1484200000 | 1995600000 | 1257833333 | 1591700000 | 0.225077492 | 1.265429972 | 0.339627672 |
| P62750 | RPL23A | | 327790000 | 478000000 | 386970000 | 835500000 | 632990000 | 522870000 | 397586666.7 | 663786666.7 | 0.058542283 | 1.669539555 | 0.739450275 |
| P83731 | RPL24 | | 423300000 | 445000000 | 401540000 | 608070000 | 715310000 | 621140000 | 423280000 | 648173333.3 | 0.003358956 | 1.531311031 | 0.614767345 |
| P61254 | RPL26 | | 334860000 | 438540000 | 430170000 | 1052600000 | 604970000 | 314920000 | 401190000 | 657496666.7 | 0.303219869 | 1.63886604 | 0.712697934 |
| P61353 | RPL27 | | 1772700000 | 2120100000 | 2358800000 | 3446300000 | 3250300000 | 2356800000 | 2083866667 | 3017800000 | 0.067935513 | 1.448173268 | 0.534234225 |
| P46776 | RPL27A | | 905480000 | 1141400000 | 1375400000 | 2368100000 | 1743800000 | 1575400000 | 1140760000 | 1895766667 | 0.052496115 | 1.661845319 | 0.732786106 |

| | | | | | | | | | | | | | |
|---|---|---|---|---|---|---|---|---|---|---|---|---|---|
| P46779 | RPL28 | | 365320000 | 457080000 | 237000000 | 395430000 | 342450000 | 225140000 | 353133333.3 | 321006666.7 | 0.712793223 | 0.909023976 | -0.137609748 |
| P47914 | RPL29 | | 0 | 0 | 10184000 | 40161000 | 22320000 | 9094700 | 3394666.667 | 23858566.67 | 0.100517906 | 7.028250196 | 2.81316555 |
| P39023 | RPL3 | | 521010000 | 442630000 | 501330000 | 949270000 | 764360000 | 550070000 | 488323333.3 | 754566666.7 | 0.086523059 | 1.545219356 | 0.627811655 |
| P62888 | RPL30 | | 707320000 | 644660000 | 754480000 | 2742100000 | 1937800000 | 1880200000 | 702153333.3 | 2186700000 | 0.006079909 | 3.114277034 | 1.638897286 |
| P62899 | RPL31 | | 47898000 | 140560000 | 103290000 | 227390000 | 209030000 | 201640000 | 97249333.33 | 212686666.7 | 0.014555881 | 2.187024418 | 1.128969328 |
| P62910 | RPL32 | | 72762000 | 125210000 | 148820000 | 512410000 | 406480000 | 182240000 | 115597333.3 | 367043333.3 | 0.065552894 | 3.175188586 | 1.666842281 |
| P49207 | RPL34 | | 0 | 0 | 38297000 | 0 | 63540000 | 42144000 | 12765666.67 | 35228000 | 0.376777969 | 2.759589524 | 1.464453689 |
| P42766 | RPL35 | | 317540000 | 401400000 | 561170000 | 1616400000 | 940430000 | 299710000 | 426703333.3 | 952180000 | 0.245861953 | 2.231480107 | 1.158000946 |
| P18077 | RPL35A | | 58849000 | 81223000 | 102700000 | 128290000 | 132030000 | 113120000 | 80924000 | 124480000 | 0.035198272 | 1.538233404 | 0.621274428 |
| Q9Y3U8 | RPL36 | | 374090000 | 296380000 | 320340000 | 300180000 | 237520000 | 208710000 | 330270000 | 248803333.3 | 0.083135536 | 0.753333131 | -0.408640115 |
| P63173 | RPL38 | | 0 | 0 | 0 | 295320000 | 384720000 | 307100000 | 0 | 329046666.7 | 0.000301796 | N/A | N/A |
| Q59GN2 | RPL39P5 | | 0 | 0 | 0 | 58159000 | 44699000 | 0 | 0 | 34286000 | 0.122883076 | N/A | N/A |
| P36578 | RPL4 | | 5849500000 | 5846400000 | 6540800000 | 8745000000 | 8039000000 | 7589100000 | 6078900000 | 8124366667 | 0.007422887 | 1.336486316 | 0.418445066 |
| P46777 | RPL5 | | 478090000 | 632250000 | 996630000 | 1270800000 | 1181900000 | 1401300000 | 702323333.3 | 1284666667 | 0.024910999 | 1.829167003 | 0.871186799 |
| Q02878 | RPL6 | | 1594100000 | 1863100000 | 2017000000 | 4196100000 | 3159000000 | 2356700000 | 1824733333 | 3237266667 | 0.061049757 | 1.774103979 | 0.827090567 |
| P18124 | RPL7 | | 973940000 | 1731700000 | 1985300000 | 4860800000 | 3738400000 | 2177300000 | 1563646667 | 3592166667 | 0.072100341 | 2.297300754 | 1.199939741 |
| P62424 | RPL7A | | 1700900000 | 2193900000 | 2091200000 | 2701600000 | 2742100000 | 1831200000 | 1995333333 | 2424966667 | 0.266405878 | 1.215319078 | 0.281335138 |
| P62917 | RPL8 | | 822370000 | 1059700000 | 1555600000 | 2735900000 | 2117900000 | 1094300000 | 1145890000 | 1982700000 | 0.186288114 | 1.730270794 | 0.790997843 |
| P32969 | RPL9 | | 405060000 | 420320000 | 460480000 | 612100000 | 636050000 | 603130000 | 428620000 | 617093333.3 | 0.000607205 | 1.439721276 | 0.525789539 |
| P05388 | RPLP0 | | 3578700000 | 3823900000 | 3822500000 | 3884100000 | 4528000000 | 5064400000 | 3741700000 | 4492166667 | 0.099160972 | 1.200568369 | 0.263717563 |
| P05386 | RPLP1 | | 9227800000 | 5554400000 | 4671400000 | 6519900000 | 6427500000 | 9807700000 | 6484533333 | 7585033333 | 0.570668307 | 1.169711519 | 0.226152768 |
| P05387 | RPLP2 | | 7884800000 | 8322200000 | 6567300000 | 5912400000 | 7028700000 | 7632300000 | 7591433333 | 6857800000 | 0.371391236 | 0.903360367 | -0.146626475 |
| P04843 | RPN1 | | 6128900000 | 5853500000 | 6391300000 | 4156200000 | 4173800000 | 3899300000 | 6124566667 | 4076433333 | 0.000331618 | 0.665587225 | -0.587300351 |
| P04844 | RPN2 | | 3571000000 | 1993100000 | 3487900000 | 2948100000 | 2558400000 | 3824600000 | 3017333333 | 3110366667 | 0.890580074 | 1.030832965 | 0.043810579 |
| P46783 | RPS10 | | 375210000 | 362980000 | 251160000 | 452490000 | 333780000 | 369890000 | 329783333.3 | 385386666.7 | 0.352045557 | 1.16860565 | 0.22478817 |
| P62280 | RPS11 | | 90920000 | 99071000 | 88034000 | 121190000 | 115360000 | 40254000 | 92675000 | 92268000 | 0.988380903 | 0.995608309 | -0.006349825 |
| P25398 | RPS12 | | 1310500000 | 1520800000 | 1021100000 | 2028000000 | 2022700000 | 1980400000 | 1284133333 | 2010366667 | 0.007561134 | 1.565543557 | 0.646663648 |
| P62277 | RPS13 | | 575270000 | 450530000 | 558100000 | 783170000 | 769240000 | 743120000 | 527966666.7 | 765176666.7 | 0.004341187 | 1.449289728 | 0.535346033 |
| P62263 | RPS14 | | 77483000 | 0 | 120790000 | 537570000 | 373520000 | 271870000 | 66091000 | 394320000 | 0.018183627 | 5.966319166 | 2.576841156 |
| P62841 | RPS15 | | 47688000 | 0 | 0 | 0 | 118840000 | 123330000 | 15896000 | 80723333.33 | 0.209536792 | 5.078216742 | 2.344321972 |
| P62244 | RPS15A | | 126360000 | 164110000 | 0 | 135510000 | 140520000 | 101370000 | 96823333.33 | 125800000 | 0.601152891 | 1.299273591 | 0.377705254 |
| P62249 | RPS16 | | 0 | 245550000 | 287870000 | 404780000 | 471810000 | 587260000 | 177806666.7 | 487950000 | 0.041080063 | 2.744272806 | 1.456423906 |
| P08708 | RPS17 | | 285630000 | 231160000 | 119620000 | 541400000 | 443480000 | 400330000 | 212136666.7 | 461736666.7 | 0.017773646 | 2.176599991 | 1.122076297 |
| P62269 | RPS18 | | 238720000 | 216990000 | 75957000 | 689810000 | 418210000 | 327830000 | 177222333.3 | 478616666.7 | 0.066154282 | 2.700656614 | 1.433310214 |
| P39019 | RPS19 | | 436830000 | 578170000 | 495860000 | 974720000 | 852340000 | 1086800000 | 503620000 | 971286666.7 | 0.0041057 | 1.928610196 | 0.94756158 |
| P15880 | RPS2 | | 174580000 | 271590000 | 0 | 321290000 | 443840000 | 458460000 | 148723333.3 | 407863333.3 | 0.04589371 | 2.742430015 | 1.455454805 |
| P60866 | RPS20 | | 526960000 | 621090000 | 429880000 | 867410000 | 771340000 | 741770000 | 525976666.7 | 793506666.7 | 0.016202336 | 1.50863473 | 0.593243544 |
| P63220 | RPS21 | | 238800000 | 227320000 | 195110000 | 359440000 | 280140000 | 445350000 | 220410000 | 361643333.3 | 0.046150761 | 1.640775524 | 0.714377876 |
| P62266 | RPS23 | | 0 | 0 | 0 | 58739000 | 139430000 | 205720000 | 0 | 134629666.7 | 0.033924828 | N/A | N/A |
| P62847 | RPS24 | | 102060000 | 103740000 | 56740000 | 0 | 75080000 | 206590000 | 87513333.33 | 93890000 | 0.923408979 | 1.072865087 | 0.101468669 |
| P62851 | RPS25 | | 369320000 | 407500000 | 488520000 | 721830000 | 666700000 | 833940000 | 421780000 | 740823333.3 | 0.006184733 | 1.756421199 | 0.812638852 |
| P62854 | RPS26 | | 0 | 80219000 | 0 | 406710000 | 350540000 | 98147000 | 26739666.67 | 285132333.3 | 0.058735273 | 10.66327179 | 3.414578261 |
| P42677 | RPS27 | | 0 | 111190000 | 0 | 0 | 0 | 52563000 | 37063333.33 | 17521000 | 0.658457028 | 0.472731361 | -1.080907519 |
| P62979 | RPS27A | | 0 | 0 | 53790000 | 137380000 | 137420000 | 85823000 | 17930000 | 120207666.7 | 0.014644179 | 6.704275888 | 2.745081519 |
| P62857 | RPS28 | | 429690000 | 433750000 | 478800000 | 339770000 | 433200000 | 500640000 | 447413333.3 | 424536666.7 | 0.666284871 | 0.948869055 | -0.075719088 |
| P23396 | RPS3 | | 326840000 | 316330000 | 435200000 | 646400000 | 655100000 | 581120000 | 359456666.7 | 627540000 | 0.003854521 | 1.745801534 | 0.80388956 |
| P61247 | RPS3A | | 772240000 | 1033600000 | 865860000 | 1400400000 | 1348200000 | 813290000 | 890566666.7 | 1187296667 | 0.216864115 | 1.333192349 | 0.414884944 |
| P62701 | RPS4X | | 297180000 | 415920000 | 435040000 | 1149000000 | 672740000 | 446330000 | 382713333.3 | 756023333.3 | 0.152338287 | 1.975429826 | 0.982166598 |
| P46782 | RPS5 | | 1610400000 | 1758300000 | 1010500000 | 1671100000 | 2076900000 | 2164200000 | 1459733333 | 1970733333 | 0.136163846 | 1.350063939 | 0.433027734 |
| P62753 | RPS6 | | 0 | 315680000 | 293250000 | 518920000 | 490520000 | 435760000 | 202976666.7 | 481733333.3 | 0.056071894 | 2.373343406 | 1.246920863 |
| P62081 | RPS7 | | 941800000 | 987330000 | 681960000 | 1210900000 | 1017900000 | 1207600000 | 870363333.3 | 1145466667 | 0.074184815 | 1.316078726 | 0.396245791 |
| P62241 | RPS8 | | 2052400000 | 1823100000 | 1525600000 | 2206300000 | 2138200000 | 2515600000 | 1800366667 | 2286700000 | 0.064169967 | 1.270130159 | 0.344976347 |
| P46781 | RPS9 | | 119610000 | 100100000 | 97328000 | 220540000 | 231130000 | 197710000 | 105679333.3 | 216460000 | 0.000789881 | 2.048271816 | 1.034407181 |
| P08865 | RPSA | | 1264100000 | 1231200000 | 1256100000 | 1295800000 | 1507500000 | 1835200000 | 1250466667 | 1546166667 | 0.13315636 | 1.236471717 | 0.30622924 |
| P10301 | RRAS | | 35104000 | 106650000 | 76647000 | 164740000 | 147350000 | 38145000 | 72800333.33 | 116745000 | 0.381417756 | 1.603632767 | 0.681343802 |
| Q9P2E9 | RRBP1 | | 3140800000 | 3229400000 | 3355600000 | 4019300000 | 3683500000 | 2083200000 | 3241933333 | 3262000000 | 0.974945919 | 1.006189722 | 0.008902358 |
| O76021 | RSL1D1 | | 40299000 | 59362000 | 89118000 | 81221000 | 82022000 | 75444000 | 62926333.33 | 79562333.33 | 0.31098132 | 1.264372626 | 0.338421706 |

| | | | | | | | | | | | | | |
|---|---|---|---|---|---|---|---|---|---|---|---|---|---|
| Q96DX4 | RSPRY1 | | 80417000 | 0 | 0 | 0 | 71947000 | 58333000 | 26805666.67 | 43426666.67 | 0.657130835 | 1.620055461 | 0.696043203 |
| Q15404 | RSU1 | | 0 | 0 | 162420000 | 0 | 39539000 | 0 | 54140000 | 13179666.67 | 0.503053687 | 0.243436769 | -2.038381005 |
| Q9Y3I0 | RTCB | | 351170000 | 193800000 | 188880000 | 295530000 | 311180000 | 341790000 | 244616666.7 | 316166666.7 | 0.263167189 | 1.292498467 | 0.370162569 |
| Q9NQC3 | RTN4 | | 253080000 | 197670000 | 877880000 | 1080800000 | 1231300000 | 2085100000 | 442876666.7 | 1465733333 | 0.055063621 | 3.309574524 | 1.726645757 |
| Q9Y224 | RTRAF | | 254340000 | 249670000 | 241440000 | 250320000 | 200740000 | 286130000 | 248483333.3 | 245730000 | 0.917745935 | 0.988919445 | -0.016075088 |
| Q96GQ5 | RUSF1 | | 114430000 | 85732000 | 127690000 | 72695000 | 110680000 | 201450000 | 109284000 | 128275000 | 0.660885989 | 1.173776582 | 0.23115783 |
| Q9Y265 | RUVBL1 | | 127530000 | 200810000 | 201700000 | 288310000 | 370550000 | 402690000 | 176680000 | 353850000 | 0.013494982 | 2.002773376 | 1.001999182 |
| Q9Y230 | RUVBL2 | | 122170000 | 134330000 | 192720000 | 417200000 | 479860000 | 408760000 | 149740000 | 435273333.3 | 0.000796987 | 2.906860781 | 1.539461978 |
| P60903 | S100A10 | | 540140000 | 752390000 | 937700000 | 1905900000 | 1383500000 | 1027200000 | 743410000 | 1438866667 | 0.067821131 | 1.935495442 | 0.95270291 |
| P31949 | S100A11 | | 111780000 | 0 | 24452000 | 126920000 | 253640000 | 160740000 | 45410666.67 | 180433333.3 | 0.056680177 | 3.973368959 | 1.990362765 |
| Q96FQ6 | S100A16 | | 116020000 | 0 | 0 | 31953000 | 56486000 | 51195000 | 38673333.33 | 46544666.67 | 0.851342772 | 1.203533873 | 0.267276747 |
| Q9NTJ5 | SACM1L | | 208390000 | 220410000 | 0 | 126490000 | 158770000 | 0 | 142933333.3 | 95086666.67 | 0.609267085 | 0.665251866 | -0.588027444 |
| Q9Y512 | SAMM50 | | 138470000 | 209530000 | 169570000 | 241030000 | 371450000 | 50037000 | 172523333.3 | 220839000 | 0.639759284 | 1.28005294 | 0.356203478 |
| Q9NR31 | SAR1A | | 0 | 0 | 109560000 | 183190000 | 232550000 | 0 | 36520000 | 138580000 | 0.26910632 | 3.794633078 | 1.923960391 |
| P49591 | SARS1 | | 377990000 | 360240000 | 297660000 | 736840000 | 583270000 | 707280000 | 345296666.7 | 675796666.7 | 0.003364274 | 1.95714796 | 0.968752827 |
| Q9Y3A5 | SBDS | | 0 | 0 | 0 | 0 | 175690000 | 206500000 | 0 | 127396666.7 | 0.118695006 | N/A | N/A |
| O15126 | SCAMP1 | | 782880000 | 791620000 | 731130000 | 840260000 | 766620000 | 1198300000 | 768543333.3 | 935060000 | 0.283874961 | 1.216665293 | 0.282932335 |
| O15127 | SCAMP2 | | 232570000 | 151610000 | 158230000 | 265570000 | 277980000 | 237620000 | 180803333.3 | 260390000 | 0.049513752 | 1.440183625 | 0.526252769 |
| O14828 | SCAMP3 | | 495870000 | 501330000 | 543970000 | 567060000 | 729520000 | 618300000 | 513723333.3 | 638293333.3 | 0.068481848 | 1.242484606 | 0.313227977 |
| Q969E2 | SCAMP4 | | 0 | 0 | 150290000 | 0 | 103920000 | 216540000 | 50096666.67 | 106820000 | 0.518031861 | 2.132277597 | 1.092395272 |
| Q14108 | SCARB2 | | 0 | 54349000 | 137660000 | 674030000 | 531130000 | 342140000 | 64003000 | 515766666.7 | 0.012263339 | 8.058476426 | 3.010507102 |
| Q8NBX0 | SCCPDH | | 452610000 | 489910000 | 569800000 | 955640000 | 764930000 | 607220000 | 504106666.7 | 775930000 | 0.063143202 | 1.53921789 | 0.622197473 |
| Q8WVM8 | SCFD1 | | 1177600000 | 891400000 | 1057400000 | 1081800000 | 1091600000 | 1102900000 | 1042133333 | 1092100000 | 0.580477465 | 1.04794652 | 0.067565093 |
| Q8WU76 | SCFD2 | | 72786000 | 63244000 | 47519000 | 41121000 | 45405000 | 0 | 61183000 | 28842000 | 0.117267894 | 0.471405456 | -1.084959639 |
| P22307 | SCP2 | | 484370000 | 566010000 | 579610000 | 245180000 | 210690000 | 186810000 | 543330000 | 214226666.7 | 0.000654101 | 0.394284628 | -1.342690632 |
| Q9HB40 | SCPEP1 | | 0 | 0 | 0 | 91183000 | 49172000 | 106950000 | 0 | 82435000 | 0.008770636 | N/A | N/A |
| Q14160 | SCRIB | | 1564500000 | 1485000000 | 1497200000 | 1339800000 | 1466500000 | 1332600000 | 1515566667 | 1379633333 | 0.05311338 | 0.910308575 | -0.135572423 |
| P18827 | SDC1 | | 90971000 | 0 | 148330000 | 0 | 0 | 0 | 79767000 | 0 | 0.138447756 | 0 | N/A |
| O00560 | SDCBP | | 0 | 0 | 105880000 | 75935000 | 138090000 | 159830000 | 35293333.33 | 124618333.3 | 0.108269842 | 3.530931243 | 1.820048728 |
| Q99470 | SDF2 | | 77302000 | 0 | 0 | 101010000 | 123190000 | 180140000 | 25767333.33 | 134780000 | 0.035462291 | 5.230653799 | 2.386991286 |
| Q9HCN8 | SDF2L1 | | 532330000 | 506730000 | 462460000 | 328540000 | 425720000 | 501180000 | 500506666.7 | 418480000 | 0.203212709 | 0.836112739 | -0.258230611 |
| Q9BRK5 | SDF4 | | 145980000 | 194440000 | 200310000 | 236720000 | 204310000 | 189870000 | 180243333.3 | 210300000 | 0.245358976 | 1.166756052 | 0.22502951 |
| Q9NRG7 | SDR39U1 | | 0 | 0 | 10416000 | 24534000 | 31533000 | 0 | 3472000 | 18689000 | 0.208969852 | 5.382776498 | 2.428350523 |
| Q96IW7 | SEC22A | | 0 | 33827000 | 0 | 39981000 | 47349000 | 96754000 | 11275666.67 | 61361333.33 | 0.076424136 | 5.441925089 | 2.444117097 |
| O75396 | SEC22B | | 1093200000 | 1105800000 | 1135700000 | 930450000 | 817930000 | 1076600000 | 1111566667 | 941660000 | 0.088864798 | 0.84714667 | -0.239316324 |
| Q15436 | SEC23A | | 825860000 | 876830000 | 704100000 | 526970000 | 759420000 | 963110000 | 802263333.3 | 749833333.3 | 0.719509008 | 0.934647393 | -0.097505901 |
| Q15437 | SEC23B | | 106640000 | 107140000 | 72274000 | 61099000 | 63497000 | 128490000 | 95351333.33 | 84362000 | 0.681881961 | 0.884749033 | -0.176659814 |
| O95487 | SEC24B | | 32041000 | 0 | 41933000 | 0 | 0 | 29075000 | 24658000 | 9691666.667 | 0.400935376 | 0.393043502 | -1.347239097 |
| P53992 | SEC24C | | 679990000 | 588420000 | 654570000 | 655980000 | 642260000 | 654550000 | 640993333.3 | 650930000 | 0.737366723 | 1.015501981 | 0.022193054 |
| O94979 | SEC31A | | 0 | 0 | 0 | 150870000 | 166060000 | 176720000 | 0 | 164550000 | 2.55443E-05 | N/A | N/A |
| P61619 | SEC61A1 | | 58069000 | 35448000 | 209510000 | 279580000 | 358460000 | 394110000 | 101009000 | 344050000 | 0.019410626 | 3.406132127 | 1.7681344 |
| P60059 | SEC61G | | 723220000 | 583910000 | 589130000 | 154010000 | 155350000 | 311640000 | 632086666.7 | 207000000 | 0.003598787 | 0.327486737 | -1.610491615 |
| Q9UGP8 | SEC63 | | 0 | 101160000 | 87150000 | 238960000 | 181860000 | 235140000 | 62770000 | 218653333.3 | 0.013087183 | 3.483405024 | 1.800498228 |
| Q9UBV2 | SEL1L | | 0 | 137800000 | 0 | 37733000 | 38402000 | 106370000 | 45933333.33 | 60835000 | 0.785751604 | 1.324419448 | 0.405360101 |
| O60613 | SELENOF | | 55800000 | 74169000 | 250130000 | 212650000 | 160990000 | 284880000 | 126699666.7 | 219506666.7 | 0.264683841 | 1.732496008 | 0.792852028 |
| Q9C0D9 | SELENOI | | 0 | 0 | 0 | 110200000 | 191840000 | 0 | 0 | 100680000 | 0.144324838 | N/A | N/A |
| O75326 | SEMA7A | | 0 | 0 | 0 | 125490000 | 184260000 | 242700000 | 0 | 184150000 | 0.005533929 | N/A | N/A |
| Q9P0V9 | SEPTIN10 | | 304880000 | 260860000 | 258580000 | 119110000 | 154730000 | 451700000 | 274773333.3 | 241846666.7 | 0.772627867 | 0.880167896 | -0.184149344 |
| Q9NVA2 | SEPTIN11 | | 742550000 | 1126100000 | 1023400000 | 2330400000 | 1502300000 | 831360000 | 964016666.7 | 1554686667 | 0.258157595 | 1.612717622 | 0.689493853 |
| Q15019 | SEPTIN2 | | 2686800000 | 3321000000 | 3392800000 | 3914000000 | 3494700000 | 4105200000 | 3133533333 | 3837966667 | 0.070621494 | 1.2248048 | 0.292551842 |
| Q99719 | SEPTIN5 | | 206870000 | 150500000 | 111730000 | 109440000 | 109450000 | 0 | 156366666.7 | 72963333.33 | 0.142430571 | 0.466616926 | -1.099689454 |
| Q14141 | SEPTIN6 | | 55510000 | 74057000 | 0 | 0 | 155060000 | 0 | 43189000 | 51686666.67 | 0.887276275 | 1.196755347 | 0.259128251 |
| Q16181 | SEPTIN7 | | 2548100000 | 2408800000 | 3155000000 | 3476900000 | 3198300000 | 3705300000 | 2703966667 | 3460166667 | 0.049793368 | 1.279663211 | 0.355764163 |
| Q92599 | SEPTIN8 | | 213890000 | 196110000 | 195130000 | 653500000 | 495590000 | 279150000 | 201710000 | 476080000 | 0.06503328 | 2.360220118 | 1.238921414 |
| Q9UHD8 | SEPTIN9 | | 1348800000 | 1438600000 | 1380900000 | 1859900000 | 1894900000 | 1284400000 | 1389433333 | 1679733333 | 0.21963053 | 1.208934098 | 0.273735602 |
| Q8NC51 | SERBP1 | | 218090000 | 236220000 | 107050000 | 239550000 | 400170000 | 289220000 | 187120000 | 309646666.7 | 0.120730923 | 1.654802622 | 0.726659149 |
| O75635 | SERPINB7 | | 0 | 0 | 59618000 | 40020000 | 0 | 0 | 19872666.67 | 13340000 | 0.798414695 | 0.671273776 | -0.575026811 |

| | | | | | | | | | | | | |
|---|---|---|---|---|---|---|---|---|---|---|---|---|
| P50454 | SERPINH1 | 18070000000 | 13389000000 | 14795000000 | 20970000000 | 22130000000 | 22297000000 | 15418000000 | 21799000000 | 0.011634003 | 1.413866909 | 0.499646322 |
| O75533 | SF3B1 | 78986000 | 72093000 | 91511000 | 132980000 | 96486000 | 0 | 80863333.33 | 76488666.67 | 0.91832886 | 0.945900491 | -0.080239676 |
| P31947 | SFN | 159650000 | 33708000 | 51715000 | 192910000 | 135850000 | 47037000 | 81691000 | 125265666.7 | 0.493279092 | 1.533408413 | 0.616742 |
| P23246 | SFPQ | 581990000 | 756530000 | 715420000 | 564270000 | 509040000 | 593660000 | 684646666.7 | 555656666.7 | 0.091097928 | 0.811596249 | -0.301165897 |
| Q9H9B4 | SFXN1 | 2145300000 | 1840400000 | 1836200000 | 1237200000 | 1334300000 | 1107100000 | 1940633333 | 1226200000 | 0.004202515 | 0.63185558 | -0.662333249 |
| Q96NB2 | SFXN2 | 99541000 | 0 | 0 | 0 | 0 | 78160000 | 33180333.33 | 26053333.33 | 0.87404291 | 0.785204087 | -0.348860413 |
| Q9BWM7 | SFXN3 | 861790000 | 1011100000 | 1091500000 | 853980000 | 962710000 | 871560000 | 988130000 | 896083333.3 | 0.28846014 | 0.906847615 | -0.141067952 |
| Q6P4A7 | SFXN4 | 105810000 | 81163000 | 0 | 0 | 62391000 | 0 | 62324333.33 | 20797000 | 0.337377973 | 0.333689891 | -1.583420113 |
| O95470 | SGPL1 | 351380000 | 600790000 | 429210000 | 488780000 | 570620000 | 418710000 | 460460000 | 492703333.3 | 0.726029081 | 1.070024179 | 0.097643397 |
| Q9P0V3 | SH3BP4 | 115220000 | 130060000 | 115530000 | 0 | 24740000 | 0 | 120270000 | 8246666.667 | 0.000307145 | 0.068567944 | -3.866321919 |
| Q9Y371 | SH3GLB1 | 0 | 0 | 0 | 153200000 | 143570000 | 0 | 0 | 98923333.33 | 0.116535171 | N/A | N/A |
| P29353 | SHC1 | 125080000 | 137820000 | 0 | 168260000 | 173780000 | 187620000 | 87633333.33 | 176553333.3 | 0.11543807 | 2.014682389 | 1.010552418 |
| Q96FS4 | SIPA1 | 124580000 | 58211000 | 93485000 | 0 | 0 | 0 | 92092000 | 0 | 0.008625912 | 0 | N/A |
| P63208 | SKP1 | 0 | 0 | 0 | 46303000 | 19008000 | 0 | 0 | 21770333.33 | 0.180526857 | N/A | N/A |
| P55011 | SLC12A2 | 1355900000 | 1186200000 | 1146600000 | 1039400000 | 942070000 | 836680000 | 1229566667 | 939383333.3 | 0.028828525 | 0.763995446 | -0.388364057 |
| Q9Y666 | SLC12A7 | 0 | 59249000 | 0 | 95324000 | 0 | 0 | 19749666.67 | 31774666.67 | 0.763987587 | 1.608871036 | 0.686048687 |
| Q9BXP2 | SLC12A9 | 46571000 | 0 | 66589000 | 163560000 | 208020000 | 303960000 | 37720000 | 225180000 | 0.015030788 | 5.969777306 | 2.577677115 |
| P53985 | SLC16A1 | 178700000 | 253870000 | 262850000 | 369630000 | 149810000 | 74045000 | 231806666.7 | 197828333.3 | 0.732170304 | 0.853419516 | -0.228672992 |
| P43007 | SLC1A4 | 127950000 | 139640000 | 156900000 | 111050000 | 0 | 143670000 | 141496666.7 | 84906666.67 | 0.270471574 | 0.60006125 | -0.736818327 |
| Q15758 | SLC1A5 | 554220000 | 1418900000 | 1965300000 | 730810000 | 958390000 | 1143400000 | 1312806667 | 944200000 | 0.437439963 | 0.719222429 | -0.475490082 |
| P12235 | SLC25A4 | 229440000 | 106510000 | 237940000 | 37185000 | 32722000 | 76091000 | 191296666.7 | 48666000 | 0.033058441 | 0.254400669 | -1.97482563 |
| P05141 | SLC25A5 | 4874000000 | 5326300000 | 4951700000 | 8754300000 | 8448000000 | 8343800000 | 5050666667 | 8515366667 | 4.911E-05 | 1.685988648 | 0.753594823 |
| P12236 | SLC25A6 | 1065200000 | 702070000 | 679990000 | 461190000 | 717340000 | 1244300000 | 815753333.3 | 807610000 | 0.976709694 | 0.990017407 | -0.014474203 |
| P11166 | SLC2A1 | 603420000 | 505910000 | 614500000 | 1109600000 | 1183000000 | 605560000 | 574610000 | 966053333.3 | 0.101465925 | 1.681233068 | 0.749519738 |
| P11169 | SLC2A3 | 0 | 0 | 73365000 | 0 | 178640000 | 101570000 | 24455000 | 93403333.33 | 0.294624908 | 3.81939617 | 1.933344572 |
| Q8NEW0 | SLC30A7 | 0 | 0 | 0 | 0 | 154670000 | 104580000 | 0 | 86416666.67 | 0.13074842 | N/A | N/A |
| Q6PML9 | SLC30A9 | 0 | 0 | 77068000 | 0 | 128490000 | 0 | 25689333.33 | 42830000 | 0.748726048 | 1.667228941 | 0.737452226 |
| L0R6Q1 | SLC35A4 | 0 | 163010000 | 259130000 | 149690000 | 178290000 | 115710000 | 140713333.3 | 147896666.7 | 0.930842188 | 1.051049415 | 0.071830499 |
| Q8TB61 | SLC35B2 | 92080000 | 125830000 | 101930000 | 93370000 | 55950000 | 109220000 | 106613333.3 | 86180000 | 0.335994913 | 0.808341671 | -0.306962873 |
| Q96K37 | SLC35E1 | 52985000 | 52893000 | 58289000 | 202450000 | 130820000 | 0 | 54722333.33 | 111090000 | 0.395628998 | 2.030066944 | 1.021527303 |
| Q9HBR0 | SLC38A10 | 52628000 | 0 | 52041000 | 0 | 0 | 0 | 34889666.67 | 0 | 0.116129033 | 0 | N/A |
| Q96QD8 | SLC38A2 | 0 | 90288000 | 119030000 | 0 | 0 | 0 | 69772666.67 | 0 | 0.123561769 | 0 | N/A |
| Q8NBW4 | SLC38A9 | 0 | 0 | 0 | 120290000 | 125220000 | 0 | 0 | 81836666.67 | 0.116276884 | N/A | N/A |
| Q9ULF5 | SLC39A10 | 75112000 | 72712000 | 0 | 68408000 | 46508000 | 0 | 49274666.67 | 38305333.33 | 0.747867921 | 0.777383916 | -0.363300836 |
| Q15043 | SLC39A14 | 1153700000 | 957490000 | 1133000000 | 352150000 | 476520000 | 476660000 | 1081396667 | 435110000 | 0.000986637 | 0.402359295 | -1.313443735 |
| P08195 | SLC3A2 | 5692700000 | 5796900000 | 6180700000 | 8641500000 | 8452000000 | 7203500000 | 5890100000 | 8099000000 | 0.009649093 | 1.3750191 | 0.459451659 |
| Q9Y6M7 | SLC4A7 | 68315000 | 24340000 | 0 | 0 | 0 | 0 | 30885000 | 0 | 0.197238976 | 0 | N/A |
| P53794 | SLC5A3 | 81998000 | 63834000 | 69269000 | 0 | 33648000 | 0 | 71700333.33 | 11216000 | 0.008268433 | 0.15642884 | -2.676421572 |
| Q9Y289 | SLC5A6 | 306730000 | 307880000 | 205610000 | 67935000 | 69112000 | 0 | 273406666.7 | 45682333.33 | 0.005087574 | 0.16708566 | -2.581340175 |
| Q01650 | SLC7A5 | 52149000 | 46242000 | 36862000 | 123240000 | 92439000 | 35039000 | 45084333.33 | 83572666.67 | 0.21611305 | 1.853696406 | 0.890404982 |
| O14745 | SLC9A3R1 | 374270000 | 338930000 | 470180000 | 516550000 | 458630000 | 478590000 | 394460000 | 484590000 | 0.102583368 | 1.228489581 | 0.296885622 |
| Q15599 | SLC9A3R2 | 276390000 | 291120000 | 242930000 | 574390000 | 308100000 | 90959000 | 270146666.7 | 324483000 | 0.718682412 | 1.20113642 | 0.264400015 |
| Q9NWH9 | SLTM | 0 | 0 | 81572000 | 88623000 | 0 | 81304000 | 27190666.67 | 56642333.33 | 0.495462624 | 2.083153533 | 1.058769173 |
| O95347 | SMC2 | 308880000 | 285640000 | 210480000 | 290830000 | 211030000 | 216370000 | 268333333.3 | 239410000 | 0.502664553 | 0.89221118 | -0.164542869 |
| Q9NTJ3 | SMC4 | 179160000 | 135730000 | 101590000 | 96484000 | 106530000 | 84063000 | 138826666.7 | 95692333.33 | 0.13864578 | 0.689293604 | -0.536809467 |
| O00161 | SNAP23 | 243790000 | 313780000 | 371150000 | 233880000 | 158610000 | 0 | 309573333.3 | 130830000 | 0.084117235 | 0.42261392 | -1.242587807 |
| O95295 | SNAPIN | 111640000 | 0 | 146330000 | 149090000 | 91549000 | 118830000 | 85990000 | 119823000 | 0.512870764 | 1.393452727 | 0.47866406 |
| Q7KZF4 | SND1 | 2784100000 | 2646100000 | 2415500000 | 2848600000 | 3453300000 | 3979100000 | 2615233333 | 3427000000 | 0.077596313 | 1.310399327 | 0.390006521 |
| O75643 | SNRNP200 | 100530000 | 109460000 | 78565000 | 0 | 0 | 0 | 96185000 | 0 | 0.000468878 | 0 | N/A |
| P08621 | SNRNP70 | 175060000 | 98020000 | 0 | 122450000 | 152950000 | 47898000 | 91026666.67 | 107766000 | 0.792390725 | 1.183894829 | 0.243540926 |
| P62314 | SNRPD1 | 0 | 0 | 0 | 172620000 | 195900000 | 0 | 0 | 122840000 | 0.117701409 | N/A | N/A |
| P63162 | SNRPN | 23959000 | 26321000 | 28513000 | 27413000 | 31735000 | 32725000 | 26264333.33 | 30624333.33 | 0.105881666 | 1.166004594 | 0.221573473 |
| Q13425 | SNTB2 | 173550000 | 104050000 | 184830000 | 0 | 218150000 | 0 | 154143333.3 | 72716666.67 | 0.349802201 | 0.471747075 | -1.083914522 |
| Q9UMY4 | SNX12 | 0 | 0 | 0 | 55002000 | 67848000 | 0 | 0 | 40950000 | 0.120447494 | N/A | N/A |
| O60749 | SNX2 | 0 | 199480000 | 172360000 | 208480000 | 134070000 | 177570000 | 123946666.7 | 173373333.3 | 0.496106205 | 1.398773666 | 0.484162541 |
| O60493 | SNX3 | 0 | 0 | 36365000 | 0 | 0 | 70764000 | 12121666.67 | 23588000 | 0.687768543 | 1.945937027 | 0.960465024 |
| Q9Y5X3 | SNX5 | 164500000 | 154100000 | 179970000 | 0 | 190010000 | 257020000 | 166190000 | 149010000 | 0.835091471 | 0.896624346 | -0.157424422 |

| | | | | | | | | | | | | | |
|---|---|---|---|---|---|---|---|---|---|---|---|---|---|
| Q9UNH7 | SNX6 | | 101040000 | 157080000 | 97726000 | 150150000 | 120360000 | 232170000 | 118615333.3 | 167560000 | 0.273361641 | 1.412633555 | 0.498387271 |
| Q9Y5X2 | SNX8 | | 0 | 51250000 | 0 | 58789000 | 0 | 0 | 17083333.33 | 19596333.33 | 0.927642707 | 1.147102439 | 0.197994233 |
| Q9Y5X1 | SNX9 | | 0 | 0 | 74446000 | 70295000 | 49022000 | 0 | 24815333.33 | 39772333.33 | 0.668217091 | 1.602732182 | 0.68053337 |
| P35610 | SOAT1 | | 0 | 0 | 112430000 | 0 | 88244000 | 117260000 | 37476666.67 | 68501333.33 | 0.579071231 | 1.827839545 | 0.87013943 |
| Q99523 | SORT1 | | 0 | 0 | 0 | 73852000 | 70103000 | 61620000 | 0 | 68525000 | 4.57796E-05 | N/A | N/A |
| Q07617 | SPAG1 | | 0 | 0 | 89470000 | 77424000 | 0 | 0 | 29823333.33 | 25808000 | 0.923807224 | 0.865362691 | -0.208623172 |
| P09486 | SPARC | | 892110000 | 1040200000 | 862010000 | 2637900000 | 3071300000 | 2984500000 | 931440000 | 2897900000 | 0.000163772 | 3.111204157 | 1.637473068 |
| Q8TB22 | SPATA20 | | 148110000 | 117900000 | 118100000 | 164170000 | 0 | 0 | 128036666.7 | 54723333.33 | 0.257997774 | 0.427403608 | -1.226329007 |
| Q15005 | SPCS2 | | 59191000 | 95970000 | 55331000 | 79737000 | 49633000 | 15100000 | 70164000 | 48156666.67 | 0.387678426 | 0.686344374 | -0.542995463 |
| Q9H2V7 | SPNS1 | | 0 | 0 | 0 | 210990000 | 194890000 | 0 | 0 | 135293333.3 | 0.116741994 | N/A | N/A |
| P35270 | SPR | | 329700000 | 368430000 | 298940000 | 253550000 | 390540000 | 315750000 | 332356666.7 | 319946666.7 | 0.793768968 | 0.962660596 | -0.054900857 |
| Q5W111 | SPRYD7 | | 51595000 | 52489000 | 40390000 | 0 | 0 | 0 | 48158000 | 0 | 0.000245323 | 0 | N/A |
| Q13813 | SPTAN1 | | 7088000000 | 7392500000 | 7295600000 | 4873700000 | 4833200000 | 4027000000 | 7258700000 | 4577966667 | 0.000761258 | 0.630686854 | -0.665004233 |
| Q01082 | SPTBN1 | | 9029100000 | 9145600000 | 8674100000 | 5918500000 | 5729100000 | 5410500000 | 8949600000 | 5686033333 | 9.12178E-05 | 0.635339382 | -0.654400646 |
| O15269 | SPTLC1 | | 321820000 | 179550000 | 230620000 | 301300000 | 297260000 | 310390000 | 243996666.7 | 302983333.3 | 0.230953604 | 1.241751937 | 0.312376997 |
| O15270 | SPTLC2 | | 104650000 | 0 | 288140000 | 83135000 | 59197000 | 76052000 | 130930000 | 72794666.67 | 0.5293199 | 0.555981568 | -0.84689104 |
| Q14534 | SQLE | | 0 | 0 | 0 | 84479000 | 96360000 | 0 | 0 | 60279666.67 | 0.117830568 | N/A | N/A |
| Q13501 | SQSTM1 | | 192940000 | 146800000 | 163690000 | 98729000 | 103500000 | 120830000 | 167810000 | 107686333.3 | 0.016226295 | 0.641715829 | -0.639993524 |
| O75044 | SRGAP2 | | 50863000 | 33632000 | 0 | 0 | 0 | 0 | 28165000 | 0 | 0.132388336 | 0 | N/A |
| P37108 | SRP14 | | 0 | 182060000 | 0 | 485780000 | 422010000 | 383510000 | 60686666.67 | 430433333.3 | 0.005440867 | 7.092716687 | 2.826338321 |
| P61011 | SRP54 | | 0 | 0 | 0 | 198520000 | 0 | 142420000 | 0 | 113646666.7 | 0.126772903 | N/A | N/A |
| Q9UHB9 | SRP68 | | 208450000 | 175670000 | 120040000 | 197570000 | 247830000 | 379330000 | 168053333.3 | 274910000 | 0.149616748 | 1.63584973 | 0.710040228 |
| O76094 | SRP72 | | 150920000 | 124780000 | 259520000 | 368320000 | 387650000 | 337290000 | 178406666.7 | 364420000 | 0.013173299 | 2.042636673 | 1.030432612 |
| P49458 | SRP9 | | 606780000 | 661310000 | 454450000 | 486040000 | 540330000 | 692040000 | 574180000 | 572803333.3 | 0.98818155 | 0.997602378 | -0.003463191 |
| P08240 | SRPRA | | 559700000 | 550980000 | 759500000 | 805320000 | 921760000 | 1127900000 | 623393333.3 | 951660000 | 0.047737538 | 1.526580329 | 0.610303507 |
| Q9Y5M8 | SRPRB | | 249080000 | 221490000 | 348780000 | 1261600000 | 1168100000 | 782810000 | 273116666.7 | 1070836667 | 0.006236485 | 3.920803076 | 1.971149184 |
| Q07955 | SRSF1 | | 36370000 | 66230000 | 72995000 | 106530000 | 76171000 | 99807000 | 58531666.67 | 94169333.33 | 0.070341859 | 1.6088613 | 0.686039957 |
| P84103 | SRSF3 | | 267280000 | 260360000 | 261880000 | 400150000 | 237090000 | 151110000 | 263173333.3 | 262783333.3 | 0.9959965 | 0.998518087 | -0.002139534 |
| P43307 | SSR1 | | 325750000 | 396340000 | 309600000 | 305180000 | 494080000 | 596990000 | 343896666.7 | 465416666.7 | 0.246096462 | 1.35336196 | 0.436547743 |
| Q9UNL2 | SSR3 | | 701640000 | 586170000 | 540360000 | 481460000 | 778090000 | 754360000 | 609390000 | 671303333.3 | 0.592461773 | 1.101598867 | 0.13959898 |
| P51571 | SSR4 | | 1008000000 | 1059300000 | 959990000 | 1490000000 | 1676200000 | 1814200000 | 1009096667 | 1660133333 | 0.002686614 | 1.64516779 | 0.718234732 |
| Q08945 | SSRP1 | | 167230000 | 212410000 | 0 | 0 | 0 | 172260000 | 126546666.7 | 57420000 | 0.468645584 | 0.453745654 | -1.140044271 |
| P50502 | ST13 | | 175590000 | 0 | 0 | 316670000 | 273680000 | 288490000 | 58530000 | 292946666.7 | 0.017316885 | 5.005068626 | 2.32338985 |
| Q14849 | STARD3 | | 0 | 0 | 0 | 34325000 | 38953000 | 40722000 | 0 | 38000000 | 3.74361E-05 | N/A | N/A |
| P42224 | STAT1 | | 0 | 99974000 | 152050000 | 87354000 | 108410000 | 0 | 84008000 | 65254666.67 | 0.752871655 | 0.776767292 | -0.364445642 |
| O95793 | STAU1 | | 127130000 | 121710000 | 171450000 | 163700000 | 134710000 | 141350000 | 140096666.7 | 146586666.7 | 0.737087059 | 1.046325156 | 0.065331254 |
| Q658P3 | STEAP3 | | 225230000 | 249160000 | 0 | 476550000 | 457660000 | 0 | 158130000 | 311403333.3 | 0.430189786 | 1.969286874 | 0.97767329 |
| Q13586 | STIM1 | | 33436000 | 28037000 | 35974000 | 39746000 | 37045000 | 0 | 32482333.33 | 25597000 | 0.625270526 | 0.788028364 | -0.343680536 |
| P31948 | STIP1 | | 185000000 | 73268000 | 171310000 | 357100000 | 214660000 | 167960000 | 143192666.7 | 246573333.3 | 0.197090369 | 1.721969002 | 0.784059172 |
| Q9Y6E0 | STK24 | | 0 | 0 | 0 | 127420000 | 214980000 | 174490000 | 0 | 172296666.7 | 0.002429696 | N/A | N/A |
| P16949 | STMN1 | | 0 | 0 | 0 | 107110000 | 124750000 | 149990000 | 0 | 127283333.3 | 0.000514748 | N/A | N/A |
| Q9Y3F4 | STRAP | | 91422000 | 0 | 0 | 261940000 | 347080000 | 269360000 | 30474000 | 292793333.3 | 0.003027455 | 9.607971823 | 3.26423192 |
| P46977 | STT3A | | 512470000 | 380820000 | 316120000 | 526410000 | 551010000 | 573620000 | 403136666.7 | 550346666.7 | 0.068202164 | 1.365161525 | 0.44907166 |
| Q8TCJ2 | STT3B | | 139480000 | 181690000 | 176850000 | 172560000 | 146900000 | 188000000 | 166006666.7 | 169153333.3 | 0.869225271 | 1.018955062 | 0.027090427 |
| O60499 | STX10 | | 58673000 | 54092000 | 0 | 0 | 28156000 | 0 | 37588333.33 | 9385333.333 | 0.251336513 | 0.249687403 | -2.001805057 |
| Q86Y82 | STX12 | | 230130000 | 262100000 | 0 | 284900000 | 257310000 | 272880000 | 164076666.7 | 271696666.7 | 0.264215452 | 1.655912886 | 0.727626778 |
| O14662 | STX16 | | 71070000 | 114090000 | 80334000 | 81487000 | 64981000 | 188100000 | 88498000 | 111522666.7 | 0.602142404 | 1.260171605 | 0.333620207 |
| Q9P2W9 | STX18 | | 30438000 | 0 | 0 | 19236000 | 52726000 | 36525000 | 10146000 | 36162333.33 | 0.136997045 | 3.564196071 | 1.833576703 |
| Q13277 | STX3 | | 69072000 | 177510000 | 95077000 | 84502000 | 87281000 | 0 | 113886333.3 | 57261000 | 0.262534626 | 0.502790794 | -0.991969859 |
| Q12846 | STX4 | | 62820000 | 66715000 | 87234000 | 190790000 | 149430000 | 152090000 | 72256333.33 | 164103333.3 | 0.003932809 | 2.271127329 | 1.183408592 |
| Q13190 | STX5 | | 90400000 | 0 | 0 | 195440000 | 296700000 | 0 | 30133333.33 | 164046666.7 | 0.219797788 | 5.444026549 | 2.444674102 |
| O15400 | STX7 | | 332590000 | 357060000 | 505530000 | 526080000 | 336920000 | 491960000 | 398393333.3 | 451653333.3 | 0.5391956 | 1.133686976 | 0.181022351 |
| Q9UNK0 | STX8 | | 269790000 | 232920000 | 296930000 | 376790000 | 282660000 | 281040000 | 266546666.7 | 313496666.7 | 0.269798837 | 1.176141764 | 0.234061963 |
| P61764 | STXBP1 | | 0 | 0 | 0 | 114500000 | 75023000 | 82685000 | 0 | 90736000 | 0.001684532 | N/A | N/A |
| Q15833 | STXBP2 | | 81064000 | 0 | 149150000 | 89928000 | 97689000 | 0 | 76738000 | 62539000 | 0.803115147 | 0.814967813 | -0.295185014 |
| O00186 | STXBP3 | | 217660000 | 147190000 | 203830000 | 366770000 | 267900000 | 622720000 | 189560000 | 419130000 | 0.100468107 | 2.211067736 | 1.144743223 |
| P53999 | SUB1 | | 126810000 | 0 | 0 | 199330000 | 135640000 | 308420000 | 42270000 | 214463333.3 | 0.059029238 | 5.073653497 | 2.343024995 |

| | | | | | | | | | | | | | |
|---|---|---|---|---|---|---|---|---|---|---|---|---|---|
| Q9Y2Z0 | SUGT1 | | 0 | 0 | 0 | 11773000 | 56283000 | 17737000 | 0 | 28597666.67 | 0.109676364 | N/A | N/A |
| Q8NBJ7 | SUMF2 | | 0 | 89081000 | 114650000 | 180380000 | 124960000 | 92056000 | 67910333.33 | 132465333.3 | 0.209930145 | 1.950591712 | 0.963911832 |
| O15260 | SURF4 | | 0 | 49942000 | 26443000 | 13524000 | 0 | 0 | 25461666.67 | 4508000 | 0.237884828 | 0.177050468 | -2.497767437 |
| O60506 | SYNCRIP | | 426040000 | 408790000 | 434210000 | 885860000 | 678760000 | 530280000 | 423013333.3 | 698300000 | 0.05622674 | 1.650775389 | 0.723143835 |
| Q8N3V7 | SYNPO | | 273490000 | 221860000 | 317110000 | 287700000 | 165170000 | 166370000 | 270820000 | 206413333.3 | 0.25974437 | 0.762179061 | -0.39179812 |
| P37802 | TAGLN2 | | 539070000 | 798000000 | 458830000 | 795900000 | 966310000 | 1144500000 | 598633333.3 | 968903333.3 | 0.061348177 | 1.61852553 | 0.694680123 |
| O15533 | TAPBP | | 0 | 72812000 | 0 | 124360000 | 126900000 | 0 | 24270666.67 | 83753333.33 | 0.286496168 | 3.450804812 | 1.786932873 |
| Q13148 | TARDBP | | 87215000 | 173360000 | 199060000 | 174760000 | 222150000 | 326010000 | 153211666.7 | 240973333.3 | 0.192305616 | 1.57281321 | 0.653347344 |
| P26639 | TARS1 | | 93597000 | 0 | 128490000 | 95846000 | 121170000 | 118240000 | 74029000 | 111752000 | 0.390224265 | 1.509570574 | 0.594138206 |
| Q9NUY8 | TBC1D23 | | 0 | 0 | 0 | 40161000 | 49617000 | 0 | 0 | 29926000 | 0.120510402 | N/A | N/A |
| Q9Y4P3 | TBL2 | | 98948000 | 98878000 | 134640000 | 349930000 | 262130000 | 155690000 | 110822000 | 255916666.7 | 0.06483901 | 2.309258691 | 1.207429798 |
| Q969Z0 | TBRG4 | | 180110000 | 460760000 | 0 | 79517000 | 0 | 214940000 | 213623333.3 | 98152333.33 | 0.478904825 | 0.459464478 | -1.121974766 |
| Q13488 | TCIRG1 | | 96364000 | 158960000 | 0 | 0 | 72603000 | 90549000 | 85108000 | 54384000 | 0.59905766 | 0.638999859 | -0.646112482 |
| P17987 | TCP1 | | 9648200000 | 8982700000 | 8774700000 | 6486300000 | 5652500000 | 5702400000 | 9135200000 | 5947066667 | 0.001073791 | 0.651005634 | -0.619258066 |
| Q9Y2W6 | TDRKH | | 0 | 58733000 | 73360000 | 0 | 33066000 | 79820000 | 44031000 | 37628666.67 | 0.852219878 | 0.854594869 | -0.22668744 |
| Q9NZ01 | TECR | | 527450000 | 382110000 | 391310000 | 289380000 | 295220000 | 197320000 | 433623333.3 | 260640000 | 0.037965209 | 0.601074665 | -0.734383881 |
| Q9P273 | TENM3 | | 448240000 | 538700000 | 571760000 | 605240000 | 794130000 | 795630000 | 519566666.7 | 731666666.7 | 0.044236801 | 1.408224803 | 0.493877658 |
| P02786 | TFRC | | 3777200000 | 3560100000 | 3981000000 | 4704800000 | 4999900000 | 4745100000 | 3772766667 | 4816600000 | 0.002392957 | 1.276675826 | 0.352392242 |
| P61812 | TGFB2 | | 97914000 | 110160000 | 108810000 | 142240000 | 108730000 | 93988000 | 105628000 | 114986000 | 0.561355722 | 1.088593933 | 0.122465901 |
| P21980 | TGM2 | | 0 | 0 | 0 | 156540000 | 93276000 | 202590000 | 0 | 150802000 | 0.008910401 | N/A | N/A |
| Q8IYQ7 | THNSL1 | | 86750000 | 94090000 | 0 | 88150000 | 98687000 | 0 | 60280000 | 62279000 | 0.965545818 | 1.033161911 | 0.047066363 |
| P04216 | THY1 | | 857750000 | 782190000 | 709150000 | 1323400000 | 1110200000 | 851810000 | 783030000 | 1095136667 | 0.09436726 | 1.39858839 | 0.483971434 |
| P31483 | TIA1 | | 38572000 | 0 | 36844000 | 0 | 0 | 0 | 25138666.67 | 0 | 0.1163253 | 0 | N/A |
| Q9NQ88 | TIGAR | | 0 | 0 | 0 | 76006000 | 113690000 | 0 | 0 | 63232000 | 0.131570991 | N/A | N/A |
| P01033 | TIMP1 | | 0 | 0 | 0 | 278320000 | 329710000 | 269990000 | 0 | 292673333.3 | 9.67953E-05 | N/A | N/A |
| Q07157 | TJP1 | | 677220000 | 547790000 | 668250000 | 660430000 | 574220000 | 322340000 | 631086666.7 | 518996666.7 | 0.364559932 | 0.822385726 | -0.28211287 |
| Q9UDY2 | TJP2 | | 215900000 | 157550000 | 128910000 | 70425000 | 146480000 | 129350000 | 167453333.3 | 115418333.3 | 0.205274016 | 0.689256708 | -0.536886691 |
| P29401 | TKT | | 0 | 0 | 0 | 85386000 | 102850000 | 118600000 | 0 | 102278666.7 | 0.000438185 | N/A | N/A |
| Q9Y490 | TLN1 | | 5676900000 | 5131100000 | 4746400000 | 5865800000 | 5801800000 | 6260900000 | 5184800000 | 5976166667 | 0.060794016 | 1.152632053 | 0.204932044 |
| Q9Y4G6 | TLN2 | | 110730000 | 105560000 | 0 | 0 | 0 | 0 | 72096666.67 | 0 | 0.116343731 | 0 | N/A |
| Q99805 | TM9SF2 | | 185620000 | 0 | 0 | 232220000 | 222160000 | 324320000 | 61873333.33 | 259566666.7 | 0.047415917 | 4.195129835 | 2.068715461 |
| Q9HD45 | TM9SF3 | | 654430000 | 295900000 | 240130000 | 374110000 | 404210000 | 997680000 | 396820000 | 592000000 | 0.463397046 | 1.491860289 | 0.577112435 |
| Q92544 | TM9SF4 | | 528800000 | 430720000 | 561860000 | 318340000 | 332750000 | 199460000 | 507126666.7 | 283516666.7 | 0.017955276 | 0.559064796 | -0.838912592 |
| P49755 | TMED10 | | 952150000 | 1172000000 | 1252500000 | 1276800000 | 861010000 | 1049100000 | 1125550000 | 1062303333 | 0.695003474 | 0.943808212 | -0.08343437 |
| Q7Z7H5 | TMED4 | | 0 | 0 | 110950000 | 0 | 0 | 59226000 | 36983333.33 | 19742000 | 0.701958644 | 0.533808022 | -0.90560711 |
| Q9Y3B3 | TMED7 | | 103780000 | 89649000 | 121130000 | 97448000 | 98822000 | 0 | 104853000 | 65423333.33 | 0.310148076 | 0.623952899 | -0.680490968 |
| Q9BVK6 | TMED9 | | 621450000 | 760120000 | 828610000 | 699880000 | 721220000 | 466270000 | 736726666.7 | 629123333.3 | 0.350491934 | 0.853944023 | -0.227786593 |
| Q9NUM4 | TMEM106B | | 0 | 0 | 0 | 0 | 75492000 | 78015000 | 0 | 51169000 | 0.116223957 | N/A | N/A |
| Q9BVC6 | TMEM109 | | 1069600000 | 1178700000 | 1162800000 | 0 | 782150000 | 0 | 1137033333 | 260716666.7 | 0.029026631 | 0.229295535 | -2.124719832 |
| Q9H061 | TMEM126A | | 242430000 | 291050000 | 222270000 | 149700000 | 190860000 | 0 | 251916666.7 | 113520000 | 0.08753448 | 0.450625207 | -1.150000079 |
| Q9HC07 | TMEM165 | | 188770000 | 173370000 | 194050000 | 406300000 | 295460000 | 197940000 | 185396666.7 | 299900000 | 0.131391742 | 1.617612686 | 0.693866217 |
| Q6NUQ4 | TMEM214 | | 147130000 | 442350000 | 162350000 | 370530000 | 456970000 | 455840000 | 250610000 | 427780000 | 0.151605546 | 1.70695503 | 0.771425051 |
| Q6PI78 | TMEM65 | | 240770000 | 189710000 | 265790000 | 215590000 | 238880000 | 113420000 | 232090000 | 189296666.7 | 0.39127321 | 0.815617505 | -0.294035356 |
| P82094 | TMF1 | | 101320000 | 90043000 | 0 | 76451000 | 0 | 105520000 | 63787666.67 | 60657000 | 0.94778526 | 0.950920502 | -0.07260336 |
| Q9NYL9 | TMOD3 | | 394960000 | 325870000 | 311250000 | 190120000 | 208220000 | 212530000 | 344026666.7 | 203623333.3 | 0.00627018 | 0.591882412 | -0.756617507 |
| P62328 | TMSB4X | | 0 | 0 | 0 | 67230000 | 0 | 43782000 | 0 | 37004000 | 0.13355086 | N/A | N/A |
| Q71RG4 | TMUB2 | | 118500000 | 0 | 0 | 0 | 71818000 | 0 | 39500000 | 23939333.33 | 0.753128673 | 0.606059072 | -0.722469677 |
| Q9H3N1 | TMX1 | | 607000000 | 644090000 | 540020000 | 658840000 | 654030000 | 690560000 | 597036666.7 | 667810000 | 0.09526099 | 1.118541016 | 0.16161816 |
| Q9Y320 | TMX2 | | 98494000 | 134480000 | 195080000 | 0 | 196120000 | 0 | 142684666.7 | 65373333.33 | 0.338547838 | 0.458166493 | -1.126056141 |
| Q96JJ7 | TMX3 | | 59858000 | 0 | 135550000 | 175360000 | 0 | 121660000 | 65136000 | 99006666.67 | 0.629980722 | 1.519999181 | 0.604070547 |
| Q9C0C2 | TNKS1BP1 | | 0 | 0 | 0 | 202700000 | 138160000 | 146530000 | 0 | 162463333.3 | 0.00131277 | N/A | N/A |
| Q92973 | TNPO1 | | 0 | 25572000 | 0 | 81976000 | 47794000 | 144320000 | 8524000 | 91363333.33 | 0.048479647 | 10.71836384 | 3.422012789 |
| O60784 | TOM1 | | 92597000 | 43999000 | 82609000 | 109020000 | 0 | 84494000 | 73068333.33 | 64504666.67 | 0.824579945 | 0.882799206 | -0.179842762 |
| Q5JTV8 | TOR1AIP1 | | 80665000 | 62814000 | 105940000 | 83444000 | 113810000 | 143940000 | 83139666.67 | 113731333.3 | 0.227545276 | 1.367955128 | 0.452020907 |
| Q8NFQ8 | TOR1AIP2 | | 0 | 57990000 | 70576000 | 64044000 | 11215000 | 32737000 | 42855333.33 | 35998666.67 | 0.809305919 | 0.840004356 | -0.251531286 |
| Q13641 | TPBG | | 0 | 0 | 0 | 135640000 | 34582000 | 0 | 0 | 56740666.67 | 0.235668694 | N/A | N/A |
| P55327 | TPD52 | | 120270000 | 0 | 0 | 289500000 | 321820000 | 219050000 | 40090000 | 276790000 | 0.009253666 | 6.904215515 | 2.787477499 |

| | | | | | | | | | | | | | |
|---|---|---|---|---|---|---|---|---|---|---|---|---|---|
| O43399 | TPD52L2 | | 156260000 | 76821000 | 136040000 | 238100000 | 302040000 | 341400000 | 123040333.3 | 293846666.7 | 0.011258026 | 2.388214163 | 1.255932216 |
| P60174 | TPI1 | | 86945000 | 52065000 | 109420000 | 357750000 | 410620000 | 406750000 | 82810000 | 391706666.7 | 0.000204377 | 4.730185565 | 2.241896782 |
| P09493 | TPM1 | | 2574100000 | 3171200000 | 3027500000 | 2141200000 | 1982900000 | 1187600000 | 2924266667 | 1770566667 | 0.028877613 | 0.605473737 | -0.723863712 |
| P06753 | TPM3 | | 228940000 | 275430000 | 426010000 | 524170000 | 337420000 | 0 | 310126666.7 | 287196666.7 | 0.895884363 | 0.926062469 | -0.110818579 |
| P67936 | TPM4 | | 460160000 | 383810000 | 414080000 | 812400000 | 398180000 | 357420000 | 419350000 | 522666666.7 | 0.520987119 | 1.246373356 | 0.317736298 |
| O14773 | TPP1 | | 957080000 | 837650000 | 676700000 | 1138500000 | 1782400000 | 1223900000 | 823810000 | 1381600000 | 0.062445422 | 1.677085736 | 0.745956444 |
| P29144 | TPP2 | | 70746000 | 102000000 | 54210000 | 56955000 | 0 | 137140000 | 75652000 | 64698333.33 | 0.807893979 | 0.85520982 | -0.225649675 |
| P12270 | TPR | | 0 | 101340000 | 176930000 | 63989000 | 36062000 | 182100000 | 92756666.67 | 94050333.33 | 0.985742375 | 1.013946886 | 0.019982081 |
| Q14258 | TRIM25 | | 151990000 | 70242000 | 83910000 | 170610000 | 154250000 | 249900000 | 102047333.3 | 191586666.7 | 0.082661706 | 1.877429428 | 0.908758678 |
| Q13263 | TRIM28 | | 167890000 | 92212000 | 136030000 | 246220000 | 163130000 | 188310000 | 132044000 | 199220000 | 0.111177898 | 1.508739511 | 0.593343741 |
| O75962 | TRIO | | 0 | 0 | 37082000 | 48965000 | 43887000 | 0 | 12360666.67 | 30950666.67 | 0.402240753 | 2.503964187 | 1.324213929 |
| Q15642 | TRIP10 | | 57928000 | 33280000 | 0 | 0 | 23941000 | 0 | 30402666.67 | 7980333.333 | 0.294102686 | 0.26248794 | -1.929676957 |
| Q15643 | TRIP11 | | 1006900000 | 564510000 | 984430000 | 561200000 | 0 | 0 | 851946666.7 | 187066666.7 | 0.047953916 | 0.21957556 | -2.18721061 |
| Q9UI30 | TRMT112 | | 103010000 | 128330000 | 0 | 0 | 165910000 | 230820000 | 77113333.33 | 132243333.3 | 0.52440791 | 1.71492176 | 0.778142758 |
| Q8TD43 | TRPM4 | | 114630000 | 45052000 | 0 | 0 | 22968000 | 0 | 53227333.33 | 7656000 | 0.253656057 | 0.143835874 | -2.797504555 |
| Q99816 | TSG101 | | 0 | 0 | 60876000 | 61110000 | 0 | 0 | 20292000 | 20370000 | 0.997965392 | 1.003843879 | 0.005534915 |
| O75954 | TSPAN9 | | 0 | 0 | 56794000 | 0 | 104520000 | 0 | 18931333.33 | 34840000 | 0.708771471 | 1.840335247 | 0.8799686 |
| Q16762 | TST | | 296740000 | 314980000 | 0 | 0 | 0 | 0 | 203906666.7 | 0 | 0.116470038 | 0 | N/A |
| Q9C0H2 | TTYH3 | | 451800000 | 258070000 | 207880000 | 554880000 | 698840000 | 755020000 | 305916666.7 | 669580000 | 0.018844287 | 2.188766004 | 1.130117727 |
| Q71U36 | TUBA1A | | 136210000 | 0 | 233450000 | 107730000 | 188800000 | 0 | 123220000 | 98843333.33 | 0.793289509 | 0.802169561 | -0.318020872 |
| P68363 | TUBA1B | | 3245500000 | 3888100000 | 3711500000 | 4626100000 | 4969500000 | 7106300000 | 3615033333 | 5567300000 | 0.07099186 | 1.540041125 | 0.622968877 |
| P07437 | TUBB | | 2050900000 | 1687600000 | 1744200000 | 1702000000 | 2220800000 | 2544700000 | 1827566667 | 2155833333 | 0.291104346 | 1.179619531 | 0.238321614 |
| P68371 | TUBB4B | | 4203700000 | 3517400000 | 3991400000 | 4697800000 | 5385900000 | 5599600000 | 3904166667 | 5227766667 | 0.017539696 | 1.339022412 | 0.421180108 |
| Q3ZCM7 | TUBB8 | | 203120000 | 0 | 0 | 119160000 | 155330000 | 0 | 67706666.67 | 91496666.67 | 0.787089548 | 1.351368649 | 0.434421291 |
| Q6IBS0 | TWF2 | | 0 | 0 | 0 | 95513000 | 82825000 | 0 | 0 | 59446000 | 0.118125967 | N/A | N/A |
| P40222 | TXLNA | | 79789000 | 68455000 | 70235000 | 175520000 | 109490000 | 70687000 | 72826333.33 | 118565666.7 | 0.211741318 | 1.628060363 | 0.703154191 |
| P10599 | TXN | | 0 | 0 | 0 | 40710000 | 0 | 92972000 | 0 | 44560666.67 | 0.173051589 | N/A | N/A |
| O95881 | TXNDC12 | | 520410000 | 799740000 | 1019300000 | 856490000 | 1067700000 | 726710000 | 779816666.7 | 883633333.3 | 0.585461065 | 1.133129582 | 0.180312853 |
| Q9BRA2 | TXNDC17 | | 0 | 0 | 0 | 53467000 | 0 | 49816000 | 0 | 34427666.67 | 0.11661331 | N/A | N/A |
| Q8NBS9 | TXNDC5 | | 2452200000 | 2764200000 | 3072000000 | 3884800000 | 2834100000 | 3249100000 | 2762800000 | 3322666667 | 0.188980394 | 1.20264466 | 0.266210439 |
| P04818 | TYMS | | 135050000 | 156050000 | 87836000 | 0 | 0 | 55344000 | 126312000 | 18448000 | 0.016873647 | 0.146051048 | -2.775455383 |
| Q01081 | U2AF1 | | 122360000 | 231670000 | 47357000 | 91217000 | 42108000 | 147680000 | 133795666.7 | 93668333.33 | 0.550276351 | 0.700084955 | -0.514398091 |
| P26368 | U2AF2 | | 129380000 | 59152000 | 75477000 | 76916000 | 0 | 116110000 | 88003000 | 64342000 | 0.58745752 | 0.731134166 | -0.451791924 |
| P22314 | UBA1 | | 327890000 | 372210000 | 297840000 | 425620000 | 542820000 | 872640000 | 332646666.7 | 613693333.3 | 0.106825606 | 1.844880454 | 0.883527334 |
| Q14157 | UBAP2L | | 0 | 0 | 0 | 122100000 | 133280000 | 96212000 | 0 | 117197333.3 | 0.00043607 | N/A | N/A |
| P62837 | UBE2D2 | | 112090000 | 0 | 13266000 | 165220000 | 48950000 | 73502000 | 41785333.33 | 95890666.67 | 0.340270486 | 2.294840295 | 1.198393755 |
| P63279 | UBE2I | | 0 | 0 | 0 | 0 | 79269000 | 92418000 | 0 | 57229000 | 0.118444373 | N/A | N/A |
| P61088 | UBE2N | | 131460000 | 58420000 | 78974000 | 152650000 | 145580000 | 177940000 | 89618000 | 158723333.3 | 0.044291703 | 1.77110997 | 0.824653794 |
| Q92575 | UBXN4 | | 142630000 | 155940000 | 188770000 | 123380000 | 162820000 | 159020000 | 162446666.7 | 148406666.7 | 0.492246865 | 0.913571634 | -0.130410239 |
| Q9Y3C8 | UFC1 | | 0 | 0 | 0 | 55712000 | 76232000 | 0 | 0 | 43981333.33 | 0.125647043 | N/A | N/A |
| O94874 | UFL1 | | 313090000 | 223390000 | 433150000 | 382520000 | 423460000 | 466210000 | 323210000 | 424063333.3 | 0.197860948 | 1.31203655 | 0.39180791 |
| P61960 | UFM1 | | 0 | 69905000 | 0 | 0 | 0 | 165290000 | 23301666.67 | 55096666.67 | 0.623219096 | 2.364494671 | 1.241531891 |
| Q9NYU2 | UGGT1 | | 500970000 | 461820000 | 381960000 | 310860000 | 555600000 | 658020000 | 448250000 | 508160000 | 0.611109738 | 1.133653095 | 0.180979234 |
| P11172 | UMPS | | 0 | 0 | 0 | 0 | 22072000 | 29732000 | 0 | 17268000 | 0.124739812 | N/A | N/A |
| Q92900 | UPF1 | | 0 | 0 | 0 | 0 | 76754000 | 29483000 | 0 | 35412333.33 | 0.188338346 | N/A | N/A |
| O60763 | USO1 | | 47186000 | 48077000 | 58312000 | 135640000 | 74637000 | 35874000 | 51191666.67 | 82050333.33 | 0.351022596 | 1.602806446 | 0.680600217 |
| Q14694 | USP10 | | 0 | 92070000 | 90828000 | 177570000 | 147600000 | 119110000 | 60966000 | 148093333.3 | 0.066738109 | 2.429113495 | 1.280429898 |
| P45974 | USP5 | | 325060000 | 297000000 | 302870000 | 306610000 | 639020000 | 925580000 | 308310000 | 623736666.7 | 0.152916343 | 2.023082828 | 1.016555387 |
| P46939 | UTRN | | 916470000 | 954260000 | 769300000 | 55044000 | 54291000 | 95043000 | 880010000 | 68126000 | 0.000151128 | 0.077415029 | -3.691242511 |
| Q8NBZ7 | UXS1 | | 0 | 0 | 18253000 | 16499000 | 91159000 | 189760000 | 6084333.333 | 99139333.33 | 0.139417247 | 16.29419821 | 4.026286458 |
| Q15836 | VAMP3 | | 231110000 | 292900000 | 239280000 | 449390000 | 331790000 | 299220000 | 254430000 | 360133333.3 | 0.099861692 | 1.415451532 | 0.501262349 |
| Q9P0L0 | VAPA | | 1631200000 | 1901100000 | 1813200000 | 1132900000 | 1316000000 | 1592400000 | 1781833333 | 1347100000 | 0.048947473 | 0.756019081 | -0.403505447 |
| O95292 | VAPB | | 1049300000 | 1209200000 | 984760000 | 916690000 | 1028000000 | 803030000 | 1081086667 | 915906666.7 | 0.150705727 | 0.847209289 | -0.239209687 |
| P26640 | VARS1 | | 0 | 246140000 | 185280000 | 453290000 | 610710000 | 451700000 | 143806666.7 | 505233333.3 | 0.016446414 | 3.513281721 | 1.812819267 |
| P50552 | VASP | | 135560000 | 222860000 | 220490000 | 142600000 | 128930000 | 51450000 | 192970000 | 107660000 | 0.102133893 | 0.557910556 | -0.841894246 |
| Q99536 | VAT1 | | 373960000 | 259220000 | 471850000 | 678380000 | 730340000 | 588310000 | 368343333.3 | 665676666.7 | 0.015991652 | 1.807217904 | 0.853770468 |
| P18206 | VCL | | 302230000 | 183270000 | 337190000 | 269050000 | 301330000 | 284810000 | 274230000 | 285063333.3 | 0.830808327 | 1.039504552 | 0.055896076 |

| | | | | | | | | | | | | | |
|---|---|---|---|---|---|---|---|---|---|---|---|---|---|
| P55072 | VCP | | 4868700000 | 4425200000 | 4439500000 | 4700900000 | 5400000000 | 5423700000 | 4577800000 | 5174866667 | 0.098380448 | 1.130426551 | 0.176867257 |
| O15240 | VGF | | 0 | 0 | 0 | 490560000 | 482370000 | 758350000 | 0 | 577093333.3 | 0.003122747 | N/A | N/A |
| P08670 | VIM | | 22944000000 | 22719000000 | 24194000000 | 31044000000 | 32387000000 | 25610000000 | 23285666667 | 29680333333 | 0.039411531 | 1.274618149 | 0.350065108 |
| Q96RL7 | VPS13A | | 115810000 | 100520000 | 0 | 0 | 0 | 0 | 72110000 | 0 | 0.118099738 | 0 | N/A |
| Q9P253 | VPS18 | | 0 | 175390000 | 165710000 | 67002000 | 127260000 | 131000000 | 113700000 | 108420666.7 | 0.934741904 | 0.953567869 | -0.068592471 |
| Q96QK1 | VPS35 | | 243980000 | 340250000 | 298530000 | 237610000 | 181250000 | 283410000 | 294253333.3 | 234090000 | 0.21266134 | 0.795538991 | -0.329995453 |
| P49754 | VPS41 | | 0 | 0 | 0 | 0 | 74001000 | 64823000 | 0 | 46274666.67 | 0.117852158 | N/A | N/A |
| Q9NRW7 | VPS45 | | 14520000 | 27025000 | 14233000 | 0 | 52667000 | 27816000 | 18592666.67 | 26827666.67 | 0.629443174 | 1.442916562 | 0.528987877 |
| Q86Y07 | VRK2 | | 0 | 116800000 | 0 | 29394000 | 0 | 0 | 38933333.33 | 9798000 | 0.508200415 | 0.251660959 | -1.990446672 |
| Q9NP79 | VTA1 | | 397730000 | 293850000 | 259460000 | 0 | 0 | 0 | 317013333.3 | 0 | 0.00158638 | 0 | N/A |
| Q96AJ9 | VTI1A | | 224100000 | 0 | 98417000 | 0 | 0 | 120950000 | 107505666.7 | 40316666.67 | 0.428614077 | 0.375018991 | -1.414964438 |
| Q9UEU0 | VTI1B | | 216030000 | 164360000 | 176840000 | 193760000 | 246600000 | 189800000 | 185743333.3 | 210053333.3 | 0.368957393 | 1.130879529 | 0.177445249 |
| A3KMH1 | VWA8 | | 0 | 0 | 87762000 | 0 | 56984000 | 0 | 29254000 | 18994666.67 | 0.783286804 | 0.64930152 | -0.623039508 |
| P23381 | WARS1 | | 0 | 0 | 0 | 51587000 | 67158000 | 56238000 | 0 | 58327666.67 | 0.000225607 | N/A | N/A |
| Q9Y6W5 | WASF2 | | 512780000 | 374170000 | 291830000 | 200300000 | 280730000 | 182750000 | 392926666.7 | 221260000 | 0.073406834 | 0.56310762 | -0.828517422 |
| Q641Q2 | WASHC2A | | 73833000 | 0 | 67642000 | 50337000 | 60557000 | 0 | 47158333.33 | 36964666.67 | 0.752347551 | 0.783841668 | -0.351365828 |
| Q9BZH6 | WDR11 | | 0 | 0 | 0 | 95593000 | 33567000 | 0 | 0 | 43053333.33 | 0.198960647 | N/A | N/A |
| Q8NI36 | WDR36 | | 0 | 0 | 0 | 0 | 32804000 | 52201000 | 0 | 28335000 | 0.136401 | N/A | N/A |
| O14980 | XPO1 | | 0 | 0 | 8602600 | 0 | 0 | 54259000 | 2867533.333 | 18086333.33 | 0.452662332 | 6.307279195 | 2.657017796 |
| Q9UBH6 | XPR1 | | 0 | 2774100 | 0 | 5423600 | 0 | 0 | 924700 | 1807866.667 | 0.686054929 | 1.955084532 | 0.967230987 |
| P13010 | XRCC5 | | 1107200000 | 976650000 | 853680000 | 827790000 | 873370000 | 799170000 | 979176666.7 | 833443333.3 | 0.128811752 | 0.851167477 | -0.232485067 |
| P12956 | XRCC6 | | 1251700000 | 1159700000 | 1210700000 | 1516000000 | 1390300000 | 801880000 | 1207366667 | 1236060000 | 0.903271098 | 1.023765219 | 0.033884899 |
| P54577 | YARS1 | | 164030000 | 269250000 | 81256000 | 109270000 | 189380000 | 54109000 | 171512000 | 117586333.3 | 0.466583596 | 0.685586626 | -0.544589129 |
| P67809 | YBX1 | | 28170000 | 47918000 | 104310000 | 279540000 | 156500000 | 79277000 | 60132666.67 | 171772333.3 | 0.149177278 | 2.856556059 | 1.514276843 |
| P16989 | YBX3 | | 526270000 | 495020000 | 461390000 | 1218900000 | 1060200000 | 527840000 | 494226666.7 | 935646666.7 | 0.103214572 | 1.893152939 | 0.920790965 |
| P07947 | YES1 | | 348280000 | 362050000 | 707950000 | 537280000 | 570110000 | 496800000 | 472760000 | 534730000 | 0.631579849 | 1.13108131 | 0.177702644 |
| Q9BWQ6 | YIPF2 | | 28799000 | 85666000 | 23627000 | 34567000 | 268440000 | 146330000 | 46030666.67 | 149779000 | 0.214560556 | 3.253895953 | 1.70216812 |
| Q969M3 | YIPF5 | | 216600000 | 0 | 0 | 0 | 0 | 235870000 | 72200000 | 78623333.33 | 0.954903041 | 1.088965836 | 0.122958693 |
| Q96EC8 | YIPF6 | | 55705000 | 43036000 | 0 | 76090000 | 79427000 | 0 | 32913666.67 | 51839000 | 0.573744611 | 1.57499924 | 0.655351133 |
| O15498 | YKT6 | | 0 | 0 | 0 | 109930000 | 100110000 | 0 | 0 | 70013333.33 | 0.116985223 | N/A | N/A |
| Q96TA2 | YME1L1 | | 406590000 | 365500000 | 367060000 | 227960000 | 196930000 | 379130000 | 379716666.7 | 268006666.7 | 0.125727592 | 0.705806961 | -0.502654435 |
| P31946 | YWHAB | | 137340000 | 74651000 | 124830000 | 234800000 | 335560000 | 218780000 | 112273666.7 | 263046666.7 | 0.021699908 | 2.342906173 | 1.228299179 |
| P62258 | YWHAE | | 88504000 | 57106000 | 0 | 143580000 | 164710000 | 98117000 | 48536666.67 | 135469000 | 0.05558807 | 2.791065174 | 1.480815813 |
| P27348 | YWHAQ | | 112660000 | 0 | 104720000 | 88800000 | 172350000 | 192830000 | 72460000 | 151326666.7 | 0.177641829 | 2.088416598 | 1.06240953 |
| P63104 | YWHAZ | | 444210000 | 698290000 | 722750000 | 1505900000 | 1333000000 | 1006500000 | 621750000 | 1281800000 | 0.018278557 | 2.061600322 | 1.043764667 |
| Q8N4Q0 | ZADH2 | | 257070000 | 0 | 281600000 | 192260000 | 275970000 | 318220000 | 179556666.7 | 262150000 | 0.444064851 | 1.459984777 | 0.545953327 |
| Q8WU90 | ZC3H15 | | 0 | 0 | 147900000 | 183870000 | 0 | 201380000 | 49300000 | 128416666.7 | 0.384579308 | 2.604800541 | 1.381172904 |
| Q7Z2W4 | ZC3HAV1 | | 150770000 | 138140000 | 178550000 | 171620000 | 200130000 | 253220000 | 155820000 | 208323333.3 | 0.120898125 | 1.336948616 | 0.418944018 |
| O95159 | ZFPL1 | | 163180000 | 244730000 | 223040000 | 391160000 | 259780000 | 205350000 | 210316666.7 | 285430000 | 0.280880155 | 1.357143989 | 0.440573795 |
| Q68DK2 | ZFYVE26 | | 0 | 0 | 0 | 35195000 | 33449000 | 0 | 0 | 22881333.33 | 0.116373791 | N/A | N/A |
| O75844 | ZMPSTE24 | | 154560000 | 31371000 | 177180000 | 127480000 | 249600000 | 187290000 | 121037000 | 188123333.3 | 0.307457988 | 1.554263022 | 0.636230666 |
| O15231 | ZNF185 | | 132430000 | 161030000 | 158680000 | 293840000 | 203150000 | 146130000 | 150713333.3 | 214373333.3 | 0.221270181 | 1.422391295 | 0.5083184 |
| O43264 | ZW10 | | 352650000 | 378830000 | 279030000 | 205850000 | 241230000 | 296500000 | 336836666.7 | 247860000 | 0.089352659 | 0.735846256 | -0.442523727 |
| Q15942 | ZYX | | 67846000 | 0 | 0 | 0 | 94707000 | 0 | 22615333.33 | 31569000 | 0.828965447 | 1.395911329 | 0.481207301 |
| P01892 | HLA-A | | 305760000 | 332900000 | 441930000 | 573710000 | 467090000 | 356700000 | 360196666.7 | 465833333.3 | 0.232834922 | 1.29327497 | 0.371029047 |
| P10319 | HLA-B1 | | 0 | 0 | 6795200 | 0 | 8962800 | 0 | 2265066.667 | 2987600 | 0.856568949 | 1.318989875 | 0.39943349 |
| P30481 | HLA-B2 | | 117730000 | 125370000 | 0 | 221900000 | 237040000 | 124460000 | 81033333.33 | 194466666.7 | 0.102518953 | 2.399835459 | 1.262935493 |
